# InFiniteLungDT, an in-vitro-learned digital twin for the prediction of material-triggered chronic neutrophilic lung inflammation: mechanism-anchored pre-validation across materials, species and exposure scenarios

**DOI:** 10.64898/2026.05.12.723437

**Authors:** Iztok Urbančič, Tilen Koklič, Hana Kokot, Boštjan Kokot, Tomasz Kolodziej, Lan Ma-Hock, Pernille Høgh Danielsen, Keld Alstrup Jensen, Markus Rehberg, Mateja Čubej Gašparin, Tereza Pahor, Fréderic Cosnier, Sarah Valentino, Carole Seidel, Christina Isaxon, Tomaž Vuk, Laurent Gaté, Andrea Haase, Robert Landsiedel, Tobias Stoeger, Ulla Birgitte Vogel, Janez Štrancar

## Abstract

Regulatory agencies require the dose–response relationship, time of onset and duration of chronic lung inflammation, which no animal-free method has yet delivered. We present the pre-validation of an in-vitro-learned digital twin (InFiniteLungDT) that predicts neutrophilic lung inflammation — adverse outcome pathway (AOP) key event KE1497, increased recruitment of inflammatory cells — as a dose- and time-resolved trajectory from acute to chronic conditions, for respirable materials and their mixtures and without knowledge of their physicochemical properties. Seventeen biological observables, nine of them corresponding to Key Events documented in AOP-Wiki, are monitored hourly for 24 hours by bi-modal live-cell confocal microscopy and translated into material-specific interaction couplings that parameterise a system of coupled first-order differential equations — the early Mode-of-Action (eMoA). Forward integration of the eMoA, with material-agnostic terms for the systemic response, yields the prediction.

Calibration used 15 materials chosen for diversity of eMoA against mouse intratracheal instillation. Pre-validation used 49 materials with matched mouse instillation data — including five industrial samples whose intrinsic properties cannot be characterised — and seven materials with rat inhalation data acquired in compliance with OECD Test Guideline (TG) 412, in total 3,094 mouse and 364 rat exposures. Predictive capacity for LOAEL/BMC is 92% for instillation (R² = 0.73) and 84% for inhalation (R² = 0.63), against the 0.55–0.73 inter-study variability range for benchmark concentrations in the US EPA ToxRefDB; for the influx trajectory it is 83% and 77%.

Because the twin delivers the full eMoA and not only the endpoint, materials can be grouped by a measured eMoA distance against a threshold derived from 22 repeat measurements of one reference material. Fewer than 3% of the 1,176 material pairs fall below it, and grouping cuts across chemistry in both directions: nanoforms of one substance may separate while chemically unrelated materials coincide. Because the criterion lies upstream of the predicted endpoint, a grouped material represented by its reference avoids the complete Test Guideline study, not only its neutrophil measurement. Three grouping scenaria, corresponding for different objectives, are illustrated.

The method is positioned as a replacement for the in vivo neutrophil-influx endpoint and, through grouping, as a reduction of animal use across inhalation test guidelines — in range-finding, Point-of-Departure derivation, grouping and read-across, and as a component of an Integrated Approach to Testing and Assessment (IATA). With a time-to-result under one week, it is also a suitable tool within the Safe-and-Sustainable-by-Design (SSbD) framework during development of new advanced materials.

## 1 Introduction

### 1.1 Comprehensive hazard information is not available for all chemicals in commerce

Despite extensive regulatory requirements for chemical safety assessment, comprehensive toxicological information is not available for all chemicals in commerce. Under REACH, information requirements increase with production or import volume. A sub-acute repeated-dose toxicity study (28 days) forms part of the standard information requirements for substances manufactured or imported at ≥10 t/year, whereas a sub-chronic (90-day) repeated-dose toxicity study is a standard information requirement at ≥100 t/year. However, these requirements do not necessarily entail testing by the inhalation route; the appropriate route is selected considering the likely route of human exposure, and the standard testing requirements may be adapted where scientifically justified. Consequently, inhalation-specific repeated-dose data may remain unavailable even for substances subject to repeated-dose toxicity information requirements. Data gaps are expected to be even greater for substances manufactured or imported at lower tonnages and for substances at an experimental or early research stage. For nanoforms and other poorly soluble particles, for which the inhalation route may be particularly relevant, the availability of route-specific repeated-dose data can therefore be especially limited.

Generating such information with current methods is demanding. Repeated-dose inhalation studies are low-throughput, costly and animal-intensive: for example, TG 413 studies with BAL and lung-burden groups require about 160 animals per substance, plus range-finding studies. Each additional substance, form or exposure scenario multiplies this effort. EU legislation also requires animal testing to be used only as a last resort. Several regulatory and scientific initiatives therefore promote new approach methodologies (NAMs), defined as any in vitro, in chemico or in silico method, or their combination, that enables next-generation risk assessment ^1^.

### 1.2 For regulatory purposes, a NAM must deliver a dose-response relationship, not only a hazard flag

A NAM-based approach for systemic toxicity was first proposed in 2007 by the US National Academy of Sciences. The original approach for classification of carcinogens by IARC was that a NAM capable of demonstrating that an adverse outcome pathway is triggered at *any* dose — even an irrelevantly high one — could be useful for regulatory purposes^1^. However, this identifies a trigger as potentially harmful without determining the dose at which harm occurs. So far, only a few NAMs have been accepted by regulatory bodies — reconstructed human epidermis methods for skin corrosion and irritation (OECD TG 431, 439) and the human Cell Line Activation Test for skin sensitisation (OECD TG 442C/D/E)^1^ — They all have been validated in a formal validation study, thus accepted and endorsed as OECD TGs. However all of them serve acute risk assessment, i.e. observing/predicting effects over timespan of a few days. **NAMs for chronic adverse outcomes (i.e. beyond 3–12 months), which carry the highest economic burden, remain absent.**

An analysis by a large group of toxicologists from multiple regulatory agencies (BfR, OECD, US EPA, NIEHS, EFSA, RIVM, NICEATM and others) identified several obstacles to the broader application of NAMs in regulatory risk assessment^2^:

A. unreliable modelling of repeated-dose toxicity, including chronic toxicity, and of different exposure scenarios;
B. lack of detailed evaluation of accuracy, within laboratory repeatability and between laboratory reproducibility; and
C. inability to deliver a quantitative dose–response relationship (quantitative hazard characterisation) and a reliable quantitative Point of Departure (PoD), with currently no recognised alternative to the long-term rodent bioassay that would simultaneously measure the apical endpoint of interest and provide dose–response information^3^.

Toxicology of particulates pose additional challenges, related to assay artefacts specific to particulates, resulting in absence of predictive, validated alternative methods that could replace animal tests for particulates^4^;

A further obstacle is the inability of current NAMs to address **mixture risk assessment** (MRA), the assessment of adverse effects arising from combined exposure to multiple substances, which regulatory authorities such as EFSA and the US EPA perform routinely^5,6^.

### 1.3 Digital twins address the static perspective of existing NAMs

One common property to practically all methods proposed as NAMs is their **static perspective**: they examine phenomena and correlations within a particular time frame, usually early responses of up to a month, when data are easier — or only then possible — to obtain. It is therefore unsurprising that there are no NAMs for risk characterization.

A methodology capable of predicting the dynamics of a complex system ahead of time is the **digital twin**^7–11^ — a virtual replica of a physical/biological/chemical system, calibrated with measured data, able to simulate the time evolution of that system into the future under various scenarios. Digital twins employ mechanistic modelling, typically a set of differential equations describing the time evolution of observables through interaction terms whose coupling constants require numerical parameterisation. As noted by De Domenico et al. (2025) biological digital twins should be based on an understanding of the underlying biological, physiological or physical mechanisms that govern the behaviour of systems in the body, reflected in the associated mathematical representation^11^. **This differs fundamentally from data-driven machine learning models, which learn patterns from data without necessarily incorporating underlying mechanisms**^11^ — a distinction that becomes decisive when predictions must be made outside the range of the training data.

The integration of mechanistic (open-box) models into digital twins also addresses the **overfitting** problem that is prevalent in systems characterised by vast parameter spaces, such as deep learning methods^10^. Namely, for the digital twin presented here, the in vitro degrees of freedom do not exceed the number of available in vivo data points (for details see Design objectives and premises in chapter 4.1). Digital twins additionally enable **scenario-based modelling**: once the Mode-of-Action (MoA) triggered by a stressor is known, different exposure patterns — single, repeated, semi-continuous — follow the same propagation and need not each be tested in life. Testing all such scenarios experimentally would be prohibitively long and expensive; multi-scenario testing of simpler digital twins is already routine in epidemic modelling, which has been used for policy decision-making.

Digital twins have nevertheless not been broadly employed in biology, toxicology or pharmacology, because of three **grand challenges** relating to the data needed for parameterisation: (1) hypothesis-based data acquisition with major knowledge gaps in the underlying biology; (2) incompleteness of data on biological observables across time points and concentrations; and (3) low data quality due to biological noise and phase delays between events at different levels of complexity^8,11^ – issues all addressed in this work.

### 1.4 What this work demonstrates

Independently and well before the work of De Domenico et al. (2025)^12^, our group developed a similar mathematical representation of a digital twin of interdependent biological quantities *q_i_*^7^. Each interaction term is — as in generalised chemical kinetics — expressed as a product of one or more state functions (analogous to concentrations or activities) and a state-independent, material-specific interaction coupling (analogous to a reaction rate constant). The general mathematical formula is given in section 3 and shown as the inset of Figure 1a.

**Figure 1.**
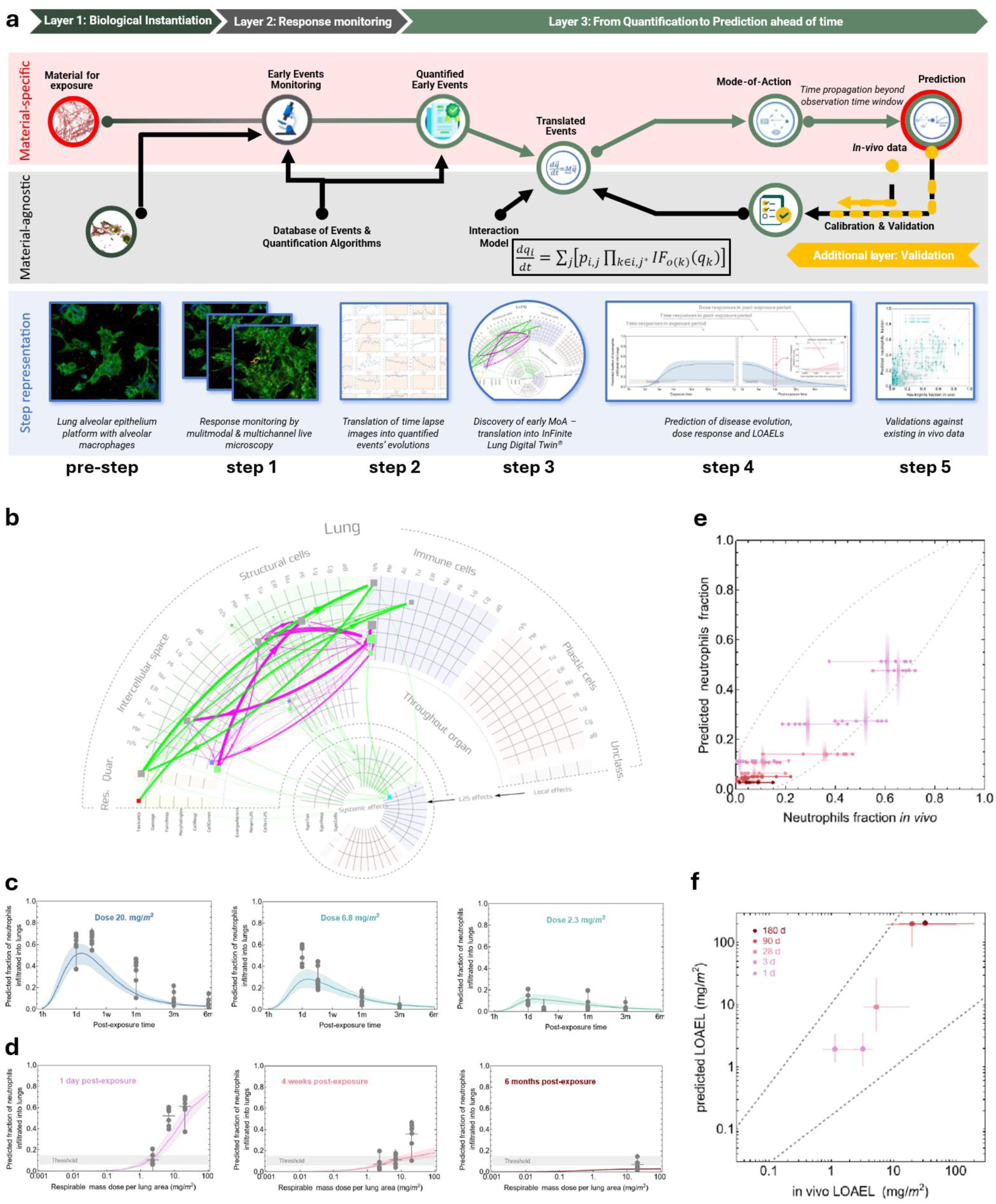
**Basic principles behind the construction, and evaluation of the output of the in vitro-learned digital twin** for chronic neutrophilic lung inflammation (InFiniteLungDT), capable of predicting the dose (LOAEL/BMC) at which inflammatory cell influx (KE1497) exceeds the pre-defined threshold, as well as the resolution time of inflammation at any dose. **a** the pipeline comprises three layers — (1) biological instantiation, (2) response monitoring, (3) quantification and prediction ahead of time — subdivided into steps that are either material-specific or material-agnostic. It starts with preparation of the biological platform (a co-culture of lung epithelial cells and lung macrophages, as well as the respective monocultures) and preparation of the material (pre-step); followed by 24 hours of time-lapse monitoring of single post-exposure responses in vitro by live-cell bi-modal confocal microscopy, in which 4 physical channels are used to acquire the evolution of the responses of the biological system (step 1); these are quantified into 17 (sub)cellular observables in a material-agnostic way, using a database of events and quantification algorithms (step 2); which are in turn translated into material-specific interaction couplings that parameterise the predefined material-agnostic interaction model and define the material-specific-triggered early Mode-of-Action (eMoA) (step 3). The mathematical representation of the digital twin is shown as an inset in the black frame, where we have postulated that an interaction term M_i,j_ can be expressed as a product of one or more state functions IF(q_k_) and the state-independent interaction coupling p_i,j_. By applying the discovered MoA (**b**) to a pre-selected exposure scenario (digital twin), we finally predict (**c**) the evolution of the in vivo responses/endpoints by time propagation (step 4, solid lines), which we calibrate — to define the best use of in vitro data and achieve the highest accuracy and reproducibility — and validate against in vivo data (step 5, gray datapoints); from time-dependent values at different doses, (**d**) dose-responses per time window are determined, which define (**f**) LOAEL/BMC values for comparison with in vivo-obtained LOAEL/BMC. **b – f** the case of intratracheal mice instillation of anatase TiO₂ nanotubes.

This work demonstrates that it is possible to:

- construct a digital twin providing long-term hazard assessment up to 6–9 months (internal numerical stability tests conducted up to 2 years; data not shown) beyond the time scale of in vitro monitoring;
- parameterise such a digital twin from 24-hour multi-channel time-lapse in vitro live cell (sub-confluent) co-culture clusters imaging, with automated early Mode-of-Action (eMoA)discovery;
- pre-validate it against a large but finite set of in vivo data, across two species and two exposure modalities;
- achieve more than 80% accuracy; and
- achieve better repeatability than the in vivo experiments themselves.

We overcome the three grand challenges as follows. (1) **Hypothesis-free but biologically relevant data acquisition** — time-lapse imaging with at least 20 time points per day instead of static data collection, which distinguishes driving from driven events, goes beyond pattern matching, reveals causality, and enables parameterisation of the full set of differential equations. (2) **Completeness of acquired data** — high-throughput live bi-modal multi-channel fluorescence and back-scatter confocal microscopy monitoring the complete evolution of 17 observables covering different functional units of cells, each proposed as, or mechanistically linked to, a Key Event (KE) in AOP-Wiki. (3) **High-quality data** — an optimised number of technical and biological replicates, noise-reducing data pre-processing algorithms, and predefined phenomenological interaction functions, all implemented to increase the significance of interaction couplings and the reliability of the resulting predictions.

## 2 Aim, context of use, and applicability domain

### Aim

InFiniteLungDT quantitatively predicts neutrophil accumulation in the lung — a systemic inflammatory response to local injury — for a delivered dose of the respirable fraction of a poorly water-soluble material, for single (instillation) or continuous/repeated (inhalation) exposure scenarios, at different doses or dose rates, for any time point during or post-exposure (shown to match in vivo data up to 9 months (internal numerical stability tests conducted up to 2 years) from the first interaction of cells with the trigger).

### Input

Predictions of the digital twin (DT) are generated on the basis of 24-hour in vitro monitoring of 17 biological observables at the level of cellular organelles, whole cells, and intercellular events, in a lung cell line co-culture model exposed to the material (Table 2, Figure 1, and Figure 2). Of note, no knowledge of material’s physico-chemical properties, such as specific surface area, functionalization or solubility is required, since material properties are reflected in the dynamics of the biological observables.

**Figure 2.**
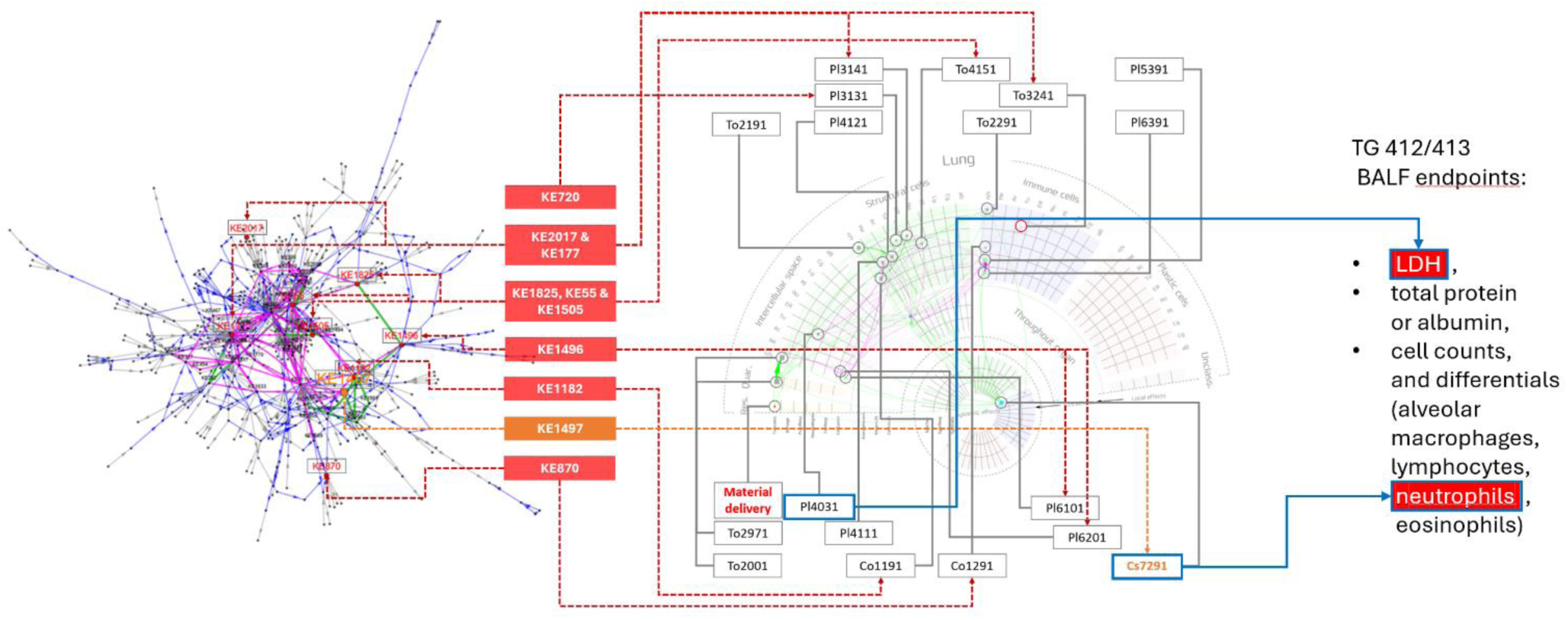
**Association map of in vitro observables monitored during in vitro-learned InFiniteLungDT** construction (middle panel) and KEs related to inflammation in AOP-Wiki (blue and grey dots on the network on the left), which are related through the 9 selected KEs (red central column), culminating in the endpoint predicted by the digital twin — KE1497 (orange). Left panel: 250 KEs related to inflammation (from AOP-Wiki) (blue), with an additional 281 upstream or downstream KEs (grey). Green lines show linear pathways connecting KEs leading to KE1497; pink cycles show cycles with KEs including KE1497 (altogether 46 KEs). Middle panel: KEs present in AOP-Wiki that are also measured in vitro by InFiniteLungDT: KE720 – disruption of microtubule dynamics; KE2017 and KE177 – ER stress and mitochondrial dysfunction; KE1825, KE55 and KE1505 – disrupted cell cycle, increased cell death and injury; KE1496 – secretion of proinflammatory mediators; KE1182 – affected epithelial cell proliferation; KE1497 – PMN influx, a robust and consistent marker of induced inflammation in vivo; KE870 – affected immune cell proliferation. For codes in the right panel, see Table 1. Rightmost panel: BALF-related OECD Test Guideline TG 412 (Subacute Inhalation Toxicity: 28-day Study) and TG 413 (Subchronic Inhalation Toxicity: 90-day Study) endpoints corresponding to the two endpoints predicted by the digital twin (red background, blue frame).

**Table 1.**
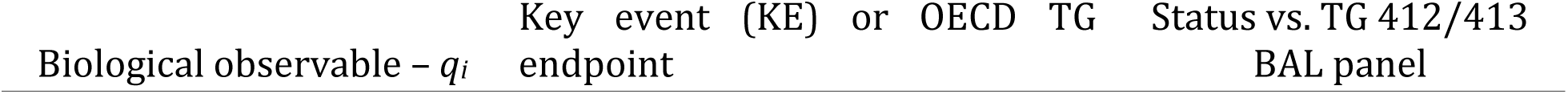
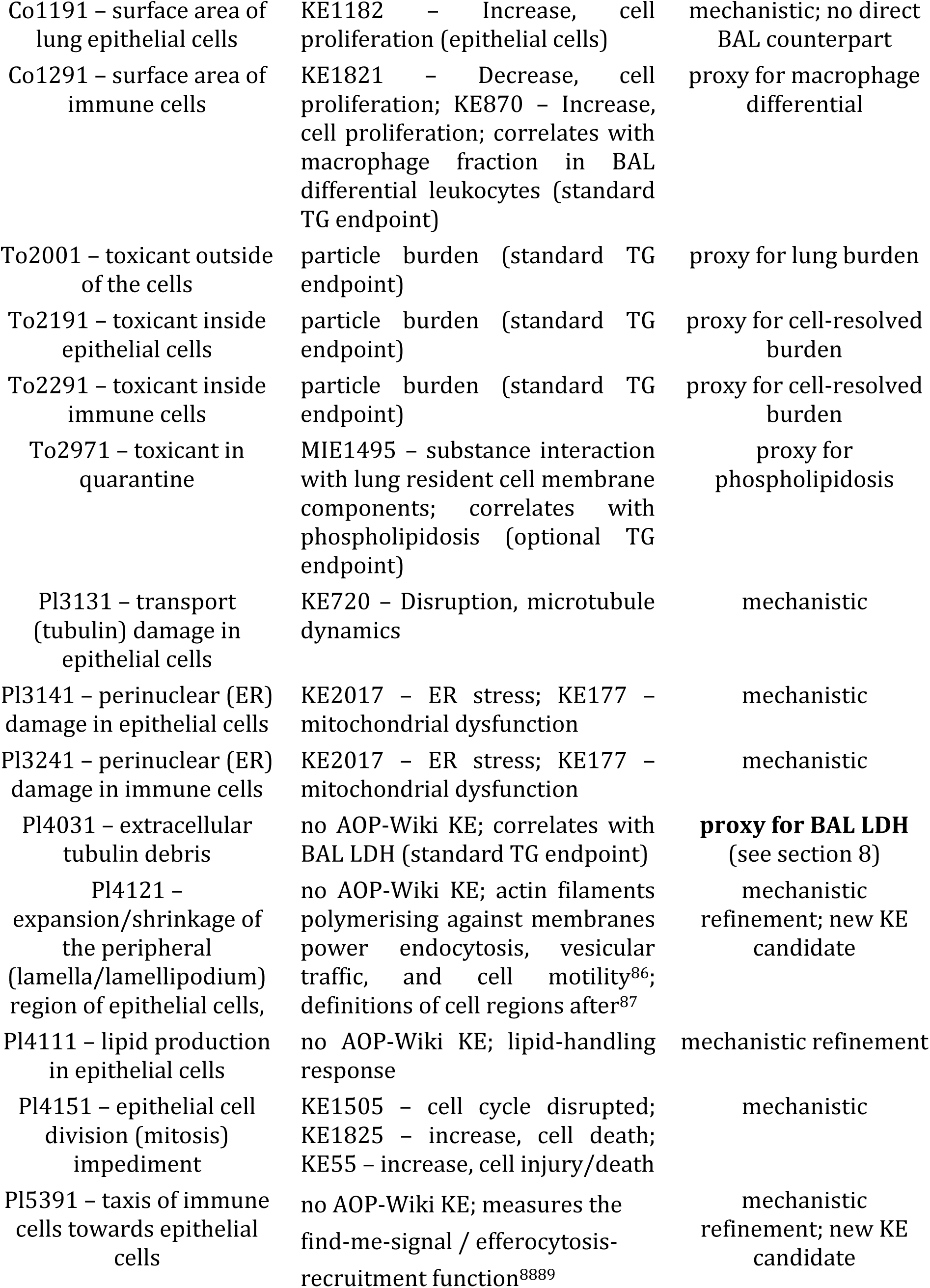
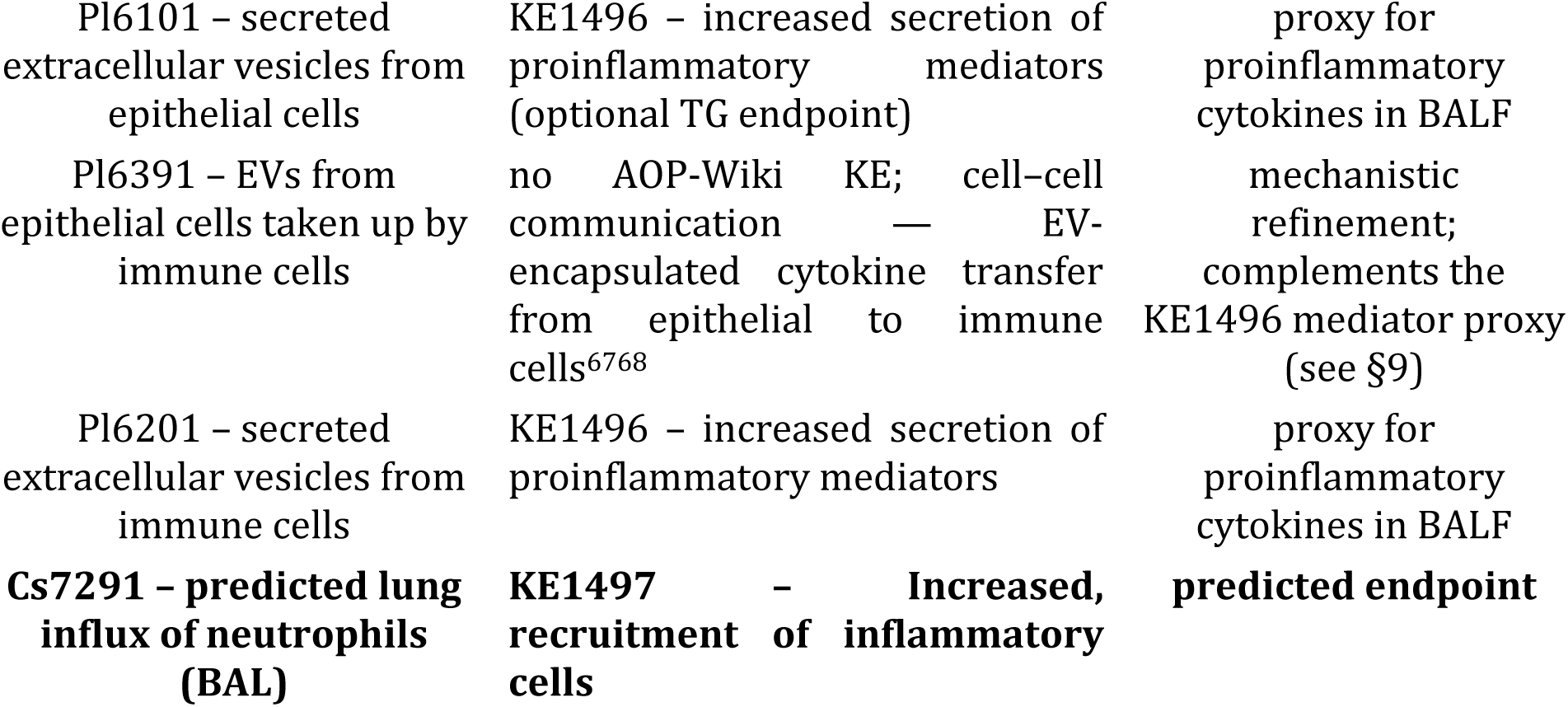
Biological observables q_i_ measured in vitro and used in the digital twin, with associated Key Events documented in AOP-Wiki or analogues identified in the standard endpoint list of OECD TGs, and their status relative to the TG 412/413 BAL panel.

**Table 2.** Species parameters used to translate between airborne concentration and delivered respirable dose per unit lung surface. The bottom row gives the resulting lung dose rate per unit airborne concentration, relative to the mouse under the same ambient conditions.

|  | Mouse | Rat | Human |
|---|---|---|---|
| Lung surface area (m<sup>2</sup>) | 0.008 | 0.3 | 50 |
| Air flow (m<sup>3</sup>/h) | 0.0015 | 0.02 | 1.5 (light work) |
| Dose rate relative to mouse under same ambient conditions | 1 | 0.4 | 0.2 |

### Predicted endpoint

KE1497 — *Increased, recruitment of inflammatory cells* — expressed as the neutrophil fraction in bronchoalveolar lavage (BAL), together with the derived dose– response curves, benchmark concentration (LOAEL/BMC), and inflammation resolution time is an OECD-endorsed key event ^13^. KE1497 is the central inflammation readout of the BAL panel prescribed by OECD TG 412 and TG 413, as it is a robust and consistent marker of particle-induced inflammation in vivo.

### Pre-validation status

The method’s accuracy figures described in sections 7.1–7.3 apply in every context below.

### Regulatory contexts of use

- range-finding and post-exposure-observation timing for OECD TG 403/412/413/452 (sections 7.1, 7.2);
- derivation of a dose–response relationship and Point of Departure for neutrophilic lung inflammation†;
- grouping and read-across, and discrimination between nanoforms of one substance (section 9);
- occupational inhalation risk assessment† — demonstrated for workplace-collected samples (section 7.3); translation to human airborne concentrations uses the dose-rate relation of section 4.5 with the human parameters of Table 2, which are not validated against human data;
- component of an Integrated Approach to Testing and Assessment in a Weight of Evidence approach — foreseen, not demonstrated here.

† Any further derivation — permissible exposure levels, health-based guidance values, with all the assessment factors they require — should follow the applicable guidance (e.g. ECHA/REACH) ⁸¹.

### Voluntary context of use

- pre-screening during early material development under the Safe-and-Sustainable-by-Design framework, including materials available only in milligram quantities and samples whose intrinsic properties cannot be meaningfully characterised. No regulatory decision attaches to this use.

The method is not a stand-alone replacement of TG 412/413, since it does not predict endpoints downstream of KE1497 (histopathology, fibrosis, proteinosis, systemic availability); however, MoA-based grouping extends the reduction beyond KE1497: when a group member is represented by its reference material, the animal study that is not performed is the complete Test Guideline study, including the endpoints the method itself does not predict (section 9).

### Applicability domain

*In physicochemical terms*, the material must be dispersible but poorly soluble in water (upper solubility cases in the tested set are ZnO100 and ZnO111 with solubility limit 1–10 mg/L and half-solubility time a few hours, SI-8) and must not be a gas or a solvent. There is no requirement for a particular dispersion protocol or characterization of nanomaterials’ physical and chemical properties ^14^, since the sedimentation and biological effects of the nanomaterial are measured within the 17 biological observables in vitro. Therefore, mixtures and cocktails are within the applicability domain provided at least one component meets this criterion regarding poor water-solubility.

*In mechanistic terms* — which is the operational definition — the domain is the set of early Mode-of-Action segments (KERs) covered by the calibration set. A material is inside the domain when every segment of its early MoA is a subset of the union of MoA segments detected during calibration. This is a **response-space** definition, not a chemical-descriptor definition, and it is quantified in section Mechanistic coverage: the method generalises beyond its calibration set. The consequence is that the applicability domain does not depend on the availability of physicochemical characterisation, except from the poor-water solubility, described above — which is what allows the method to be applied to industrial or environmental samples whose intrinsic properties cannot be meaningfully determined (as shown in section 7.3).

## 3 Mathematical structure of the digital twin

The digital twin is a system of coupled first-order ordinary differential equations. Following our earlier formulation^15^,

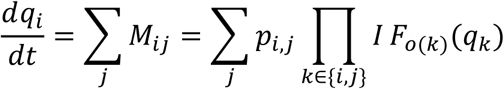

where an interaction term *M_i,j_* is expressed as the product of one or more state functions *IF(q_k_)* and the state-independent interaction coupling *p_i,j_*:

- the entire product has the form of generalised chemical kinetics, where the interaction coupling is the analogue of a reaction constant and the state functions describe how concentrations — more generally activities — affect the reaction rate;
- each state function *IF(q_k_)* is one of a set of predefined, coupling-independent functions, where *k* may be *i* (the affected observable) or *j* (the driving observable);
- the predefined state functions extend beyond the linear and quadratic forms of chemical kinetics to biologically relevant forms such as exponentially saturated linear and quadratic responses, threshold-based responses that generate phase delays, and exponentially decaying responses;
- the diagonal term (i.e. self-regulation) is implicitly included in the sum.

Three properties of this structure carry directly into the interpretation of the results.

### Causal direction is encoded, not assumed

The coupling matrix *p_i,j_* is **asymmetric**: *p_i,j_* ≠ *p_j,i_*. The magnitude and sign of each element state how strongly, and in which direction, observable *j* drives the change in observable *i*. Upstream and downstream position in the Key Event sequence is therefore an *output* of the fit to the measured dynamics, not an input assumption (though couplings between some response classes are precluded, i.e. *p_ij_* set to 0, due to mechanistic implausibility and model restriction to direct effects; e.g. excreted vesicles cannot directly cause material intake by cells). This is only possible because the observables are monitored as time series with hourly resolution: a static snapshot cannot distinguish a driving from a driven event. **Time propagation is mechanistic, not extrapolative pattern matching.** The equations are integrated forward from the in-vitro-parameterised state. Long-time behaviour — including the *resolution* of inflammation after the driving perturbation decays — emerges from the structure of the differential equations, which tend to restore equilibrium in the absence of a trigger, rather than purely from any long-time training data. This is why the method predicts both the rise and the fall of neutrophil influx up to 9 months post-exposure in mice and 6 months post-exposure in rats (sections 7.1 and 7.2), and it is the property that most sharply distinguishes the approach from pattern-matching machine learning-based NAMs, for which extrapolation of empirical non-linear terms (e.g. high-degree polynomials) is likely to diverge from reality.

## 4 Methodology

### 4.1 Design objectives and premises

The following objectives were set: delivery of a toxicologically relevant output able to support regulatory decisions (dose–response curves, LOAEL/BMC, endpoints identical or analogous to those used in the corresponding OECD TGs); applicability to previously untested material, i.e. the ability to unravel a Mode-of-Action without relying on external mechanistic data; reliance on data acquired exclusively using cellular systems in vitro (i.e. eschewing the need for material phys-chem characterization); delivery in a high-throughput-compatible, resource-efficient mode; and prediction for a time window beyond the time scale of the data acquisition.

Our solution to these objectives is based on four **premises**. They are design choices, and they pre-empt several of the questions most often asked of this class of method.

#### Premise 1 — 24 hours of multi-channel live-cell monitoring is sufficient to feed the digital twin

The duration is chosen to cover three early response phases and their time windows: organelle structural changes due to physical interaction (0–4 h), subcellular changes due to biological responses including transcription and expression (2–12 h), and cellular changes due to cell–cell communication, interaction and motility/taxis (6–24 h). Monitoring the local responses of small cellular clusters (with starting confluency levels being below 70% for lung epithelial cells (LA-4) and below 10% for ling macrophages (MH-S) with the ratio of LA-4 to MH-S cell areas (in co-cultures) being in the range 1:30 to 1:6, for full details see the SI-1 Standard operating procedure (SOP) in the supplement) over the 24-hour window gathers enough information to resolve the material-specific interaction couplings and thereby predict local responses together with systemic neutrophil influx from times as short as one hour to times as long as several months after the first interaction.

#### Premise 2 — the biological platform is optimised for the rate and diversity of early responses, not for structural mimicry

The platform consists of submerged mono- and co-culture of murine lung epithelial cells (LA-4) and murine alveolar macrophages (MH-S) in a cell-type surface ratio of 20:1, used as cellular clusters before full confluence. The two cell lines were chosen based on their phenotypic characteristics. The MH-S cell line consists of contact-uninhibited, adherent cells that closely mimic primary alveolar macrophages by retaining typical morphology, phagocytic capacity, increased IL-1 production upon LPS stimulation, and polarization after exposure to stimuli. The LA-4 cell line replicates key features of type II pneumocytes, including contact-inhibited growth, a polygonal morphology, lamellar bodies, and high phospholipid synthesis (more about the baseline phenotype of the cells in the supplement SI-9). The preference for cellular clusters over a tight epithelium is deliberate: a well-evolved epithelium, whose primary function is to maintain a tight barrier, changes as late as possible and therefore has the *lowest* sensitivity to the expression of responses; cellular clusters, being structurally unevolved, are far more dynamic and provide the highest sensitivity while retaining reasonable resemblance of biological responses. As the in vivo outcome is not part of the observed system but is predicted in silico based on the monitored inputs, the biological relevance of the in vitro platform is established **by validation against in vivo data** — the platform is relevant **if the propagated responses match** those observed in animals — and not by anatomical fidelity or direct one-to-one in vitro-to-in vivo response mapping (e.g. expressing same cytokine levels). Because the digital twin is parameterised by *responses* and not by exposure modality, and because submerged exposure allows the delivered dose to be measured per region of interest (section 4.5), submerged culture is the appropriate in vitro cell system for the inflammation-onset Key Events that this method targets.

#### Premise 3 — observables are selected at the functional (organelle) level

Seventeen observables from six classes covering six distinguishable functional units of the cell (Table 2) provide a compromise between what can be observed live in vitro and what is known in vivo as a Key Event connected to other KEs in AOP-Wiki. Selection is **a priori**, from the AOP-Wiki mapping, and is not the result of data-driven feature selection — a distinction that matters for the overfitting question, since no observable was retained or discarded on the basis of how well it improved the prediction. Two further constraints apply: the experiment must interfere minimally with cell dynamics (no fixation, unlike in traditional Cell Painting)^16^, and the number of observable pairs (key event relationships) — the in vitro degrees of freedom — must not exceed the number of available in vivo data points. This is why the method deliberately refocuses from the molecular level (i.e. following all protein and lipid species) to the functional level (i.e. organelles), where the number of degrees of freedom raised to the number of time windows is more than ten orders of magnitude lower and, critically, of the same order as the number of available in vivo data points. The same strategy is used in phenotypic drug discovery^17^, where molecular richness is deliberately traded for a tractable number of monitored units.

#### Premise 4 — Mathematically, a digital twin can be implemented as a system of differential equations assembled from Interaction terms, expressed as products of one or more state-dependent material-agnostic interaction functions and the state-independent material-specific or state-independent material-agnostic interaction couplings (see equation in section 3), whereas in vitro early evolution data can be used to reliably parameterize the material-specific interaction couplings and in vivo data can be used during calibration to parameterise the material-agnostic interaction couplings

Based on the above, the calibration phase revealed and validation phase confirmed that in vitro monitoring of the 17 observables under the defined constraints reliably parameterise all 162+ 2 non-zero material-specific interaction terms, while in vivo mouse response data covering time windows from day 1 to month 6 (and up to 9 months) after single exposure to different doses parameterise the remaining 35 material-agnostic terms. All parameterisations are performed by singular value decomposition. This guarantees **hypothesis-free early Mode-of-Action determination**: any possible, even yet undiscovered, biological pathway at the functional level can be devised, provided the associated events are among the monitored observables (empirical indirect connections over hidden variables may as well be proposed).

### 4.2 Biological observables, their AOP anchoring, and their relation to the TG 412/413 BAL panel

The 17 observables and their AOP-Wiki Key Events are listed in Table 2. The observables span microtubule/transport disruption (KE720), endoplasmic-reticulum stress and mitochondrial dysfunction (KE2017, KE177), disrupted cell cycle and cell injury/death (KE1505, KE1825, KE55), secretion of proinflammatory mediators (KE1496), substance interaction with lung resident cell-membrane components (MIE1495), epithelial and immune cell proliferation (KE1182, KE870, KE1821), immunotaxis, extracellular-vesicle traffic in both directions between the two cell types, and cell-type-resolved material burden.

Three observables — extracellular tubulin debris (Pl4031), expansion/shrinkage of the peripheral (lamella/lamellipodium) region of epithelial cells (Pl4121), and lipid production (Pl4111) — have no direct AOP-Wiki KE. They are **mechanistic refinements**, with an independent literature anchors and, in two cases, a correspondence with a standard TG endpoint. Extracellular tubulin debris (Pl4031) might arise because loss of plasma-membrane integrity leads to misfolding, crosslinking and aggregation of cellular proteins including tubulins^18^, and because necrotic cell death — in contrast to apoptosis, in which microtubules reorganise into a cortical cocoon that preserves membrane integrity^19^ — releases cell remnants and debris^20^; since BAL lactate dehydrogenase (LDH) and double-stranded DNA release also reflect loss of membrane integrity^21^, extracellular tubulin debris is the in vitro analogue of the BAL LDH readout. We confirm this in section 8 in the case of two materials (MWCNT - Mitsui7 and carbon black - Printex90. The peripheral lamellipodial/lamellar region is a branched actin network extending several µm behind the leading edge (ref. Mueller 2017), expansion and shrinkage of this region constitute a direct, mechanistically interpretable readout of material uptake and of membrane perturbation — which is why Pl4121 is retained as a mechanistic refinement despite having no AOP-Wiki Key Event. Lipid production (Pl4111) corresponds to increased lipid synthesis and surfactant excretion by lung epithelial cells ^7,22^ and correlates with the phospholipidosis-type readout listed as an optional endpoint in the TGs.

Table 1 maps each observable to its counterpart in the TG 412/413 BAL panel where such a counterpart exists. The mapping is deliberately conservative. The NAM method predicts the neutrophil-influx readout (KE1497). It offers proxies for parts of the remaining panel: extracellular tubulin debris for BAL LDH, cell-type surface fractions for macrophage cell count, extracellular vesicle (EV) secretion for the cytokine/mediator panel, and toxicant-in-quarantine for phospholipidosis. Formal quantitative validation of the proxy relationships against in vivo BAL LDH and differential counts across a material subset is reported for LDH and dsDNA in in section 8.

### 4.3 Implementation: steps 1 to 5

The set-up of the digital twin for each material relies on five steps (Figure 1a):

**Pre-step** — preparation of the biological platform and of the material dispersion in a standard 96-well plate.

#### Step 1 — monitoring

24 hours of time-lapse monitoring of single exposure responses in vitro by live-cell bi-modal confocal microscopy. Four physical channels of two modalities are used: fluorescence, with CellTracker Green (to label epithelial cells), CellMask Deep Red (membranes), Abberior LIVE 550 Tubulin (tubulin fibres) and CellTracker Orange (immune cells); and label-free back-scatter, for detection of the material itself. Labels were selected from more than 100 candidates for minimal interference with cell dynamics and for compatibility with a standard four-channel confocal microscope. Acquisition is performed in six exposure scenarios (mono- and co-cultures, exposed and control), at 1-hour time resolution, in multiple planes over a large multi-region-of-interest (ROI) field of view at 500 nm spatial resolution, with 3 technical replicates, delivering over 5,000 images per material-treated sample or vehicle-exposed control (in total, approximately 41,500 images per fully utilised 24-hour monitoring run and approximately 168,000 quantified observable time evolutions per day) in each of the typical 2–3 biological repeats.

##### Imaging quality assurance

Any phototoxicity is accounted for in the baseline dynamics of material-unexposed cells. Focus drift is eliminated by the hardware autofocus unit. MH-S macrophages, which are partly in suspension, are handled by acquiring a defined z-stack rather than a single plane and by projecting within a fixed axial window, so that cells transiently leaving the focal plane are not scored as lost. Cell type segmentation, and the surface-versus-internal discrimination criterion for material localisation are reported in the Supplementary Information (SI-5).

##### Seeding density

The SOP specifies a working range for epithelial seeding density. Before exposure and imaging (1-day post seeding), it is again checked that cell confluency is within the specified range (uncompliant samples are discarded), and best reference value matching ROIs selected.

#### Step 2 — quantification

Material-agnostic translation of the image stacks into the 17 (sub)cellular observables, using a fixed database of events and quantification algorithms. No step in the quantification is material-specific, which is what allows the pipeline to be applied to an unknown material. Cell surface area of each cell type is one of the main observables quantified from the images, and an essential input to the model to accurately determine the confluency-independent coupling constants. Wherever mechanistically meaningful in the context of the interaction terms of differential equations, cell-response observables are first normalized to the time-dependent confluency levels, to yield ‘per-cell type’ response intensities.

#### Step 3 — translation

Automated discovery of the material-specific early Mode-of-Action (patented^15^) by translating the quantified observables into the parameterisation of the system of differential equations — approximately 200 material-specific interaction couplings per material. A noise-reducing algorithm is used to derive the time derivatives.

#### Step 4 — time propagation and endpoint evaluation

The material-specific MoA is expanded by the material-agnostic couplings describing in vivo responses, and endpoints are predicted by forward integration, for the exposure scenario of interest (single instillation; repeated or semi-continuous inhalation with post-exposure recovery), yielding time-resolved endpoint trajectories for a chosen dose/dose rate. From a set of runs at different doses, dose–response predictions for each post-exposure time window are generated, from which LOAEL/BMC are read out as values where the dose-response crosses the threshold inflammation level (10% neutrophils). We chose to report the resolution time (i.e. post-exposure time when neutrophils level falls below the threshold) for a 3-fold LOAEL/BMC dose.

#### Step 5 — calibration and validation

The material-agnostic couplings are derived by calibration against in vivo neutrophil influx following single instillation in mice for 15 calibration materials. Materials with in vivo data are split into orthogonal calibration and validation sets; the rules of engagement of in vitro and in vivo data are tuned on the calibration set only and tested blind on the validation set, iterating until both sets deliver consistent quality descriptors. All steps except step 5 are performed for every tested material; step 5 was performed once, with results now being applied to any new material being tested.

A robust SOP has been developed to automate the workflow, improve reproducibility and enable inter-laboratory transferability; transferability has been exercised within the HEurope nanoPASS project from Infinite to the Jožef Stefan Institute. The SOP is organised in three layers; layer 1 is presented schematically in the supplement (SI-1).

### 4.4 Selection of the calibration set: orthogonality of Mode-of-Action

The 15 calibration materials were **not** selected as a random set. They were selected to be **as different as possible in in vivo responses** (i.e. low–high LOAEL, short–long resolution time) to ensure the broadest coverage of endpoint profiles, as well as in **triggered early Mode-of-Action**, i.e. to maximise coverage of the Key Event Relationship network permitted by the model, so that the material-agnostic couplings are constrained by mechanistically diverse ways of both triggering and resolving inflammation. Random selection of the calibration materials, disregarding the in vitro measured mode of action, would be a weaker, not a stronger, design.

### 4.5 Dose metrics and cross-modality dose translation

Interspecies dose conversion requires normalisation of the delivered dose, ideally expressed per unit tissue surface (µg/cm² of lung epithelium or of cell-culture surface)^23,24^. We therefore **measure**, rather than assume, the amount of material that reaches the surface of the epithelial cells. This follows the recommendation of Schmid and Cassee that “cell/lung delivered particle dose should be determined and normalisation of dose to cell/lung surface area is essential for extrapolation of dose levels from cell culture or animal models to humans”^24^, and the finding of Donaldson et al (2008). that surface-area burden is the key dose metric for the elicitation of inflammation in rat lungs by poorly soluble particles^25^.

Natively to the in vitro experiment, dose is measured directly by back-scatter microscopy through four observables, in material-surface (pixel) units per epithelial-barrier surface (pixel). The delivered dose is thus the dose (nanomaterial surface area) that actually interacts with the cell clusters, resolved in time, in three dimensions (through optical sections and between cell types), and in response space (e.g. material associated with damage).

#### On the choice of dose metric

Because the delivered dose is measured per region of interest rather than inferred from the nominal administered amount, the response surface of the digital twin is, to first order, **independent of the dose metric chosen for reporting**. The same measurement yields surface-to-surface dose directly, and — because the in vitro experiment is also sensitive to properties such as surface-to-mass ratio — permits translation into mass-to-lung surface units (mg/m²). Mass-to-surface dose is used as the primary reporting metric, because it can be read efficiently from the in vitro experiment, can be estimated for instillation experiments (a known mass is delivered into a lung of known surface, bypassing breathing-related complications), and is the metric in which the calibration anchor is expressed. Number-based dose can be derived from the same measurement where the primary particle size distribution is resolvable but is not used for reporting because it is ill-defined for aggregates and for the industrial samples in section 7.3.

Where dose is delivered per animal, the lung surface area used for normalisation is **0.008 m²** for a pair of lungs of adult female C57BL/6 mice^26^ and **0.3 m²** for rat lungs^27^. These values may vary by up to a factor of 1.8 in rats, and by up to a factor of 2.5 in humans^28^; this variability propagates directly into the predicted dose axis in addition to the reported prediction uncertainty bands.

#### Instillation-to-inhalation dose translation

For inhalation experiments with repeated semi-continuous delivery, the digital twin uses respirable dose *rates* rather than a single delivered dose. Rates in mg/m²/day are translated into airborne concentration in mg/m³ as

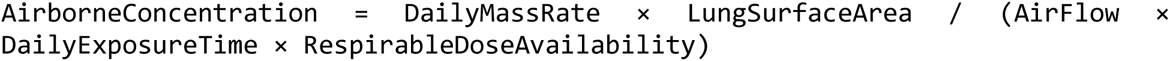

using a typical rat lung air flow of 0.02 m³/h (2.1 ml tidal volume, 166 breaths/min respiratory rate), the lung surface areas above, and the occupational exposure pattern of 6 h/day for 5 days/week. Due to limited information on airborne particle size distribution, required for models like MPPD^90^, we estimated respirable dose availability from the in vitro-detected particle size distribution and results verified with the available material-matched mouse instillation and rat inhalation in vivo datasets. The same relation is used to translate a prediction to conditions other than those assumed here, taking into account interspecies variation in lung surface area and breathing parameters (Table 2). Because airborne concentration is the natural dose representation for inhalation, it is reported as the secondary dose metric for inhalation scenarios. The uncertainty in this translation — dominated by respirable dose availability and by lung surface area — is not propagated into the inhalation prediction bands of Figure 3 d-f and may be the principal reason why inhalation accuracy (84% for LOAEL/BMC) is lower than instillation accuracy (92%).

**Figure 3.**
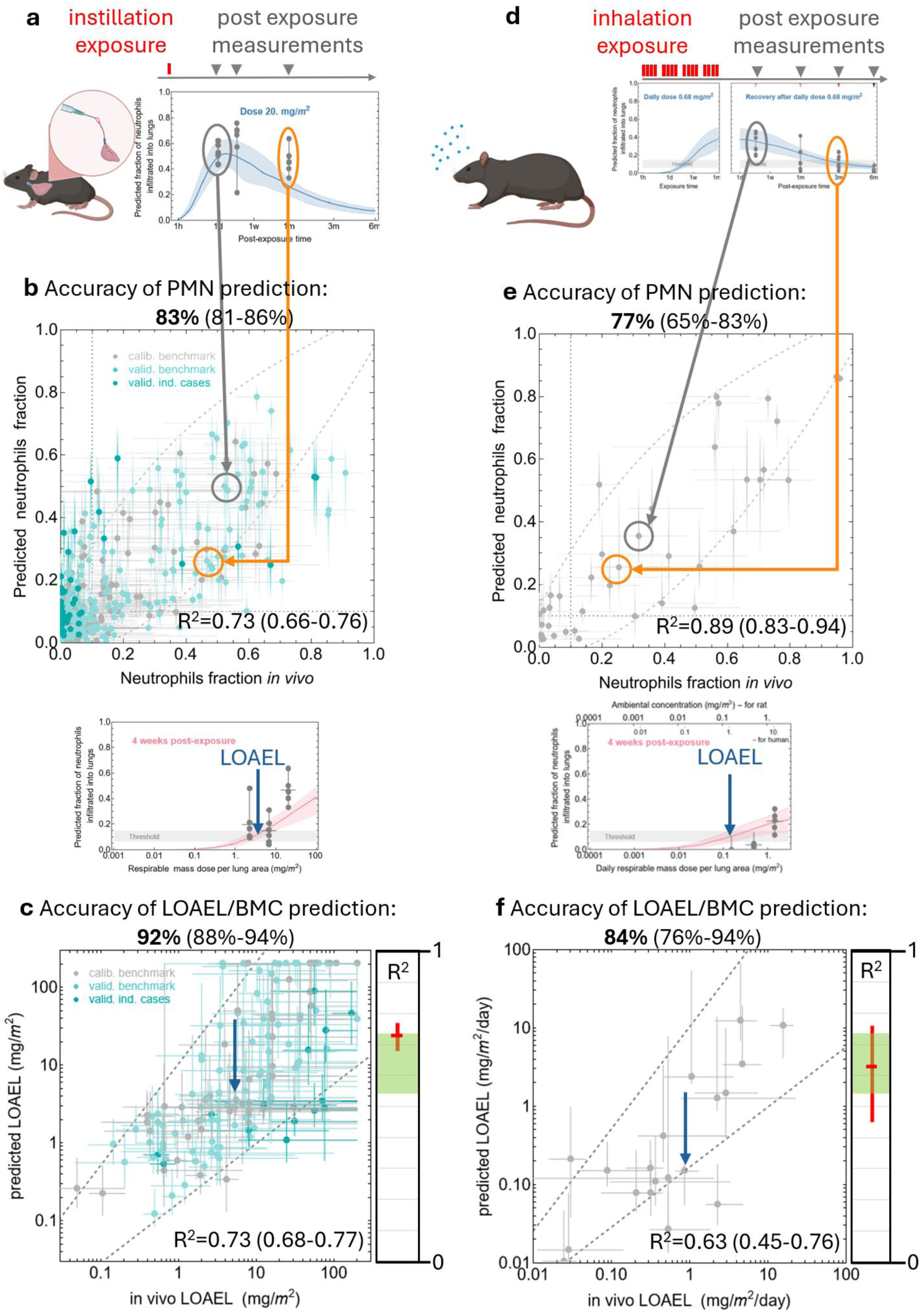
Accuracy (predictive capacity) of the in vitro-learned digital twin InFiniteLungDT for instillation and inhalation (left and right columns, respectively). **a** Examplary data set of neutrophil influx versus post-exposure time, at one dose, for one of the 49 tested materials, with predicted neutrophil influx (blue line with 80% percentile band) overlaid with the neutrophil influx measured in mice after intratracheal instillation (grey circles representing individual mice). b Accuracy of predicted influx of neutrophils, matching in vivo values within the experimental variability band (dashed region) in 83% of cases after intratracheal mice instillation of 44 benchmark and 5 industrial materials at all doses and time points (together nearly 400 data points). c Accuracy of predicted LOAEL/BMCs of lung influx of neutrophils above the inflammation threshold, for different post-exposure observation times, compared with matched in vivo values within the experimental variability band (dashed regions) is 92%; d An example of prediction of neutrophil influx by InFiniteLungDT, based on in vitro measurements of the same material, for nose-only inhalation exposure of rats according to TG 412. e Accuracy of predicted influx of neutrophils, matching in vivo values within the experimental variability band (dashed regions) is 77% for nose-only rat inhalation. f Accuracy of predicted LOAEL/BMCs of lung influx of neutrophils above the inflammation threshold, for different post-exposure observations, compared with matched in vivo values within the experimental variability band (dashed regions) is 84% for nose-only inhalation. The panels on the right of the graphs c and f shows that the R² for goodness-of-fit of these data is 0.73 and 0.63, with a 10–90 percentile range of 0.68–0.77 and 0.45–0.76 (red), for mice instillation and rat inhalation, respectively which is right in the middle of the range of 0.55 to 0.73 (green) that, according to the US EPA Toxicity Reference Database (ToxRefDB) for variability in LOAEL values, meets the US EPA expectation on NAMs^81^.

### 4.6 In vivo reference data: provenance and quality

Most of the in vivo reference data were assembled from studies performed with consistent methodology over the last 15 to 20 years. Mouse instillation data (C57BL/6J BomTac, single intratracheal instillation, up to 80 animals per material, doses spanning at least three levels, post-exposure times from 1 day to 6 months and in selected cases to 9 months) were collected by the National Research Centre for the Working Environment, Denmark (NRCWE), within several EU projects^29,30^ and national projects, using a single BAL protocol and a single administering laboratory ^31,32,32^. Rat inhalation data (nose-only, OECD TG 412 compliant) were contributed by Institut National de Recherche et de Sécurité (INRS)^33,34^ and by BASF. Additional in vivo BAL cytokine and cell damage data for the mechanistic test in section 8 were contributed by Helmholtz Center Munich. In vitro measurements were performed by Infinite and, for transferability, by the Jožef Stefan Institute. Materials for the industrial cases were collected and distributed within the nanoPASS project, with Lund University and other partners.

In total, **3,869 single mouse exposures and 589 rat exposures** were used. This includes **3,094 single mice and 364 rats exposed to different materials**, including approximately 775 vehicle-exposed mouse and 225 rat controls. All in vitro measurements were performed on material from the same batch as the corresponding in vivo experiment — which is the practice the JRC Nanomaterials Repository was established to enable, and which removes batch identity as a confounder, or on material collected and distributed simultaneously to the in vivo and in vitro laboratories. Experimental details about the in vivo and in vitro data collection are provided in supplement (SI-6).

### 4.7 Definition of the inflammation threshold, of LOAEL/BMC, and of accuracy

#### Inflammation threshold

The LOAEL/BMC is defined as the dose at which the predicted neutrophil fraction exceeds a significance threshold. That threshold depends on the variability of the neutrophil fraction in vehicle-exposed control animals and was set to 10%. The empirical distribution of the BAL neutrophil fraction across the 775 vehicle exposed control mice and 225 control rats in the database distributed on different post-exposure time points, and from which these values are derived makes the threshold an empirical property of the reference dataset rather than a modelling choice.

#### In vivo LOAEL

Conventionally, the in vivo LOAEL is the defined as the lowest dose with a statistically significant effect above the control level. Because few of the assembled experiments have enough data points at enough doses for this rule to be applied directly, the most probable *range* for the in vivo LOAEL/BMC was estimated by fitting the 10th and 90th percentiles of the in vivo data and determining the doses at which those percentiles cross the threshold. The resulting range is what is plotted as the in vivo variability band in Figure 3c and 3f.

#### Definitions used in the accuracy calculations

- The *animal variability band* (in vivo experimental error band) is the area covering 95% of all animal data points at each mean neutrophil-influx value, or LOAEL/BMC. Note that for the neutrophils fraction, the values of which are bound within 0–1, the variability band is broadest in the middle of the range, while for unbound LOAEL values the in vivo uncertainty is highest at high LOAELs, i.e. for conditions where little effect is seen even at the highest evaluated doses.
- A prediction is scored as *correct* when it falls within that band. “Equivalent” therefore has a defined meaning: inside the empirical animal variability band, which for LOAELs corresponds on average to within a factor of about 3 in either direction.
- *Accuracy* is the percentage of predicted values (neutrophil influx, or LOAEL) falling within the band of in vivo data. It uses a continuous-space definition, in contrast to sensitivity and specificity (i.e. true positive and true negative rates, respectively), which use binary definitions relative to a predefined threshold. Both are reported.
- As another continuous alternative, the goodness-of-fit metric R² is reported. For LOAELs, values are log-transformed first, since they span several orders of magnitude.
- Confidence intervals on all accuracy percentages and R² values are obtained by bootstrap resampling of materials (not of individual data points, which are not independent within a material), and are reported alongside the point estimates in Figure 3 and in the supplement (SI-2b and SI-3b).

We note that obtaining a substantially better R² for the in vitro data compared to the in vivo variability floor would indicate overfitting — fitting with too many free parameters — rather than a better method.

### 4.8 Materials

In total **53 materials** were involved in this work: **44 benchmark materials** and **9 industrial samples**. Of the 9 industrial samples, 5 have matched in vivo instillation data and are included in the instillation validation statistics, giving **49 materials with mouse instillation data**; **7 materials** (4 industrial and 3 benchmark) have rat inhalation data. The entire material set comprises (for full material specifications, including JRC and NIST codes see supplement SI-4 Material characterization table):

- **3 spherical isometric magnetite alloys**: NiFe₂O₄ N022 (25 nm), NiZnFe₄O₈ N020 (20 nm), ZnFe₂O₄ N021 (22 nm);
- **20 carbonaceous nanomaterials**: **2 graphene oxides** (GO N058, 1.7 × 2500 nm, highly hydroxylated sheet; GO N059, 1.7 × 1500 nm, reduced); **15 multi-walled carbon nanotubes** (MWCNT NM-401, 67 × 4000 nm; N006 (Mitsui-7), 74 × 5700 nm; N040, 22 × 500 nm; N041, 27 × 1000 nm, –OH; N042, 30 × 700 nm, –COOH; N043, 27 × 800 nm; N044, 33 × 1300 nm, –OH; N045, 30 × 1550 nm, –COOH; N046, 29 × 700 nm; N047, 23 × 500 nm, –OH; N048, 18 × 1600 nm, –COOH; N061, 16 × 700 nm, –NH₂; N062, 9 × 450 nm; N063, 14 × 350 nm, more –OH; N064, 7 × 2000 nm, more –COOH); **3 single-walled carbon nanotubes** (SWCNT N054, 17 × 17500 nm, –COOH; N055, 14 × 2000 nm; N056, 13 × 2000 nm, –OH);
- **5 clays** (phyllosilicates): HNTNN (61 × 250 nm, elongated halloysite), HNTNNE (27 × 100 nm, etched halloysite), Nanoclay 3000 (300 nm sheets, organoclay functionalised with dialkyldimethyl ammonium), Nanoclay 9 (300 nm sheets, montmorillonite functionalised with benzalkonium chloride), Nanoclay Bent (650 nm bentonite flakes);
- **5 combustion-generated particles**: Diesel N1650 (24 nm, heavy-duty truck exhaust particles, NIST1650), Diesel N3 (22 nm), Exhaust N1 (15 nm, rapeseed methyl ester), Exhaust N5 (21 nm, hydrotreated vegetable oil)
- **3 carbon blacks** Flamruss 101 (95 nm carbon black), Printex 90 (14 nm carbon black), Printex XE2B (30 nm carbon black);
- **9 (semi)metal oxides**: TiO₂ large (16–28 nm anatase/rutile spheres), TiO₂ N001 (10 nm rutile), TiO₂ N025 (38 nm rutile), TiO₂ N030 (12 nm rutile), TiO₂ nanocubes (15 × 20 nm anatase), TiO₂ nanotubes (10 × 100 nm anatase), ZnO 110 (70–110 nm uncoated zincite), ZnO 111 (60–90 nm, coated with hydrophobic triethoxycapryl silane), crystalline SiO₂: DQ12 (225 nm quartz);
- **9 industrial materials** provided by nanoPASS partners and by industrial partners of Infinite: nanoplastics, WEEE recyclate, clinker, cement, NiMo particles, and 4 undisclosed materials with undetermined intrinsic properties.

Specific surface area (BET), material density, nominal dimensions, ranked dissolution of the core material and ranked release potential of toxic metals are tabulated per material in the supplement (SI-4), together with the calibration/validation set assignment. Solubility of both ZnO materials set the edge of the stated applicability domain;its treatment is addressed in section 10.3.

## 5 The digital twin and its output: worked example

By applying the in vitro discovered MoA (Figure 1 b) to a pre-selected exposure scenario – single intratracheal instillation of a material (in this exemplary case, data for anatase TiO₂ nanotubes are shown) the digital twin predicts the temporal evolution of the neutrophil fraction in the lung for any chosen dose (coloured curves and bands in Figure 1 c), which can be compared directly against the individual-mouse in vivo values (gray datapoints). From predictions at various doses, dose–response curves are generated for each post-exposure time window (Figure 1 c). Comparing all values of lung neutrophil fraction shows that most of the predicted values lie within the in vivo variability band (Figure 1 e). Reading each dose– response curve at the inflammation threshold yields the benchmark dose or LOAEL/BMC for that post-exposure time window (Figure 1 d). Comparing the LOAELs/BMCs for anatase TiO₂ nanotubes measured in mice and predicted by the digital twin, all five values — one per time window — lie within the in vivo determined diagonal area (Figure 1 f), even though the TiO₂ nanotubes were excluded from the calibration set. This example illustrates the regulatory relevant outputs that the method delivers: an in vitro determined material specific early mode of action (eMoA network) and a dose–response curves of neutrophilic lung inflammation for LOAEL/BMC determination.

## 6 Mechanistic anchoring of the observables in the AOP framework

*Input relevance:* Analysis of AOP-Wiki identified 250 Key Events associated with the keyword *inflammation*, together with 281 KEs directly upstream or downstream (Figure 2, left panel). Among them, KE1497 — neutrophil influx — is a particularly robust and consistent marker of particle-induced inflammation in vivo, and it is the central endpoint predicted by InFiniteLungDT. KE1497 is part of the OECD-endorsed AOP173 ^35^. To predict KE1497 we monitor 17 observables corresponding to 9 KEs (Table 1). This visualization attests that the chosen observables are broadly spread over the known mechanistic space, providing the **basis for the method’s sensitivity to material-specific triggering of the material-agnostic adverse outcome**. Already within existing toxicological knowledge, there are 6 linear pathways from the selected KEs to KE1497, and 12,953 cycles comprising KE1497 and at least one of the monitored KEs.

Noteworthy, among the monitored variables, InFiniteLungDT provides **hypothesis-free early Mode-of-Action analysis** for each tested material; any possible, even as yet undiscovered (thus not yet included in the AOP analysis above), biological pathway at the functional level can be found – direct or indirect via hidden variables. Observing a limited set of functional response of cells at a higher-level (cell organelles and intercellular interactions), which themselves have already integrated inputs across numerous molecular entities (e.g. individual cytokines) thus ensures high sensitivity of the approach.

### Endpoint coverage

The right-hand panel of Figure 2 shows what the method delivers against the BAL panel determined according to TG 412 and TG 413: 1) one endpoint is predicted and pre-validated, some are proxied, some are not covered. Thus, as shown in the figure below the InFiniteLungDT is not a complete substitute for TGs 412/413. Rather, it could support the replacement of an in vivo inhalation study in situations where systemic toxicity is not expected, for example based on available data from other routes of exposure, and the primary concern is local pulmonary toxicity. In addition, the method could provide mechanistic information to support grouping and read-across of materials and may help reduce animal studies.

## 7 Predictive capacity against in vivo data

### 7.1 Mouse instillation: the calibration anchor

The right comparator for a method that predicts an animal endpoint is not perfect agreement with animal data (i.e. fit quality R²∼1) but **the reproducibility of the animal experiment itself**. According to the US EPA Toxicity Reference Database, the inter-study variability of LOAEL/BMC values corresponds to R² in the range 0.55–0.73^81^. This is the band within which a NAM meets the US EPA expectation, and it is the band against which the numbers below should be read. Note that predictions close to R²∼1 are probably indicative of overfitting of a particular limited dataset.

Figure 3 a–c shows the accuracy of InFiniteLungDT predictions against intratracheal instillation in C57BL/6J BomTac mice for 49 materials. The data were collected by NRCWE using consistent methodology within several EU projects over the last 15–20 years, with up to 80 animals per material, at different doses and post-exposure time points from 1 day to 6 months (and to 9 months in selected cases). ^29,36–65^

One exemplary data set, at one of the doses, for one of the 49 materials is shown in Figure 3 a: the predicted neutrophil influx (curve with uncertainty band) describes the in vivo measurement (gray datapoints) within the error bars of the in vivo data. Across all materials, at approximately 3 doses and 3 to 5 post-exposure times — nearly 400 data points in total — the predicted versus measured comparison of Figure 3b gives R² of 0.73, and places **83%** of predictions inside the in vivo variability band. Binary evaluation relative to the inflammation threshold gives **74% specificity** and **84% sensitivity**. Predicted values show about one third lower variability (i.e. better repeatability) than the in vivo data. All scores are similar for the calibration and validation subsets (gray and green datapoints, respectively), indicating no overfitting.

The correlation of in-vivo-derived and predicted LOAELs/BMCs across all materials used in calibration and validation (Figure 3 c) has **R² = 0.73** for goodness-of-fit, close to the upper bound of the ToxRefDB range, with **92%** of predicted LOAELs/BMCs inside the in vivo variability band indicated by dashed gray lines. Predicted LOAEL/BMCs deviate from in-vivo-determined LOAELs/BMCs on average by less than a factor of 3 in either direction. The quality of prediction was similar across material families — alloys, metal oxides, clays, carbonaceous materials and combustion condensate cocktails. The plots for individual post-exposure time windows (SI-11) do not indicate any notable bias.

The best empirical evidence against overfitting remains that calibration and validation scores are comparable, and that R² does not exceed the variability floor of the reference data.

### 7.2 Rat inhalation according to OECD TG 412: the regulatory validation anchor

One objection is routinely and correctly raised against instillation-calibrated methods: instillation is a bolus rather than an aerosol, with vehicle effects and focal high lung burden. This concern is addressed here by data rather than by argument.

Once calibrated, the digital twin can predict responses for any exposure pattern — continuous, repeated, with post-exposure recovery — because the exposure scenario enters only as the forcing term of the differential equation system and not through any of the couplings, i.e. the material-specific MoA information coming from the in vitro experiment is assumed to apply to any exposure type, but – in the first approximation – also a similar animal species (with lung and breathing parameters properly adjusted).

To confirm applicability to OECD Test Guidelines, Figure 3 d shows the neutrophil influx measured at three post-exposure times in rats continuously exposed to 1.4 mg/m³ MWCNT (JRC NM-401) for 28 days according to TG 412^33^. The predicted BAL neutrophil fraction is shown for the exposure window and for the following post-exposure period, up to 6 months. **This is not a fit**: it is a prediction computed from the early Mode-of-Action extracted from the in vitro measurements, compared against 66 rats including controls, measured previously by Gaté et al. (2019)^33^. (It is also the worked example of the instillation-to-inhalation dose translation of section 4.5: the mouse instillation calibration is expressed in mass-to-surface units, and the rat airborne concentration of 1.4 mg/m³ follows from the dose-rate relation.)

Across all 7 materials with rat inhalation data, accuracy is **77%** for the neutrophil-influx trajectory (Figure 3 e) and **84%** for the LOAEL/BMCs (Figure 3 f), with **R² = 0.63** (10–90 percentile 0.45–0.76) — inside the ToxRefDB 0.55–0.73 green band. Predictions for individual materials, with all in vivo data points, are shown in the supplement (SI-2 and SI-3).

The two-tier structure should be read as intentional: **instillation is the calibration anchor**, because the delivered dose is known and the dose–time matrix is dense enough to constrain the material-agnostic couplings; **inhalation is the regulatory validation anchor**, because it is the modality of the Test Guidelines, that allow both rats, mice and other rodents. Neither alone would be sufficient. That parameters calibrated on mouse instillation predict rat inhalation to within the animal variability band is the strongest available evidence that the digital twin encodes response biology rather than the idiosyncrasies of the exposure route. As more rat (and eventually human) inhalation data become available and if any discrepancies are observed, the interspecies variability may also be integrated into the systemic response-describing material-agnostic terms of the digital twin’s next generation. Of note, neutrophil influx in rats following inhalation and instillation of two different carbon nanotubes was on the same dose-response curves 1-3 and 28 days after end of exposure ^66^. Furthermore, mice and rats had very similar neutrophil influx and pulmonary transcriptional response following MWCNT exposure (two different MWCNTs) by intratracheal instillation ^39^.

### 7.3 Industrial real-world cases

Five of the nine industrial samples — two collected along a cement production line, one sampled in a recycling plant for waste from electrical and electronic equipment (WEEE), artificially degraded 3D-printed plastics, and an engineered bimetal nanomaterial —are included in the statistics above. For these samples the quantity collected is typically too small to permit detailed characterisation of intrinsic material properties (which in turn can be very heterogeneous across the sample), yet sufficient for parallel in vitro and in vivo experiments. Thus, we measured the same batch of collected real-world air-borne materials with the digital twin method and in vivo mouse instillation experiments. The accuracy of predicted neutrophil fraction is similar to benchmark materials (valid. ind. cases, dark green data points, in Figure 3b and 3c). This is the clearest demonstration that the applicability domain of the method is not defined by physicochemical characterization of materials. The in vitro learned digital twin method seems to be the only existing method applicable to incompletely characterised real-world samples, for which a descriptor-based method cannot be applied even in principle.

### 7.4 Mechanistic coverage: the method generalises beyond its calibration set

Even though we calibrated the digital twin on a finite set of materials, it applies to any other, here untested, material – as demonstrated above with validation materials (both benchmark and industrial).

Considering all possible causal relationships among the biological events described by the differential equations, the model spans a very large number of early adverse outcome pathways leading to KE1497, assembled from approximately **200 permitted Key Event Relationships**. If every material triggered only a single pathway, an impractical number of materials would be needed to cover this network. In fact, every material triggers a *subnetwork* considerably larger than a single specific linear AOP — materials are less mechanistically specific than usually anticipated — so a relatively small number of well-chosen materials can cover the accessible mechanistic space. On average a material covers 16% of KERs, ranging from 10% to20%, as can be seen in the supplement (SI-2).

As quantified in Figure 4a, the 15 calibration materials together cover **92%** of the permitted KERs; across all 49 materials the coverage rises to **98%**; only **6%** of KERs present across the full set are absent from the calibration set, and these are shown explicitly in the right-hand panel. As prediction of the neutrophilic inflammation was comparably accurate for validation as for calibration materials, and largely within the in vivo variability band, the additional few KERs did not have considerable impact on the final prediction. A material outside the calibration set is therefore inside the applicability domain whenever its early MoA is largely covered by the union of the calibration MoAs — a condition satisfied with high probability at 92% coverage, and verifiable per material rather than assumed.

**Figure 4.**
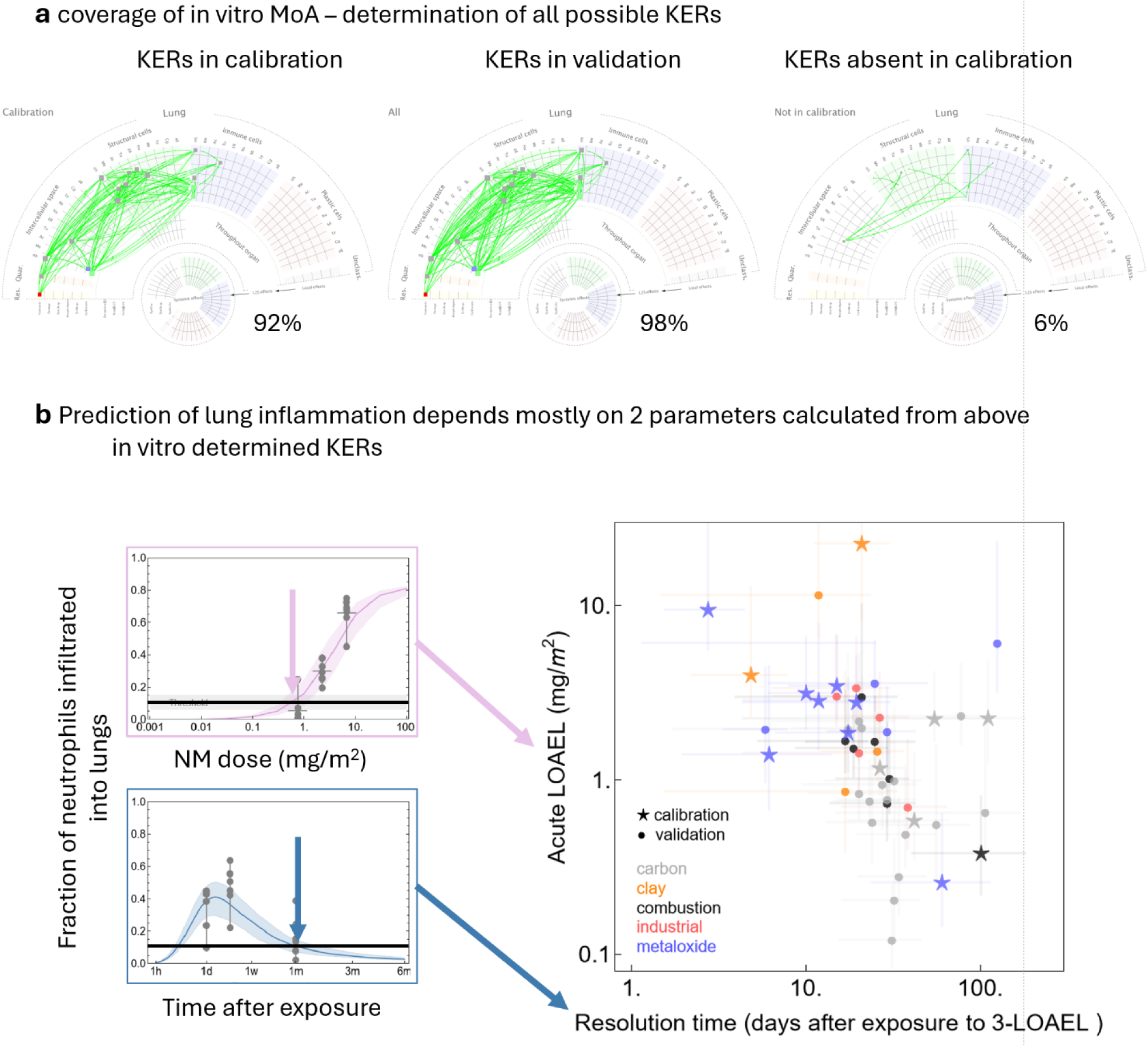
**InFiniteLungDT works for materials outside the calibration database** due to high coverage of the MoA by the calibration materials, without the need to know intrinsic material properties. **a** KERs identified in the in vitro MoA of the 15 calibration materials (left) and of all 49 materials (centre) cover 92% and 98% of all possible KERs of the model, respectively; those not significantly present among the calibration materials are also shown (right). **b** Neutrophilic inflammation is characterized by 2 features: acute LOAEL/BMC and resolution time. For calibration, we selected materials that widely cover this 2-dimensional space, while the validation materials and industrial cases lie distributed within, confirming that the digital twin should be able to predict neutrophilic lung inflammation for any future unknown material within this feature space.

Figure 4b provides the complementary empirical check in the space of *predictions* rather than of mechanisms. The kinetics of predicted neutrophilic inflammation is described by two characteristics: the acute LOAEL/BMCs and the resolution time (determined at 3-fold LOAEL/BMC dose). The calibration materials were chosen so that they broadly cover this 2-dimensional feature space, while the values for the validation materials and for the industrial cases lie distributed within the calibration span — there is no region of this two-parameter space that the calibration set fails to populate and into which validation materials fall. This is the reason to expect the digital twin to predict neutrophilic lung inflammation for a future unknown material, and it is a statement that can be tested for any new material before its prediction is used.

Generalisation is argued at the level of **mechanism coverage**, which is measurable in the same units for a calibration material and for an uncharacterised industrial sample, rather than at the level of **chemical similarity**, which cannot be evaluated when the chemistry is unknown, or if the tested material is a heterogeneous mixture. Chemistry-level overlap between calibration and validation sets — five or six material families across both — would be a legitimate concern for a chemistry-anchored model but is not an issue for our approach.

### 7.5 The effective concentration phase space is continuous, not a single point

A related question concerns the concentration at which the in vitro experiment is performed. Each experiment is run at a single *nominal* concentration, which invites the objection that couplings nonlinear in dose — macrophage saturation, lysosomal overload — could not be resolved, and that dose–response prediction would then be an extrapolation from a single point.

The parameterisation does not, in fact, sample a single point in concentration. Three properties of the measurement make the sampled space continuous:

1. **The two biological replicates per material are run at different concentrations**, by design, so that even the nominal dose is not single-valued. For one material in vitro response is monitored for 24 hours with 2 subsequent monitoring runs representing 2 biological replicates each comprising 3 technical replicates.
2. **The delivered dose is measured per region of interest** (i.e. a technical replicate), by back-scatter, and it varies substantially between cell clusters within one field of view. Each ROI is therefore an independent observation at its own delivered concentration. Thus, we obtain the concentration dependent response of biological observables, i.e. we measure the biological response as a function of time and concentration of delivered material.
3. **The response is dynamical.** Delivered dose per ROI evolves over the 24 hours by sedimentation, uptake, redistribution between cell types, and quarantine; the fit sees the whole trajectory, not an endpoint at a fixed concentration.

The couplings are consequently constrained across a continuous **effective concentration phase space**, spanning the range in which the dose–response relation has appreciable slope, rather than at one nominal value. The state functions *IF(q_k_)* of section 3 include exponentially saturated forms precisely to represent the finite capacity of biological responses such as macrophage uptake saturation, and the saturation parameters are identifiable from the ROI-resolved and time-resolved dose variation. A direct visualisation of the sampled phase space per ROI is provided in SI-10.

## 8 Test of the Mode-of-Action prediction against in vivo BAL cytokine release and cellular damage

### 8.1 The MoA prediction matches in vivo BAL bulk measurements of cytokine release and cellular damage

The digital twin has been calibrated to predict the primary inflammation endpoint – neutrophils influx. It does so on the basis of the in vitro-determined MoA for each material, and predicts each observable’s in vivo time course as well, but for these predictions very few reference in vivo data exist for direct comparison. To best demonstrate the value of the Mode-of-Action provided by the digital twin, we qualitatively compared the predicted MoAs for exposure to two types of carbon-based nanomaterials with in vivo data on mechanistically related events: Printex90 (20 µg) and Mitsui7 MWCNTs (15 µg). We exposed C57BL/6J mice to these two nanoparticles intratracheally and measured cytokines released into bronchoalveolar lavage fluid (BALf) at 0.5, 6 and 28 days after exposure. All the up-regulated measured cytokines — CCL2 (chemokine (C-C motif) ligand 2, also known as monocyte chemoattractant protein-1, MCP-1), CXCL1, IL33 and lipocalin-2 (LCN2) — were increased more prominently in mice exposed to Mitsui7 (Figure 5 c, LCN2 and CCL2, others not shown).

**Figure 5.**
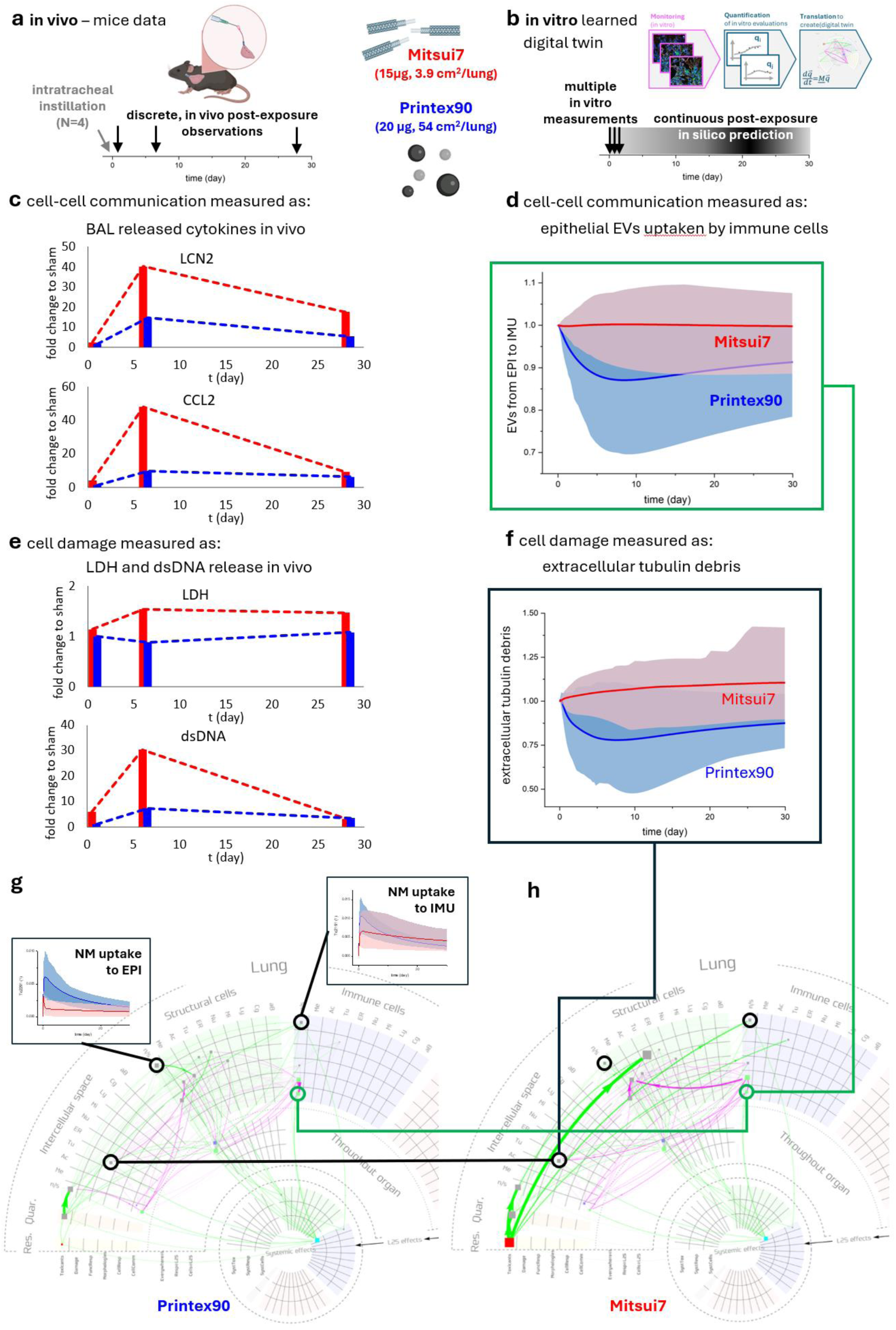
Comparison of the Mode-of-Action (MoA) prediction by the In Finite in vitro-learned digital twin after exposure to two types of carbonaceous nanoparticles with in vivo mice data: Printex90 (a carbon black nanoparticle) and multi-walled carbon nanotubes (MWCNTs – Mitsui7). a Schematic presentation of the intratracheal exposure in vivo experiment. b Schematic presentation of the in vitro-learned digital twin. c Cell–cell communication measurement in bronchoalveolar lavage (BAL cytokines) after exposure of mice to Printex90 (blue) and Mitsui7 MWCNTs (red) at three different post-exposure time points (0.5, 6 and 28 days). d Digital twin prediction of extracellular vesicles released by epithelial cells (EPI) and taken up by immune cells (IMU). e Cell plasma membrane damage measured as extracellular LDH activity and dsDNA release in BAL in vivo at three discrete time points, with dashed lines connecting the measurements to guide the eye. f Digital twin prediction of extracellular tubulin cell debris. g Mode of action (MoA) for Printex90 exposure. h MoA for Mitsui7 MWCNTs exposure in vitro at the same concentrations as in the in vivo experiment, highlighting main differences in the mode of action between the two materials. Insets in g – material uptake to epithelial cells (EPI, or AEC(alveolar epithelial cell)) and macrophages (IMU, or MΦ) show higher uptake of Printex90 but slower clearance of Mitsui7 from the macrophages

We observed a similar difference between the two tested nanomaterials in the amount of extracellular vesicles (EVs) released from epithelial cells and taken up by immune cells, Pl6391 — EVs from EPI to IMU (Figure 6d). While the cytokines shown in Figure 5 c detected by ELISA in BAL were free and not EV-entrapped, the concordance is in line with the measurements of Fitzgerald et al. (2018), showing that cytokines are not only released as soluble factors, but can be packaged and released inside or on the surface of EVs, to execute cell–cell communication^67^. They evaluated the relative amounts of different cytokines in the free (soluble) fraction and in the EV fraction for different cell types: in one system a cytokine was released mostly in soluble form, whereas in another the same cytokine was released in EV-associated form. Most notably, they showed that cytokines released by immune cells (monocytes and T cells) are mostly released in an EV-associated form. Cell–cell communication via EV-encapsulated cytokines represents a paradigm shift in immunology and molecular biology beyond the traditional view of signaling via soluble cytokines ^68^.

**Figure 6.**
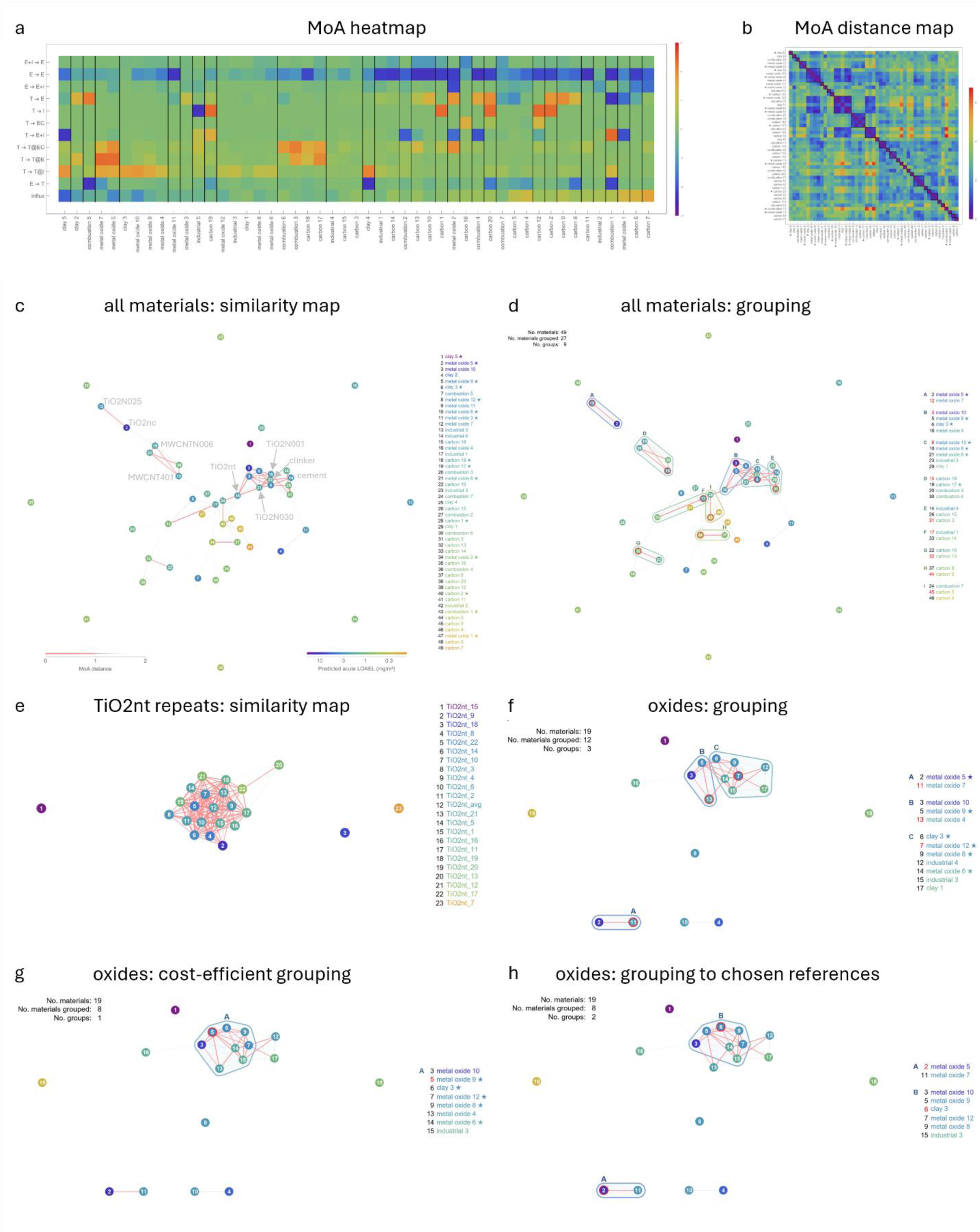
Grouping and discrimination of materials by early Mode-of-Action (eMoA). Material identities are anonymised by family, except where annotated in panel c; stars mark the 15 calibration materials. **a** Effective in vitro-determined Mode-of-Action by classes of observables (rows: driving → driven functional compartment, E-epithelial cells, I-immune cells, T-toxicant and damage, EC-extracellular, influx-rate of neutrophils recruitment; columns: materials), colour-coded in units of uncertainty. **b** Map of eMoA distances between all material pairs. **c** Similarity map of all 49 materials: red links join pairs below the grouping threshold, grey links pairs above it, with line visibility disappearing with distance; node colour encodes the predicted acute LOAEL/BMC (mg/m^2^). Annotations identify selected materials and show that nanoforms of one substance can separate while chemically unrelated materials can coincide. **d** One possible grouping of all materials at the threshold: 27 of 49 materials fall into 9 groups; the chosen reference material of each group, which can replace the other members in an animal test, is labelled in red. **e** Similarity map of 22 independent repeat measurements of one reference (benchmark) material (anatase TiO2 nanotubes), demonstrating repeatability: all but three repeats form a single cluster below the threshold. **f–h** Three groupings of the same 19 oxide-containing materials under different objectives: **f** all materials that can be grouped (12 of 19 in 3 groups); **g** the most cost-efficient grouping, maximising the number of registered materials per unit of testing investment at an in-vitro-to-in-vivo cost ratio of 1:20 (8 of 19 in 1 group); **h** grouping to chosen references, e.g. materials already registered or already tested in vivo (6 materials attach to 2 references).

Another clear distinction between Mitsui7 and Printex90 exposure in vivo is that Mitsui7 induces much more release of double-stranded DNA (dsDNA) and lactate dehydrogenase activity (LDH) than Printex90 (Figure 5 e). Both are routinely considered a sign of cell damage and plasma membrane disruption. The same might hold true for the in vitro monitored, and in silico modelled, tubulin debris release. Macrophages are supposed to internalise and degrade such extracellular debris actively, to maintain tissue homeostasis. As expected, extracellular tubulin debris Pl4031, predicted by the digital twin, is also elevated in the Mitsui7-versus Printex90-exposed case (Figure 5 f). One can see that levels of extracellular tubulin debris are higher after exposure to Mitsui7 than to Prinex90, both in vitro (Figure 5 f), and in vivo (Figure 5 e). During regulated cell death such as apoptosis, microtubules reorganise to form a cortical ring or cocoon-like structure during the cell-shrinking process, preserving plasma membrane integrity^19^. Necrosis, by contrast, is an overarching term describing cell-death modalities caused by extreme adverse conditions in which cells lose structural integrity, and rupture of plasma membrane, resulting in spills of internal necrotic cell remnants, or debris^20^. Moreover, Samson et al (2012). showed that cell death and loss of plasma membrane integrity lead to misfolding, crosslinking and aggregation of cellular proteins including tubulins^18^. Since LDH and dsDNA release into bronchoalveolar lavage fluid or other extracellular fluids reflects loss of membrane integrity^21^, it should not be surprising that extracellular tubulin debris predicted by the digital twin correlates with LDH activity and dsDNA release after exposure to Mitsui7 in vivo. To further confirm the type of sudden cell death, we used the necroptosis inhibitor necrostatin and an interleukin-1 (IL1) inhibitor in bone-marrow-derived macrophages in vitro, which indicated that Mitsui7-triggered CCL2 release was dependent on necroptosis and IL1R signalling (manuscript under revision).

### 8.2 The MoA yields mechanistic insight beyond what is measurable in vivo

The in vitro-extracted MoA also offers additional insight into the mechanism of action beyond what can be measured in vivo in exposed mice. Namely, the MoA shows that Printex90 is taken up by both epithelial cells and macrophages more than Mitsui7 (Figure 5 g, insets: NM uptake to EPI and NM uptake to IMU, respectively), in agreement with the LDH and dsDNA data showing that Printex90 does not cause sudden cell death and can therefore be taken up by the cells in greater amounts. We also observe that clearance of Mitsui7 from immune cells is slower than clearance of Printex90. This is in agreement with biokinetics of carbon black and multi-walled carbon nanotubes reviewed by Hadrup et al. (2024) ^69^, where clearance of Printex90 from lungs of female B6C3F1 mice ^70^ faster than clearance of Mitsui7 from lungs of male C57BL/6 J mice exposed in a whole-body inhalation system ^71^. Also the predicted clearance of total nanomaterial by the digital twin agrees with the observation of lung burden clearance in vivo (SI-12). The majority of clearance half-times, predicted by our model for the 49 nanomaterials included in this study, lie between 20 and 50 days, which is consistent with measured in vivo half-times from different in vivo studies performed with similar materials reviewed in Hadrup et al. (2024 and 2025) ^69,72^. However, the lack of in vivo data with enough post exposure time points that allow accurate quantification of lung clearance for these exact materials precludes a more precise comparison.

### 8.3 Value of MoA information

The in vitro-learned digital twin methodology presented here offers three unique advantages with respect to other currently available methods:

### Differential cell type-specific uptake is measured, not neglected

Material burden inside epithelial cells (To2191) and inside immune cells (To2291), which cannot be discerned in vivo but are important in triggering systemic inflammation, are separate observables with separate material-specific couplings, and Figure 5 g/h shows them differing between two materials. Uptake kinetics resolved by cell type is an output of the method.

### One observable without an AOP-Wiki Key Event is validated against standard in vivo readouts

Extracellular tubulin debris (Pl4031), predicted in vitro, tracks BAL LDH and dsDNA measured in vivo in the same experiment. This observable therefore represents a NAM for a new KE candidate for AOP-Wiki.

### The sign of a coupling is interpretable

The negative coupling between Printex90 uptake and extracellular debris, and the positive coupling between the same uptake and cortical actin area, are predictions about mechanism that were not put into the model and that turn out to be consistent with independent in vivo damage markers. This is what the asymmetry of *p_i,j_* (section 3) provides: a directional (cause and effect), experimentally verifiable mechanistic claim per material rather than a similarity score.

## 9 Grouping and read-across by early Mode-of-Action

### 9.1 What a grouping method has to deliver

Reduction of animal testing by grouping and read-across is explicitly foreseen in REACH Annex XI, Section 1.5, which allows toxicological properties to be predicted from structural similarity, in vitro profiles or in silico data. For nanoforms the practical obstacle is that each nanoform is assumed to be different a priori and must therefore be supported by its own dataset. A registrant placing twenty nanoforms of one substance on the market in the 10– 100 t/year band may need a repeated-dose study on every one of them — of the order of €6 million and more than 3,000 animals for a single information requirement^83^. Characteriser-based frameworks developed to identify when several nanoforms may be treated as one, built on the Annex VI identifiers (size, shape, crystallinity, surface area, surface treatment), are workable but deliberately conservative: the thresholds they use are constrained by the sparseness of the hazard data behind them, and the final report of that work recommends the integration of New Approach Methodologies — in vitro, in silico and mechanistic data — to bridge precisely this gap^83^.

Grouping by intrinsic physicochemical properties alone is insufficient to predict the severity of pulmonary inflammation, because clear links between individual characteristics and toxic effects remain unestablished^77^. The ECETOC Nano Task Force proposed, in the DF4nanoGrouping framework, that the specific mode of action resulting in a toxic effect is the crucial element for grouping^78^, and that meaningful grouping should integrate intrinsic characteristics, life-cycle changes, biokinetics (whether the material reaches the target site) and observed biological effects including cellular uptake, intracellular trafficking and toxic responses^77,78^. In our own dataset, neither material family nor specific surface area organises the observed LOAELs/BMCs and resolution times into coherent groups (SI-7).

Two requirements define a usable grouping method and pull in opposite directions:

- sensitivity (discrimination) — materials that are not the same, including different nanoforms of the same source material, must not all be grouped; and
- robustness (repeatable similarity) — independent repeats of the same material must be recognised as similar (note that even the same nanomaterial might elicit different biological response as it ages, aggregates, oxidizes, or changes its phys-chem characteristics by any other means).

The methodology described below demonstrates that these two requirements are balanced on measured data.

### 9.2 The grouping metric and the definition of the threshold

InFiniteLungDT addresses the ECETOC proposal directly: it measures 17 observables that **are** biological effects — uptake, redistribution between cell types, direct damage or indirect structural changes to subcellular organelles by or due to the physical presence of the material, intercellular signalling by extracellular vesicles, and recognition of damaged epithelial cells by immune cells — and it addresses life-cycle changes implicitly, since a material sampled at any stage of its life cycle is characterised by its actual biological response rather than by the properties of its pristine form.

For grouping, each material is represented by its **effective early Mode-of-Action**: the material-specific interaction couplings of section 3, aggregated into classes of observables — which functional compartment drives which — and scaled to their own uncertainty (Figure 6a). The distance between any two materials is the distance between these representations (Figure 6b) is obtained as described in SI-13; it was not optimized or fitted for the purpose of grouping, i.e. it is derived directly from experimentally determined MoA.

The distance threshold for grouping was **determined by** (i) **minimizing grouping, i.e. maximizing discrimination, across the 49 materials** of the study, and (ii) **maximizing grouping across 22 independent repeated measurements of a single reference material** over 2 years to achieve the highest robustness against experimental variability including preparation of material, its ageing, oxidation and aggregation. All the distances are normalized to this threshold, which became the unit of the distance axis in every panel of Figure 6. A **group** is defined with respect to a selected reference material, whereas every material in the group has its eMoA distance to this selected reference below 1.

### 9.3 Repeatability: the method recognises a material as itself

The distance definition from above also defines the **repeatability**: two measurements are reliable repeats of the same material if their eMoA distance is below 1. Of the 22 independent repeat measurements for the same reference material (anatase TiO2 nanotubes), 19 lie within the threshold distance of one another and form a single densely connected cluster (Figure 6e). The three outliers might indicate the upper estimation limit for the empirical false-negative rate of the procedure — the rate at which the method would decline to group a material with itself. As noted above, failing to do so might also indicate real differences originating for example in undetected ageing and oxidation, or insufficiently handled aggregation. Because the repeats span independent biological runs, the figure reports within-laboratory repeatability in exactly the units in which grouping decisions are taken.

As a result of the repeatability study described above, the threshold is now used also as a **quality-control filter on the individual measurement**: a repeat lying further than the threshold from the other repeats of the same material (for example of the benchmark material) is flagged as non-representative, and can be identified and rejected before its prediction is used.

### 9.4 Grouping across all 49 materials

From the matrix of pairwise distances (Figure 6b), the corresponding similarity map was constructed (Figure 6c), which shows that among 1176 possible pairs of the 49 materials, less than 40 material pairs (i.e. 3%) have MoA similar below the threshold (indicated by red lines in Figure 6c). These 27 materials can be placed into 9 groups (Figure 6d), while 22 remain ungrouped – an expected result for materials of such variety, and in line with conservative approach for regulatory read-across without over-grouping.

Among the grouped materials, we can identify some with previously known similarities: MWCNTN006 and MWCNT401 (grouped also by an independent IATA-based assessment^84^), TiO2N001 and TiO2N030 (reportedly two batches of the same material with very similar properties, SI-4; also serves as an independent repeatability validation), and clinker and cement (which typically contains about 70% clinker^85^). However, the presented grouping clearly **cuts across chemistry in both directions**. (i) **Chemically different materials can fall into one group**: three TiO2 variants lie in the dense central region together with cement and clinker. (ii) On the other hand, **the same chemistry does not guarantee MoA similarity**: TiO2 nanocubes, despite being derived from the very same anatase source material and having very similar XRD spectrum as TiO2 nanotubes^29,36–65^, show a clearly distinct MoA, and group with rutile TiO2N025 instead. Five nanoforms of the same substance thus distribute over two well-separated regions of the map, while three of them coincide with chemically unrelated industrial materials.

Across the whole set, no material family shows a unique or qualitatively distinct Mode-of-Action that would separate it from the rest, while chemically dissimilar materials can share one — consistent with the repeated finding that no “nanospecific” mechanism of action has been linked to a physicochemical property across in vivo and in vitro studies^77,78^.

For the 18 single- and multi-walled carbon nanotubes of the study (SI-13), the grouping identifies 5 pairs of similar materials. Nanotubes of nominally the same chemistry and morphology class are therefore not one mechanistic group. An independent, characteriser-based analysis carried out for ECHA on multi-walled carbon nanotubes reached a comparably conservative conclusion for the same material class, finding acceptable variation in 6 of 18 framework challenges^83^; the two approaches use entirely different evidence and converge on a similar degree of groupability.

### 9.5 Three regulatory scenarios, worked on the oxide subset

While similarity depends exclusively on the experimentally derived MoA, grouping depends on the choice of reference materials and the regulatory objective. To illustrate, we present three grouping scenarios (Figure 6f–h) together with their associated reference material selection, of the same 19 oxide-containing materials of the study, obtained from the same distances and the same threshold, under three different objectives:

**Maximal grouping (Figure 6f)** could be used when one wants to register all the given materials, for which every material would either require its own animal test or to be read-across against another reference material. In this scenario, 19 oxide material would fall into 3 groups comprising 12 materials, and 7 remain ungrouped (their distances to any other material are larger than the threshold). This scenario would allow all 19 materials registered relying on 10 animal studies (reference materials indicated in red) instead of 19.

**Cost-efficient grouping (Figure 6g)** allows a registrant to minimize the total cost per registered material by reducing the costs of required animal tests, selecting reference materials that represent the largest group(s). In this scenario, every material would still have been assessed with InFinite method, but the in vivo test would be run only for the selected group reference materials. At an in vitro-to-in vivo cost ratio of 1:20, the optimum is a single group of 8 members: one animal study would result in registration of eight materials, with the remaining 11 left unregistered. Each additional group adds an animal study whose cost outweighs the marginal members it brings in. The cost ratio is an explicit parameter of the optimisation, so a registrant or an authority can re-run it with their own cost structure (e.g. including expected market returns), and the ratio at which the preferred grouping changes is itself informative. This scenario can stimulate the registration of more novel materials, all of them long-term safety-assessed, which is the ultimate goal of the regulatory framework.

**Read-across from already in vivo-tested reference materials (Figure 6h)**, typically registered materials, stimulates reuse of the animal datasets by attaching new materials to these references where the distance permits. In this scenario, 2 such reference materials have been selected for demonstration, to which 6 out of 17 remaining materials could be grouped/read-across.

### 9.6 Scope of the claim

The InFiniteLungDT method enables grouping based on the observed biological effects, exactly what the DF4nanoGrouping framework calls for. While it does not suggest to replace/substitute any characterisation or the substance-identity assessment required by Annex VI; it does provide a mean to determine similarity of early Mode-of-Action between materials supporting grouping and read-across as suggested by Annex XI. For sets of similar nanoforms, which require evidence beyond Annex VI, the eMoA distance together with the predicted LOAEL/BMC and resolution time is exactly such evidence — and, because it is derived from a measured response rather than from a descriptor, it can be delivered also for materials whose descriptors cannot be determined, including the industrial real-world samples and their cocktails (section 7.3).

**MoA-based grouping enriches the single predicted endpoint.** While the digital twin predicts a limited number of endpoints per TG panel together with LOAELs, they are all derived from much richer (upstream) early Mode-of-Action. Materials close in eMoA share their predicted LOAEL/BMC (see colour coded nodes in Figure 6c,d), whereas materials with indistinguishable predicted LOAEL/BMC can be found across the whole map, mostly ungrouped: a similar Mode-of-Action implies a similar endpoint, while a similar endpoint does not necessarily imply a similar Mode-of-Action. MoA similarity between two materials is thus a stronger claim than agreement of their predicted neutrophil influx. As similar Mode-of-Action reasonably implies similarity of other outcomes delivered during a TG animal study (histopathology, lung burden, differential cell counts, etc.; not only BAL neutrophil measurement), all these endpoints need to be tested only once per group, and similarity of the early Mode-of-Actions of the tested materials within the identified group provides scientific ground for the standard read-across^79^.

## 10 Discussion

Here, we describe a new testing paradigm in nanotoxicology that addresses the pressing need in the community to optimize and validate novel animal replacement alternatives, i.e. new approach methodologies (NAMs), in nanomaterials (NMs) toxicity testing ^73^. Namely, current regulatory decision-making still relies on animal testing, which is expensive and time consuming. For example, the 2-year rat carcinogenicity study costs around $4 million and requires more than 500 animals for risk assessment of a single substance^73^. Due to high amount of new NMs being produced, it is believed that the only path forward for the toxicological community is to find alternatives that will reduce or replace animal testing.

### 10.1 Regulatory context of use – reduction, not replacement of complete TGs

InFiniteLung in vitro learned digital twin could be used in regulatory setting in the following contexts: i) for dose range-finding and for LOAEL/BMC determination for neutrophilic lung inflammation for TG 412 and 413, ii) as an IATA component, iii) for derivation of a dose– response relationship and a PoD, which can be divided by uncertainty factors to obtain health-based guidance values in line with ECHA guidance ^74^, and iv) as support for grouping, read-across and the selection of the material to be tested.

1. According to Chung (2022), the OECD TGs for inhalation toxicity testing 412 (28-day) and 413 (90-day) were recently revised to evaluate lung inflammation after rodent inhalation studies with insoluble nanomaterials. This increased the number of testing animals needed to conduct bronchoalveolar lavage (BAL) fluid analysis and nanomaterial lung burden measurements to 120 for TG 412 and 160 for TG 413 ^75^, although current guidelines allow doing BAL and lung burden and histopathology in the same animal. In addition, a typical dose range-finding study generally uses 3 to 6 males and 3 to 6 females per tested concentration, and 1 to 3 exploratory concentrations plus a control, resulting additional 12 to 48 test animals. Using InFiniteLungDT-enabled grouping, some studies may be waived, especially those not expected to cause systemic toxicity. More importantly, it provides scientifically sound data to perform the initial risk assessment for the different modifications of nanomaterials without animal testing, which have small production volume (e.g. research samples).
2. InFiniteLungDT can be employed as quantitative, dose-resolving component for the inflammation-onset and inflammation-resolution determination in an integrated approach with other methods. It pairs naturally with future NAMs covering later stages of the AOPs containing inflammatory cell influx as a key event (KE1497).
3. InFiniteLungDT predicts neutophilic lung inflammation as a percentage of neutrophil in BAL dependent on material concentration which can be used for determination of LOEAL/Benchmark Concentration (BMC). Since the InFiniteLungDT predicts BMC derived from rat inhalation studies with 84% accuracy, it’s LOAEL/BMC prediction could be used as a Point of Departure (PoD) to calculate human equivalent concentrations and health-based exposure limits. This can be used in industrial setting for pre-screening of materials available only in milligram quantities, workplace air-sample assessment, and for mixture risk assessment, since it doesn’t depend on characterization of material properties.
4. Animal inhalation studies remain necessary for endpoints this method does not deliver: histopathology and morphometry, systemic availability and distribution, and any outcome downstream of KE1497 — fibrosis, proteinosis, atherosclerosis, carcinogenicity. Sustained neutrophil influx is a precursor of several of these outcomes and a protective basis for a PoD, but a precursor is not a substitute. However, the MoA-based grouping, demonstrated in section 9, aims precisely at reducing the need for animal tests also for endpoints other than KE1497, as a similar in vitro-detected MoA lays scientific grounds to expect an overall similar in vivo response. The study that is then not performed is the complete Test Guideline study for that material, and not only its neutrophil measurement, so the reduction reaches endpoints the method itself does not predict. That grouping is performed on the early Mode-of-Action rather than on the predicted endpoint is what makes this expectation defensible, since it is the stricter of the two criteria (section 9).

### 10.2 Comparison with other approaches

**Machine-learning NAMs and QSARs** excel at interpolation and struggle with extrapolation to dissimilar substances — which are, by definition, those outside the training distribution — and they cannot readily address mixtures. Forecasting further into the future than any trajectory seen in training is not possible: even models designed for temporal dependence degrade as the prediction horizon leaves the training range. A digital twin does not extrapolate in this sense at all. It integrates equations of motion forward, and the horizon is limited by the validity of the mechanism, not by the reach of the training data. Unlike other approaches, the InFiniteLungDT can predict the dose- and time-dependent endpoint of interest for various exposure scenarios.

**Organ-on-chip and air–liquid interface (ALI) systems** are structurally more faithful, yet they do not readily provide conclusions outside the time frame of the observation without an accompanying in vitro-to-in vivo translation. They also face difficulties with reproducibility and dose quantification and maintaining stable cell hydration and viability. The digital twin based on simplified, but robust biological in vitro system is restricted to predefined biological observables which it propagates far beyond the observation window, and is thus faster, more resource-efficient and more high-throughput-compatible.

The InFiniteLungDT excels over **high-content screening and Cell Painting** in two respects: it avoids probes that interfere with cell dynamics and avoids fixation altogether; and it replaces holistic morphological “painting” with a biologically relevant, predefined, organelle-focused observable set, in order not to produce a high-dimensional dataset of uncertain informative value. The deliberate reduction of dimensionality is what keeps the number of degrees of freedom commensurate with the number of available in vivo data points (section 4.1, Premise 3).

### 10.3 The applicability domain

The applicability domain of InFiniteLungDT is limited a) experimentally by the solubility of materials, and b) by the limited toxicity MoA coverage.

a) We successfully applied the method to two highly soluble materials (50-60% in cell medium after 24 hours, SI-8) **ZnO 110 and 111** ^76^. Two things make this edge case tractable rather than disqualifying. First, the delivered dose is measured continuously by back-scatter, so the *disappearance* of particulate material during the experiment is observed (and automatically integrated into MoA) rather than assumed. Second, ion-releasing zinc oxide nanoparticles significantly affect cell organelle structure, triggering severe structural and functional damage ^77,78^, thus the ion-release component of the response enters the MoA through the same couplings as any other perturbation, without a solubility-specific model.

**The applicability domain extends to** mixtures and cocktails (material and adsorbed chemicals) are in domain provided at least one component satisfies the solubility criterion, because the method never requires the composition to be resolved. Seven combustion-generated cocktails in the validation set confirm this.

b) The method is applicable precisely where descriptor-based methods fail, namely to materials whose descriptors cannot be measured; instead, it depends on the in vitro measured biological response that is measured for every material the method is applied to. The approach would completely fail for substances triggering KE1497 via pathways that do not at all reflect in the chosen set of observables. These being rather cell organelle-level cellular responses (e.g. proliferation, material co-localisation with organelles, cell reshaping, division impediment, excretion of debris and vesicles, etc. – see Table 2), though, such complete misses are highly unlikely. The limitation that calibration materials cover 92% of the available MoA space, has been tested with validation materials, for which predictions were not substantially worse (see section 7.4).

### 10.4 Human relevance

First, we attested the human relevance of the responses in murine cells (LA-4, MH-S) in Brown et al. (2024) where we documented whole-genome duplication and associated genomic signatures in tumours from a cohort of more than 1,800 patients, and demonstrated that combustion by-products are capable of inducing whole-genome duplications in cell culture, with mutational signatures in patient tumours consistent with exposure to those carcinogens^79^. LA-4 cells under particle exposure reproduce the same phenomenon. This is a peer-reviewed correspondence between the cellular response of our murine cell platform and a phenotype documented in human patients, established at the level of the underlying cellular response rather than reproducing a model that would reproduce human morphology or all the human specific biochemical pathways. InFiniteLungDT is, at least to our knowledge, the only in vitro cell model capable of reproducing this observation in human patients.

Second, the demonstration that mouse-instillation-calibrated parameters predict rat inhalation responses (section 7.2) is another empirical evidence that the digital twin methodology can be applied across species without re-engineering in vitro cellular platform. The same approach used here can therefore be used to validate prediction of human inflammation once human reference data of comparable quality exists. The method is for inflammation prediction in the healthy lung, not for modelling COPD, asthma or other disease states.

Third, amongst the seventeen observables defined beforehand from mechanistic reasoning and availability what can be monitored live at the level of cellular functional units, nine were found to correspond to Key Events documented in AOP-Wiki for inflammation (Table 1), an independent convergence and a consistency check on the platform’s human relevance.

Finally, the InFiniteLungDT’s prediction for human effects would take into account different lung surface area and breathing parameters (Table 2). As solid data on interspecies variation of other influencing systemic factors become available (e.g. resolution kinetics, material clearance, macrophage turnover, neutrophil half-life, surfactant/corona), these can be integrated into the model to further improve human applicability.

## 11 Conclusions

Here we describe a testing paradigm in nanotoxicology that addresses the community’s need to optimise and validate animal replacement alternatives for (chronic) toxicity testing of substances^73^. Until now, there has been no animal-free alternative method for predicting chronic inflammation that can estimate the toxicant concentration, the timing of onset and the duration of inflammation, as required by regulatory agencies. “Increased recruitment of inflammatory cells” (KE1497) is a frequently reported event following exposure to materials^80^; it is routinely measured in vivo, with detailed guidance in OECD TG 412 and 413, and until now it could not be assessed in quantitative manner in vitro^73^.

We developed the in-vitro-learned digital twin InFiniteLungDT, which predicts KE1497 as accumulation of neutrophils in the lung, together with its dose dependence and its resolution up to chronic condition. We validated the prediction against the largest available consistently acquired in vivo instillation dataset — 49 materials, 3,094 mouse exposures —, and against rat inhalation data acquired in compliance with OECD TG 412 for 7 materials, 364 rat exposures, contributed by three partner laboratories. The predictive capacity of LOAEL/BMC determination is **92% (R² = 0.73) for instillation** and **84% (R² = 0.63) for inhalation**; the correlation between predicted and in-vivo-determined LOAELs/BMCs lies at the upper limit of the expected range for NAMs given the variability of in vivo data (R² = 0.55–0.73, ToxRefDB)^81^. Predictions are 1.5 times more repeatable than the animal experiments they are compared against. The 15 calibration materials, selected for diversity of endpoint kinetics and of triggered Mode-of-Action, cover 92% of the Key Event Relationships permitted by the model. The remaining 34 validation materials show similar prediction accuracy, even though some of them stretch the MoA beyond the calibrated coverage — the basis on which the method is expected to apply to materials outside its calibration set. We confirmed this on 7 industrial samples, collected on-site at milligram quantities and composed of mixture of materials, whose intrinsic properties cannot be characterised. A detailed SOP has been developed and its transferability exercised between two laboratories (data not shown).

On the basis of the demonstrated capability, the following abilities define the context of use for regulatory risk assessment of inhalation toxicity:

1. determination of a quantitative dose–response relationship for benchmark dose determination, usable as a Point of Departure that can be divided by uncertainty factors to obtain health-based guidance values in line with EFSA guidance^82^ — a PoD derived from chronic neutrophilic lung inflammation is protective for human risk assessment because it precedes more severe adverse outcomes;
2. prediction of the apical endpoint of regulatory relevance (neutrophilic lung inflammation as KE1497), while downstream morphometric and pathophysiological outcomes remain outside the predictive scope;
3. prediction of responses for different exposure scenarios (single and continuous/repeated);
4. prediction of responses to an unknown, not previously characterised material, without knowledge of its intrinsic properties;
5. prediction of responses to mixtures of poorly water-soluble substances whose intrinsic physicochemical properties are difficult to characterise meaningfully;
6. prediction of neutrophilic lung inflammation as a post-exposure observation aligned with OECD TG 412 and TG 413, and beyond (e.g. TG 452);
7. prediction of the early Mode-of-Action, including early biological effects and molecular initiating events which, for many stressors such as nanomaterials and physical agents, which are not classical molecule–receptor interactions; and
8. grouping of materials by their measured biological effects rather than by their intrinsic properties, against a threshold fixed by the repeatability of the method itself, so that one representative material per mechanistic group is carried into animal testing — sparing the other members the complete Test Guideline study and not only the one endpoint the method predicts — or an already tested material serves as the read-across reference for the others, while the same criterion discriminates between forms of one substance.

With automated monitoring of up to six samples per day per microscope and automated translation into an inflammation-relevant Mode-of-Action, the material-specific potency for triggering long-term lung inflammation is delivered in **less than a week** after receipt of the sample, for effects up to 6–9 months (internal numerical stability tests conducted up to 2 years) ahead of time. This method can significantly reduce the usage of test animals per material across the relevant Test Guidelines. Before legal recognition, InFiniteLungDT can already reduce animal use through faster and more resource-efficient range-finding and post-exposure-timing studies, use as an IATA component, grouping and differentiation of forms of materials, and pre-screening where only small sample quantities are available – i.e. sampled in the environment, or early in the R&D phase as part of the voluntary Safe-and-Sustainable-by-Design (SSbD) framework.

## Supporting information

inhalation dose-dependent plots

inhalation time-dependent plots

instilation dose-dependent plots

instilation time-dependent plots

## 12 Acknowledgements

The authors thank Joakim Pagels (Lund University) and Andreas Holländer (Fraunhofer Institute for Applied Polymer Research) for providing materials used in this study; Tomaž Ličina and Nika Kozoderec (both Infinite LCC), and Eva Cerkvenik, Cody Tripp (JSI) for technical support and help in transferability study.

## 13 Author contributions

### Project conceptualisation

Janez Štrancar, Tilen Koklič, and Iztok Urbančič generated the initial overarching ideas or formulated the research goals, provided high-level guidance, and institutional resources.

### Pre-validation supervision

Ulla Birgitte Vogel and Tobias Stöger guided the regulatory-relevant pre-validation process

### Experimental Design

Janez Štrancar, Ulla Birgitte Vogel, Tobias Stöger, Iztok Urbančič, Tilen Koklič, Hana Kokot, Boštjan Kokot, Tomasz Kolodziej, Christina Isaxon, Tomaž Vuk, Laurent Gaté, and Robert Landsiedel structured the specific workflows, methods, or experimental milestones.

### Material and data – acquisition and analysis

Hana Kokot, Boštjan Kokot, and Tomasz Kolodziej conducted analysis using confocal live cell in vitro microscopy; Markus Rehberg & Tobias Stöger team members (LDH, dsDNA in vitro cell assays); Pernille Høgh Danielsen, Keld Alstrup Jensen, Ulla Birgitte Vogel and Markus Rehberg & Tobias Stöger team members conducted in vivo intratracheal mice instillation experiments; Lan Ma-Hock, Laurent Gaté, Fréderic Cosnier, Sarah Valentino, and Carole Seidel conducted in vivo rat inhalation experiments; Christina Isaxon, Mateja Čubej Gašparin, and Tereza Pahor sampled and characterized the industrial materials.

### Computational Analysis

Janez Štrancar and Iztok Urbančič built theoretical models or ran digital twin (DT) calculations.

**Manuscript writing**: Janez Štrancar, Iztok Urbančič, and Tilen Koklič drafted the core text,

**Manuscript editing**: Ulla Birgitte Vogel, Lan Ma-Hock, Robert Landsiedel, Tobias Stöger, Markus Rehberg, Andrea Haase, Keld Alstrup Jensen, and Laurent Gaté substantively revised the scientific narrative.

## Declaration of generative AI and AI-assisted technologies

During the preparation of this work the authors used Claude (Anthropic) to assist with drafting and editing of parts of the manuscript. All underlying material — experimental data, analyses, figures and their interpretation — was produced by the authors, who supplied it to the tool, directed its use throughout, and reviewed and edited all resulting text. No generative AI was used to generate, analyse or interpret data, or to produce figures. The authors verified every factual statement and reference and take full responsibility for the content of the publication.

