## Supplementary information for "InFiniteLungDT, an in-vitro-learned digital twin for the prediction of material-triggered chronic neutrophilic lung inflammation: mechanism-anchored pre-validation across materials, species and exposure scenarios"

#### Abbreviations and terminology

- InFiniteLungDT – in-*vitro*-learned digital twin for predicting long-term time-evolved dose-dependent neutrophilic lung inflammation triggered by inhaled poorly-water-soluble substances
- EPI – lung epithelial cells - Murine epithelial lung tissue cell line (LA-4), such as ATCC cat.nr. CCL-196
- IMU – immune cells such as lung macrophages - murine alveolar lung macrophages (MH-S), such as ATCC cat.nr. CRL-2019
- TUB – microtubular / tubulin transport systems
- ACT – cytoskeleton actin-based system
- ER – endoplasmic reticulum
- PNU – perinuclear region with ER and ribosomes
- EV – extracellular vesicles
- MoA – Mode-of-Action in terms of functional units - subcellular structures and cell-cell interactions; not to be confused with molecular-level Mechanism-of-Action
- LOAEL/C – lowest-observed-adverse-effect-level/concentration
- BMD/C – benchmark dose (threshold dose/concentration)
- AOP – adverse outcome pathway
- MIE – Molecular Initiating Event, a MIE is a direct interaction between a stressor (material) and a molecular target (e.g., direct damage of cellular organelle, such as plasma membrane, a vital, semipermeable barrier enclosing all cells, separating the internal cytoplasm from the external environment, for example lipid wrapping of a nanoparticle[1])
- eAOP – early adverse outcome pathway, sequency of biological events (subcellular structures and cell-cell interactions) leading from MIE to a KE, for example KE1497
- KE – Key Event
- KE1497 – Key Event - recruitment of inflammatory cells, endorsed by WPHA/WNT as part of AOP173 listed in AOP-Wiki, and relevant for assessing risk to human health
- KER – Key Event Relationship, scientifically supported causal link between two "Key Events" (KEs) in an Adverse Outcome Pathway (AOP), showing how one measurable biological event (upstream KE) triggers or leads to the next (downstream KE), helping to build a causal chain from a chemical's initial interaction (Molecular Initiating Event, MIE) to a final adverse outcome (AO). Key event relationships (KERs) explain the causal linkages between KEs and form the basis of using AOPs for predictive toxicology [2]
- OECD – Organization for Economic Co-operation and Development
- WPHA – OECD Working Party on Hazard Assessment
- WNT – Working Group of the National Coordinators for the Test Guidelines Program
- TG – Test Guideline
- NAM – New Approach Methodology
- SOP – Standard Operation Procedure
- JRC – European Commission's Joint Research Centre
- DA – Defined Approach, a formalized decision-making approach consisting of a fixed data interpretation procedure used to interpret data from a defined set of information elements
- AOP – Adverse Outcome Pathway
- AOP-Wiki – Collaborative Adverse Outcome Pathway encyclopedia developed in the context of projects that have been accepted for inclusion in the OECD Workplan
- GIVIMP – Guidance Document on Good In Vitro Method Practices

- Prevalidation (Preliminary validation) – the first stages of the validation process. Ideally the optimization and characterization of the test method have been completed, and transferability is established, and within-lab variability is assessed [3]
- Ring Trial – Also termed e.g., “inter laboratory comparison”, “ring-study” or “round-robin”. Perform the same study, using the same test materials (blind-coded) according to the same protocol
- Within-laboratory variability or intra-laboratory variability – How well a test result is reproduced in the same lab using the same equipment based on repetitive testing
- Transferability – evaluation of the within and between laboratory reproducibility
- Between-laboratory variability – or inter-laboratory variability. How well a test result is reproduced between laboratories based on repetitive testing in a ring trial
- Reliability – within-laboratory reproducibility (WLR) and between-laboratory reproducibility (BLR)
- Accuracy – how often a model gets it right (correct predictions/total), the proportion of total predictions that were correct (True Positives + True Negatives / Total Predictions).
- Predictivity, Predictive capacity, or predictive accuracy – how well a prediction model predicts **new, unseen data** - a reference outcome (effects in humans and/or animals). A model can have high accuracy on training but low predictive capacity if it's overfit (only good on known data). A model with good predictive capacity has high accuracy on new data
- Predictive model **calibration** – adjusting a model’s output to achieve the highest accuracy on training data. The training (**calibration**) data is used to define the model
- Predictive model **validation** – evaluating predictive model’s performance – assessing performance on unseen data with metrics (accuracy, $R^2$) to check for overfitting, ensuring it generalizes well beyond training, and confirming its real-world predictive power [4].

Why does InFiniteLungDT biological platform comprise mouse epithelial and immune cell lines? ... 91
Why are cell lines in InFinite biological platform grown at sub-confluent conditions? ... 91
Is InFinite biological platform relevant to human health? ... 92
Why have we selected multi-channel bi-modal live cell microscopy? ... 92
Why is digital twin superior to pattern-matching AI approach? ... 93
Single versus continuous exposure – why were data from mice instillation experiments instead of rodent inhalation experiments used for the creation of the digital twin? ... 94
Why were the 44 nanomaterials selected for the creation of the InFiniteLung digital twin? 94
Reliability of the prediction regarding the nanomaterial assay interference ... 95
SI-17 In vitro measurements and time dependent quantifications of biological observables for each nanomaterial used in calibration and validation of InFiniteLungDT ... 96
References ... 97

#### SI-1 Standard operating procedure (SOP) (layer 1 with details)

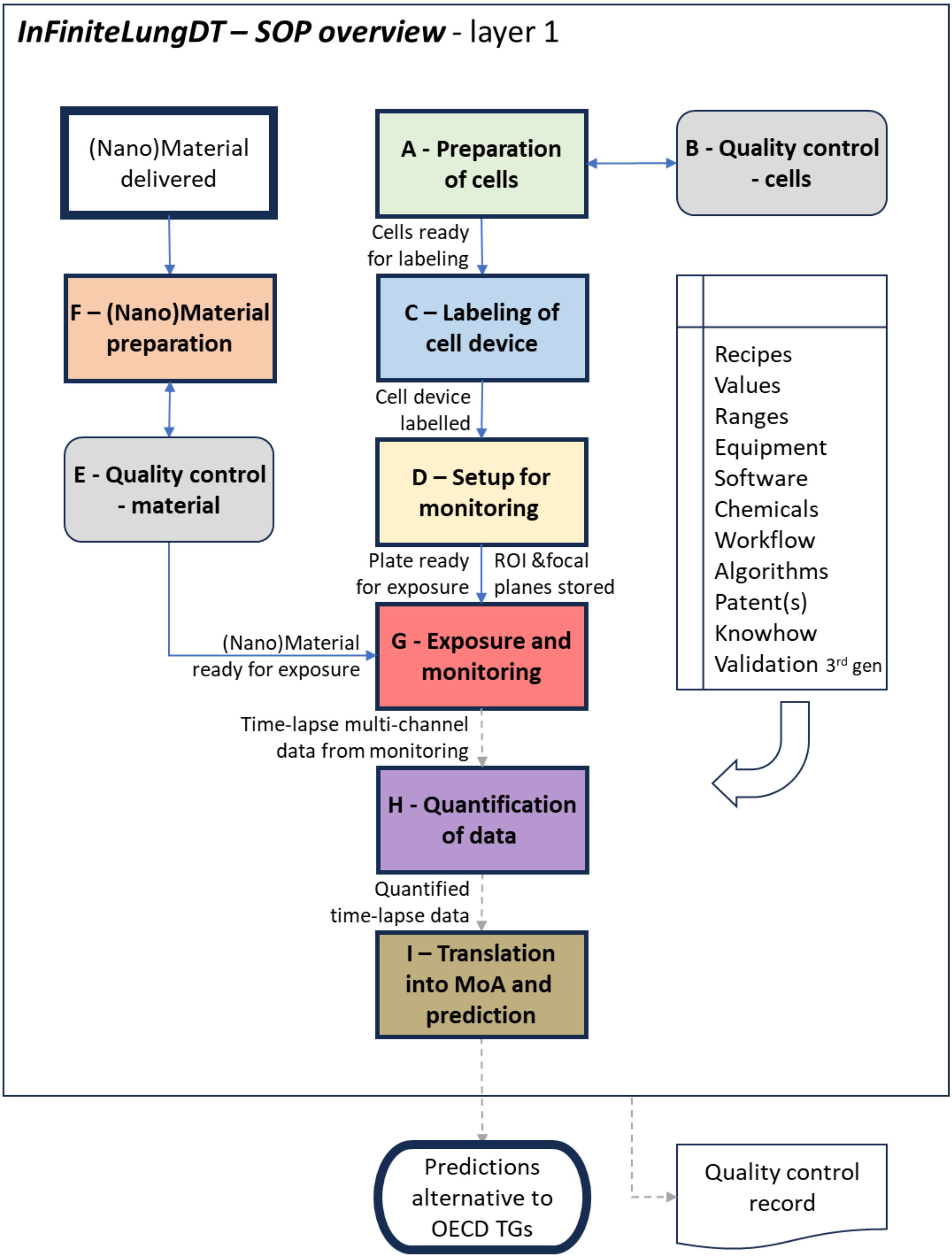

*FigureS 1.* ***SOP overview.*** *In finite platform at a glance – from in vitro monitoring to a long-term prediction within a digital twin (steps in gray boxes), that gains its relevance through Calibration and validation step, as explained in this document (box with the yellow glow).*

#### Cell cultures maintenance & quality checks

**Aim: to provide reliable and reproducible high-quality biological material**

**Key elements:**

- **Cell type selection**: mice alveolar epithelium LA-4 and mice alveolar macrophages MH-S
- **Cell culturing:** according to the good cell culture practices (GCCP), subculturing at confluency levels of 70-100% for LA-4 and 60-80% for MH-S
- **Microbial contamination testing:** at every subculturing.
- **Mycoplasma test:** at predefined intervals, at least during cell bank preparation
- **Morphology test:** according to cell line characteristics, focusing on
  - Cell shape
  - Arrangement/clustering with other cells
  - Spread on the surface
  - Adherence
  - Specific criteria for particular cell line
- **Maximum passage number:** 20
- **Banking procedure**: Cells from original vial, received from cell culture supplier, are placed in culture and harvested when they reach recommended confluency. Cells are then frozen according to standard cryopreservation method using DMSO and serum as cryoprotectants at slow cooling rate (-1 °C/minute). Cell bank usually consists of 10 to 20 vials of 1 mL, each usually containing 1-5 million cells. Stocks are stored in vapor phase of liquid nitrogen in cryostorage vessel.
- **Label uptake test:** ability of cells to be fluorescently labelled, tested for every service
- **Nanoquarantining test**: Standard response to a benchmark material with known MoA (see Kokot et al. Adv.Mat.2020[5]) to search for biologically covered material aggregates

**Workflow: Block A (7 steps) + Block B (11 steps)**

**Result:**

- Cell lines' general biological compliance
- Cell lines' compliance with InFiniteLungDT method

#### Cell device preparation

**Aim: to create a cell device**

**Key elements:**

- **Cell seeding for microscopy / monitoring:**
  - Cells are seeded in 96-well plate in central 60 wells, excluding the border wells
  - Cells are seeded according to scenaria: 2x monocultures + 1 coculture
  - Cells are seeded at lower cell confluency levels to maintain responsiveness of cell groups and to guarantee that confluency levels will not exceed the supplier's levels anytime during monitoring, with starting confluency levels

being below 70% for LA-4 and below 10% for MH-S with the ratio of LA-4 to MH-S cell areas (in co-cultures) being in the range 1:30 to 1:6
    - Cell confluency is automatically quantified before microscopy imaging to select the most suitable tiles (Field-of-Views) for imaging as well as during image quantification to feed the digital twin creation process with the actual cell confluency levels that were monitored for responses
- **Cell labelling:**
    - Labelling enables direct discrimination and identification of
        - cell type identity (LA-4 vs MH-S), intracellular and extracellular compartments
        - membranes (plasma membranes, endoplasmic reticulum)
        - transport fibres (tubulin)

      as well as indirect assessment of
        - shape of cytoskeleton (actin related)
        - state of nucleation (mono-, bi-, multinucleation, nucleus related)
    - All labels are selected as live-cell-imaging compatible and multi-channel imaging compatible labels meaning that they
        - do not interfere significantly with cell functions during monitoring interval, guaranteeing (but not limited to) undisrupted cell integrity, cell division, cell growth, cell motility, cell taxis, cell transcription and expression processes, cell communication, and cell contact
        - do not biodegrade significantly
        - do not require photoexcitation at phototoxic light current densities or wavelengths
        - do not change spectral properties after binding or partitioning
        - do allow unambiguous image unmixing (image-to-channels decomposition)
- **Labels used:**
    - CellTracker™ Green CMFDA (ThermoFischer Scientific, *Cat No. C2925*) @ 7.5 µM
    - Cell Tracker™ Orange CMRA (ThermoFischer Scientific, *Cat No. C34551*) @ 10 µM
    - Abberior LIVE 550 Tubulin (Abberior, *Cat. No. LV550*) @ 100 nM
    - CellMask Deep Red (ThermoFischer, Cat. No. C10046) @ 0.5 µg/mL
- **Labelling check of each well within the sample:** verification whether fluorescent labels were properly applied to each well within the plate

**Workflow: Block C (8 steps)**

**Result:**
- Labelled cell device
- Verified cell labelling

#### Nanomaterial preparation & QC

**Aim: To provide well-characterised dispersion of material**

**Key elements:**
- **Preparation of stable material dispersion:**
    - Dispersing the particulates of material in buffer such as bicarbonate buffer by sonication complying with nanohazard protection measures
    - For poorly-dispersible materials, a biologically compatible surfactant is used

- Stability is checked via assessing aggregation speed.
- **Characterisation & Quality checks:**
  - Direct concentration of material measured by QCM
  - Suspension stability – lack of sedimentation
  - Material aggregation – measured via backscattered microscopy, optionally SEM
  - Microbial contamination test

**Workflow: Block E (9 steps) + Block F (10 steps)**

**Result:**

- Well-dispersed nanomaterial, ready to be tested

#### Exposing cell device to materials, microscopy observation

**Aim: To provide proper material testing environment**

**Key elements:**

- **Monitoring run:** control, benchmark material, up to 5 tested materials, each monitoring run uses different biological replicate
- **Material testing conditions** – cell devices (prepared according to block of steps (A, B & C) exposed with each material (prepared according to block of steps E & F) monitored for 24 hours with 2 subsequent monitoring runs representing 2 biological replicates each comprising 3 technical replicates
- **Control:** cell device (prepared according to block of steps (A, B & C) exposed to buffer only (without material) monitored for 24 hours in 2 technical replicates for each monitoring run
- **Benchmark material:** cell device (prepared according to block of steps (A, B & C) exposed to dispersion of $TiO_2$ nanotubes (prepared according to block of steps E & F) monitored for 24 hours in 3 technical replicates, used to quality test the entire methodology (cell devices, microscopy setup, quantification, digital twin creation and prediction) including dynamic range of all responses, replacing positive control, relying on known & published MoA (*see: Kokot et al. Adv.Mat.2020*[5]), exhibiting confirmed significant responses in acute, subacute and subchronic postexposure times even after single-dose exposure
- **Dosing:** Optimization of material dose for in-vitro monitoring with respect to local material-surface-to-cell-surface ratio (targeting ratio around 1)
- **Microscopy preparation phase** involves:
  - Stabilization of environmental conditions enabling further cell growth (temperature, $CO_2$ and humidity)
  - Tile selection according to optimal cell confluency level and according to optimal cell type ratio (in case of co-cultures)
  - Focal plane(s) determination
  - Laser power selection
- **Automated microscopy** run for 24 hours in 4 physical channels: one back-scattered channel (at 488 nm) and 3 fluorescent channels (excitation at laser wavelengths of 488nm, 561nm and 640 nm):

- In the back-scattered channel, the material in tracked in label-free way delivering the mask of material for every image;
- In the first fluorescent channel, cell tracker is used to detect interior of epithelial cells;
- In the second fluorescent channel, second cell tracker is used to detect interior of immune cells together with tubulin label that is used to detect tubulin fibres in both type of cells;
- In the third fluorescent channel, membrane probe (cell mask) is used to localize all membrane-based cellular structures.

**Workflow: Block D (11 steps) + G (6 steps)**

**Result:**

- Cell device exposed to material suspensions
- Time lapsed 4-channel images of exposed/control cells, for every replicate/material

#### SI-2a Time-course and dose-response predictions for instillation exposure versus in vivo (mice) data – all 49 materials

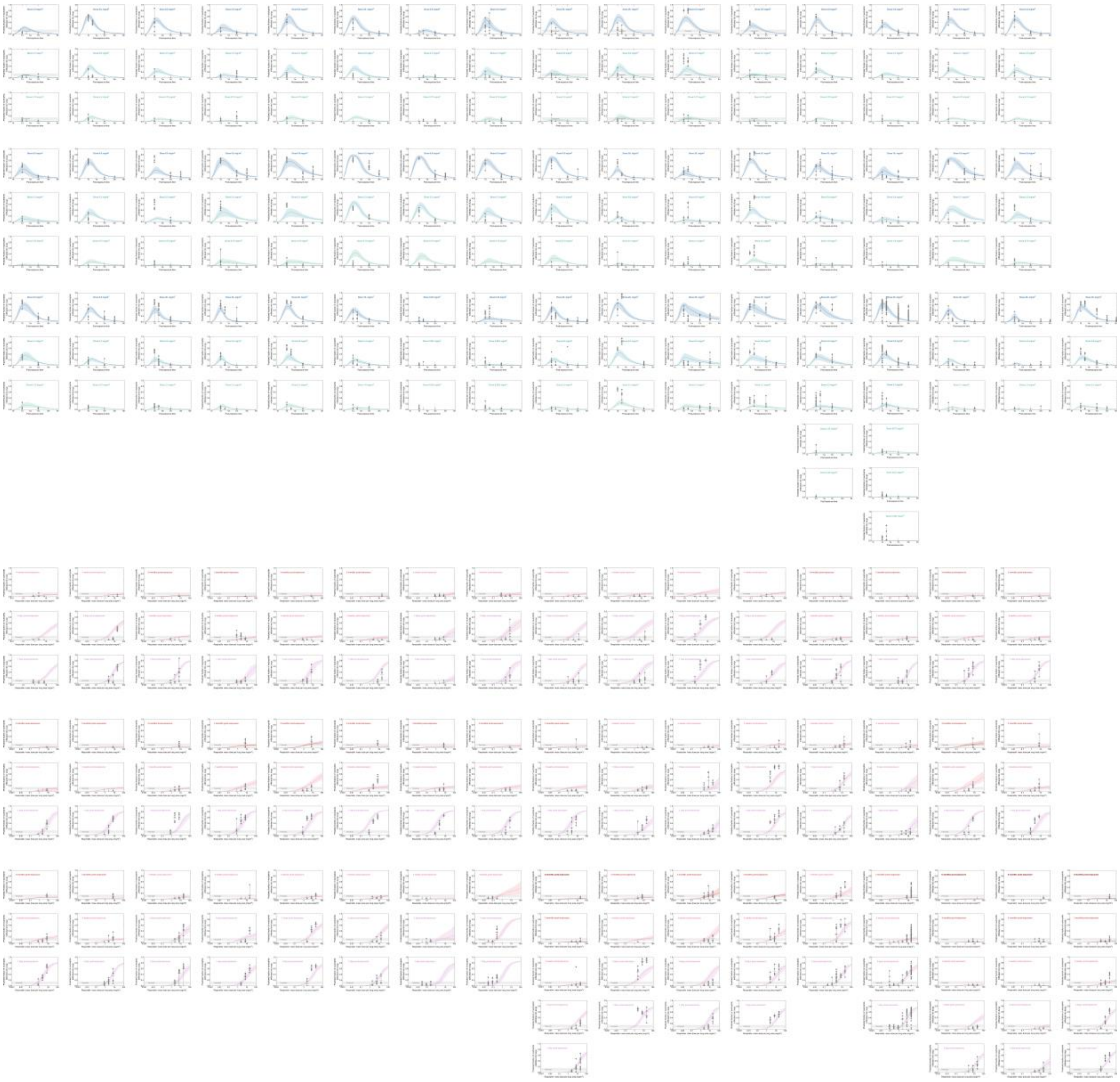

*FigureS 2. Predictions of time dependent (blue lines) and dose dependent (red lines) neutrophil influx versus instillation exposure in vivo (mice) data points for all 49 individual materials with all in vivo data points (grey circles). For higher resolution images see supplemental files: instilation_time-dependent_HiRes_plots.zip, and instilation_dose-dependent_HiRes_plots.zip.*

### SI-2b Time-course and dose-response predictions for instillation exposure versus in vivo (mice) data – per material

##### Carbon 1

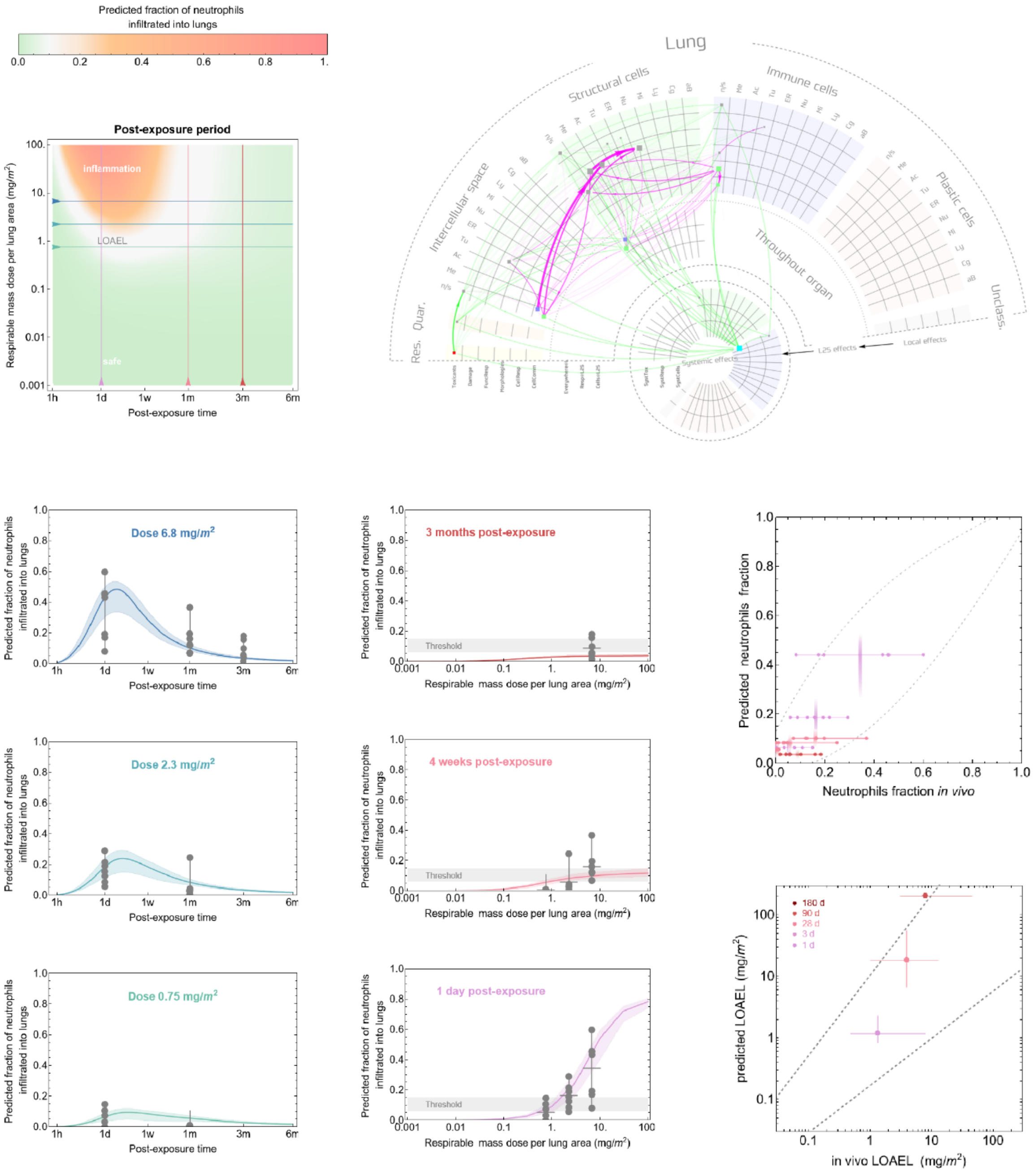

### Carbon 2

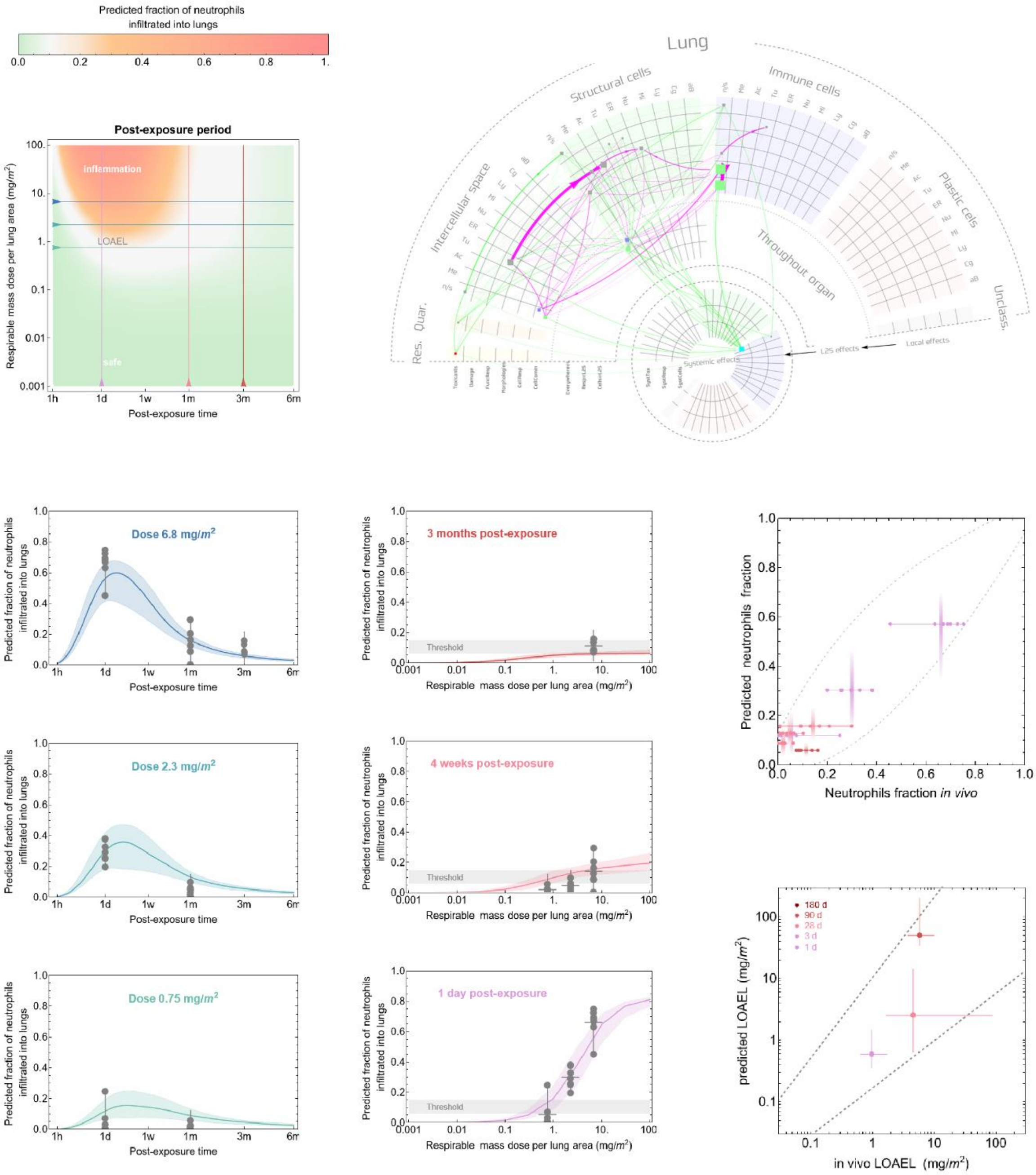

Predicted fraction of neutrophils infiltrated into lungs
Post-exposure period
inflammation
LOAEL
safe
Respirable mass dose per lung area (mg/m²)
Post-exposure time
Lung
Structural cells
Immune cells
Intercellular space
Plastic cells
Throughout organ
Res. Quar.
Unclass.
Systemic effects
L2S effects
Local effects
Dose 6.8 mg/m²
Dose 2.3 mg/m²
Dose 0.75 mg/m²
3 months post-exposure
4 weeks post-exposure
1 day post-exposure
Threshold
Predicted neutrophils fraction
Neutrophils fraction in vivo
180 d
90 d
28 d
3 d
1 d
predicted LOAEL (mg/m²)
in vivo LOAEL (mg/m²)

#### Carbon 3

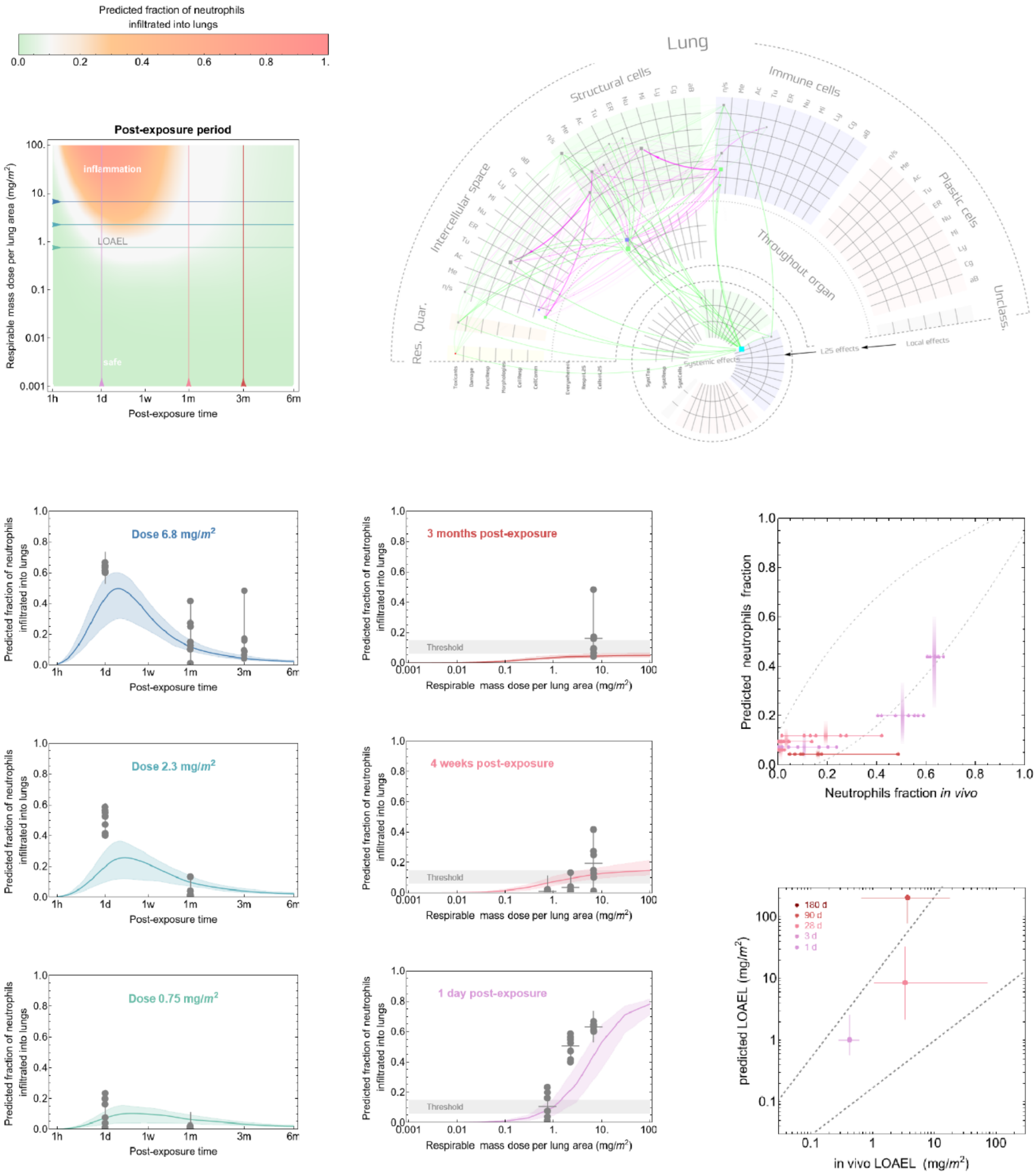

Predicted fraction of neutrophils infiltrated into lungs
0.0 0.2 0.4 0.6 0.8 1.
Post-exposure period
inflammation
LOAEL
safe
Respirable mass dose per lung area (mg/m²)
Post-exposure time
1h 1d 1w 1m 3m 6m
Lung
Structural cells
Immune cells
Plastic cells
Intercellular space
Throughout organ
Res.
Quar.
Unclass.
Systemic effects
L2S effects
Local effects
Dose 6.8 mg/m²
Dose 2.3 mg/m²
Dose 0.75 mg/m²
Predicted fraction of neutrophils infiltrated into lungs
3 months post-exposure
4 weeks post-exposure
1 day post-exposure
Threshold
Respirable mass dose per lung area (mg/m²)
Predicted neutrophils fraction
Neutrophils fraction in vivo
180 d
90 d
28 d
3 d
1 d
predicted LOAEL (mg/m²)
in vivo LOAEL (mg/m²)

#### Carbon 4

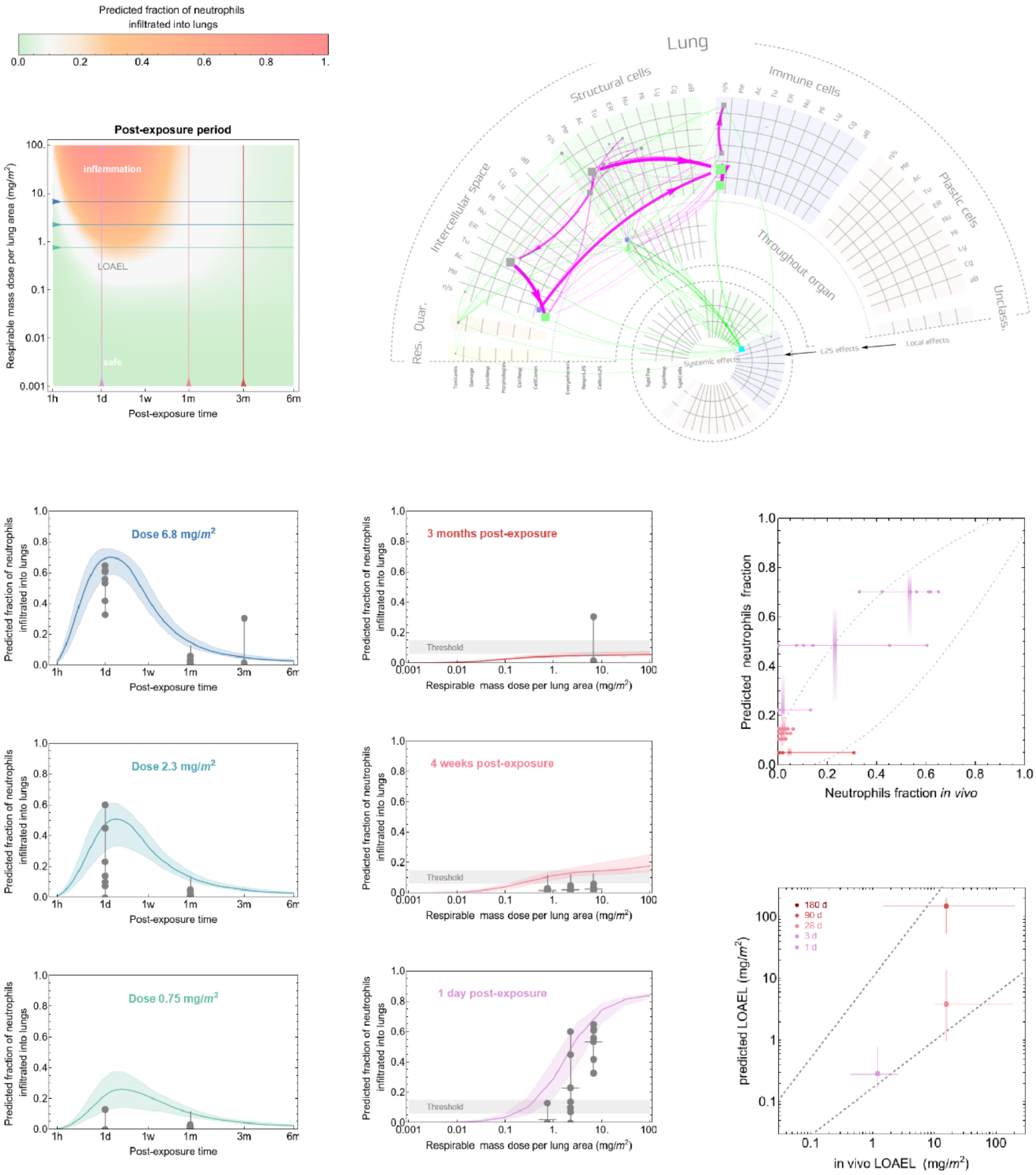

Predicted fraction of neutrophils infiltrated into lungs
0.0 0.2 0.4 0.6 0.8 1.
Post-exposure period
inflammation
LOAEL
safe
Respirable mass dose per lung area (mg/m²)
Post-exposure time
1h 1d 1w 1m 3m 6m
Lung
Structural cells
Immune cells
Intercellular space
Throughout organ
Plastic cells
Unclass.
Res.
Quar.
Systemic effects
L2S effects
Local effects
Dose 6.8 mg/m²
Dose 2.3 mg/m²
Dose 0.75 mg/m²
3 months post-exposure
4 weeks post-exposure
1 day post-exposure
Threshold
Predicted fraction of neutrophils infiltrated into lungs
Respirable mass dose per lung area (mg/m²)
Predicted neutrophils fraction
Neutrophils fraction in vivo
180 d
90 d
28 d
3 d
1 d
predicted LOAEL (mg/m²)
in vivo LOAEL (mg/m²)

#### Carbon 5

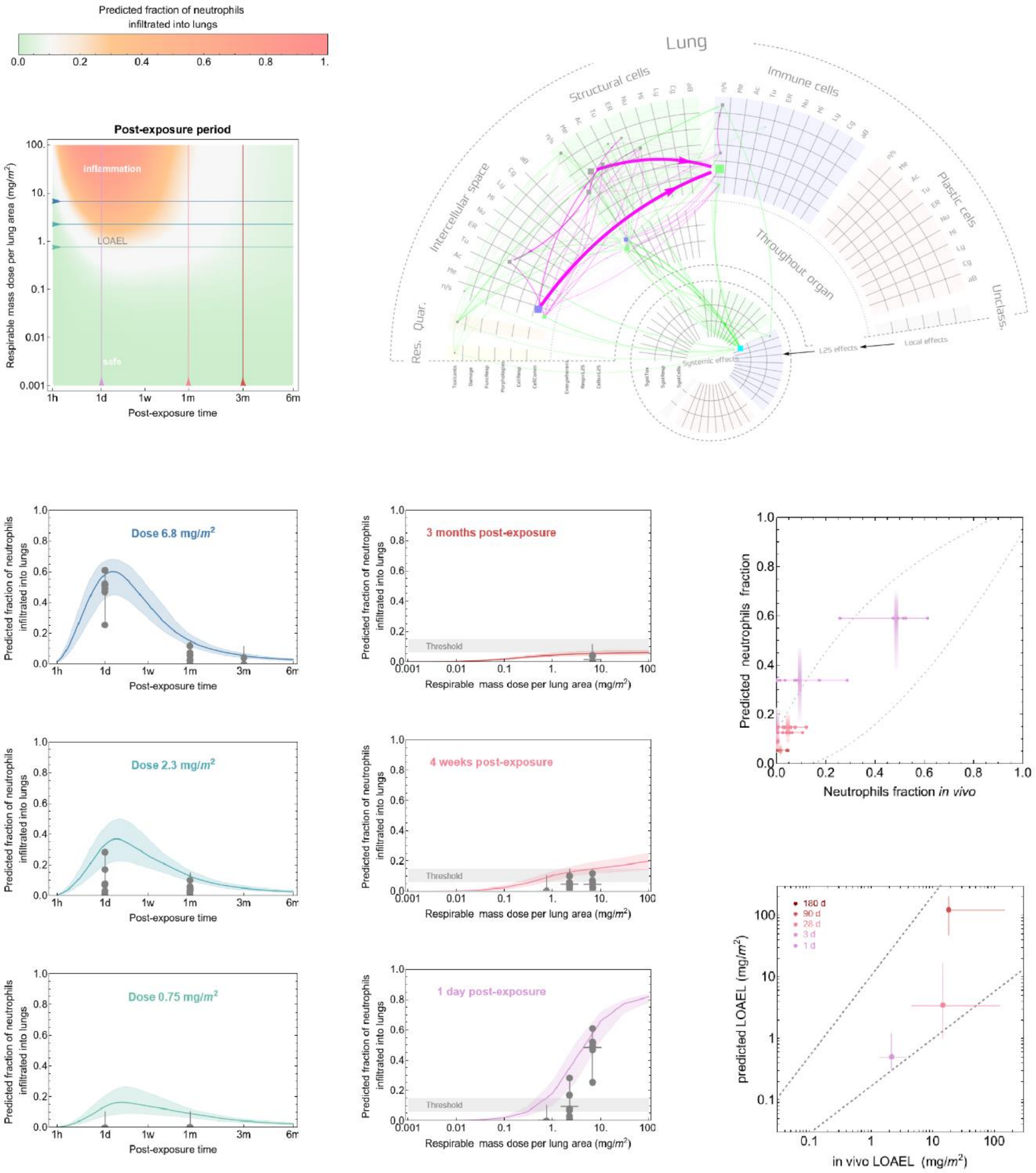
Predicted fraction of neutrophils infiltrated into lungs
Post-exposure period
inflammation
LOAEL
safe
Respirable mass dose per lung area (mg/m²)
Post-exposure time
Lung
Structural cells
Immune cells
Intercellular space
Plastic cells
Throughout organ
Res.
Quar.
Unclass.
Systemic effects
L2S effects
Local effects
Dose 6.8 mg/m²
Dose 2.3 mg/m²
Dose 0.75 mg/m²
3 months post-exposure
4 weeks post-exposure
1 day post-exposure
Threshold
Predicted neutrophils fraction
Neutrophils fraction in vivo
180 d
90 d
28 d
3 d
1 d
predicted LOAEL (mg/m²)
in vivo LOAEL (mg/m²)

**Carbon 6**

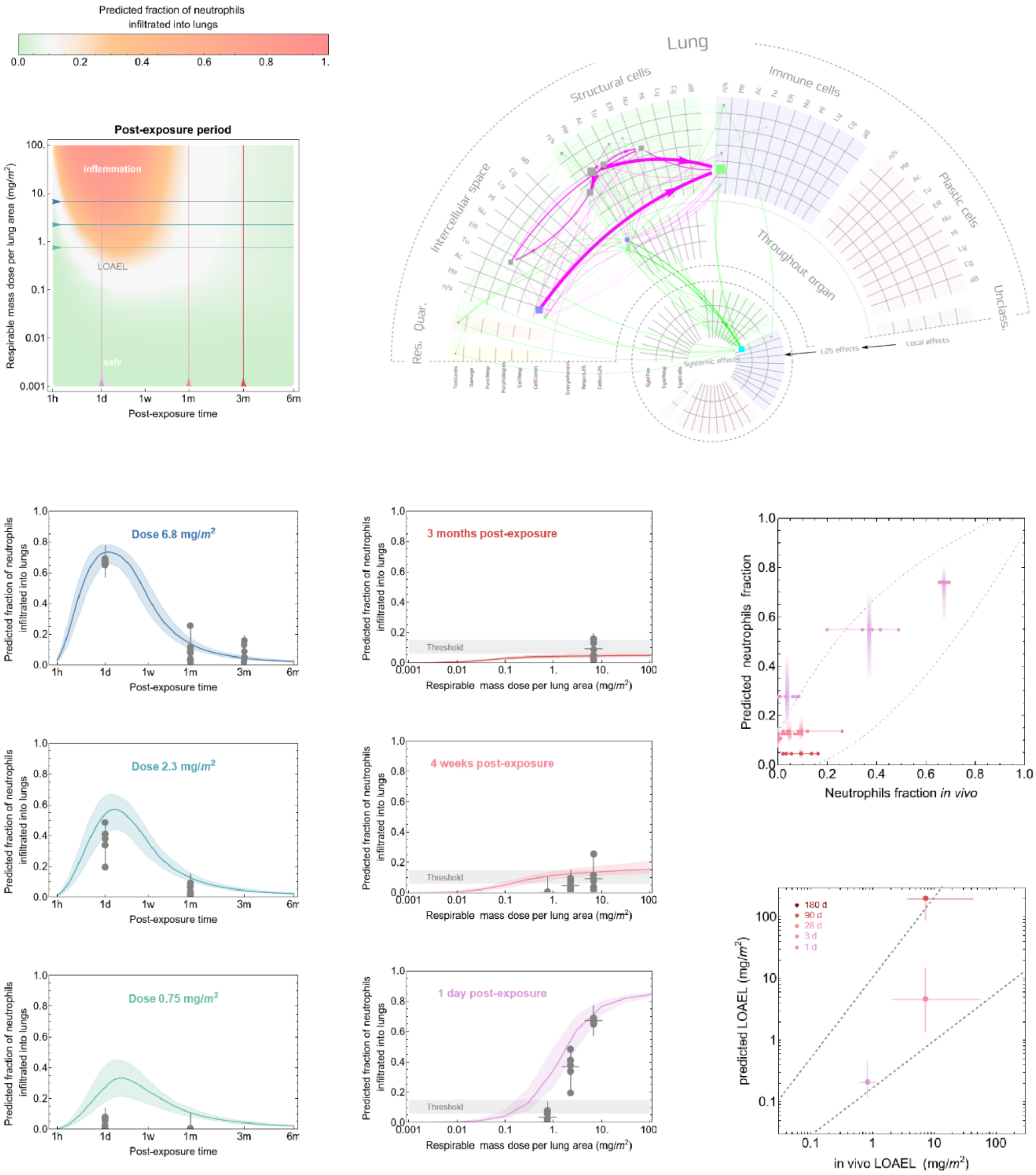

Predicted fraction of neutrophils infiltrated into lungs
Post-exposure period
inflammation
LOAEL
safe
Respirable mass dose per lung area (mg/m²)
Post-exposure time
Lung
Structural cells
Immune cells
Intercellular space
Plastic cells
Throughout organ
Res.
Quar.
Unclass.
Systemic effects
L2S effects
Local effects
Dose 6.8 mg/m²
Dose 2.3 mg/m²
Dose 0.75 mg/m²
3 months post-exposure
4 weeks post-exposure
1 day post-exposure
Threshold
Predicted fraction of neutrophils infiltrated into lungs
Respirable mass dose per lung area (mg/m²)
Predicted neutrophils fraction
Neutrophils fraction in vivo
180 d
90 d
28 d
3 d
1 d
predicted LOAEL (mg/m²)
in vivo LOAEL (mg/m²)

#### Carbon 7

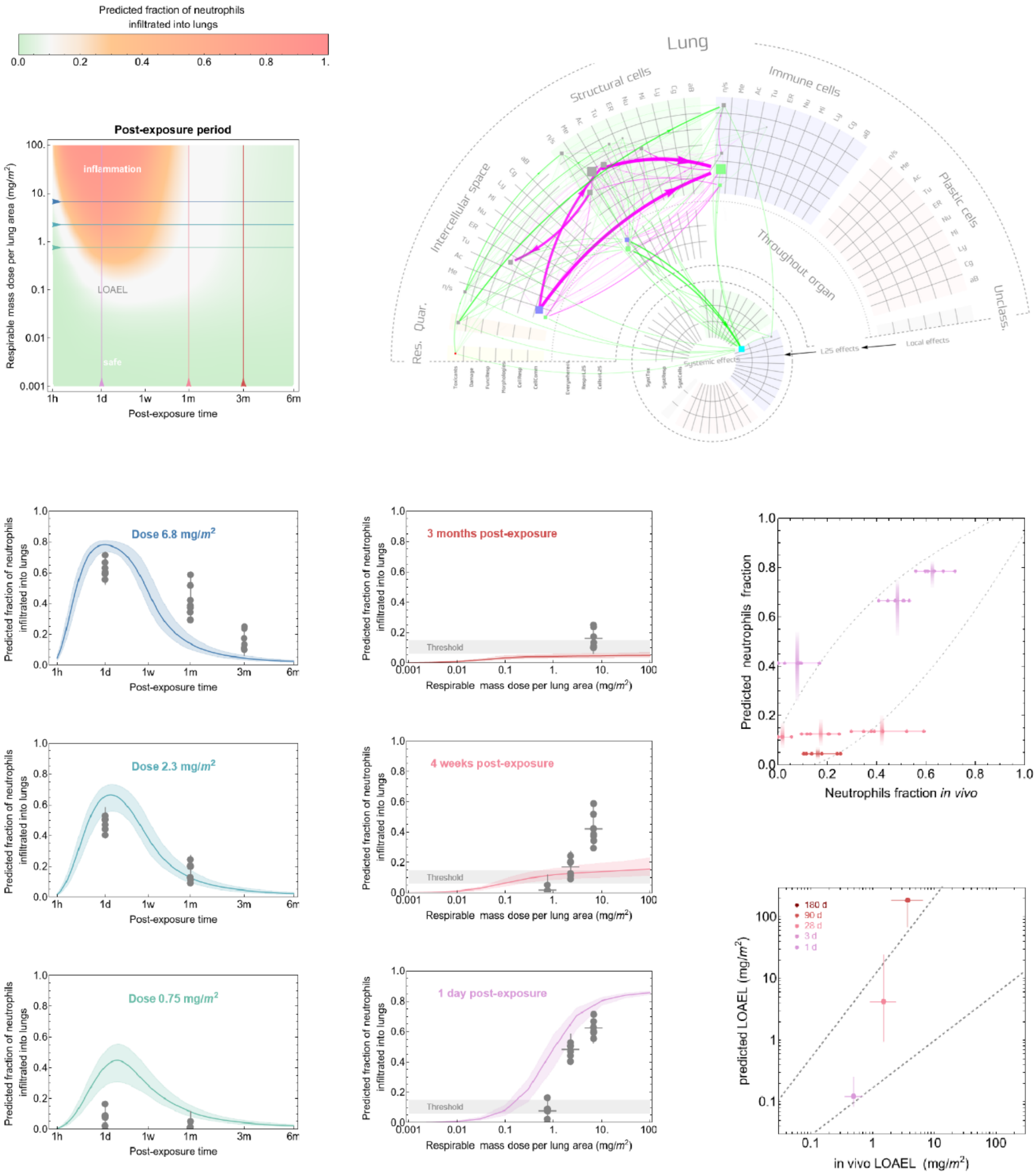
Predicted fraction of neutrophils infiltrated into lungs
Post-exposure period
inflammation
LOAEL
safe
Respirable mass dose per lung area (mg/m²)
Post-exposure time
Lung
Structural cells
Immune cells
Intercellular space
Throughout organ
Plastic cells
Res.
Quar.
Unclass.
Systemic effects
L2S effects
Local effects
Dose 6.8 mg/m²
Dose 2.3 mg/m²
Dose 0.75 mg/m²
3 months post-exposure
4 weeks post-exposure
1 day post-exposure
Threshold
Predicted neutrophils fraction
Neutrophils fraction in vivo
180 d
90 d
28 d
3 d
1 d
predicted LOAEL (mg/m²)
in vivo LOAEL (mg/m²)

### Carbon 8

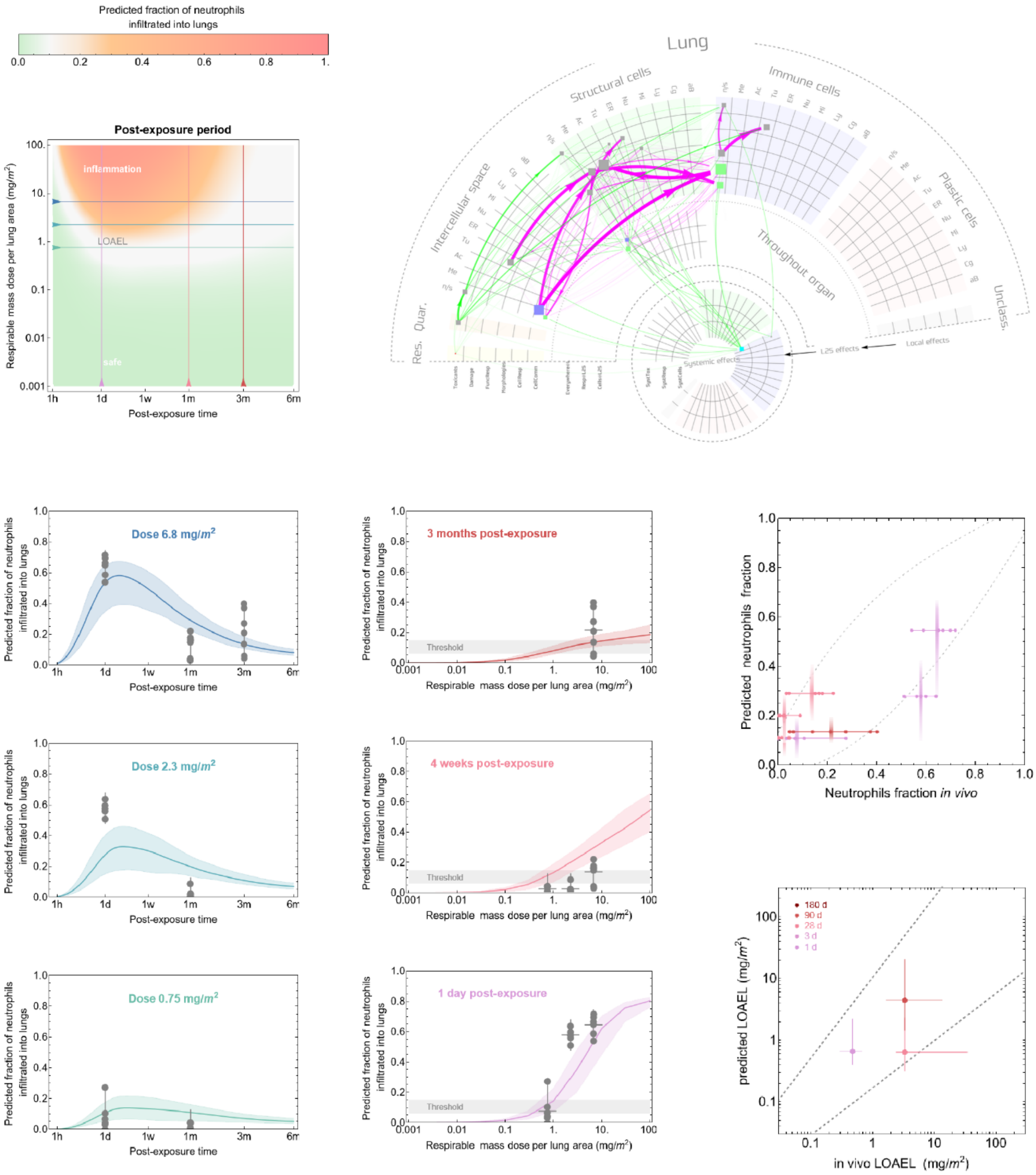
Predicted fraction of neutrophils infiltrated into lungs
Post-exposure period
inflammation
LOAEL
safe
Respirable mass dose per lung area (mg/m²)
Post-exposure time
Lung
Structural cells
Immune cells
Intercellular space
Plastic cells
Throughout organ
Res.
Quar.
Unclass.
Systemic effects
L2S effects
Local effects
Dose 6.8 mg/m²
Dose 2.3 mg/m²
Dose 0.75 mg/m²
3 months post-exposure
4 weeks post-exposure
1 day post-exposure
Threshold
Predicted neutrophils fraction
Neutrophils fraction in vivo
180 d
90 d
28 d
3 d
1 d
predicted LOAEL (mg/m²)
in vivo LOAEL (mg/m²)

### Carbon 9

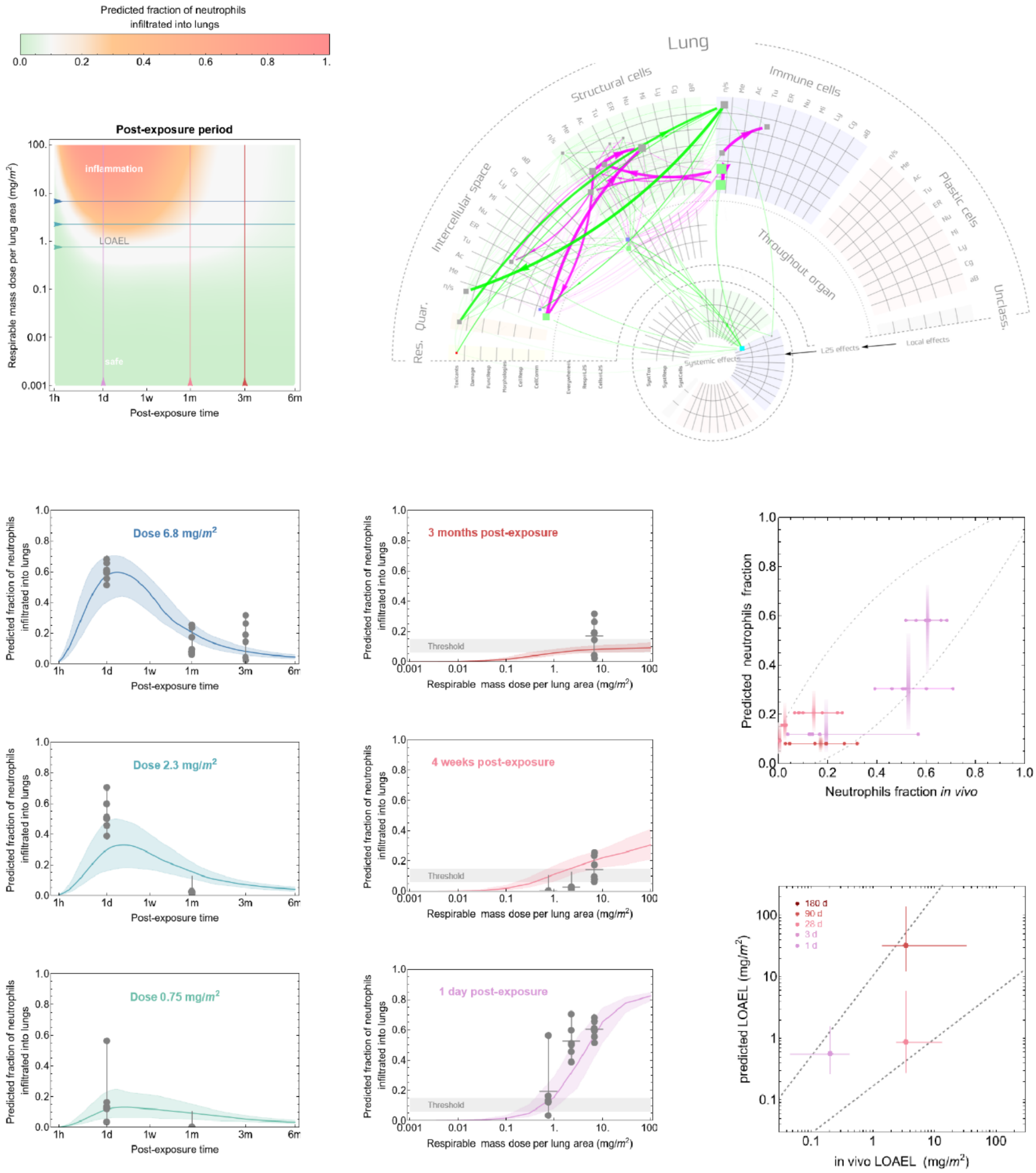

Predicted fraction of neutrophils infiltrated into lungs
0.0
0.2
0.4
0.6
0.8
1.
Post-exposure period
inflammation
LOAEL
safe
Respirable mass dose per lung area (mg/m2)
Post-exposure time
1h
1d
1w
1m
3m
6m
Lung
Structural cells
Immune cells
Intercellular space
Plastic cells
Throughout organ
Unclass.
Res.
Quar.
Systemic effects
L2S effects
Local effects
Dose 6.8 mg/m2
Dose 2.3 mg/m2
Dose 0.75 mg/m2
3 months post-exposure
4 weeks post-exposure
1 day post-exposure
Threshold
Predicted fraction of neutrophils infiltrated into lungs
Respirable mass dose per lung area (mg/m2)
Predicted neutrophils fraction
Neutrophils fraction in vivo
180 d
90 d
28 d
3 d
1 d
predicted LOAEL (mg/m2)
in vivo LOAEL (mg/m2)

### Carbon 10

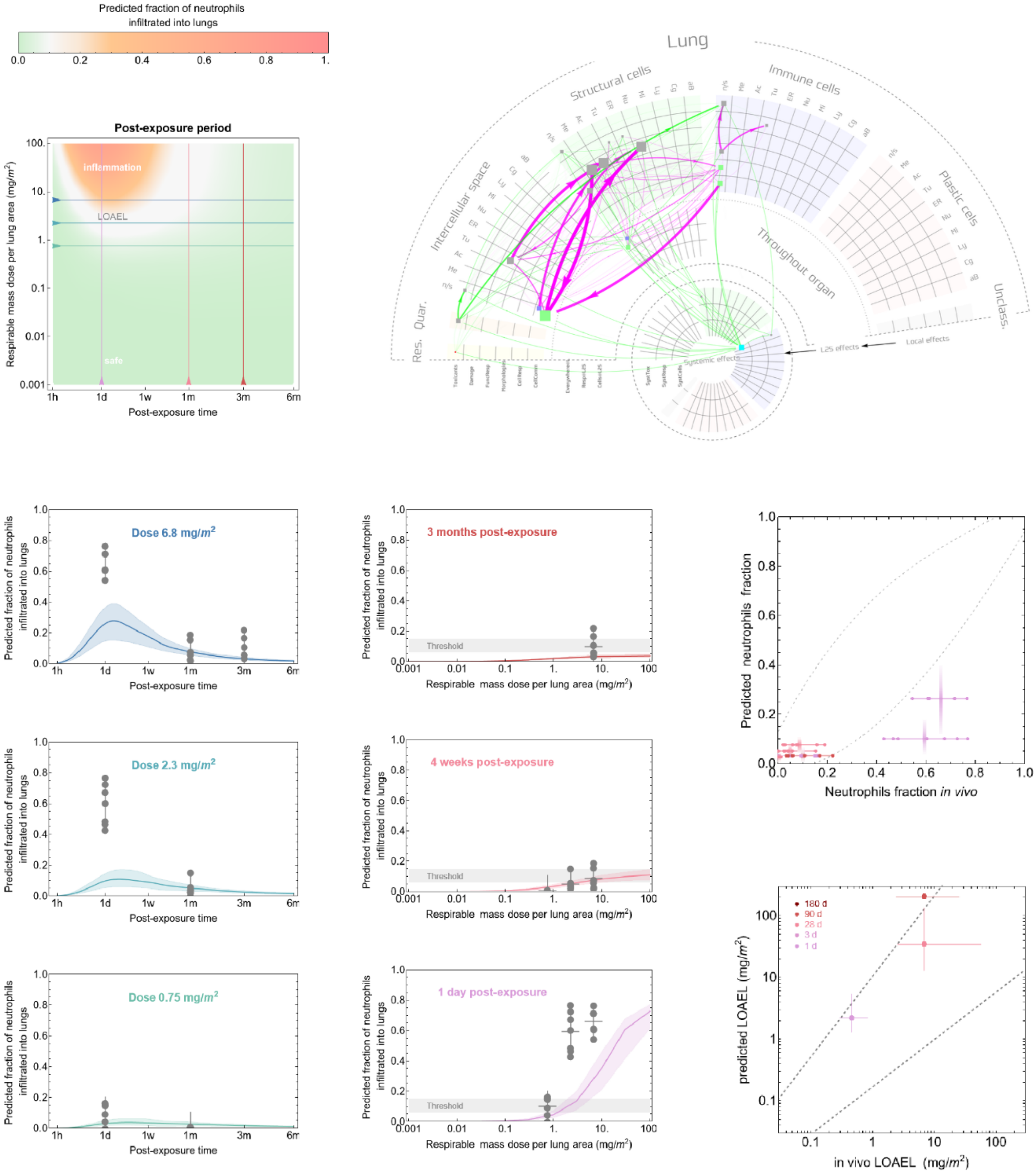

Predicted fraction of neutrophils infiltrated into lungs
0.0
0.2
0.4
0.6
0.8
1.
Post-exposure period
inflammation
LOAEL
safe
Respirable mass dose per lung area (mg/m²)
Post-exposure time
1h
1d
1w
1m
3m
6m
Lung
Structural cells
Immune cells
Plastic cells
Intercellular space
Throughout organ
Res.
Quar.
Unclass.
Local effects
L2S effects
Systemic effects
Dose 6.8 mg/m²
Dose 2.3 mg/m²
Dose 0.75 mg/m²
Predicted fraction of neutrophils infiltrated into lungs
3 months post-exposure
4 weeks post-exposure
1 day post-exposure
Threshold
Respirable mass dose per lung area (mg/m²)
Predicted neutrophils fraction
Neutrophils fraction in vivo
180 d
90 d
28 d
3 d
1 d
predicted LOAEL (mg/m²)
in vivo LOAEL (mg/m²)

#### Carbon 11

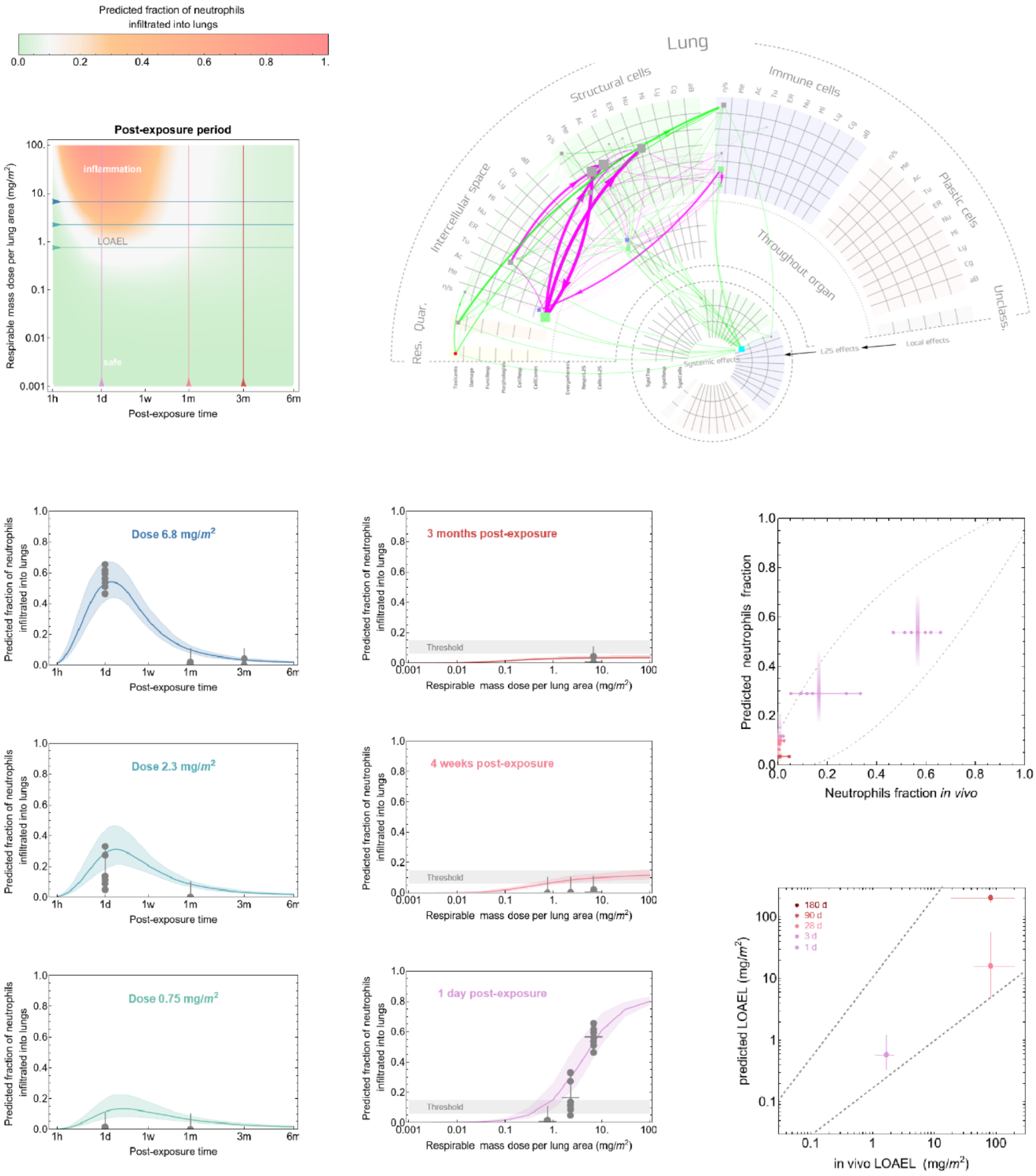
Predicted fraction of neutrophils infiltrated into lungs
Post-exposure period
inflammation
LOAEL
safe
Respirable mass dose per lung area (mg/m²)
Post-exposure time
Lung
Structural cells
Immune cells
Intercellular space
Throughout organ
Plastic cells
Res.
Quar.
Unclass.
Systemic effects
L2S effects
Local effects
Dose 6.8 mg/m²
Dose 2.3 mg/m²
Dose 0.75 mg/m²
3 months post-exposure
4 weeks post-exposure
1 day post-exposure
Threshold
Predicted neutrophils fraction
Neutrophils fraction in vivo
180 d
90 d
28 d
3 d
1 d
predicted LOAEL (mg/m²)
in vivo LOAEL (mg/m²)

#### Carbon 12

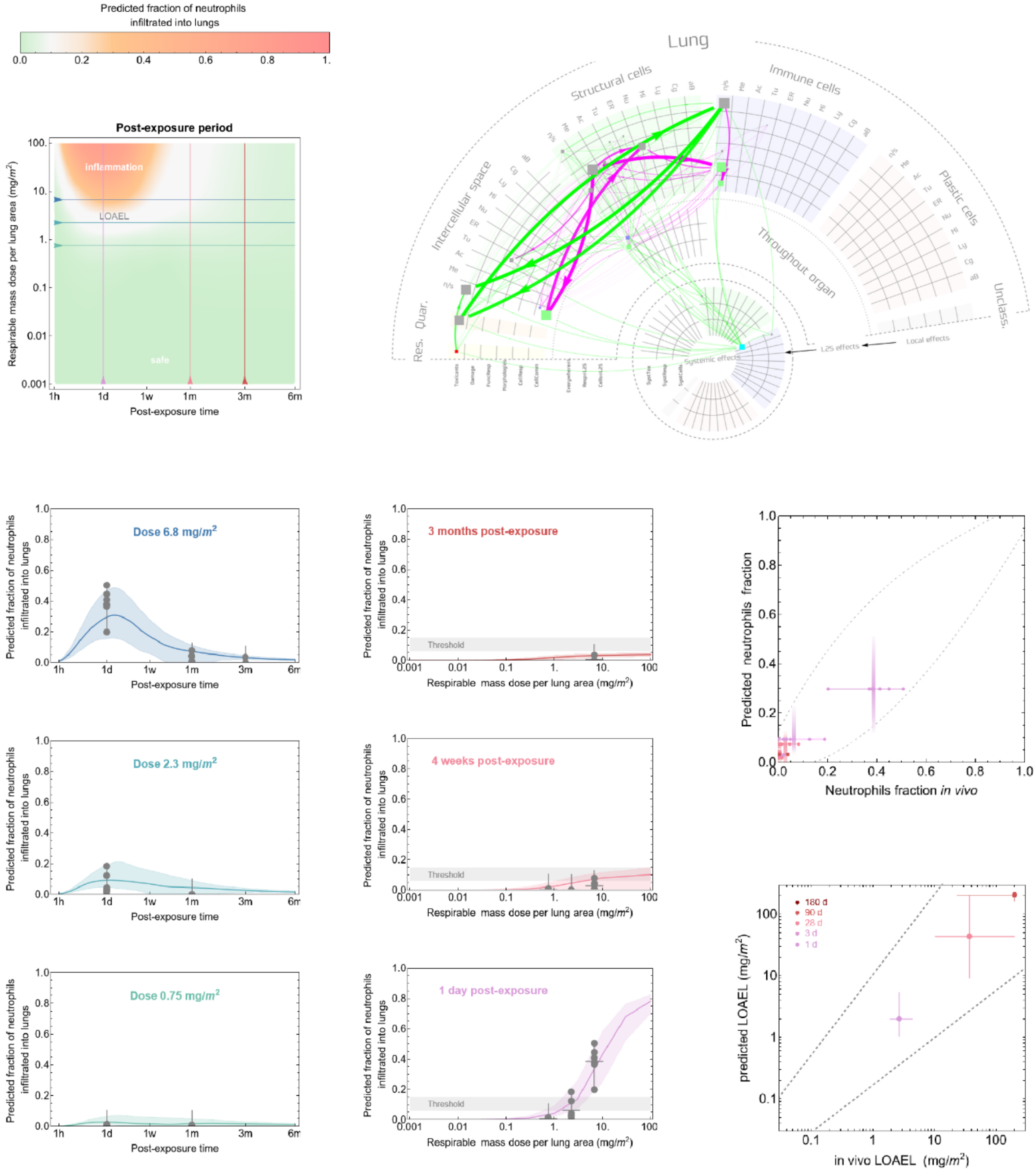

Predicted fraction of neutrophils infiltrated into lungs
Post-exposure period
inflammation
LOAEL
safe
Respirable mass dose per lung area (mg/m²)
Post-exposure time
Lung
Structural cells
Immune cells
Intercellular space
Plastic cells
Throughout organ
Res. Quar.
Unclass.
Systemic effects
Local effects
Dose 6.8 mg/m²
Dose 2.3 mg/m²
Dose 0.75 mg/m²
3 months post-exposure
4 weeks post-exposure
1 day post-exposure
Threshold
Predicted neutrophils fraction
Neutrophils fraction in vivo
180 d
90 d
28 d
3 d
1 d
predicted LOAEL (mg/m²)
in vivo LOAEL (mg/m²)

#### Carbon 13

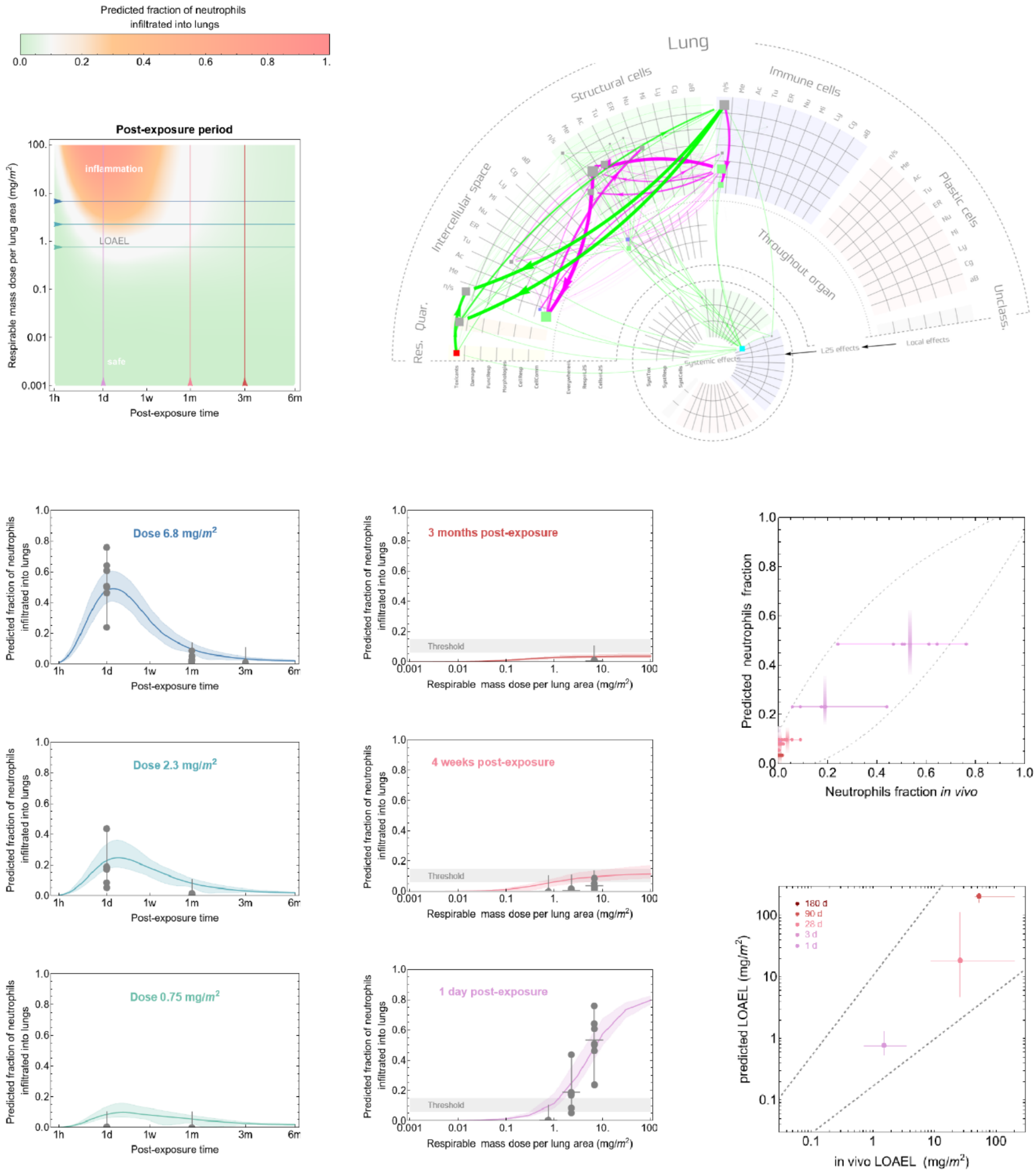

Predicted fraction of neutrophils infiltrated into lungs
Post-exposure period
inflammation
LOAEL
safe
Respirable mass dose per lung area (mg/m²)
Post-exposure time
Lung
Structural cells
Immune cells
Intercellular space
Plastic cells
Throughout organ
Res.
Quar.
Unclass.
Systemic effects
L2S effects
Local effects
Dose 6.8 mg/m²
Dose 2.3 mg/m²
Dose 0.75 mg/m²
3 months post-exposure
4 weeks post-exposure
1 day post-exposure
Threshold
Predicted neutrophils fraction
Neutrophils fraction in vivo
180 d
90 d
28 d
3 d
1 d
predicted LOAEL (mg/m²)
in vivo LOAEL (mg/m²)

**Carbon 14**

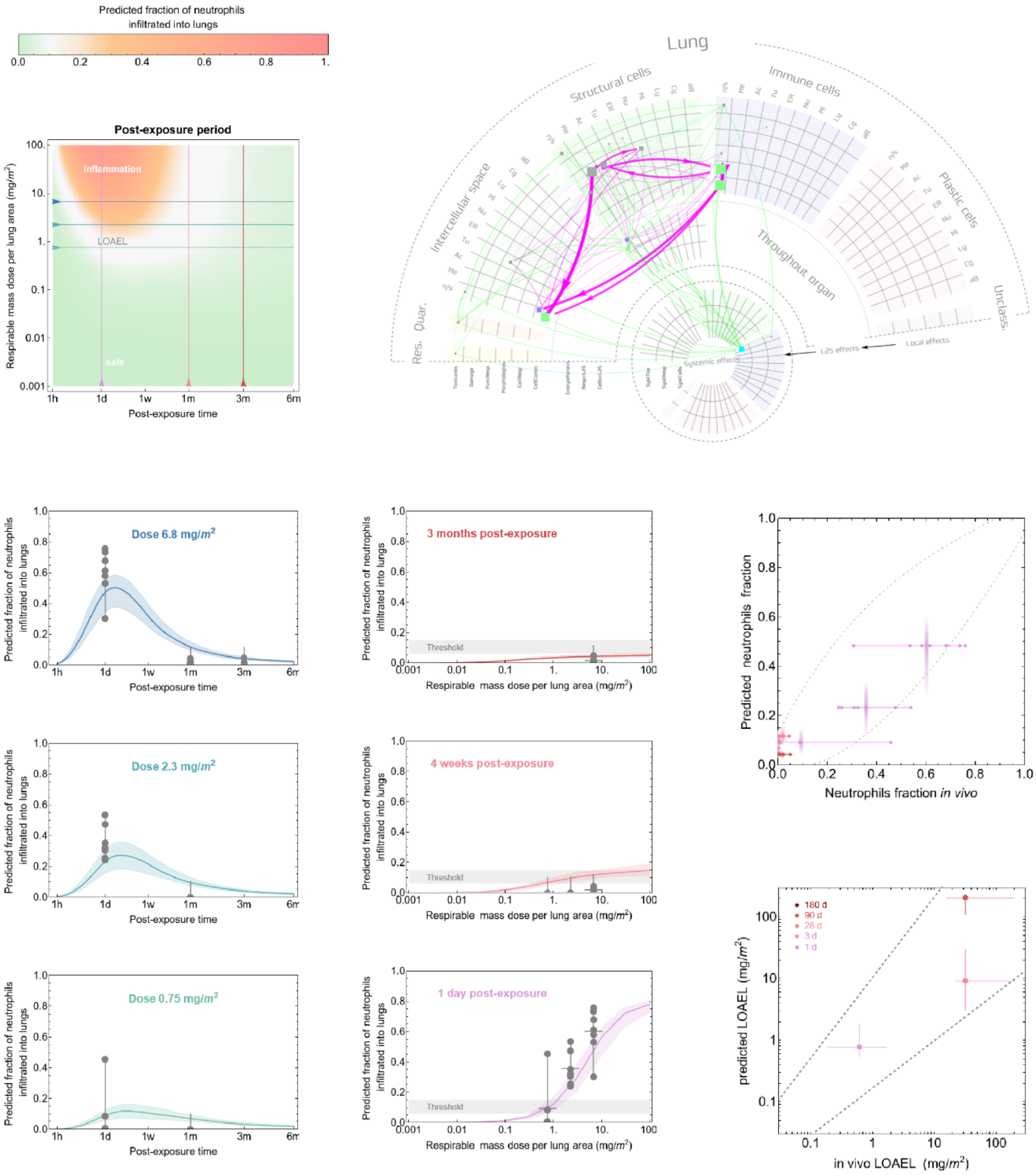

Predicted fraction of neutrophils infiltrated into lungs
Post-exposure period
inflammation
LOAEL
safe
Respirable mass dose per lung area (mg/m²)
Post-exposure time
Lung
Structural cells
Immune cells
Intercellular space
Throughout organ
Plastic cells
Res.
Quar.
Unclass.
Systemic effects
L2S effects
Local effects
Dose 6.8 mg/m²
Dose 2.3 mg/m²
Dose 0.75 mg/m²
3 months post-exposure
4 weeks post-exposure
1 day post-exposure
Threshold
Predicted neutrophils fraction
Neutrophils fraction in vivo
180 d
90 d
28 d
3 d
1 d
predicted LOAEL (mg/m²)
in vivo LOAEL (mg/m²)

**Carbon 15**

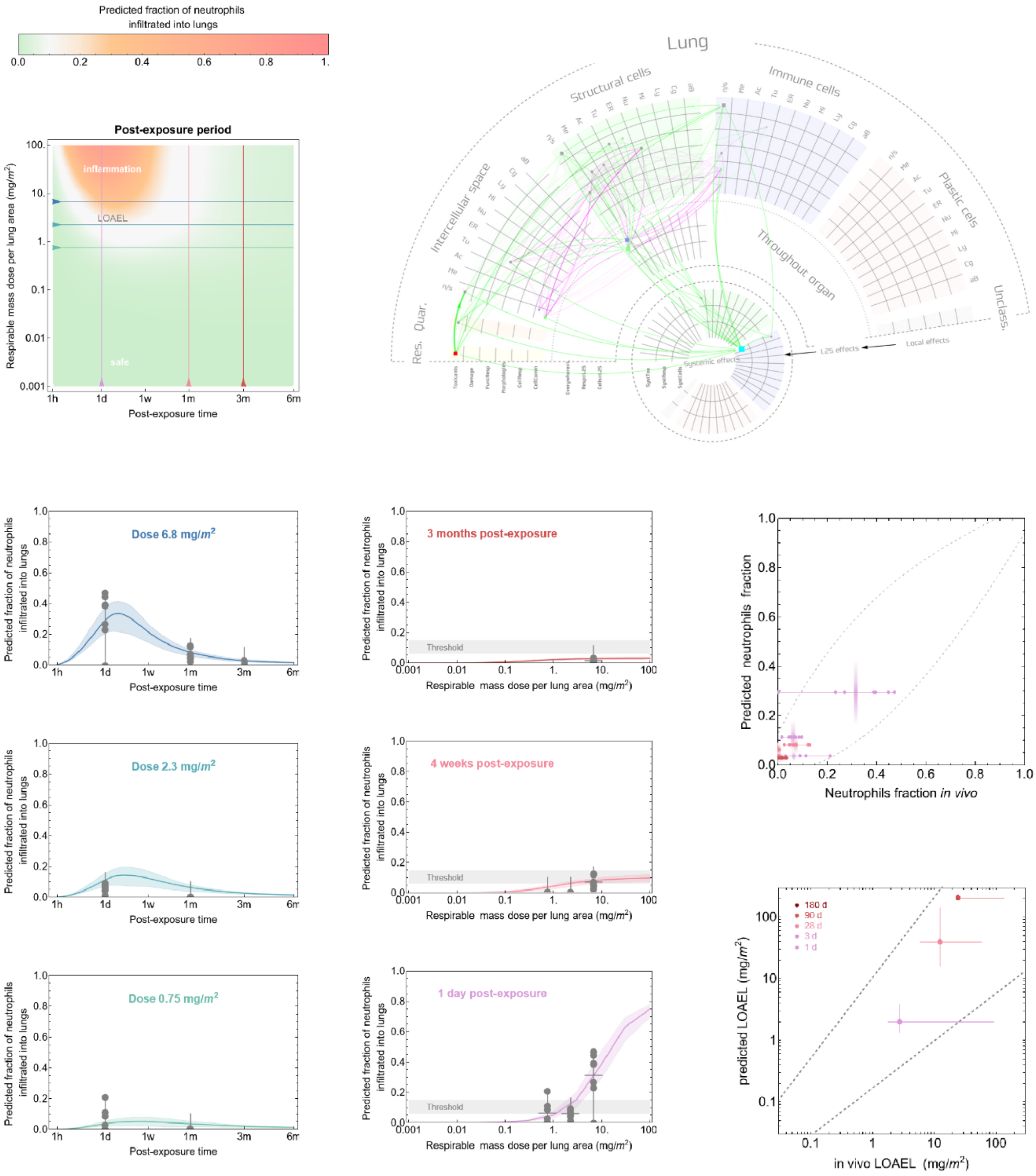

Predicted fraction of neutrophils infiltrated into lungs
Post-exposure period
inflammation
LOAEL
safe
Respirable mass dose per lung area (mg/m2)
Post-exposure time
Lung
Structural cells
Immune cells
Intercellular space
Throughout organ
Plastic cells
Res.
Quar.
Unclass.
Systemic effects
L2S effects
Local effects
Dose 6.8 mg/m2
Dose 2.3 mg/m2
Dose 0.75 mg/m2
3 months post-exposure
4 weeks post-exposure
1 day post-exposure
Threshold
Predicted fraction of neutrophils infiltrated into lungs
Predicted neutrophils fraction
Neutrophils fraction in vivo
180 d
90 d
28 d
3 d
1 d
predicted LOAEL (mg/m2)
in vivo LOAEL (mg/m2)

#### Carbon 16

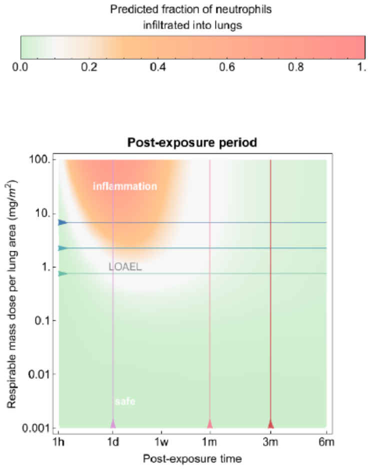
Predicted fraction of neutrophils
infiltrated into lungs
0.0
0.2
0.4
0.6
0.8
1.
Post-exposure period
inflammation
LOAEL
safe
Respirable mass dose per lung area (mg/m$^2$)
100.
10.
1.
0.1
0.01
0.001
1h
1d
1w
1m
3m
6m
Post-exposure time

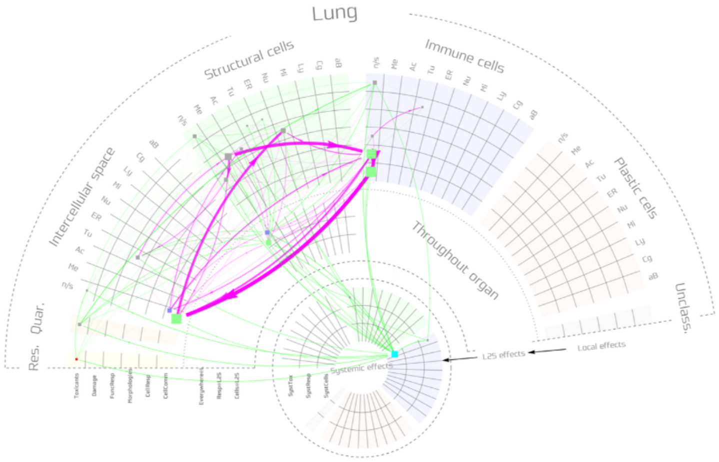
Lung
Structural cells
Immune cells
Intercellular space
Plastic cells
Throughout organ
Res.
Quar.
Unclass.
Systemic effects
L2S effects
Local effects

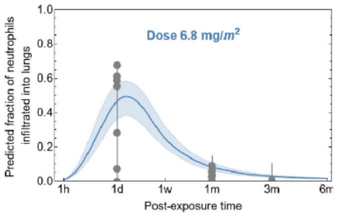
Dose 6.8 mg/$m^2$
Predicted fraction of neutrophils
infiltrated into lungs
1.0
0.8
0.6
0.4
0.2
0.0
1h
1d
1w
1m
3m
6m
Post-exposure time

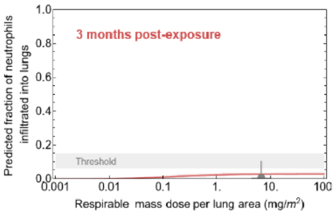
3 months post-exposure
Predicted fraction of neutrophils
infiltrated into lungs
Threshold
0.001
0.01
0.1
1.
10.
100
Respirable mass dose per lung area (mg/$m^2$)

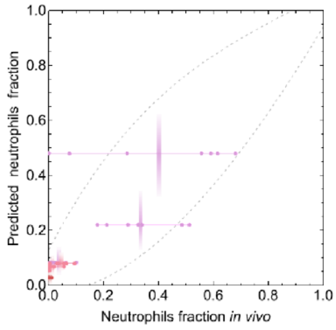
Predicted neutrophils fraction
1.0
0.8
0.6
0.4
0.2
0.0
0.0
0.2
0.4
0.6
0.8
1.0
Neutrophils fraction *in vivo*

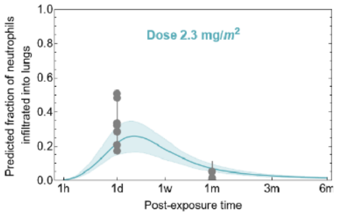
Dose 2.3 mg/$m^2$
Predicted fraction of neutrophils
infiltrated into lungs
1.0
0.8
0.6
0.4
0.2
0.0
1h
1d
1w
1m
3m
6m
Post-exposure time

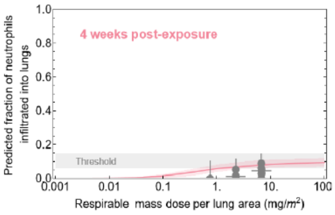
4 weeks post-exposure
Predicted fraction of neutrophils
infiltrated into lungs
Threshold
0.001
0.01
0.1
1.
10.
100
Respirable mass dose per lung area (mg/$m^2$)

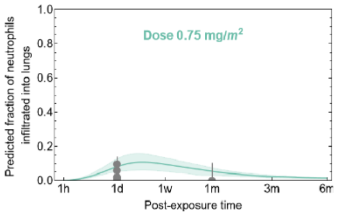
Dose 0.75 mg/$m^2$
Predicted fraction of neutrophils
infiltrated into lungs
1.0
0.8
0.6
0.4
0.2
0.0
1h
1d
1w
1m
3m
6m
Post-exposure time

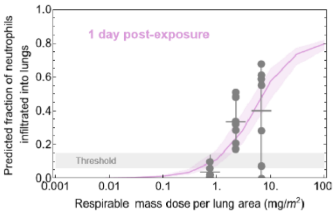
1 day post-exposure
Predicted fraction of neutrophils
infiltrated into lungs
Threshold
0.001
0.01
0.1
1.
10.
100
Respirable mass dose per lung area (mg/$m^2$)

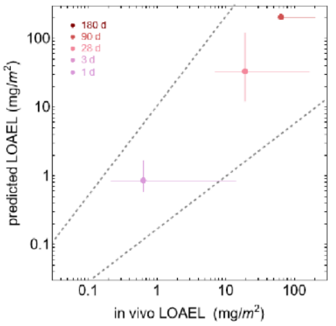
180 d
90 d
28 d
3 d
1 d
predicted LOAEL (mg/$m^2$)
100
10
1
0.1
0.1
1
10
100
in vivo LOAEL (mg/$m^2$)

### Carbon 17

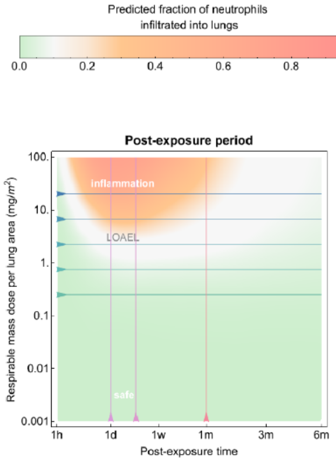

Predicted fraction of neutrophils infiltrated into lungs
0.0
0.2
0.4
0.6
0.8
1.
Post-exposure period
inflammation
LOAEL
safe
Respirable mass dose per lung area (mg/m²)
Post-exposure time
1h
1d
1w
1m
3m
6m

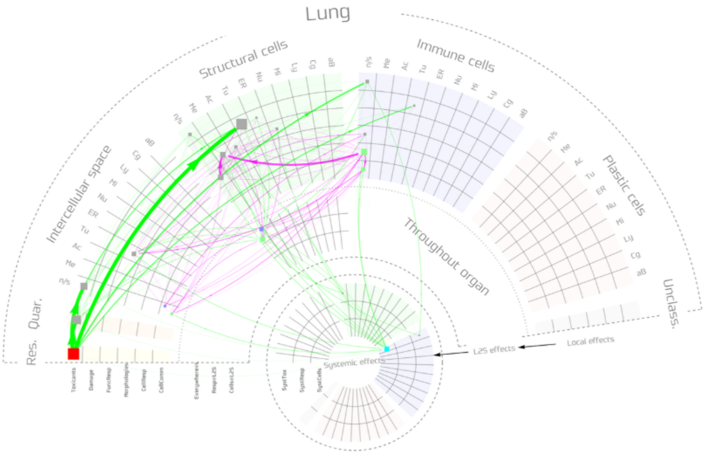

Lung
Structural cells
Immune cells
Plastic cells
Intercellular space
Throughout organ
Quar.
Res.
Unclass.
Systemic effects
L2S effects
Local effects

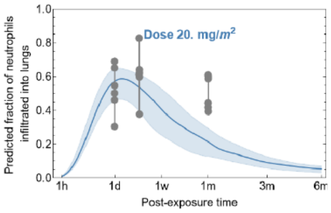

Dose 20. mg/m²
Predicted fraction of neutrophils infiltrated into lungs
Post-exposure time

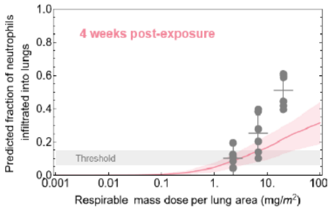

4 weeks post-exposure
Threshold
Predicted fraction of neutrophils infiltrated into lungs
Respirable mass dose per lung area (mg/m²)

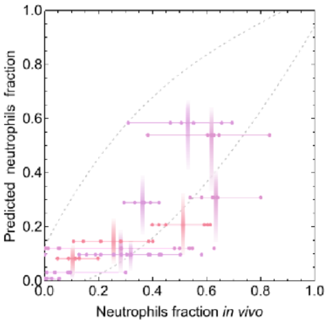

Predicted neutrophils fraction
Neutrophils fraction in vivo

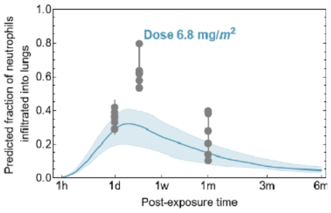

Dose 6.8 mg/m²
Predicted fraction of neutrophils infiltrated into lungs
Post-exposure time

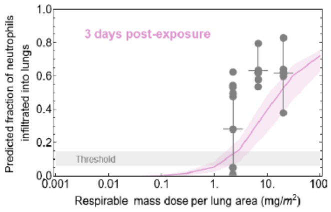

3 days post-exposure
Threshold
Predicted fraction of neutrophils infiltrated into lungs
Respirable mass dose per lung area (mg/m²)

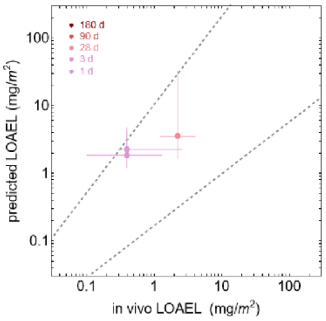

180 d
90 d
28 d
3 d
1 d
predicted LOAEL (mg/m²)
in vivo LOAEL (mg/m²)

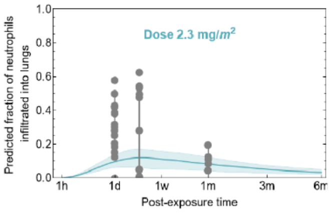

Dose 2.3 mg/m²
Predicted fraction of neutrophils infiltrated into lungs
Post-exposure time

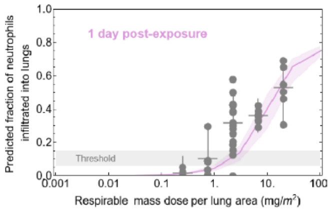

1 day post-exposure
Threshold
Predicted fraction of neutrophils infiltrated into lungs
Respirable mass dose per lung area (mg/m²)

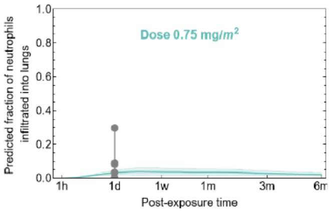

Dose 0.75 mg/m²
Predicted fraction of neutrophils infiltrated into lungs
Post-exposure time

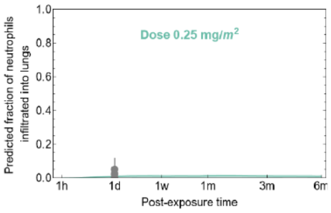

Dose 0.25 mg/m²
Predicted fraction of neutrophils infiltrated into lungs
Post-exposure time

### Carbon 18

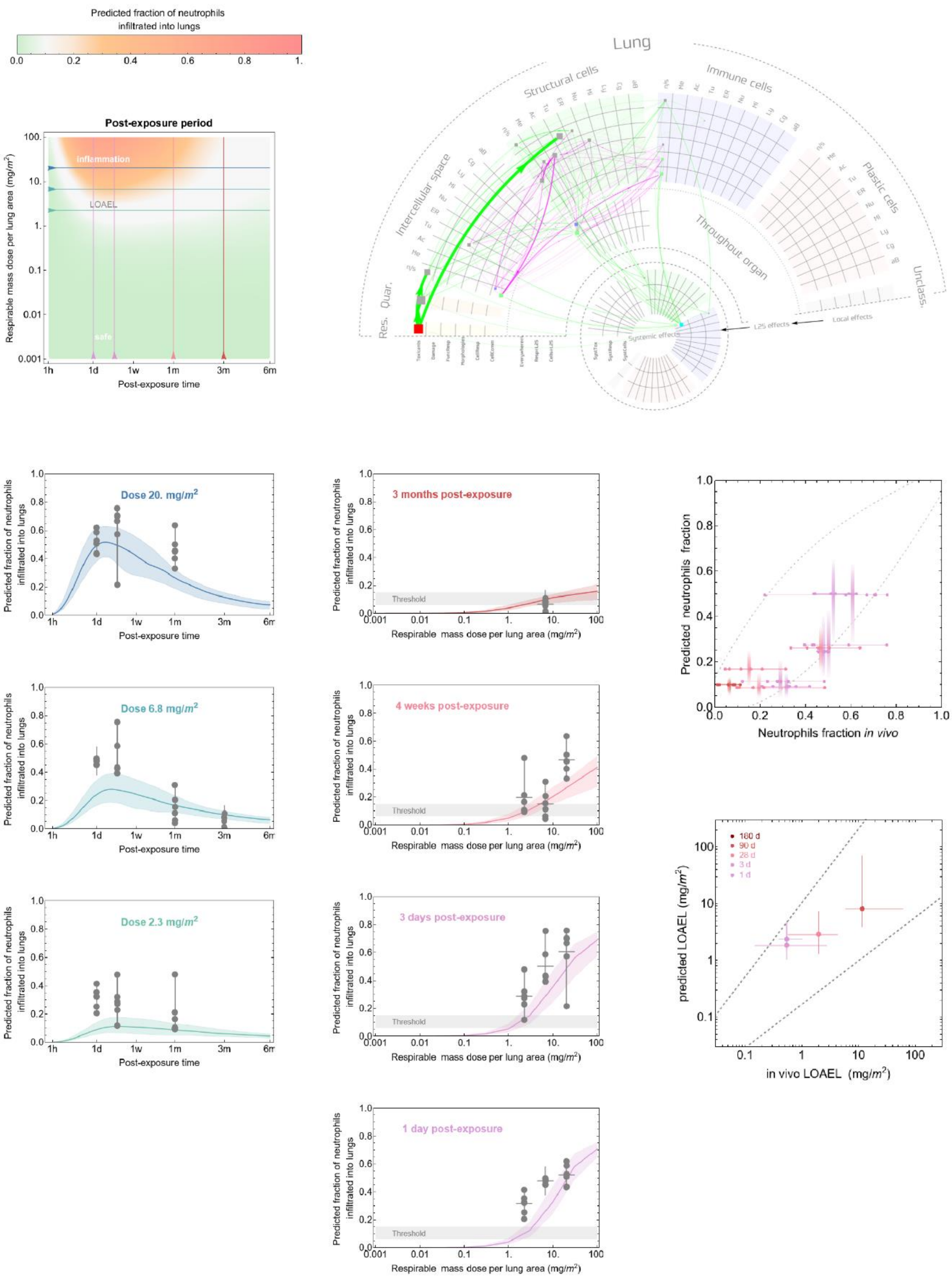

Predicted fraction of neutrophils infiltrated into lungs
Post-exposure period
inflammation
LOAEL
safe
Respirable mass dose per lung area (mg/m²)
Post-exposure time
Lung
Structural cells
Immune cells
Intercellular space
Plastic cells
Throughout organ
Res.
Quar.
Unclass.
Systemic effects
L2S effects
Local effects
Dose 20. mg/m²
Dose 6.8 mg/m²
Dose 2.3 mg/m²
3 months post-exposure
4 weeks post-exposure
3 days post-exposure
1 day post-exposure
Threshold
Predicted neutrophils fraction
Neutrophils fraction in vivo
180 d
90 d
28 d
3 d
1 d
predicted LOAEL (mg/m²)
in vivo LOAEL (mg/m²)

#### Carbon 19

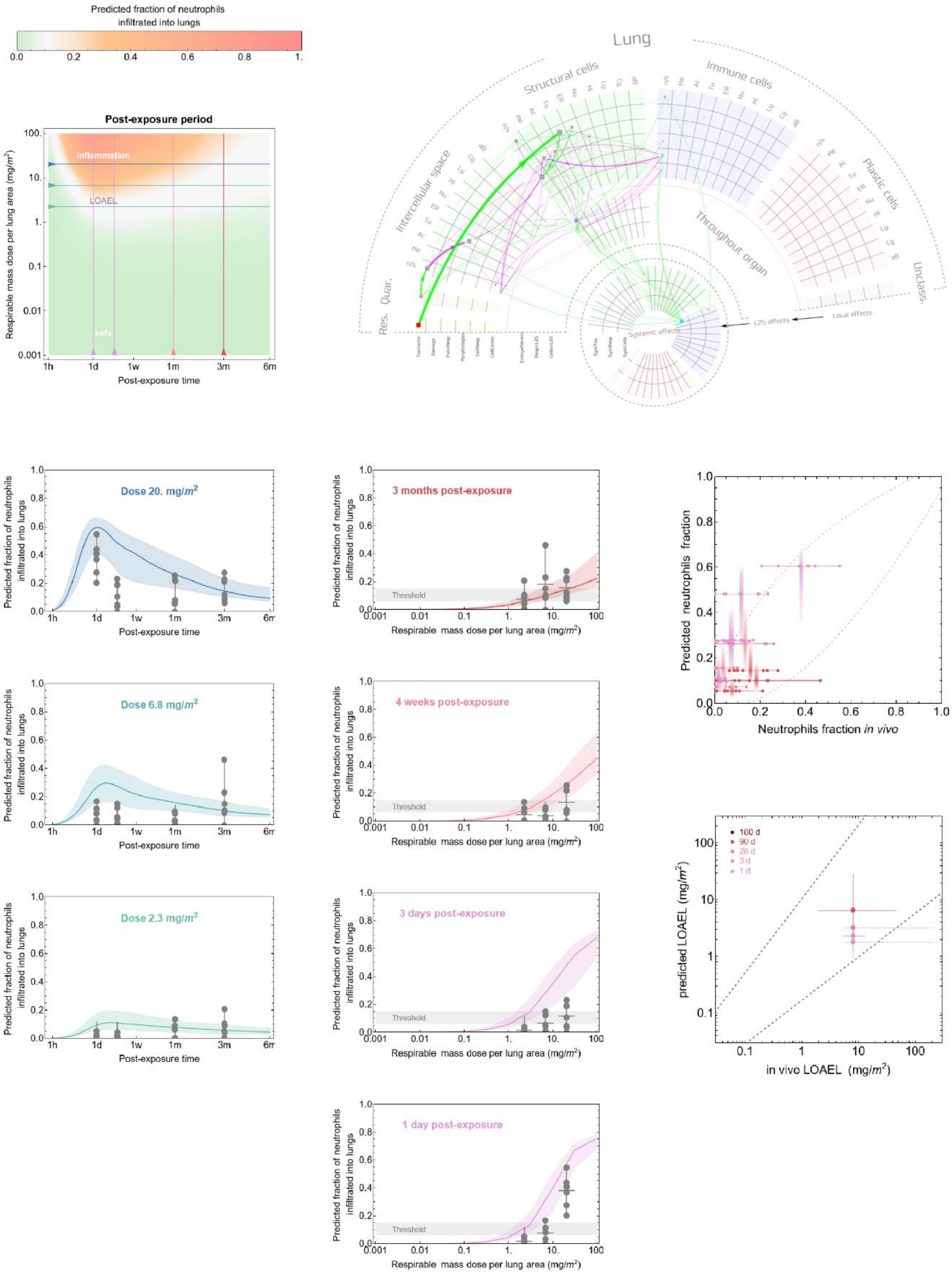

Predicted fraction of neutrophils infiltrated into lungs
0.0 0.2 0.4 0.6 0.8 1.
Post-exposure period
inflammation
LOAEL
safe
Respirable mass dose per lung area (mg/m²)
Post-exposure time
1h 1d 1w 1m 3m 6m
Lung
Structural cells
Immune cells
Intercellular space
Plastic cells
Throughout organ
Res.
Quar.
Unclass.
L2S effects
Local effects
Systemic effects
Dose 20. mg/m²
Dose 6.8 mg/m²
Dose 2.3 mg/m²
Predicted fraction of neutrophils infiltrated into lungs
3 months post-exposure
4 weeks post-exposure
3 days post-exposure
1 day post-exposure
Threshold
Respirable mass dose per lung area (mg/m²)
Predicted neutrophils fraction
Neutrophils fraction in vivo
180 d
90 d
28 d
3 d
1 d
predicted LOAEL (mg/m²)
in vivo LOAEL (mg/m²)

#### Carbon 20

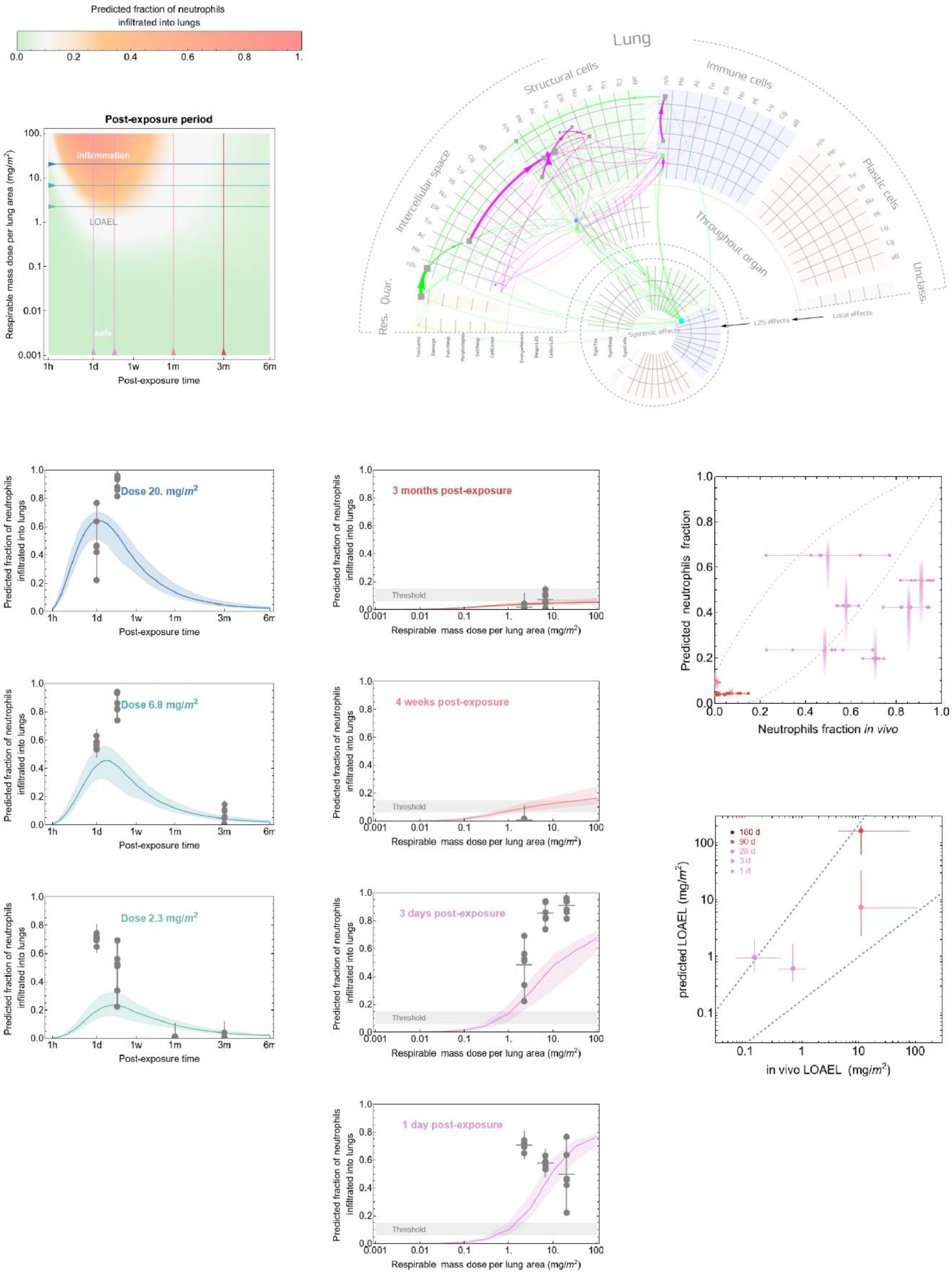

Predicted fraction of neutrophils infiltrated into lungs
0.0 0.2 0.4 0.6 0.8 1.
Post-exposure period
inflammation
LOAEL
safe
Respirable mass dose per lung area (mg/m²)
Post-exposure time
1h 1d 1w 1m 3m 6m
Lung
Structural cells
Immune cells
Intercellular space
Plastic cells
Throughout organ
Unclass.
Res.
Quar.
Systemic effects
L2S effects
Local effects
Dose 20. mg/m²
Dose 6.8 mg/m²
Dose 2.3 mg/m²
3 months post-exposure
4 weeks post-exposure
3 days post-exposure
1 day post-exposure
Threshold
Predicted neutrophils fraction
Neutrophils fraction in vivo
180 d
90 d
28 d
3 d
1 d
predicted LOAEL (mg/m²)
in vivo LOAEL (mg/m²)

### Clay 1

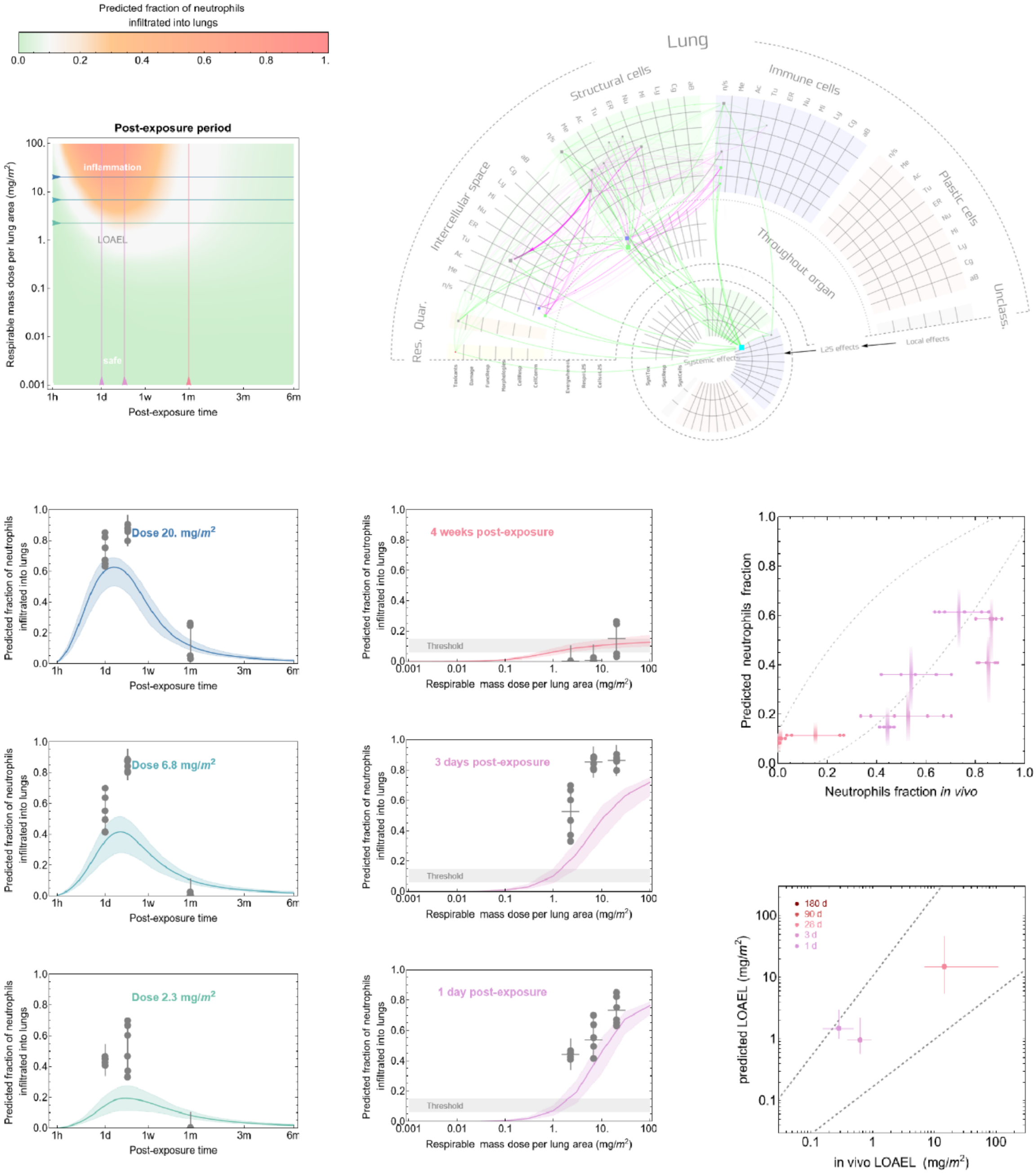

Predicted fraction of neutrophils infiltrated into lungs
0.0
0.2
0.4
0.6
0.8
1.
Post-exposure period
inflammation
LOAEL
safe
Respirable mass dose per lung area (mg/m²)
Post-exposure time
1h
1d
1w
1m
3m
6m
Lung
Structural cells
Immune cells
Plastic cells
Intercellular space
Throughout organ
Res.
Quar.
Unclass.
Systemic effects
L2S effects
Local effects
Dose 20. mg/m²
Dose 6.8 mg/m²
Dose 2.3 mg/m²
4 weeks post-exposure
3 days post-exposure
1 day post-exposure
Threshold
Respirable mass dose per lung area (mg/m²)
Predicted neutrophils fraction
Neutrophils fraction in vivo
180 d
90 d
28 d
3 d
1 d
predicted LOAEL (mg/m²)
in vivo LOAEL (mg/m²)

#### Clay 2

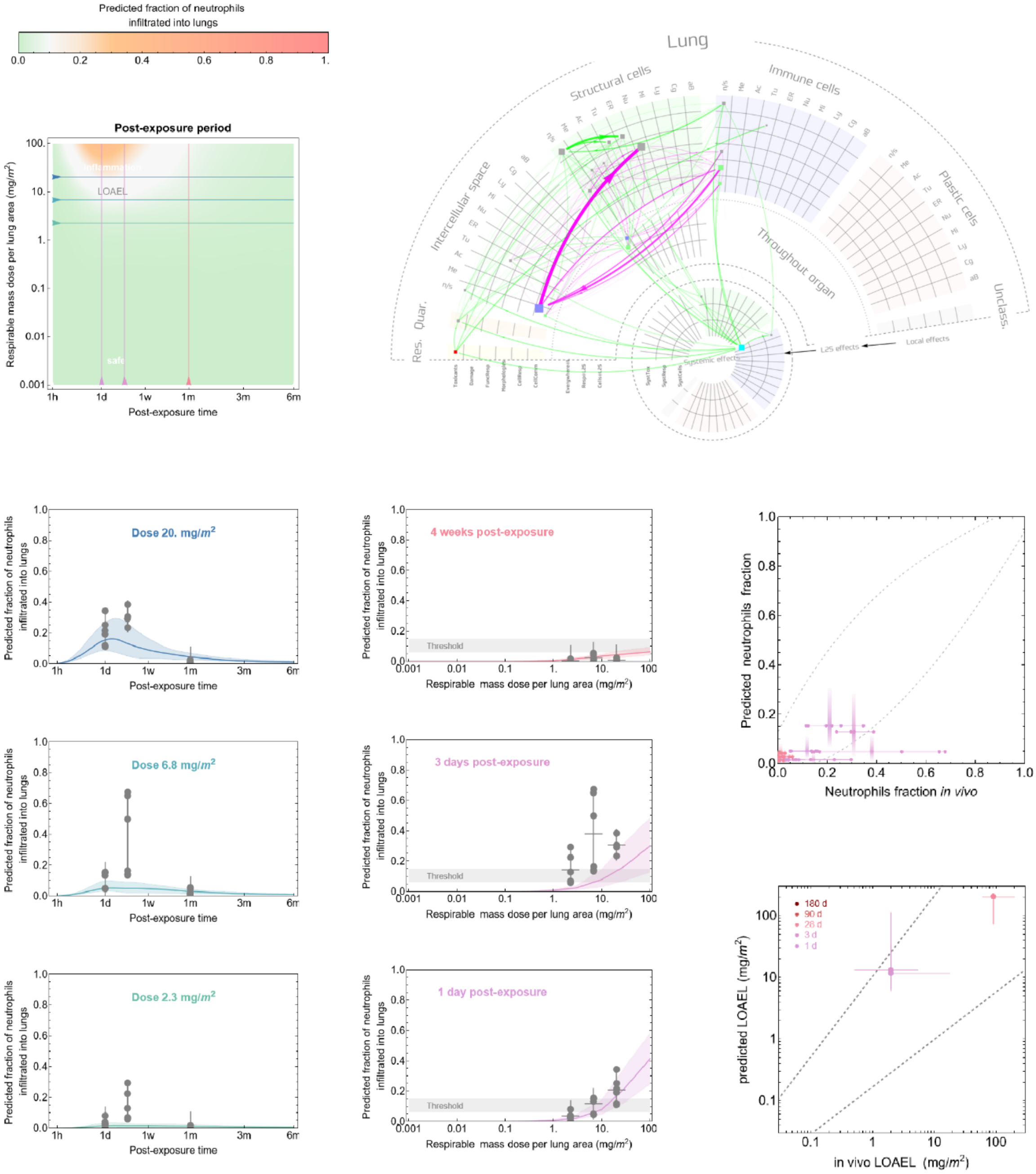

Predicted fraction of neutrophils infiltrated into lungs
0.0
0.2
0.4
0.6
0.8
1.
Post-exposure period
Respirable mass dose per lung area (mg/m²)
inflammation
LOAEL
safe
1h
1d
1w
1m
3m
6m
Post-exposure time
Lung
Structural cells
Immune cells
Intercellular space
Plastic cells
Throughout organ
Res. Quar.
Unclass.
Systemic effects
L2S effects
Local effects
Dose 20. mg/m²
Dose 6.8 mg/m²
Dose 2.3 mg/m²
4 weeks post-exposure
3 days post-exposure
1 day post-exposure
Threshold
Respirable mass dose per lung area (mg/m²)
Predicted neutrophils fraction
Neutrophils fraction in vivo
180 d
90 d
28 d
3 d
1 d
predicted LOAEL (mg/m²)
in vivo LOAEL (mg/m²)

### Clay 3

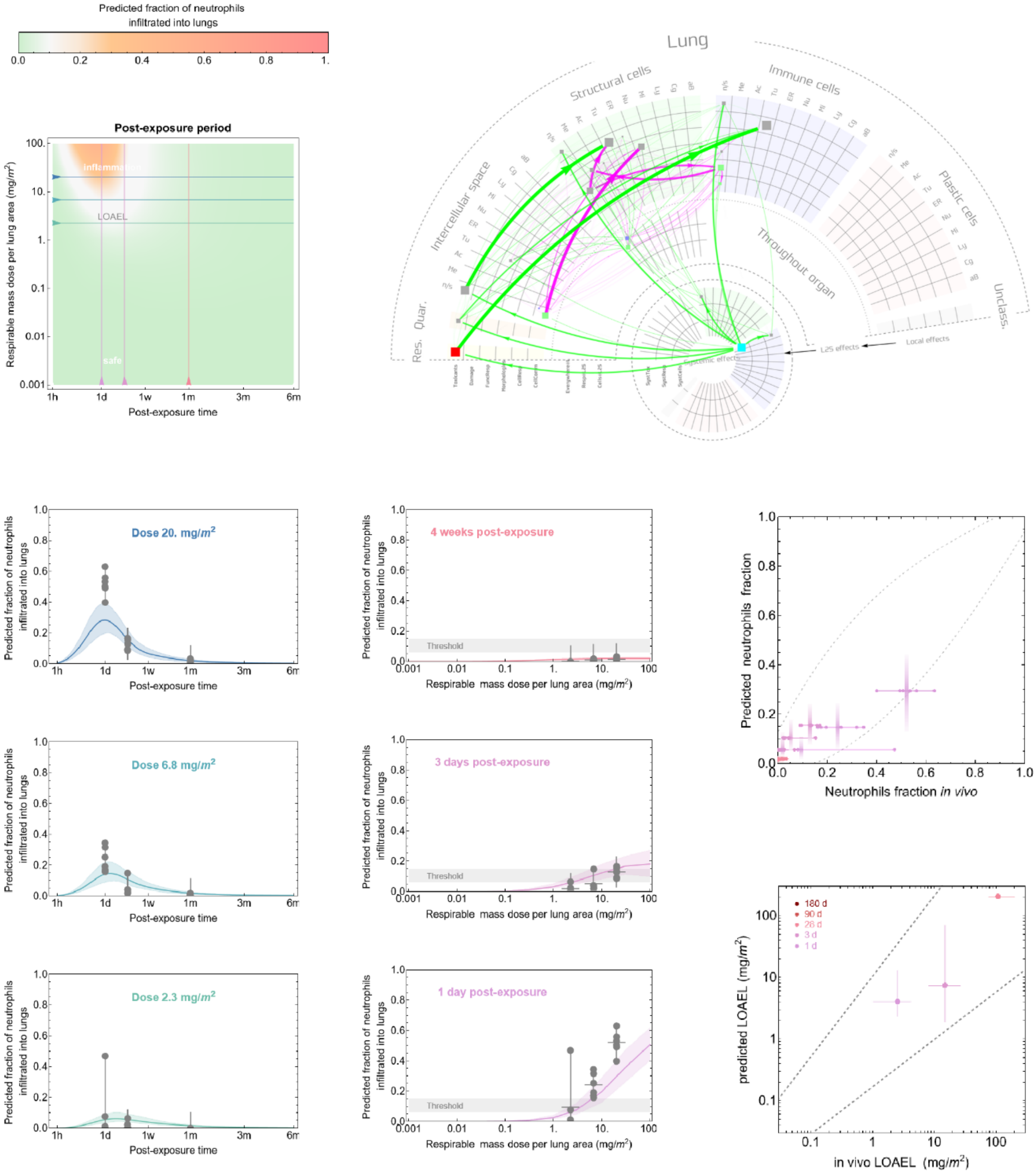

Predicted fraction of neutrophils infiltrated into lungs
0.0 0.2 0.4 0.6 0.8 1.
Post-exposure period
inflammation
LOAEL
safe
Respirable mass dose per lung area (mg/m²)
Post-exposure time
Lung
Structural cells
Immune cells
Intercellular space
Plastic cells
Throughout organ
Res.
Quar.
Unclass.
L2S effects
Local effects
Systemic effects
Dose 20. mg/m²
Dose 6.8 mg/m²
Dose 2.3 mg/m²
4 weeks post-exposure
3 days post-exposure
1 day post-exposure
Threshold
Predicted neutrophils fraction
Neutrophils fraction in vivo
180 d
90 d
28 d
3 d
1 d
predicted LOAEL (mg/m²)
in vivo LOAEL (mg/m²)

#### Clay 4

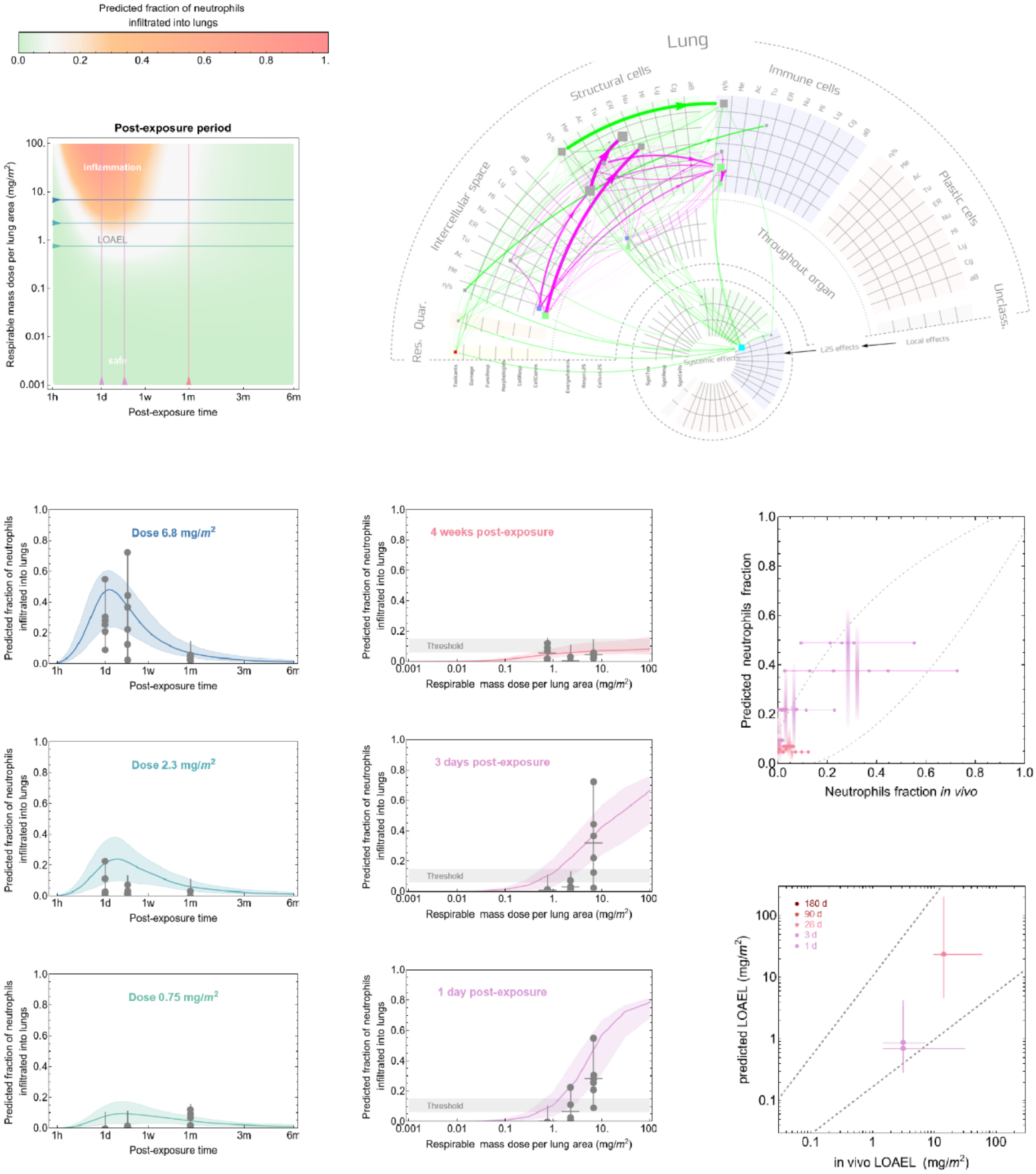

Predicted fraction of neutrophils infiltrated into lungs
Post-exposure period
Respirable mass dose per lung area (mg/m²)
inflammation
LOAEL
safe
Post-exposure time
Lung
Structural cells
Immune cells
Intercellular space
Throughout organ
Plastic cells
Res.
Quar.
Unclass.
Systemic effects
L2S effects
Local effects
Dose 6.8 mg/m²
Dose 2.3 mg/m²
Dose 0.75 mg/m²
4 weeks post-exposure
3 days post-exposure
1 day post-exposure
Threshold
Respirable mass dose per lung area (mg/m²)
Predicted neutrophils fraction
Neutrophils fraction in vivo
180 d
90 d
28 d
3 d
1 d
predicted LOAEL (mg/m²)
in vivo LOAEL (mg/m²)

#### Clay 5

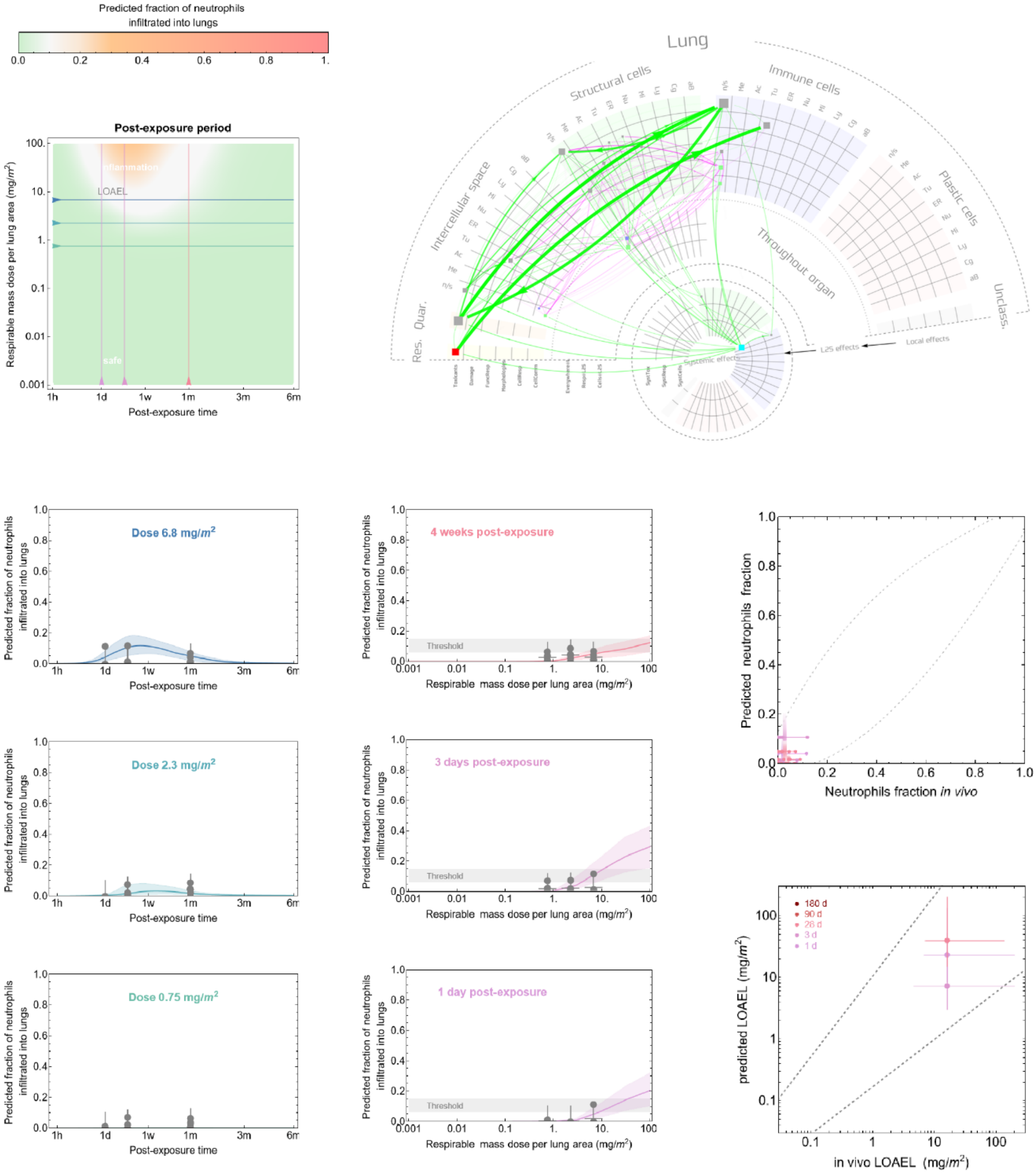

Predicted fraction of neutrophils infiltrated into lungs
0.0 0.2 0.4 0.6 0.8 1.
Post-exposure period
Inflammation
LOAEL
safe
Respirable mass dose per lung area (mg/m²)
Post-exposure time
1h 1d 1w 1m 3m 6m
Lung
Structural cells
Immune cells
Intercellular space
Plastic cells
Throughout organ
Unclass.
Res. Quar.
Systemic effects
L2S effects
Local effects
Dose 6.8 mg/m²
Dose 2.3 mg/m²
Dose 0.75 mg/m²
4 weeks post-exposure
3 days post-exposure
1 day post-exposure
Threshold
Respirable mass dose per lung area (mg/m²)
Predicted neutrophils fraction
Neutrophils fraction in vivo
180 d
90 d
28 d
3 d
1 d
predicted LOAEL (mg/m²)
in vivo LOAEL (mg/m²)

**Combustion 1**

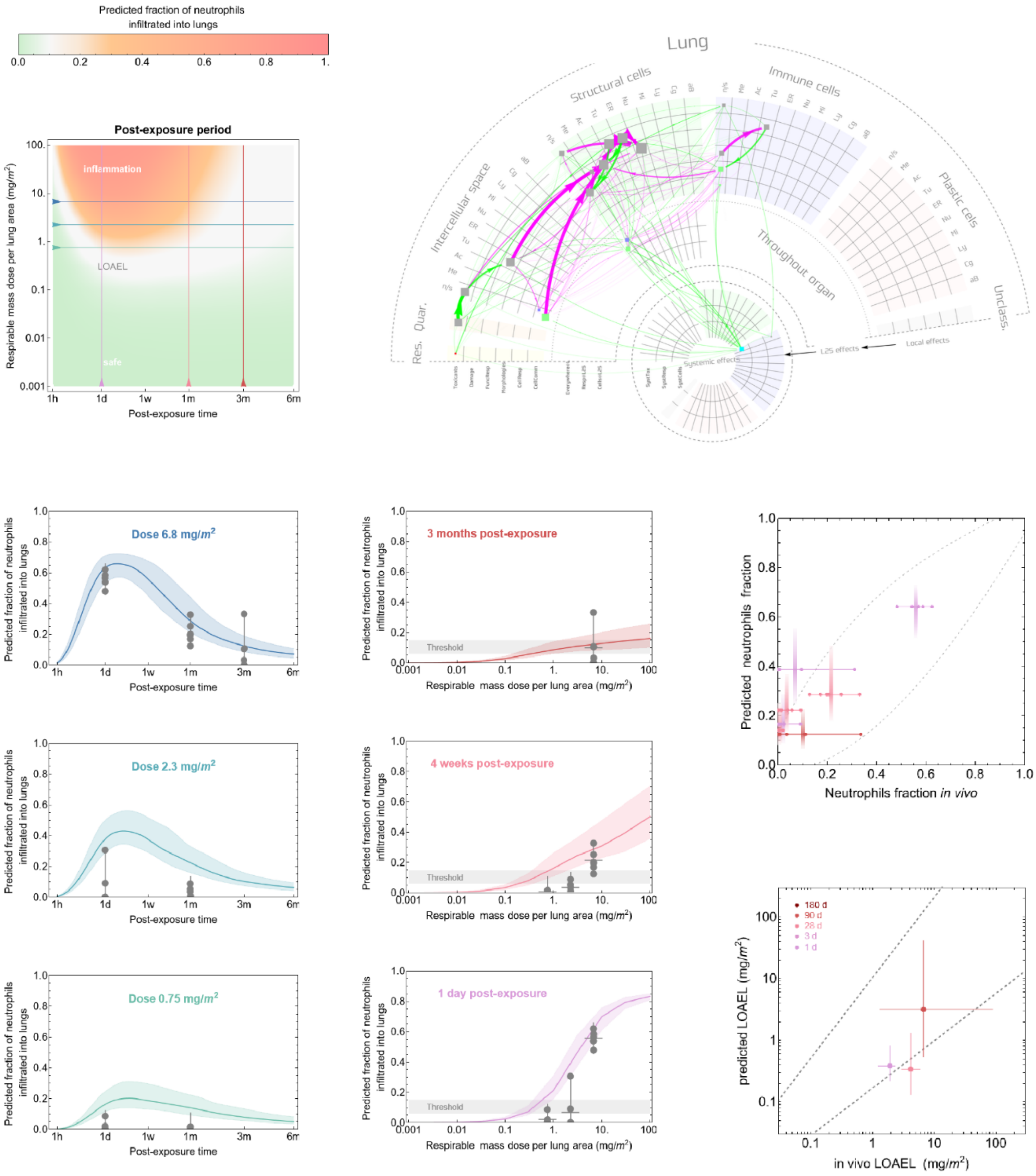

**Combustion 2**

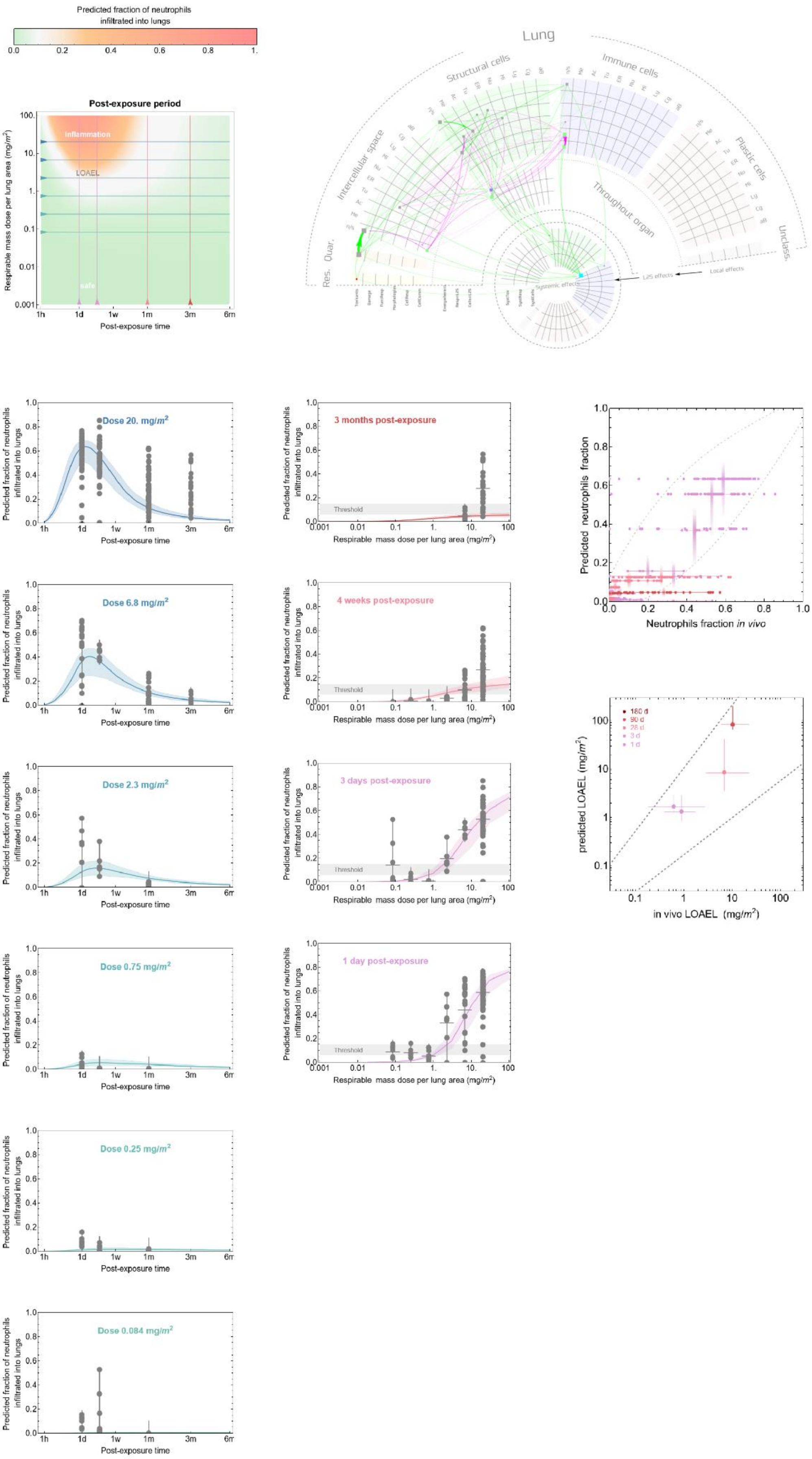

#### Combustion 3

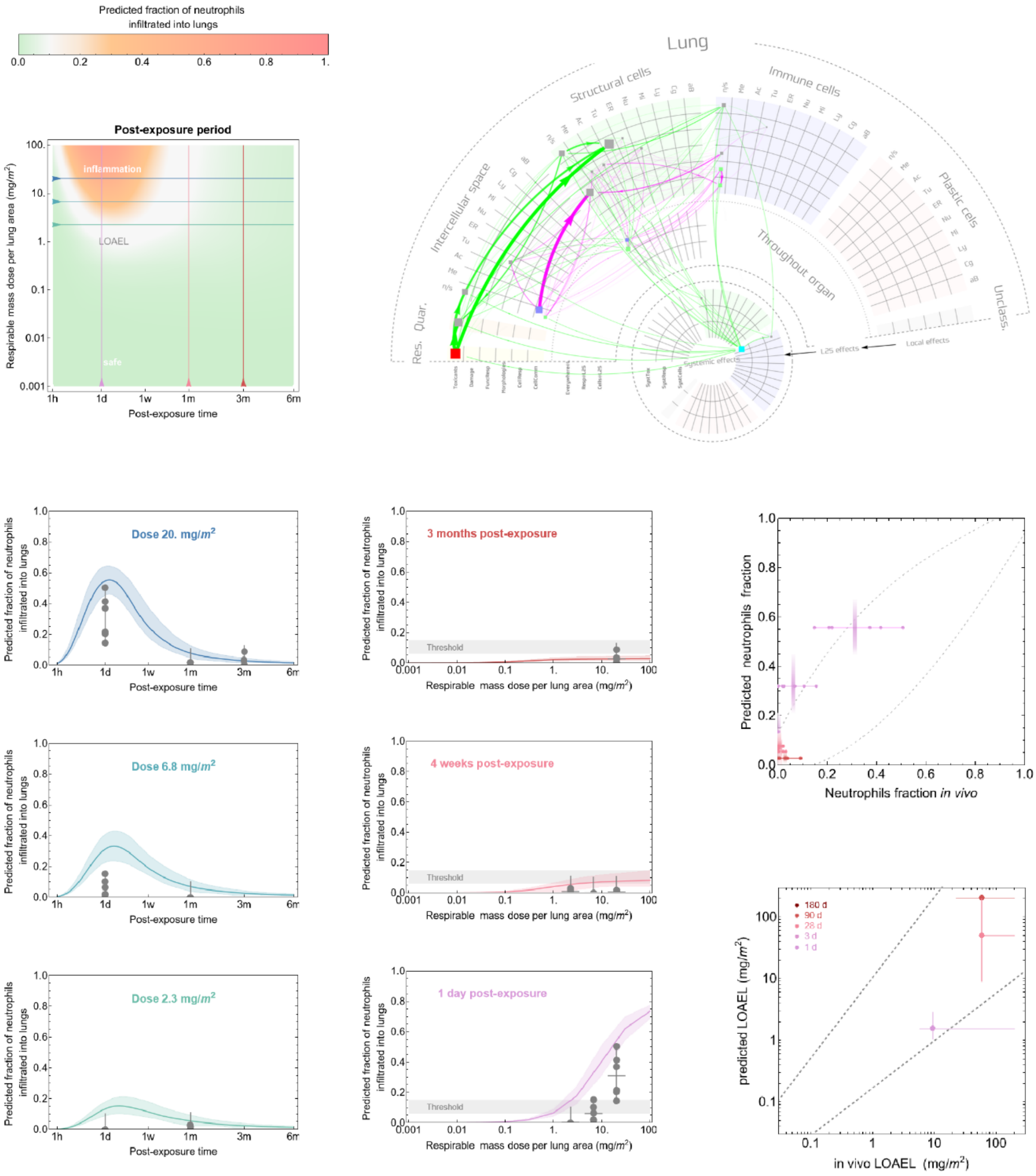

**Combustion 4**

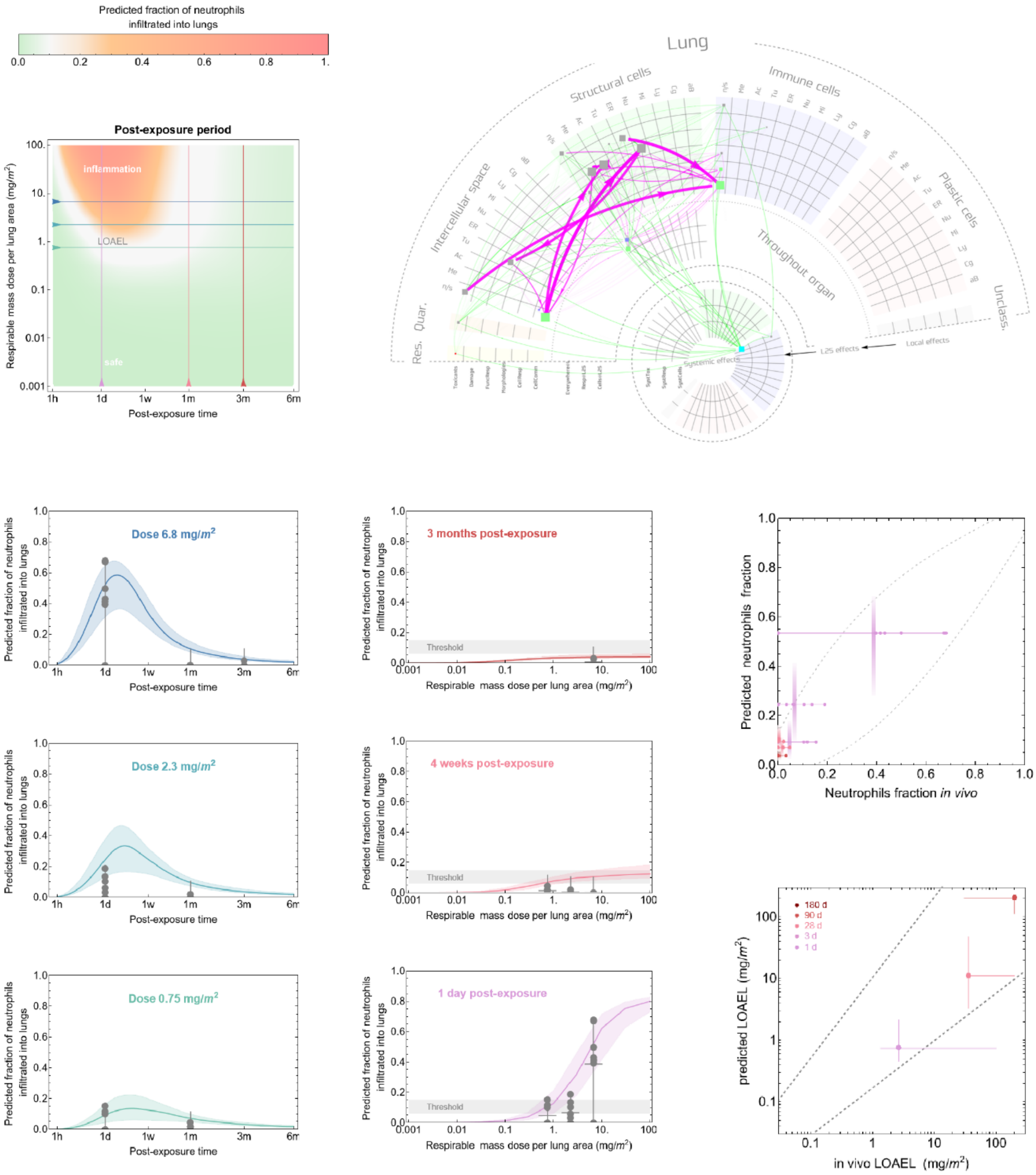

#### Combustion 5

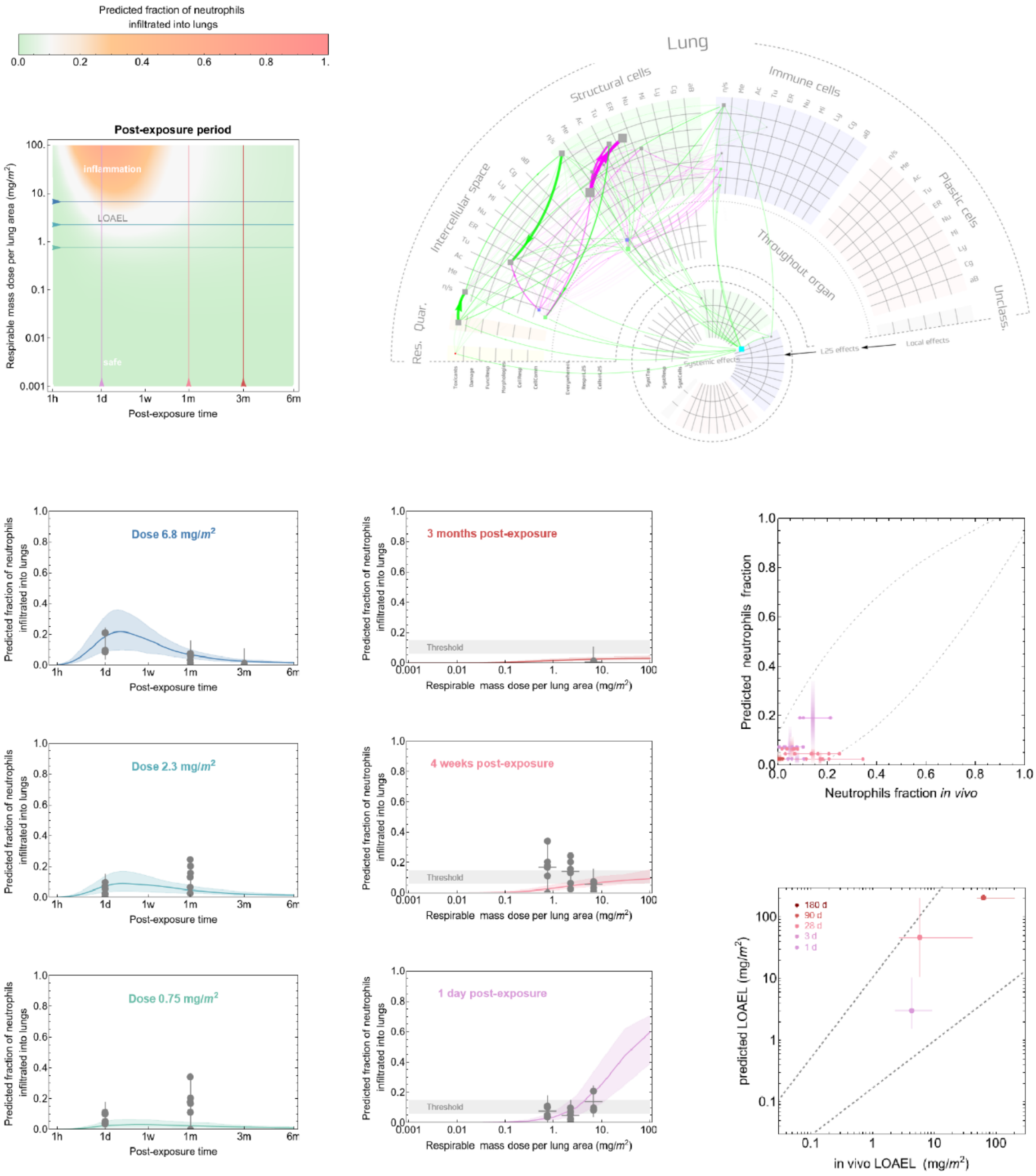

#### Combustion 6

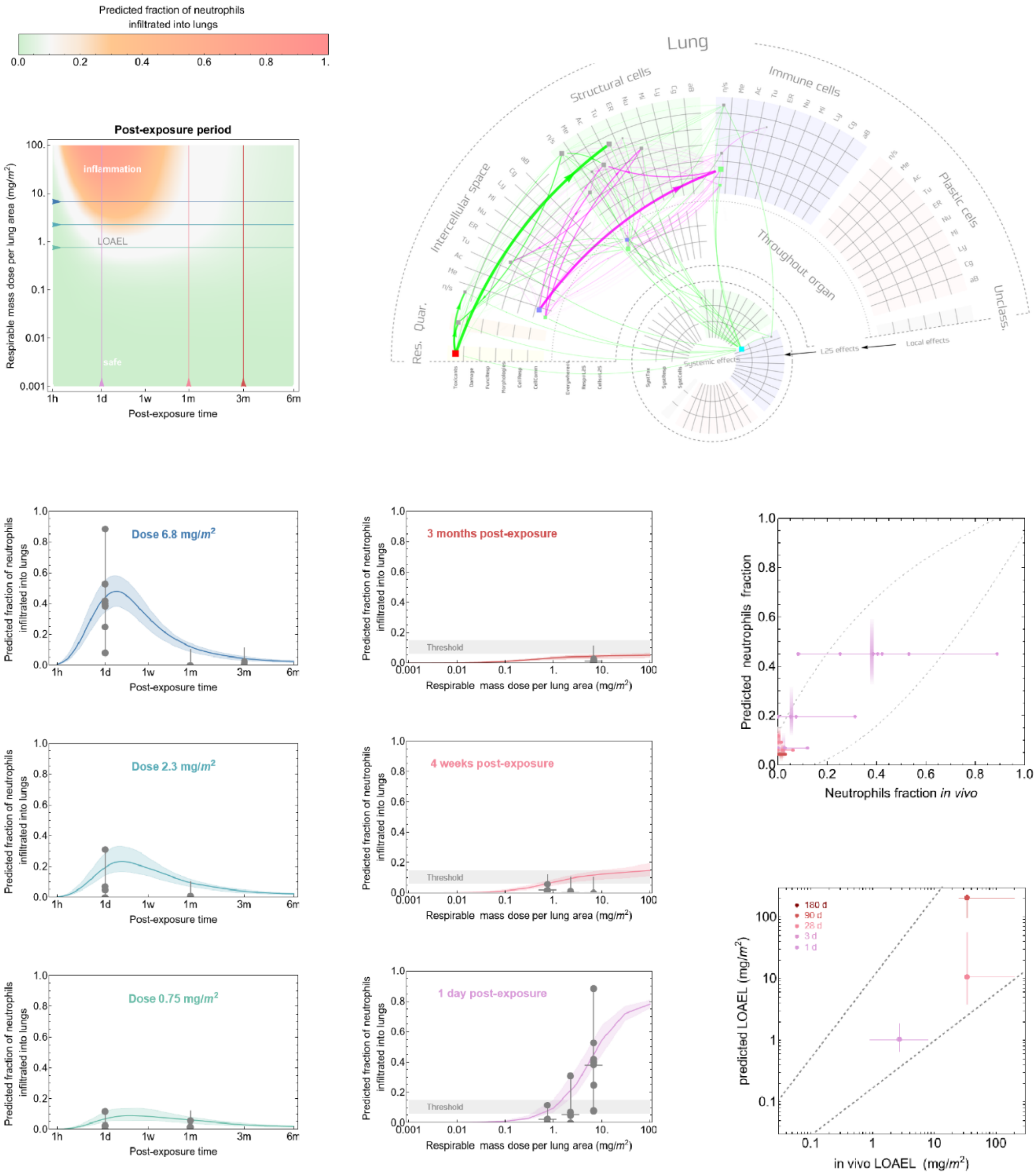

### Combustion 7

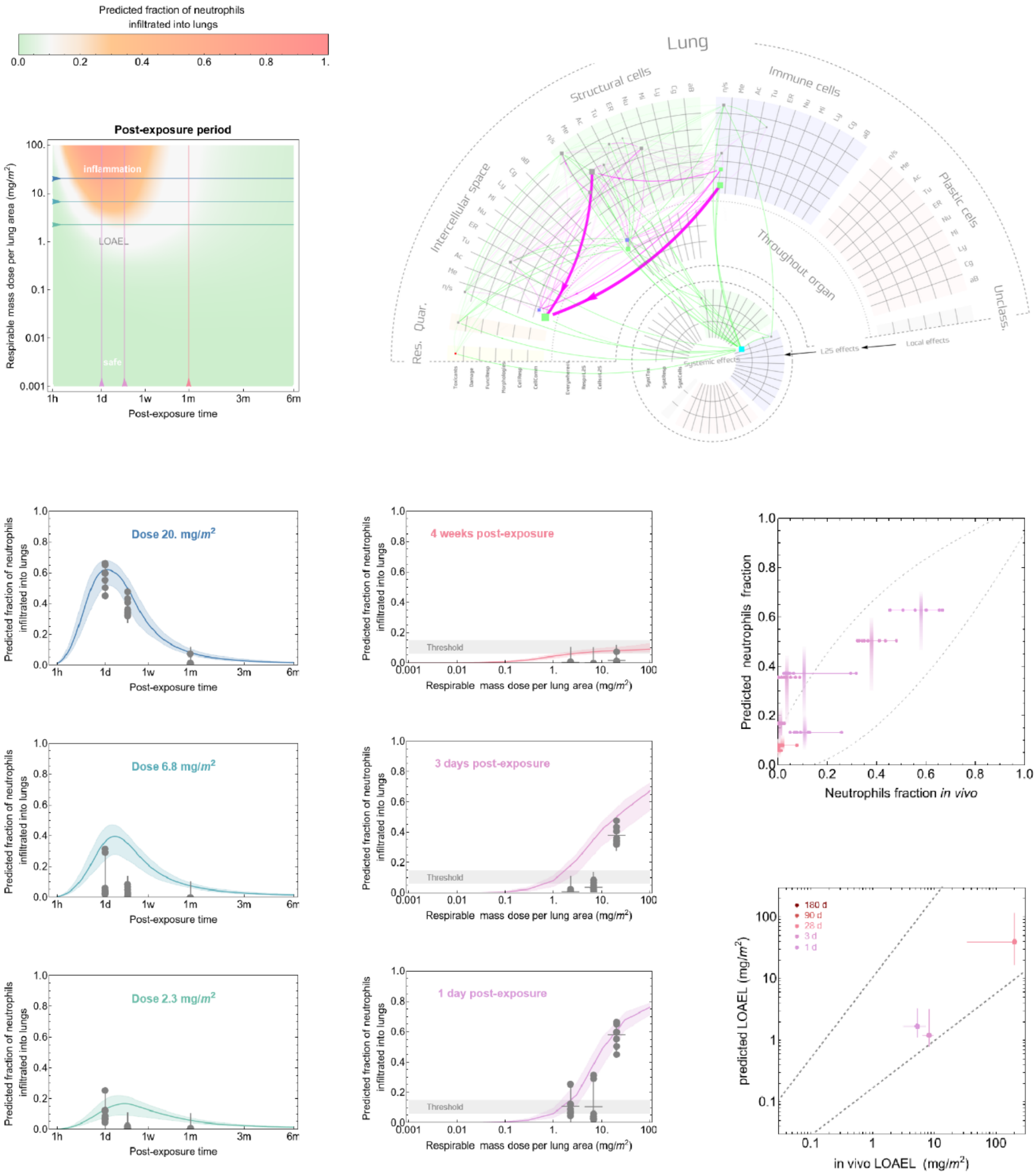

#### Industrial 1

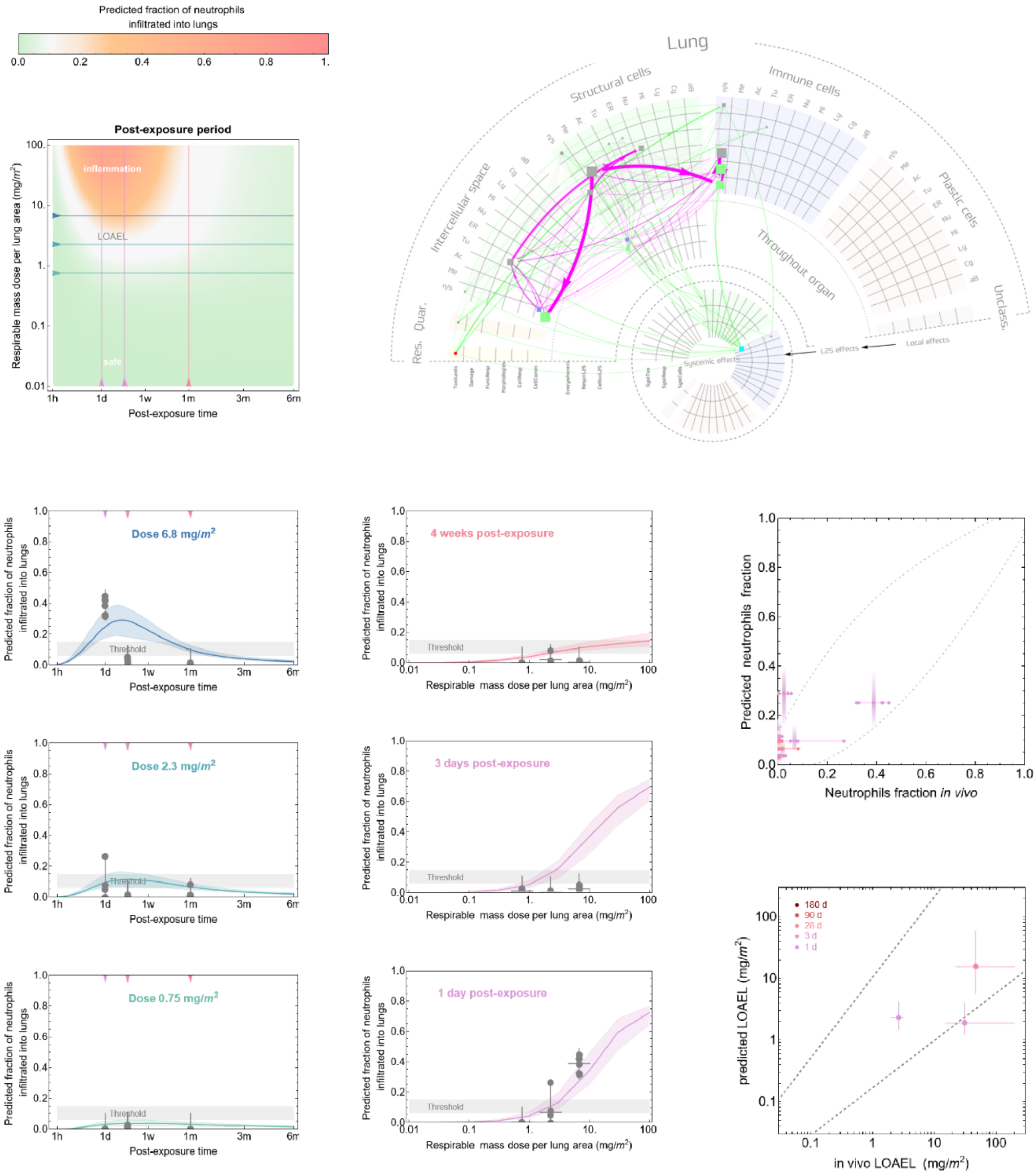

#### Industrial 2

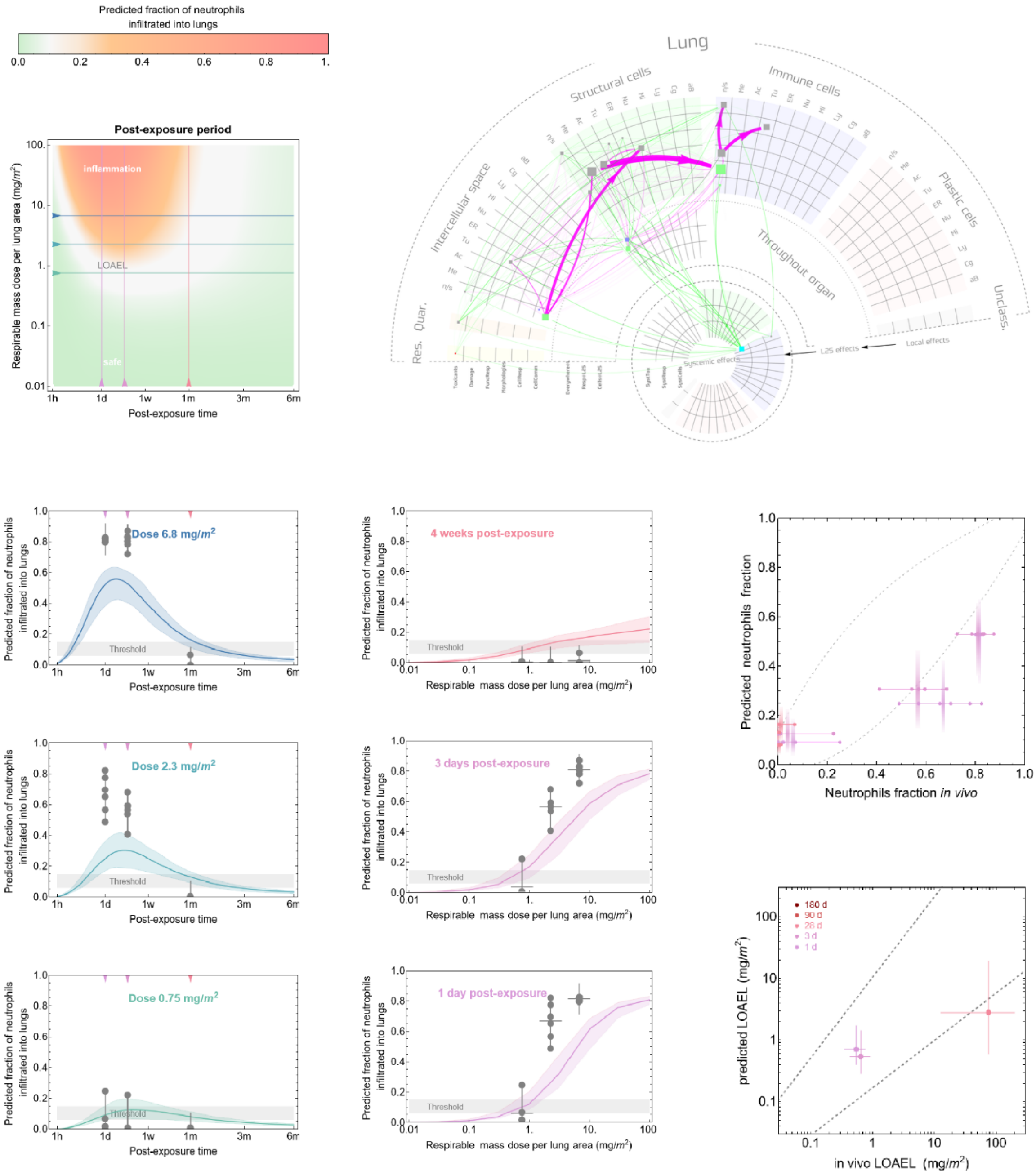

#### Industrial 3

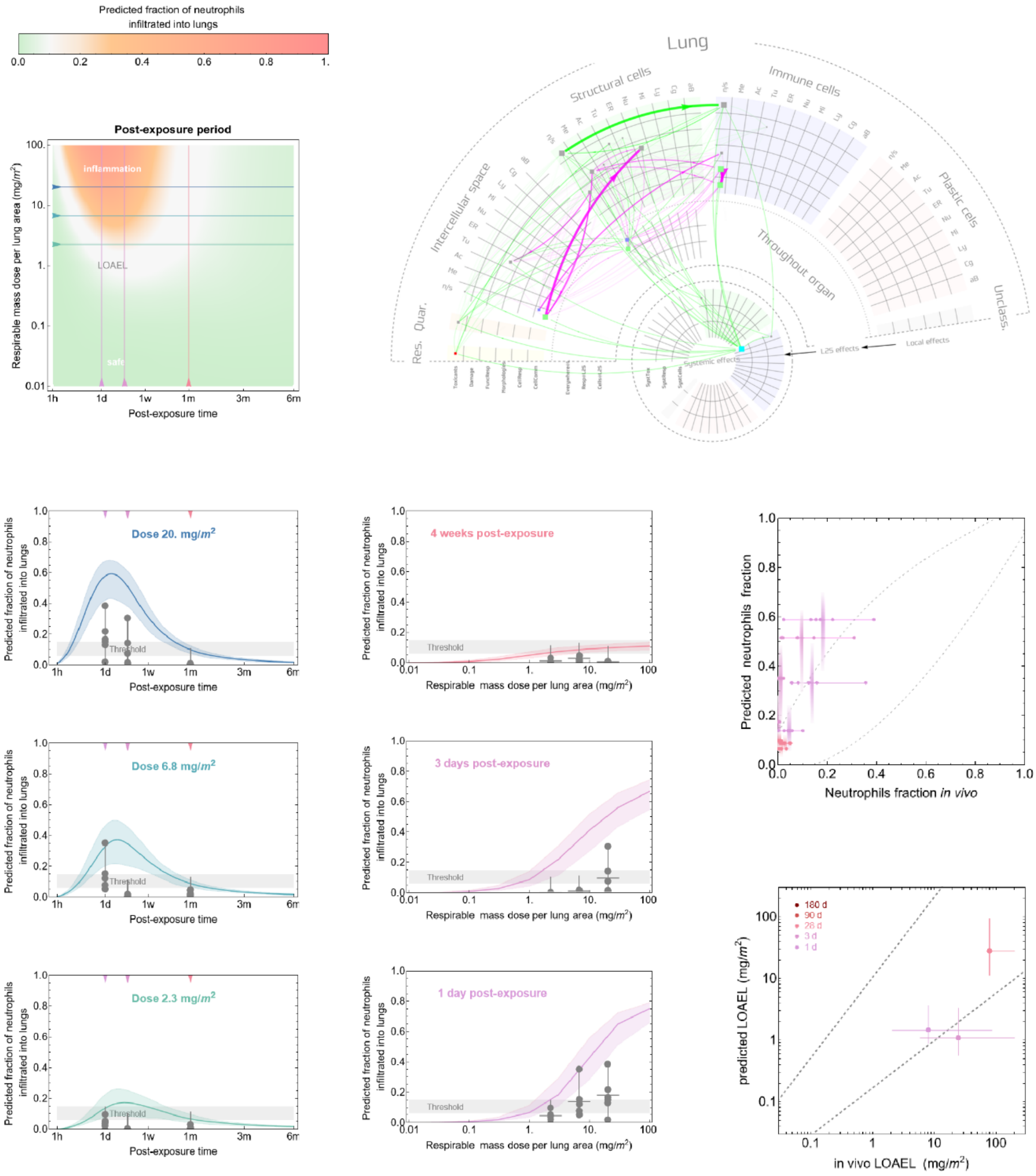

#### Industrial 4

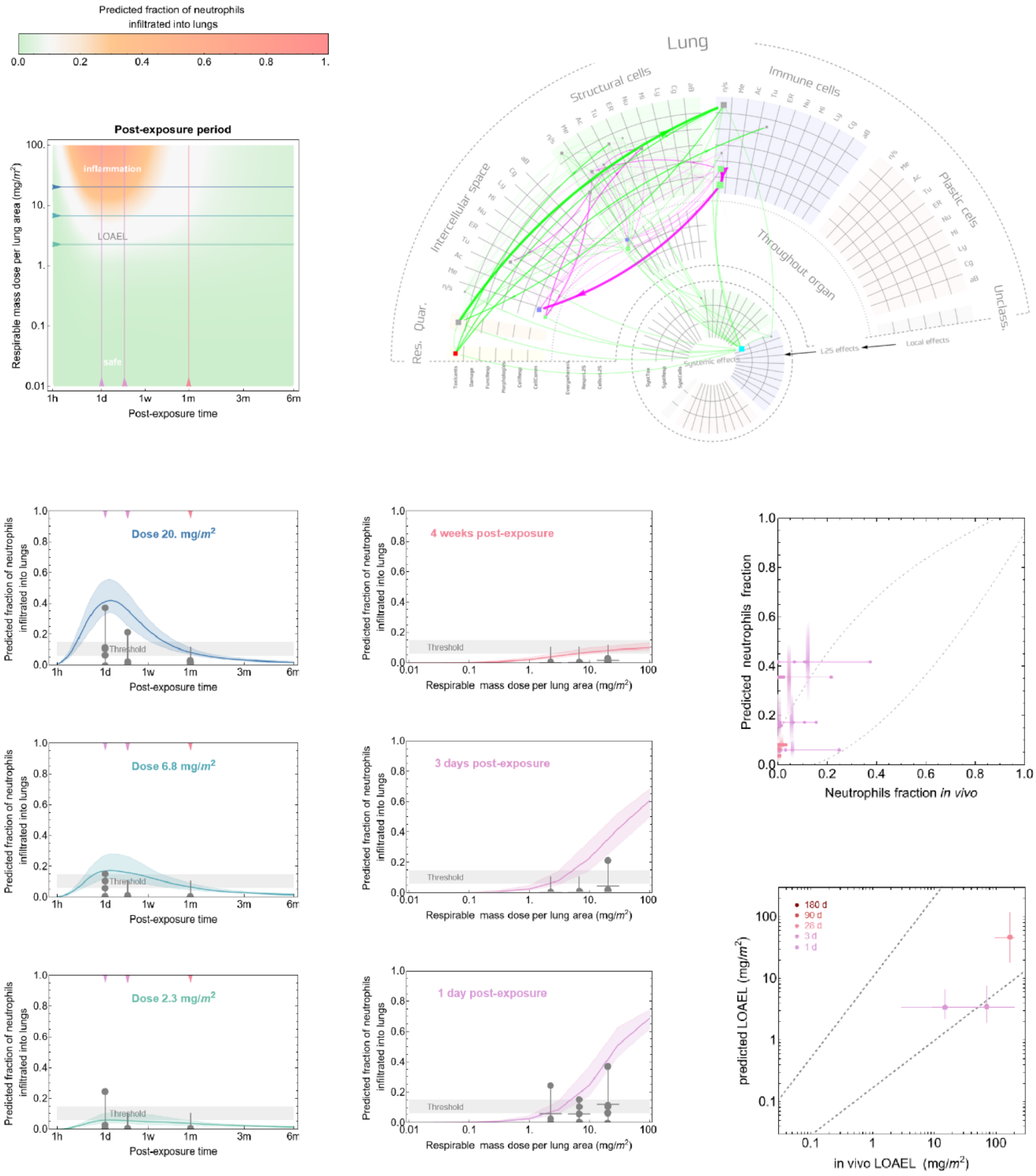

#### Industrial 5

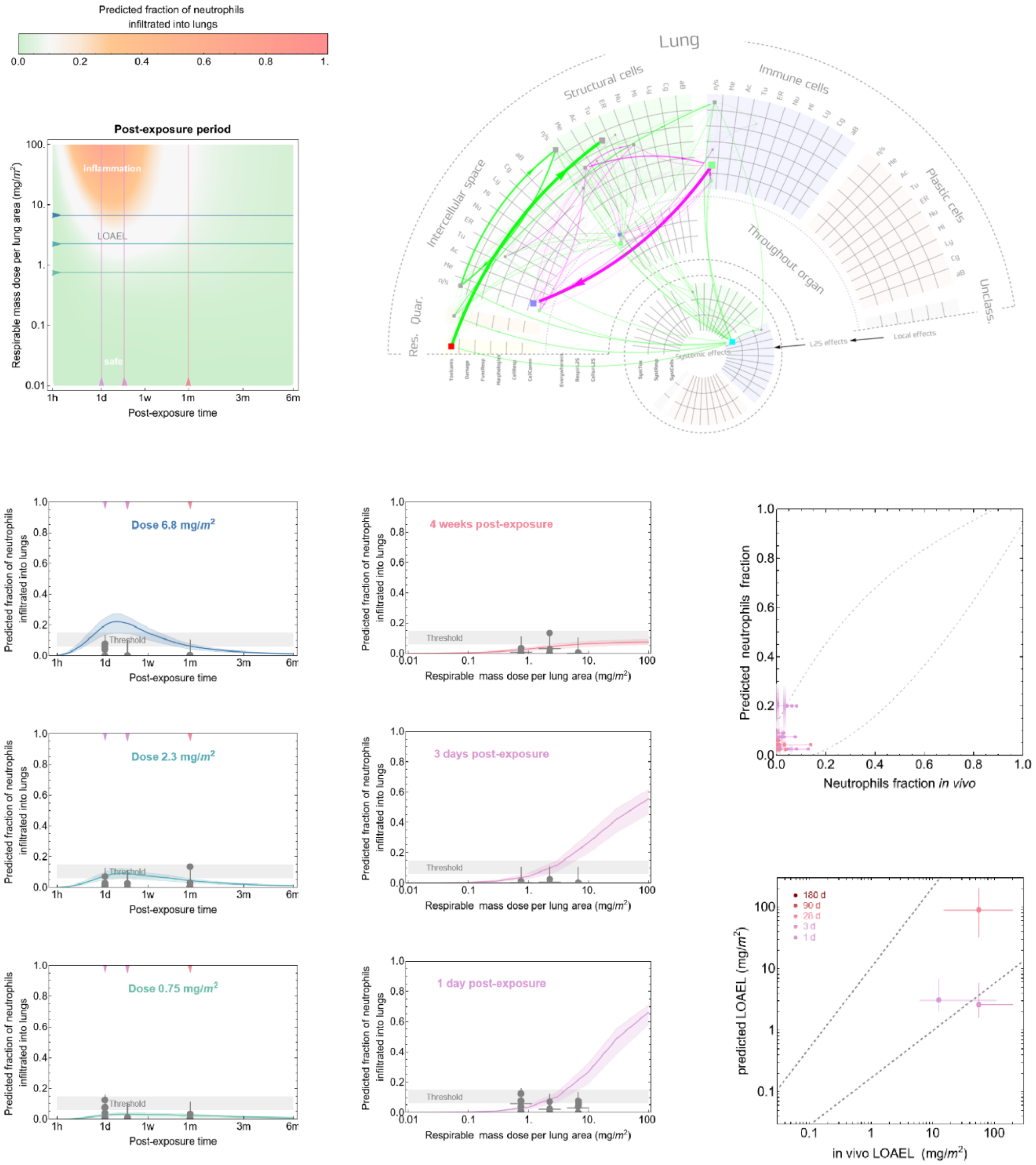

#### Metal oxide 1

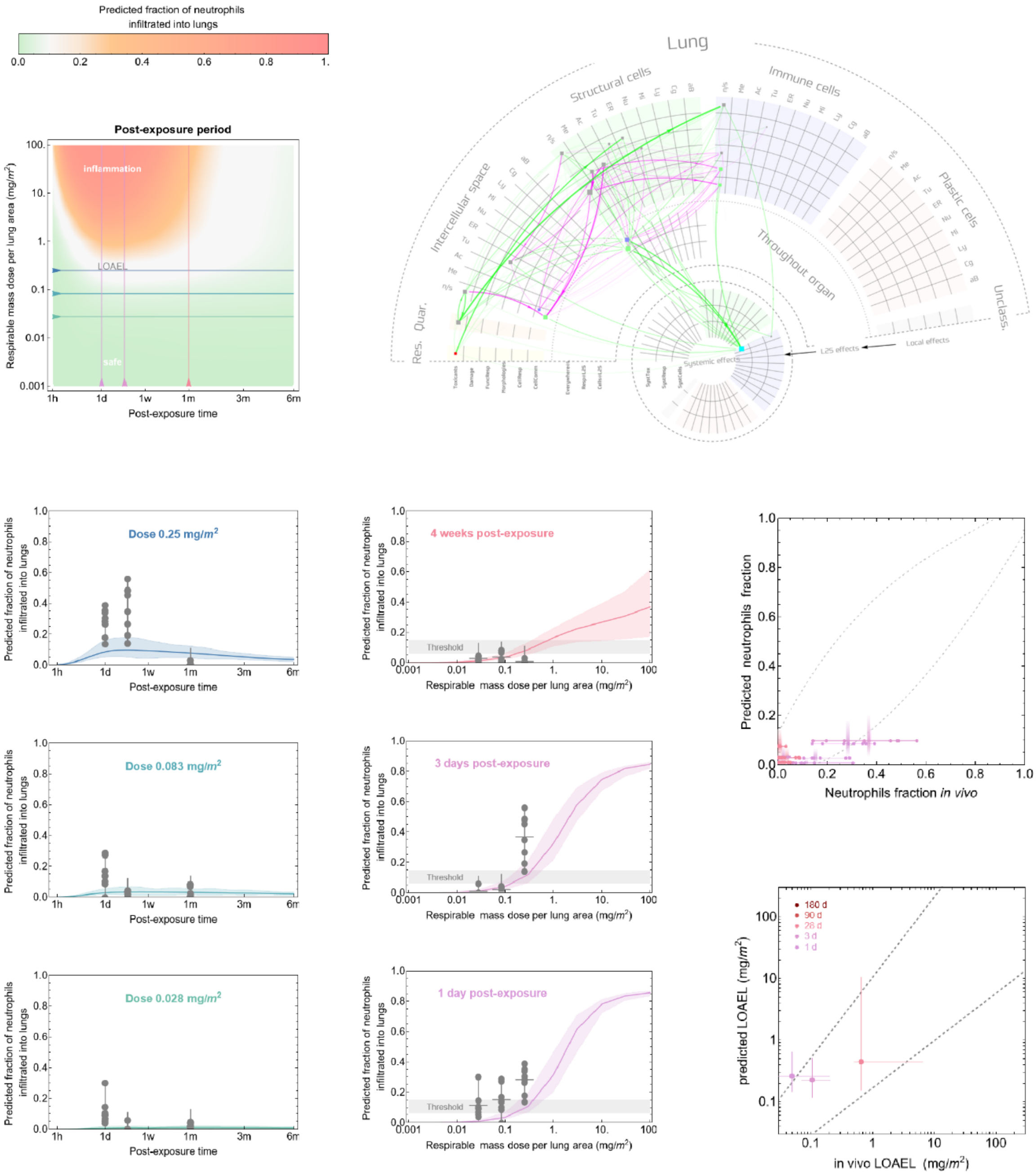

**Metal oxide 2**

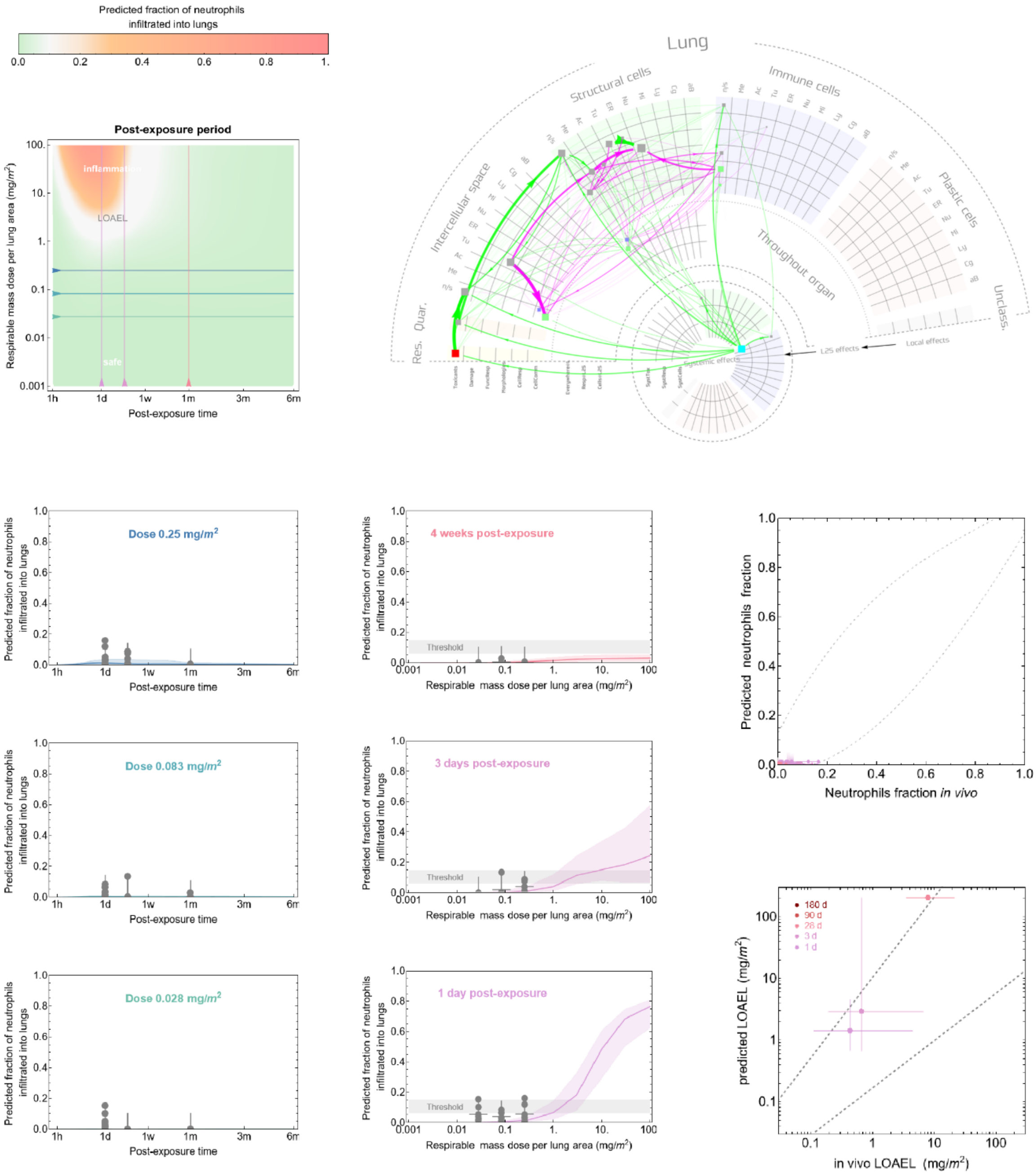

### Metal oxide 3

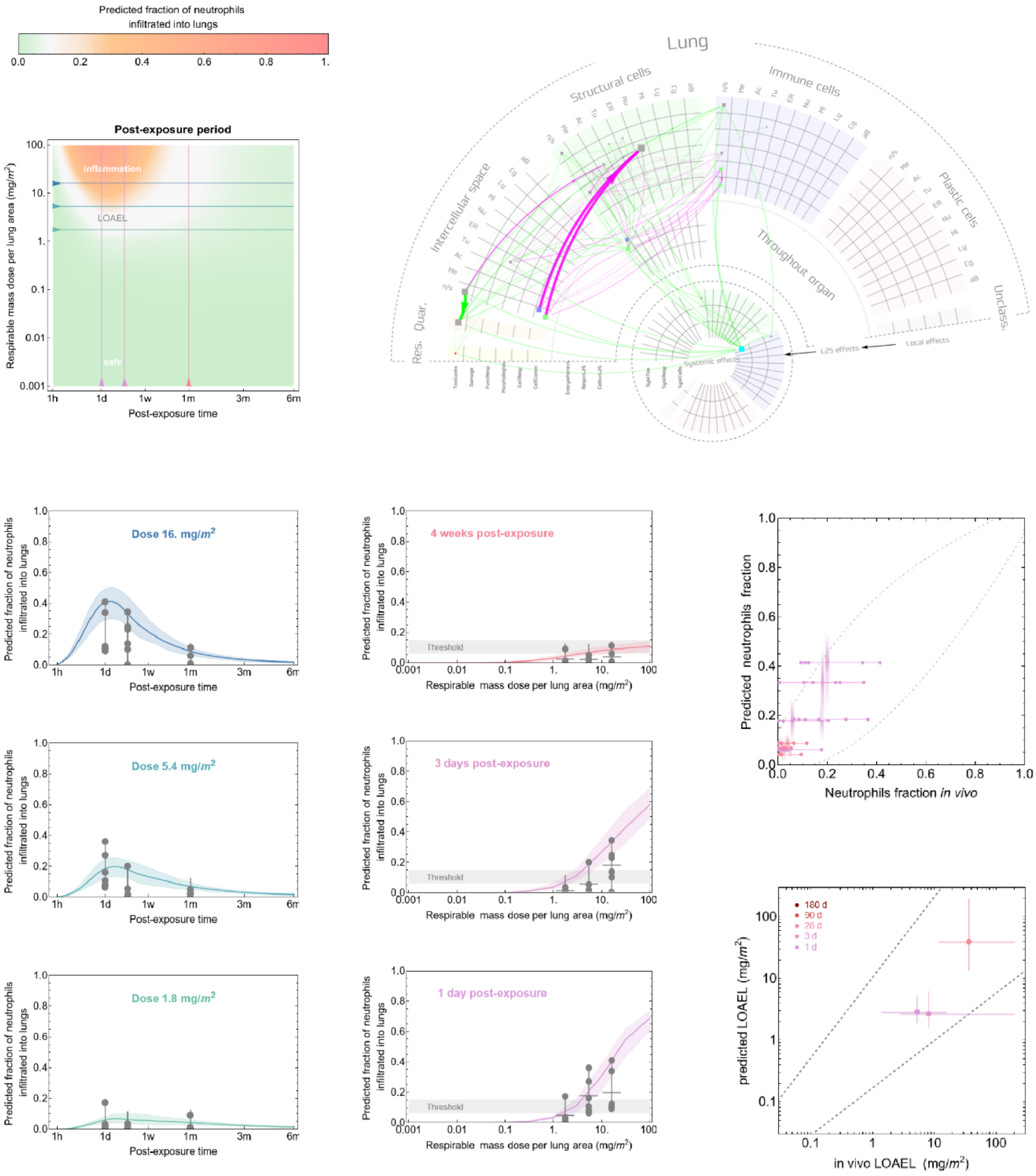

### Metal oxide 4

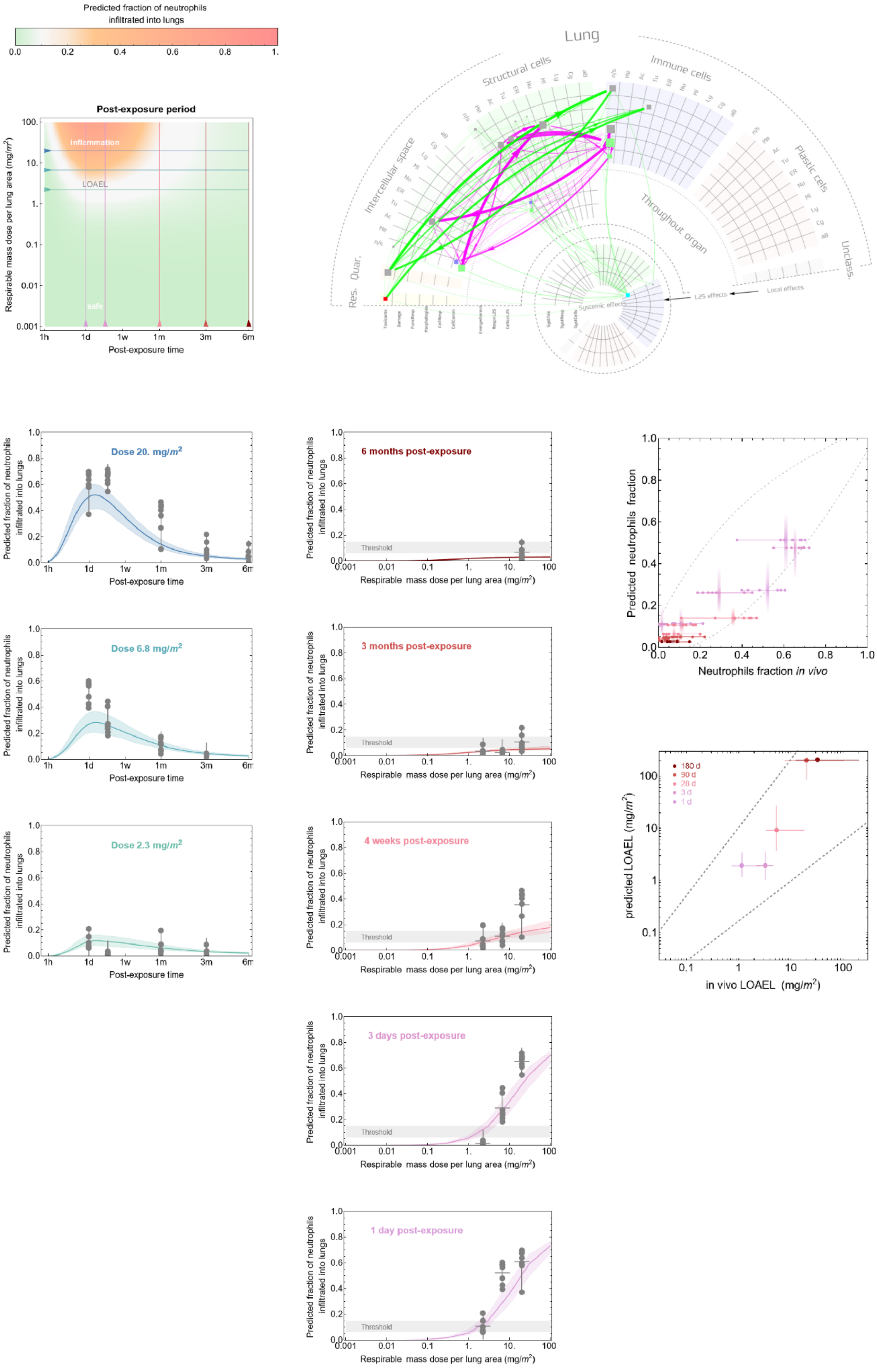

#### Metal oxide 5

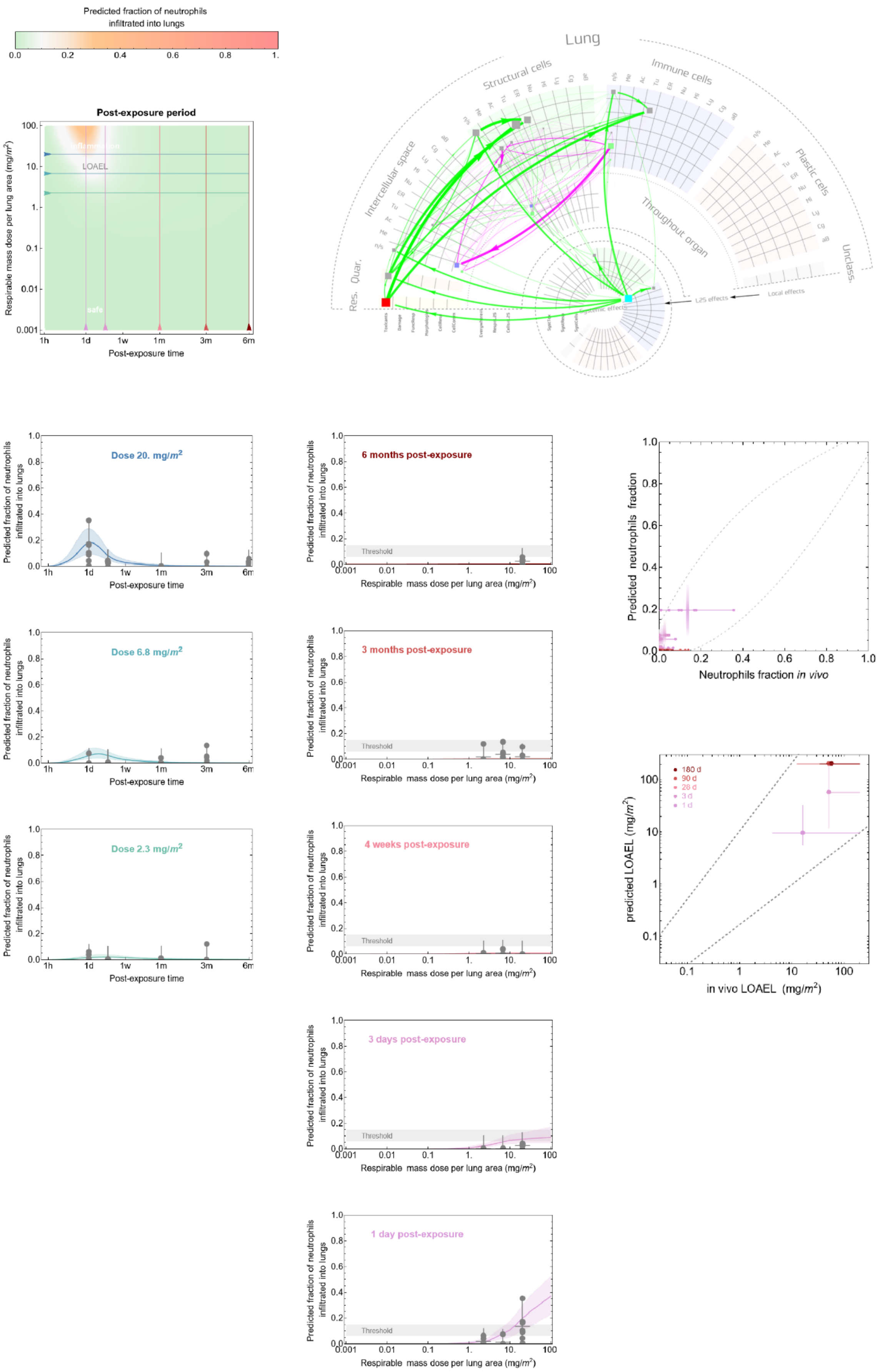

#### Metal oxide 6

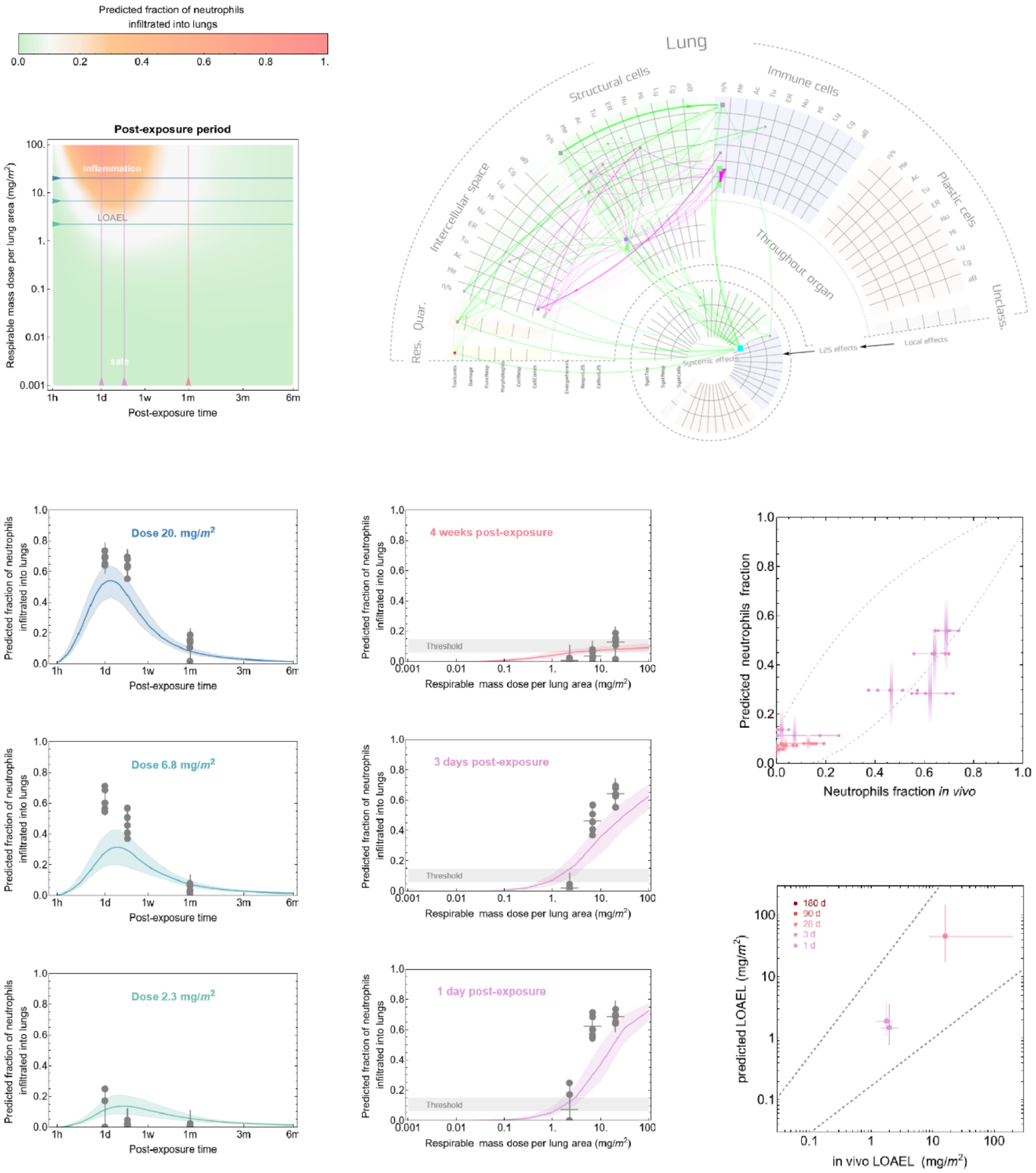

#### Metal oxide 7

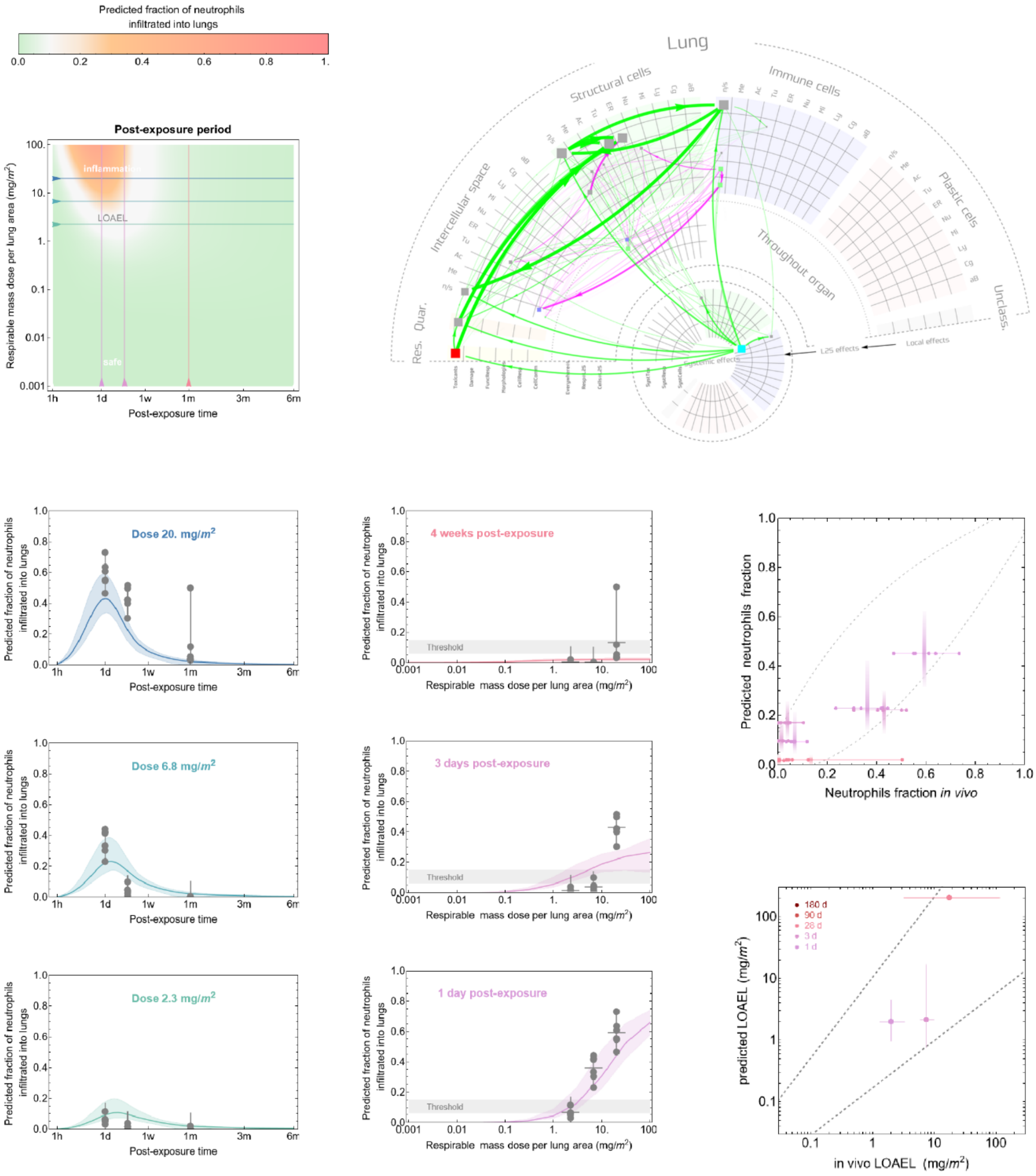

### Metal oxide 8

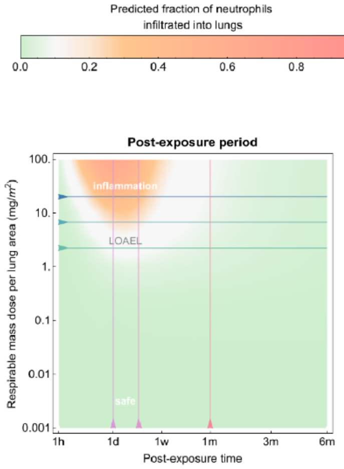

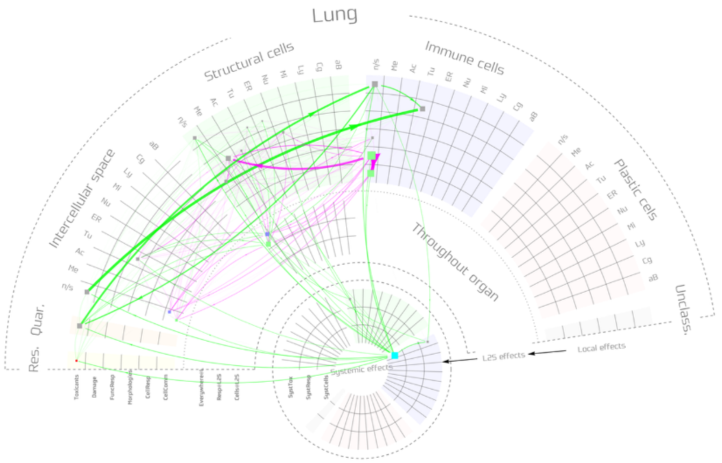

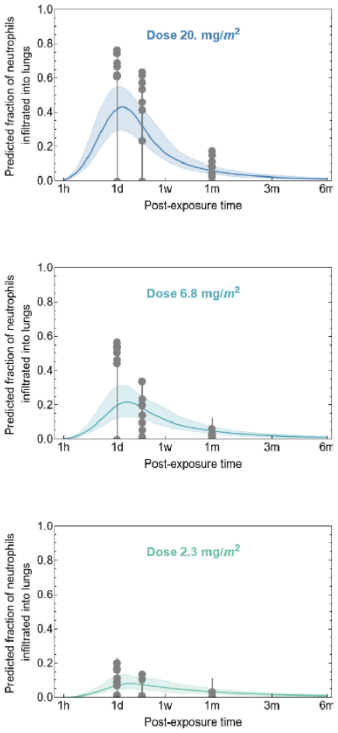

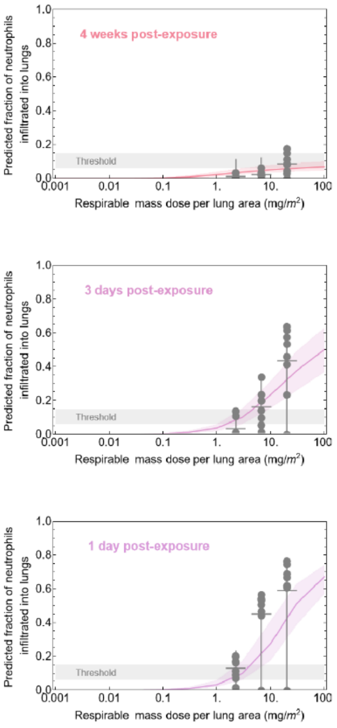

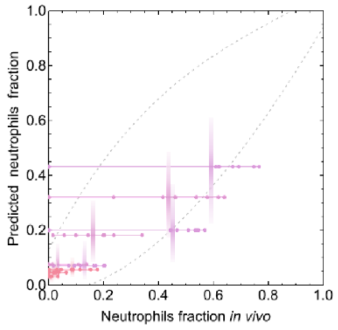

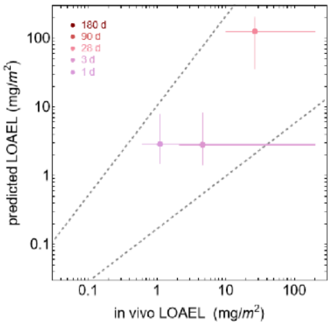

### Metal oxide 9

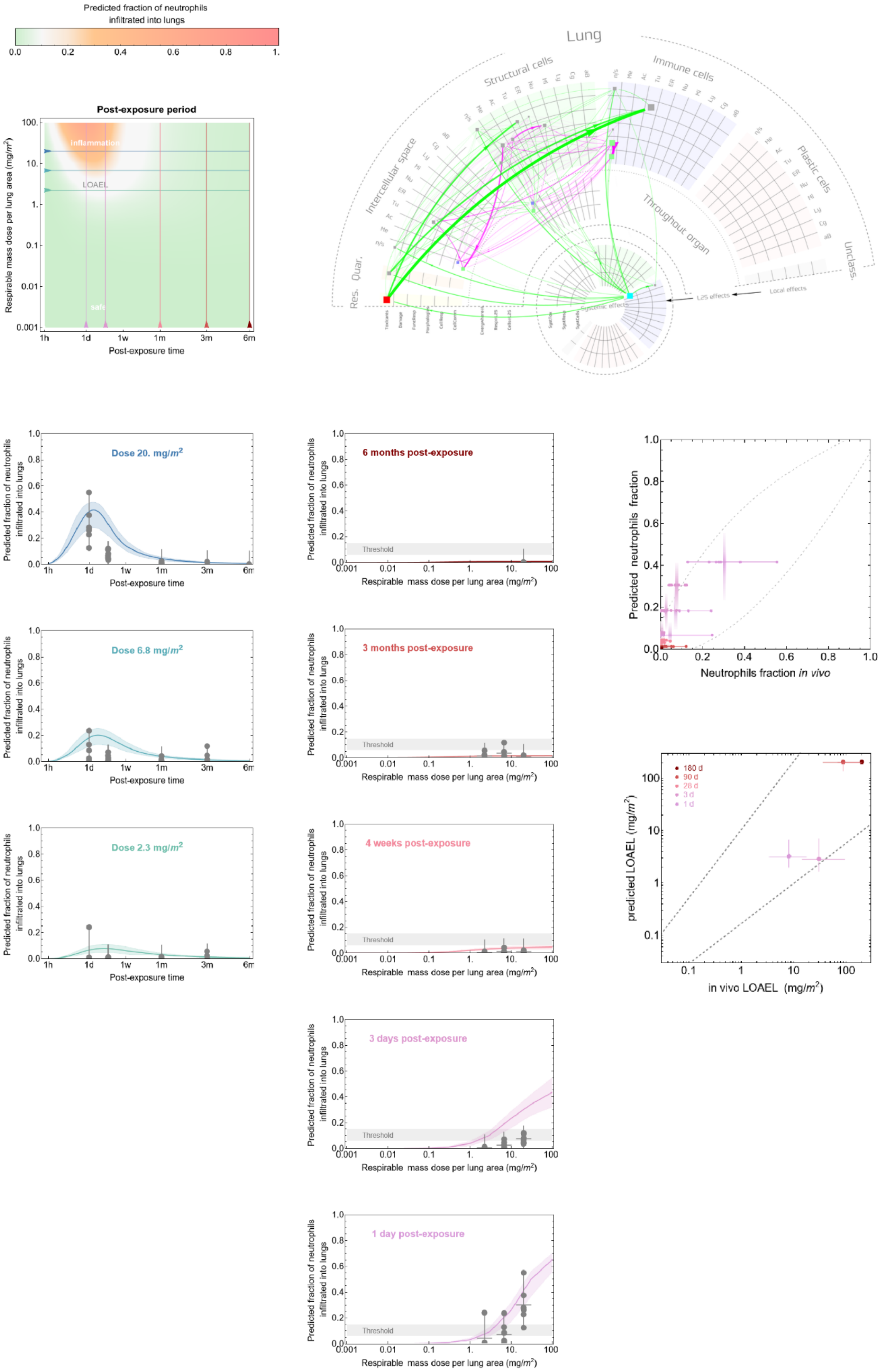

### Metal oxide 10

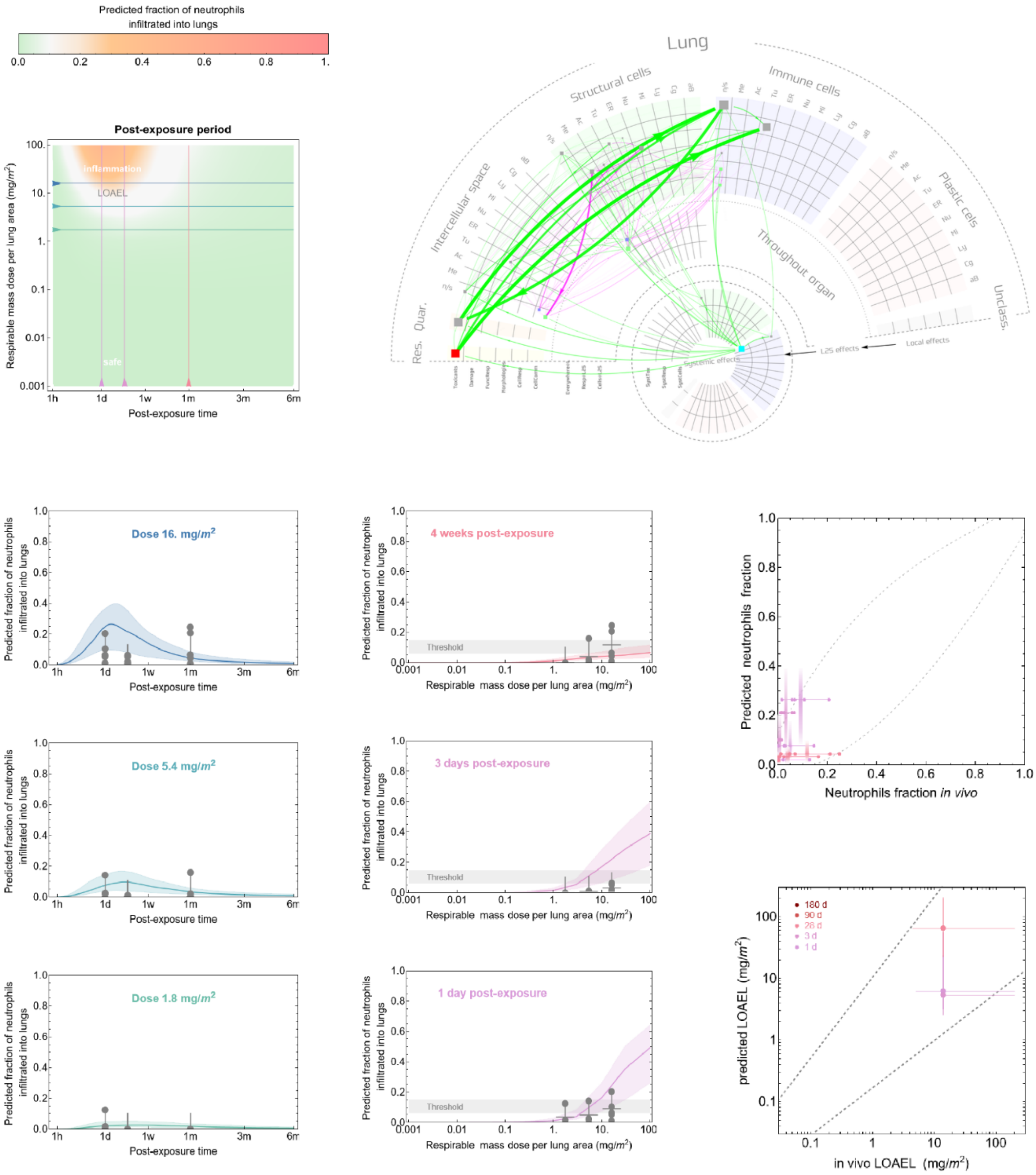

#### Metal oxide 11

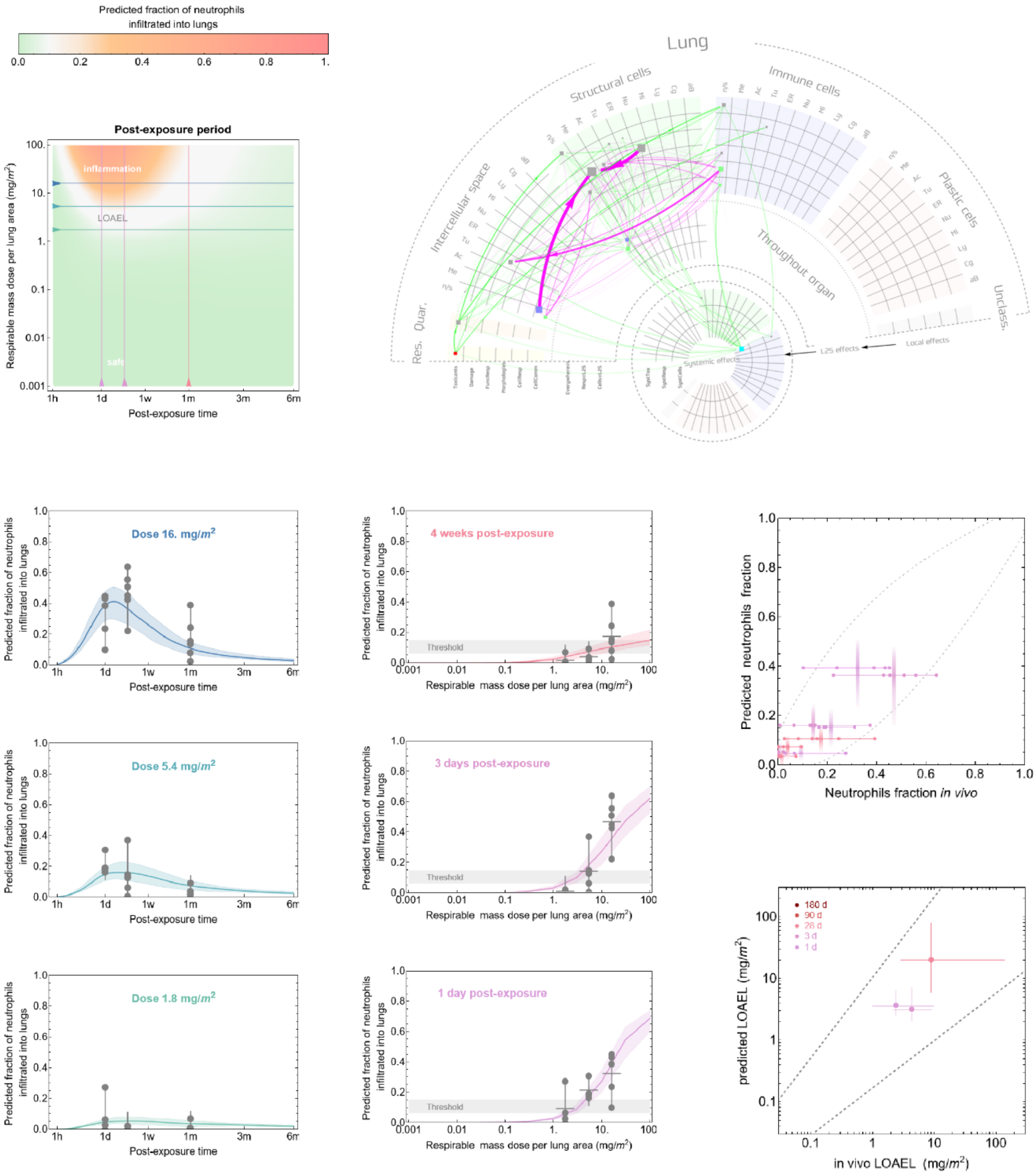

### Metal oxide 12

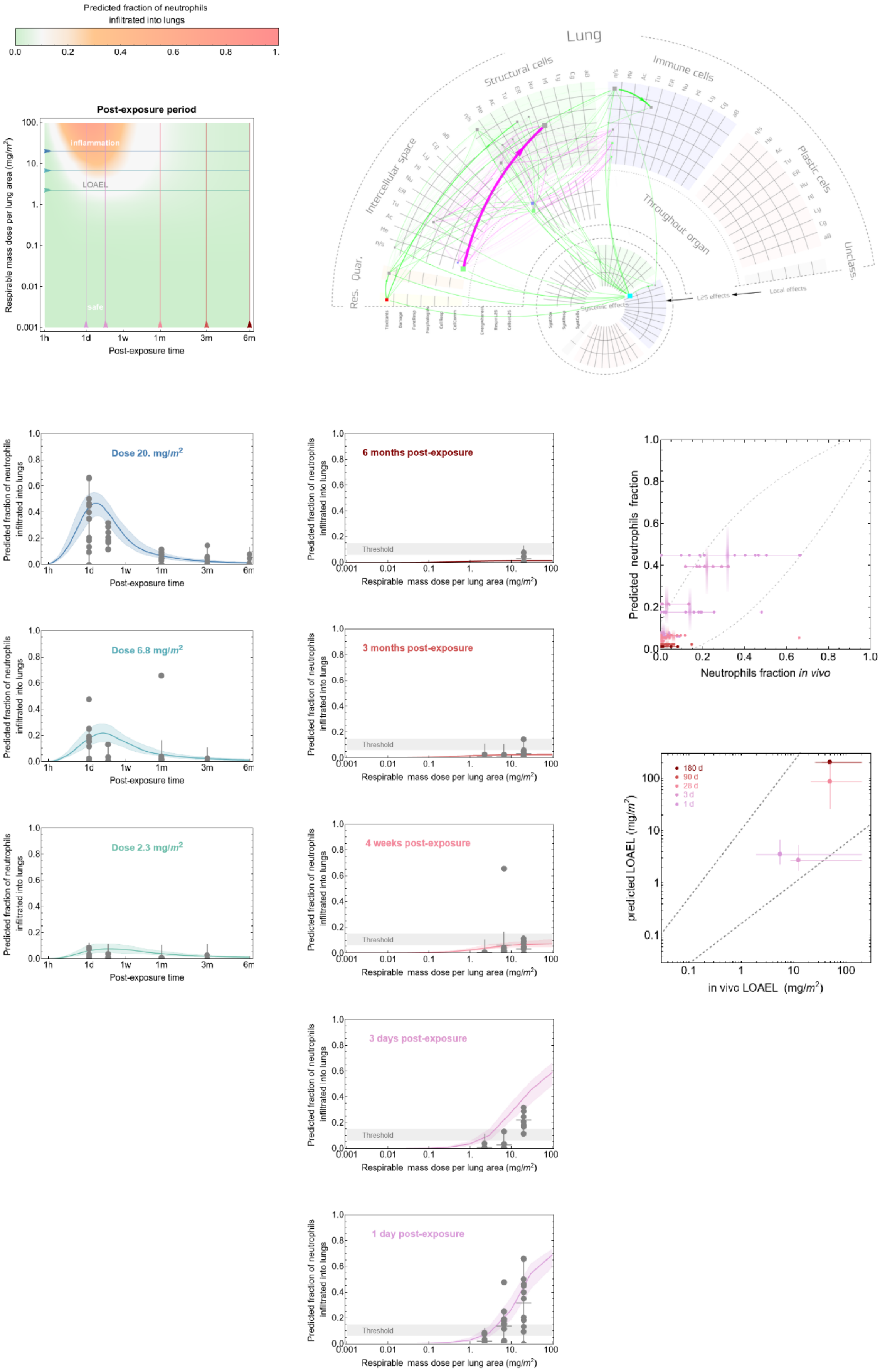

### SI-3aTime-course and dose-response predictions for inhalation exposure versus in vivo (rat) data – all 7 materials

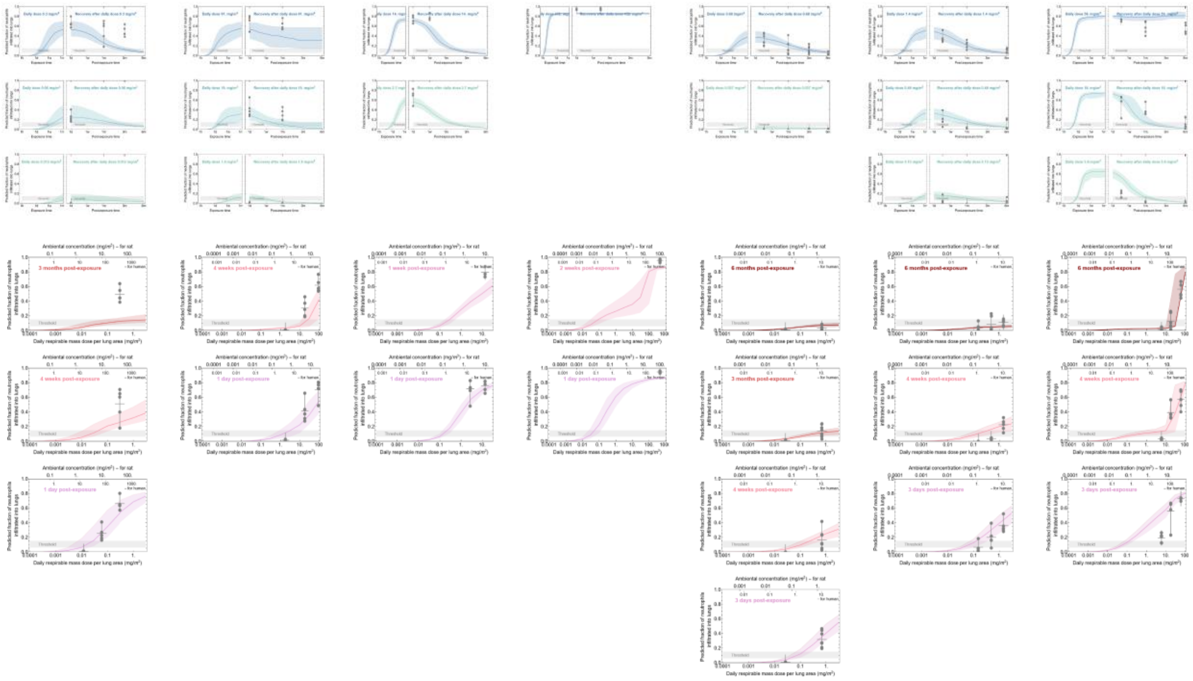

*FigureS 3. Predictions of time dependent (blue lines) and dose dependent (red lines) neutrophil influx versus inhalation exposure in vivo (rat) data points for all 7 individual materials test according to OECD test guidelines with all in vivo data points (grey circles). For higher resolution images see supplemental files: inhalation_time-dependent_HiRes_plots.zip, and inhallation_dose-dependent_HiRes_plots.zip.*

### SI-3b Time-course and dose-response predictions for inhalation exposure versus in vivo (rat) data – per material

**Carbon 17**

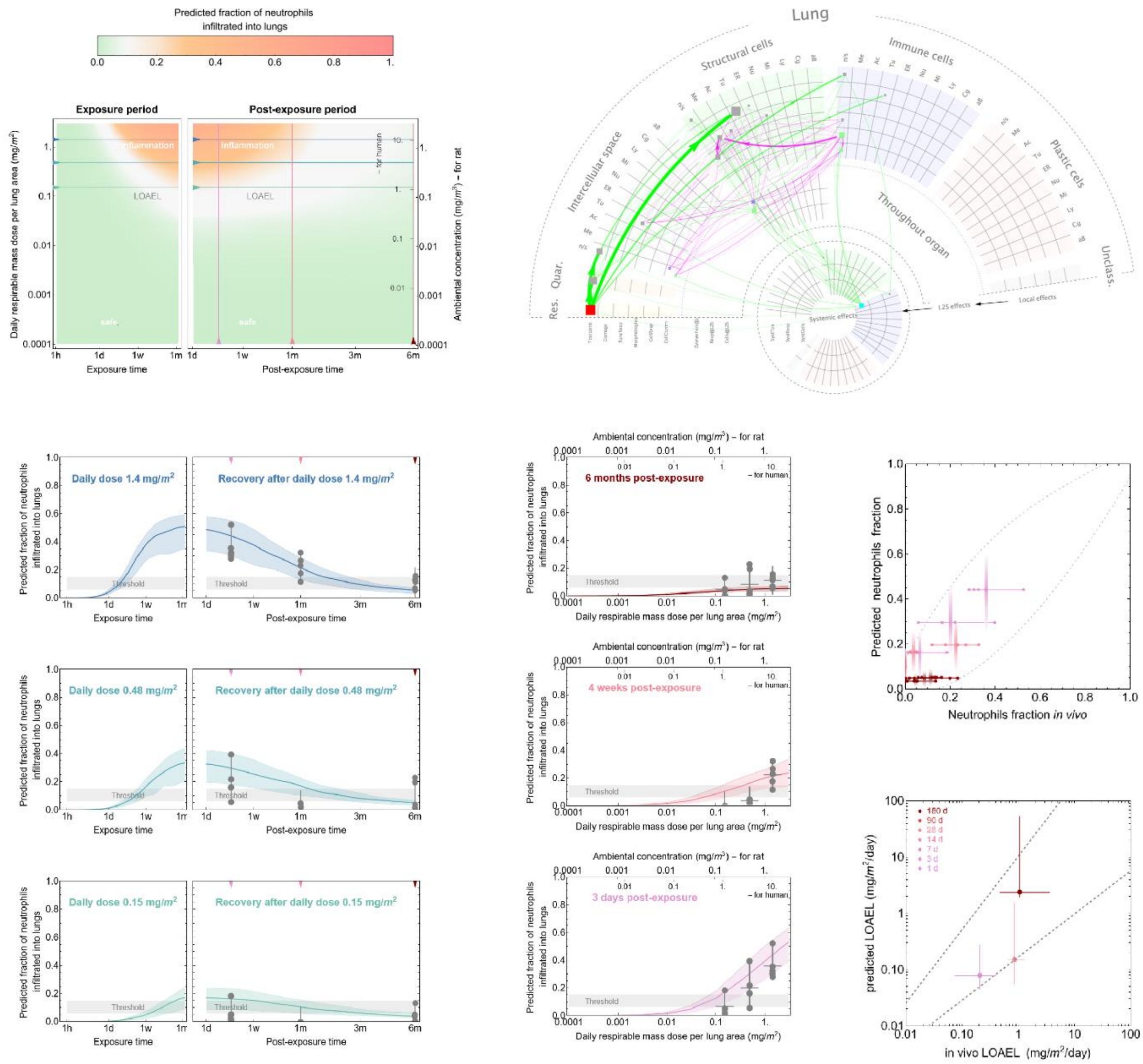

#### Carbon 18

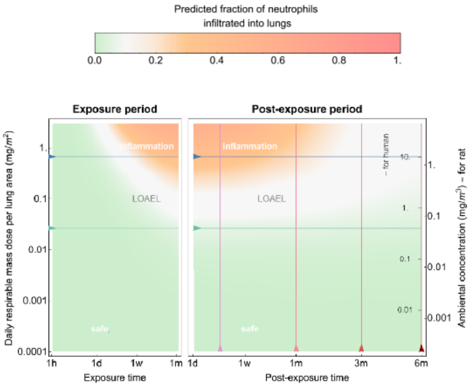
Predicted fraction of neutrophils infiltrated into lungs
0.0 0.2 0.4 0.6 0.8 1.
Exposure period
Post-exposure period
inflammation
LOAEL
safe
Daily respirable mass dose per lung area (mg/m²)
Ambiental concentration (mg/m³) – for rat
– for human
Exposure time
Post-exposure time

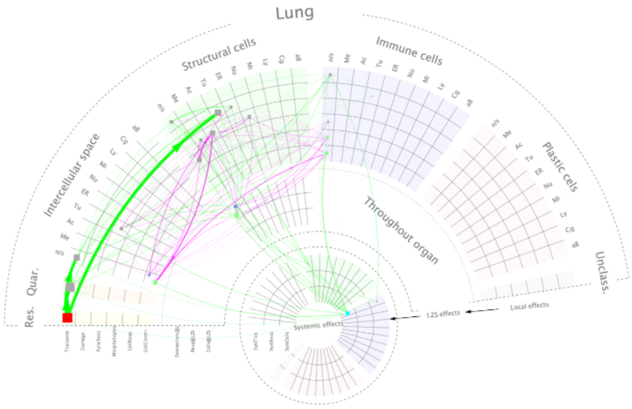
Lung
Structural cells
Immune cells
Plastic cells
Intercellular space
Throughout organ
Unclass.
Quar.
Res.
Systemic effects
Local effects

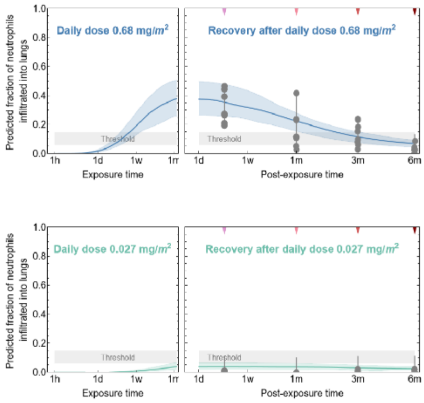
Daily dose 0.68 mg/m²
Recovery after daily dose 0.68 mg/m²
Daily dose 0.027 mg/m²
Recovery after daily dose 0.027 mg/m²
Threshold
Predicted fraction of neutrophils infiltrated into lungs
Exposure time
Post-exposure time

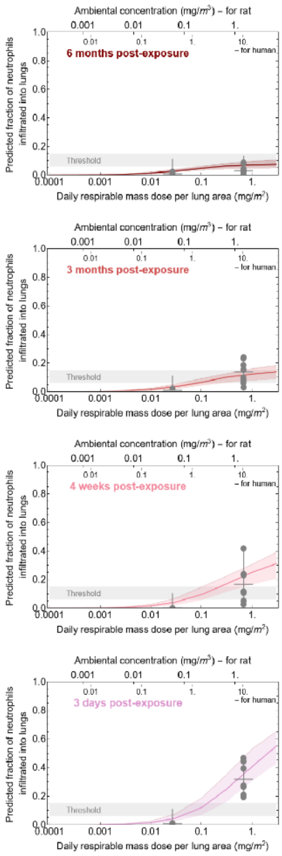
Ambiental concentration (mg/m³) – for rat
– for human
6 months post-exposure
3 months post-exposure
4 weeks post-exposure
3 days post-exposure
Threshold
Predicted fraction of neutrophils infiltrated into lungs
Daily respirable mass dose per lung area (mg/m²)

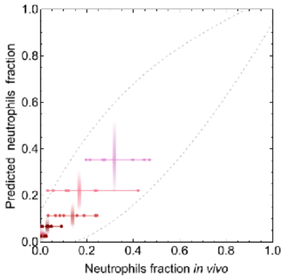
Predicted neutrophils fraction
Neutrophils fraction in vivo

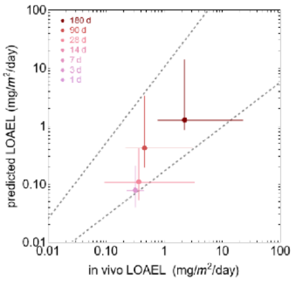
180 d
90 d
28 d
14 d
7 d
3 d
1 d
predicted LOAEL (mg/m²/day)
in vivo LOAEL (mg/m²/day)

**Combustion 2**

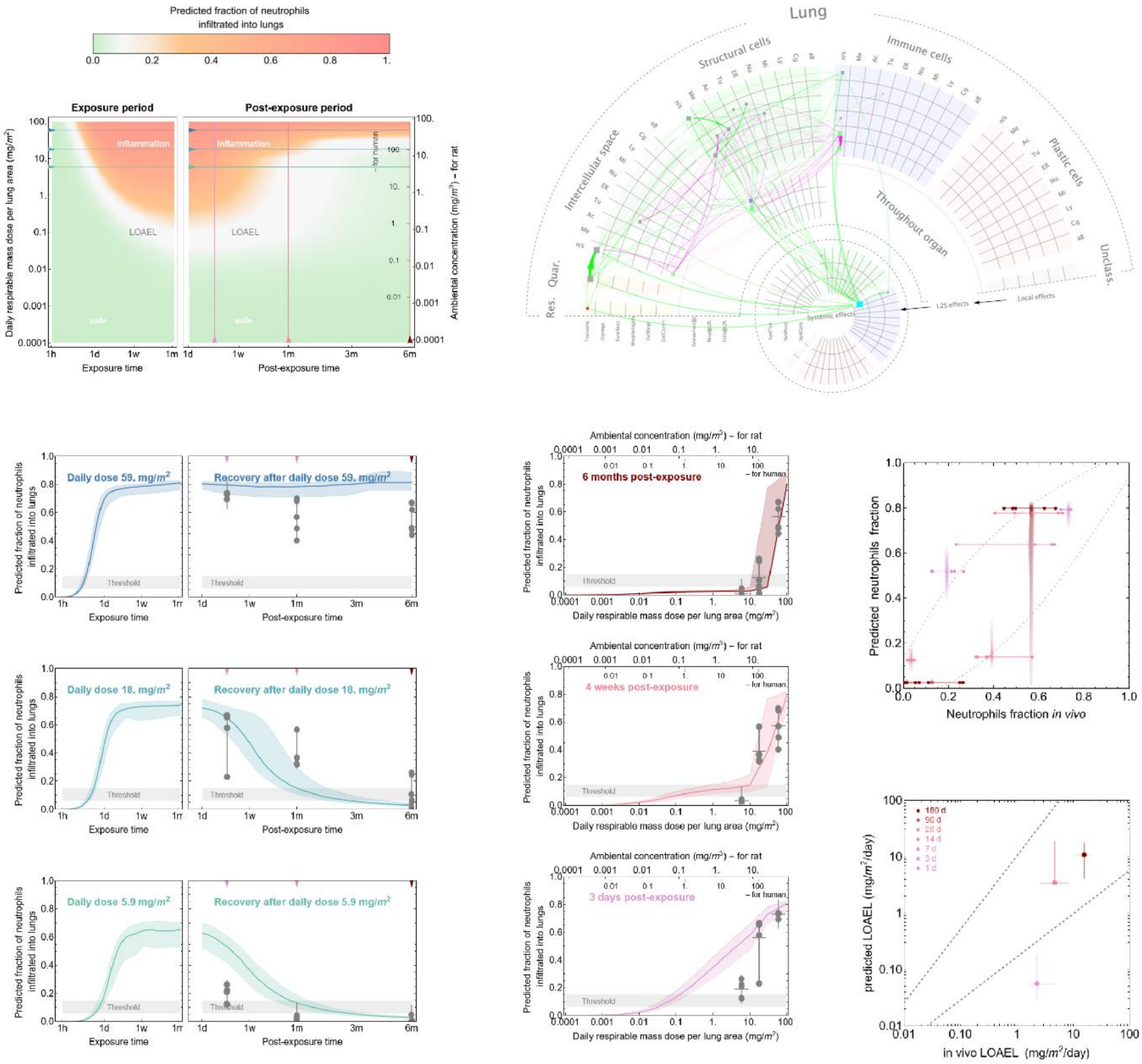

**Industrial 6**

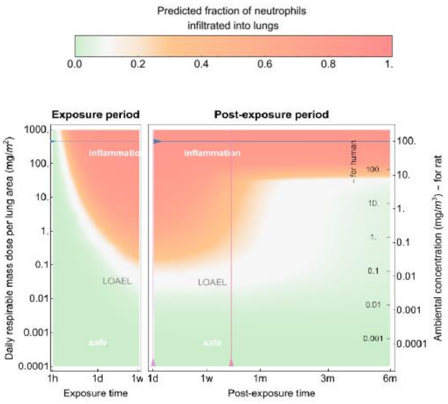

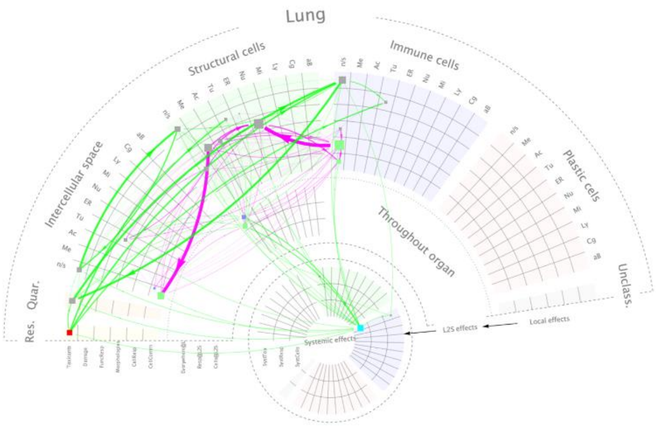

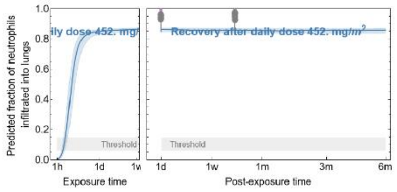

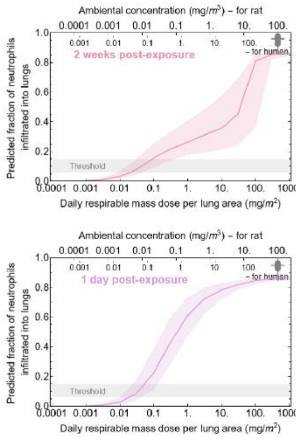

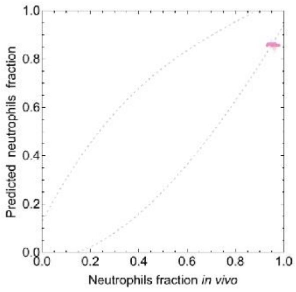

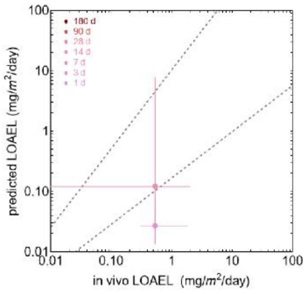

**Industrial 7**

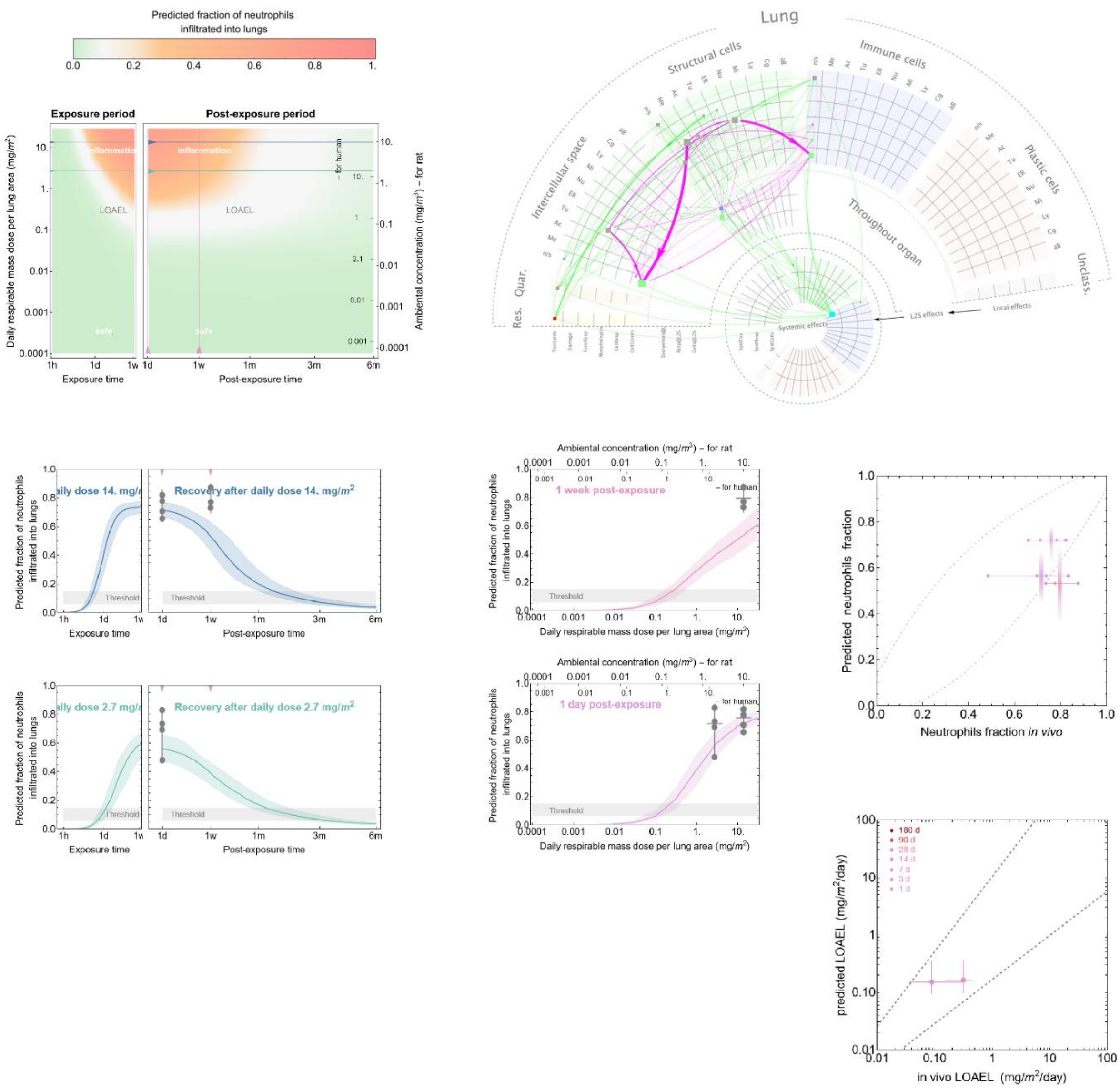

#### Industrial 8

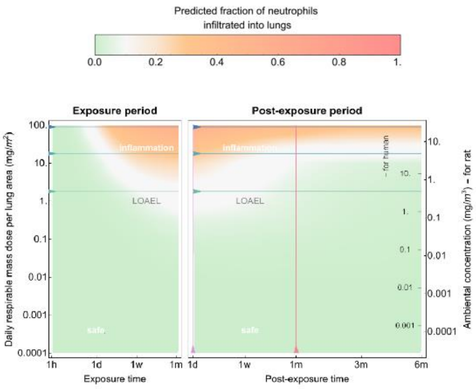

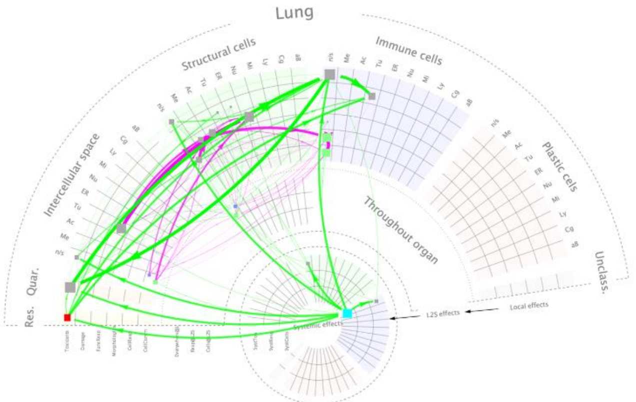

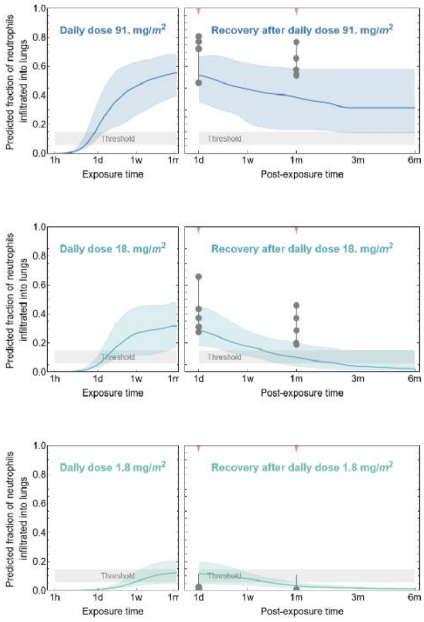

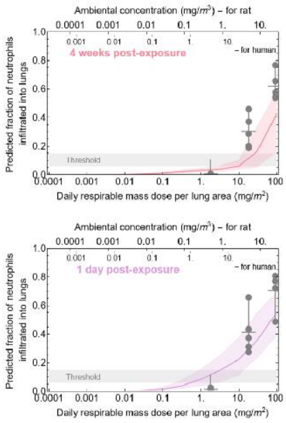

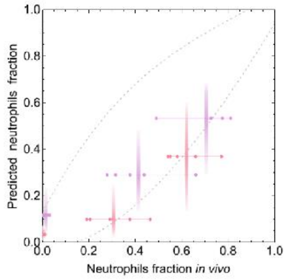

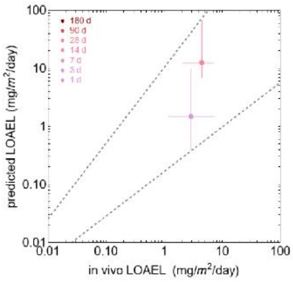

#### Industrial 9

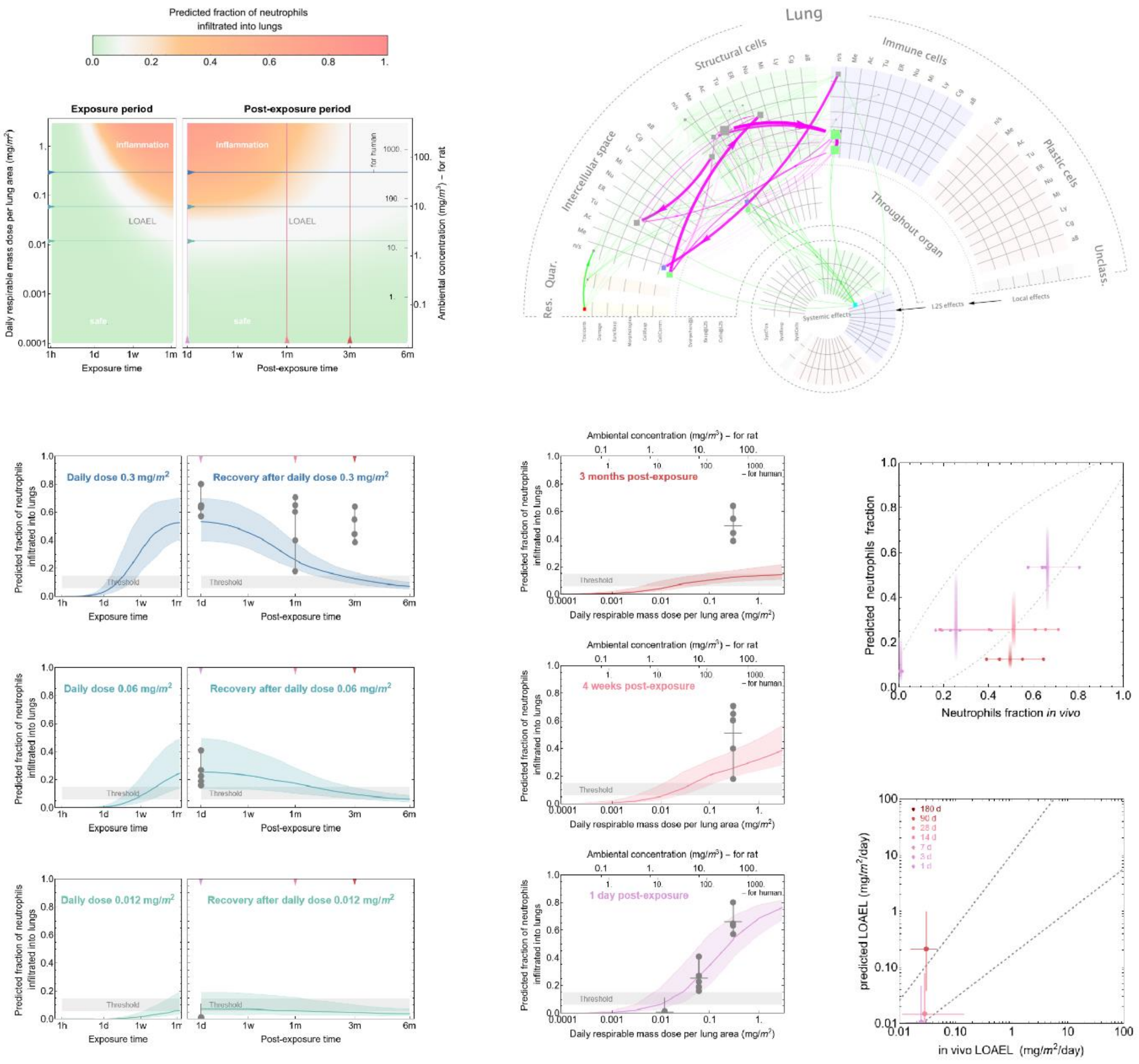

#### SI-4 Material characterization table

*Table 1. List of 44 benchmark nanomaterials used for the calibration and validation of the In Finite in vitro-learned digital twin (InFiniteLungDT). *In vivo values where data for 3 months were not available were substituted with data for 1 month after exposure. JRC in "other names" column – denotes reference nanomaterials provided by the JRC Nanomaterials Repository, which hosts a list of nanomaterials with their main physico-chemical characteristics, including particle size and surface area. More detailed characteristics of nanomaterials and in vivo data are provided in references cited next to nanomaterial name.*

| Nanomaterial name | Chemistry and morphology | Material family | set | other names | Source of NMs | NM size DxL [nm] | BET [$m^2/g$] | max (Ranked Dissolution core material in GS, PSF) | Ranked release (potential) of toxic metals in PSF | NM material density (not bulk) [$g/cm^3$] |
|---|---|---|---|---|---|---|---|---|---|---|
| **Diesel N1650**[6] | diesel exhaust particles from a heavy-duty diesel truck | combustion | validation | Diesel NIST 1650, DEP SRM 1650b | National Institute of Standards and Technology (NIST) | 24 | 108 | | | 0.8 |
| **Diesel N3**[7] | diesel exhaust particles | combustion | validation | DIESEL 9,8% O2, DEP9.7 | NRCWE | 22 | 152 | | | |
| **DQ12**[8] | $SiO_2$ crystalline | | calibration | crystalline silica | NRCWE | 225 | 10 | 2 | 0 | 2.6 |
| **Exhaust N1**[7] | rapeseed methyl ester exhaust | combustion | validation | RME 13% O2, RME 13 | NRCWE | 15 | 222 | | | |
| **Exhaust N5**[7] | hydrotreated vegetable oil exhaust | combustion | validation | HVO 13% O2, HVO13 | NRCWE | 21 | 160 | | | |

| | | | | | | | | | | |
|---|---|---|---|---|---|---|---|---|---|---|
| **Flamruss 101**[9] | carbon black, spherical | combustion | validation | Carbon black, FL101, aka lamp black 101 | NRCWE - The Grolman Group, Denmark | 95 | 24 | | | 1.7 |
| **GO N058**[10,11] | graphene sheet, highly hydroxylated | carbonaceous | validation | graphene oxide; GO-4-250 Graphenea, "GO" | NRCWE - Graphenea, San Sebastian, Spain | 1.7 x 2500 | 700 | 0 | 0 | 2.3 |
| **GO N059**[10,11] | reduced Graphene Oxide sheet | carbonaceous | calibration | Graphene oxide reduced; "rGO" | NRCWE - Graphenea, San Sebastian, Spain | 1.7 x 1500 | 411 | 0 | 0 | 2.3 |
| **HNTNN**[12] | elongated halloysite; phyllosilicate | clays | calibration | Halloysite Nanotubes, Natural Nano, HNT200 | NRCWE - Natural Nano, NY, USA | 61 x 250 | 25.6 | 2 | 0 | 2.3 |
| **HNTNNE**[12] | elongated halloysite - etched; phyllosilicate | clays | validation | Halloysite Nanotubes, Etched | NRCWE - Natural Nano, NY, USA | 27 x 100 | 128 | 1 | 0 | 2.3 |
| **MWCNT 401** [13–20] | Multi-walled carbon nanotubes - thick and long | carbonaceous | validation | NM-401, "CNTLarge" | JRC, Producer IO-LE-TECNanomaterials (CP-0006-SG) | 67 x 4000 | 18 | 0 | 2 | 2.1 |
| **MWCNT N006** [14,19,21–24] | Multi-walled carbon nanotubes | carbonaceous | calibration | Mitsui-7, lot #061220-24, MWNT-7, | Mitsui, Japan | 74 x 5700 | 26 | 0 | 3 | 2.1 |

| | | | | | | | | | | |
|---|---|---|---|---|---|---|---|---|---|---|
| | | | | MWCNT XNRI-7 | | | | | | |
| **MWCNT N040** [14,15,17,25] | Multi-walled carbon nanotubes | carbonaceous | validation | Cheaptubes kode: sku-30102 | NRCWE - Cheaptubes, VT, USA | 22 x 500 | 150 | 0 | 3 | |
| **MWCNT N041** [14,15,17,25] | Multi-walled carbon nanotubes with OH | carbonaceous | validation | Cheaptubes kode: sku-30202 | NRCWE - Cheaptubes, VT, USA | 27 x 1000 | 152 | 0 | 3 | 2.1 |
| **MWCNT N042** [14,15,17,25] | Multi-walled carbon nanotubes with COOH | carbonaceous | validation | Cheaptubes kode: sku-30302 | NRCWE - Cheaptubes, VT, USA | 30 x 700 | 141 | 0 | 3 | |
| **MWCNT N043** [14,15,17,25] | Multi-walled carbon nanotubes | carbonaceous | validation | Cheaptubes kode: sku-30107 | NRCWE - Cheaptubes, VT, USA | 27 x 800 | 82 | 0 | 4 | |
| **MWCNT N044** [14,15,17,25] | Multi-walled carbon nanotubes with OH | carbonaceous | validation | Cheaptubes kode: sku-30207 | NRCWE - Cheaptubes, VT, USA | 33 x 1300 | 74 | 0 | 4 | |
| **MWCNT N045** [14,15,17,25] | Multiwalled carbon nanotubes with COOH | carbonaceous | validation | Cheaptubes kode: sku-30137 | NRCWE - Cheaptubes, VT, USA | 30 x 1550 | 119 | 0 | 4 | |
| **MWCNT N046** [14,15,17,25] | Multi-walled carbon nanotubes | carbonaceous | validation | Cheaptubes kode: sku-30111 | NRCWE - Cheaptubes, VT, USA | 29 x 700 | 223 | | | |
| **MWCNT N047** [14,15,17,25] | Multi-walled carbon nanotubes with OH | carbonaceous | validation | Cheaptubes kode: sku-30112 | NRCWE - Cheaptubes, VT, USA | 23 x 500 | 216 | | | |

| | | | | | | | | | | |
|---|---|---|---|---|---|---|---|---|---|---|
| **MWCNT N048** [14,15,17,25] | Multi-walled carbon nanotubes with COOH | carbonaceous | validation | Cheaptubes kode: sku-30113 | NRCWE - Cheaptubes, VT, USA | 18 x 1600 | 185 | | | |
| **MWCNT N061** [15,26] | Multi-walled carbon nanotubes with NH2 | carbonaceous | validation | MWCNT (TNMN2); TimesNano | NRCWE - Timesnano, China | 16 x 700 | 170 | | | |
| **MWCNT N062** [15,26] | Multi-walled carbon nanotubes, pristine | carbonaceous | validation | MWCNT (TNM1) | NRCWE - Timesnano, China | 9 x 450 | 443 | | | |
| **MWCNT N063** [15,26] | Multi-walled carbon nanotubes with more OH | carbonaceous | validation | MWCNT (TNMH1) | NRCWE - Timesnano, China | 14 x 350 | 426 | | | |
| **MWCNT N064** [15,26] | Multi-walled carbon nanotubes with more COOH | carbonaceous | validation | MWCNT (TNMC1) | NRCWE - Timesnano, China | 7 x 2000 | 445 | | | |
| **Nanoclay 3000** [27] | sheets, organoclay; phyllosilicate; functionalized with dialkyldimethyl ammonium | clays | calibration | Nanofil SE3000; Nanoclay 3000 is IJS internal name | NRCWE - Sud-Chemie AG, Germany | 300 | 2.7 | 3 | 0 | 2.3 |
| **Nanoclay 9** [27] | sheets, Montmorillonite phase; phyllosilicate; functionalized with benzalkonium chloride | clays | validation | Nanofil 9; Nanoclay 9 is IJS internal name; Cloisite 10 A | NRCWE - Sud-Chemie AG, Germany | 300 | 24 | 2 | 0 | 2.3 |

| | | | | | | | | | | |
|---|---|---|---|---|---|---|---|---|---|---|
| **Nanoclay Bent** [27] | sheets, bentonite, organoclay; phyllosilicate; small ‘blade’-like flakes | clays | validation | same as JRC NM-600 "Bentonite" | NRCWE - anonymous company | 650 | 74 | 1 | 0 | 2.3 |
| **$NiFe_2O_4$ N022** [28] | spherical isometric magnetite | alloys | validation | stock nr 4110FY | NRCWE - NanoAmor, Houston, USA | 25 | 86.9 | 2 | 4 | 5.4 |
| **$NiZnFe_4O_8$ N020** [28] | spherical isometric magnetite | alloys | validation | stock nr 4115FY | NRCWE - NanoAmor, Houston, USA | 20 | 104 | 2 | 4 | 5.4 |
| **Printex 90** [17,29,30] | carbon black, spherical | combustion | validation | Carbon black | NRCWE - Evonik/Degussa AG | 14 | 182 | 0 | 0 | 1.7 |
| **Printex XE2B** [9] | carbon black, spherical | combustion | calibration | Carbon black | NRCWE - Grolman Group, Denmark | 30 | 1142 | 0 | 0 | 1.7 |
| **SWCNT N054** [15,26] | single-walled carbon nanotubes with COOH | carbonaceous | validation | SWCNT (TNSC) | NRCWE - TimesNano, China | 17 x 17500 | 371 | 0 | 4 | 1.3 |
| **SWCNT N055** [15,26] | single-walled carbon nanotubes | carbonaceous | calibration | SWCNT (TNSS) | NRCWE - TimesNano, China | 14 x 2000 | 453 | 0 | 4 | 1.3 |
| **SWCNT N056** [15,26] | single-walled carbon nanotubes with OH | carbonaceous | calibration | SWCNT (TNSSH) | NRCWE - TimesNano, China | 13 x 2000 | 373 | 0 | 4 | 1.3 |

| | | | | | | | | | | |
|---|---|---|---|---|---|---|---|---|---|---|
| **$TiO_2$ large** [8] | TiO2 anatase/rutile nanospheres | metal-oxide | calibration | NM-2_TiO2-A100, MKN-TiO2-A100, MKN-A100, TiO2 NM-2 | NRCWE - MK Nano, ON, Canada | 16-28 | 74 | 0 | 0 | 4.0 |
| **$TiO_2$ N001** [14,31,32] | spherical rutile | metal-oxide | calibration | TiO2NP10 | NRCWE - NanoAmor, TX, USA | 10 | 99 | 1 | 0 | 4.2 |
| **$TiO_2$ N025** [32,33] | spherical rutile | metal-oxide | validation | Lot 2010-9; TiO2 NP38; "TiO2 38nm" | NRCWE - NaBond, Shenzhen, China | 38 | 28 | 0 | 0 | 4.2 |
| **$TiO_2$ N030** [32–34] | rutile | metal-oxide | calibration | "TiO2 10.5nm" | NRCWE - NanoAmor, TX, USA | 12 | 139 | | | 4.2 |
| **$TiO_2$ nanocubes**[8] | TiO2 anatase nanocubes | metal-oxide | calibration | TiO2-NCs | Polona Umek, IJS | 15 x 20 | 97 | | | 3.8 |
| **$TiO_2$ nanotubes**[8] | TiO2 anatase nanotubes | metal-oxide | validation | TiO2-NTs | Polona Umek, IJS | 10 x 100 | 154 | | | 3.8 |
| **$ZnFe_2O_4$ N021**[28] | spherical isometric magnetite | alloys | calibration | stock nr 5710FY | NRCWE - NanoAmor, TX, USA | 22 | 87.7 | 2 | 4 | 5.4 |
| **ZnO 110** [35,36] | ZnO uncoated; spherical zincite | metal-oxide | calibration | NM-110, BASF Z-Cote | JRC | 70-110 | 12 | 5 | 5 | 5.6 |

| | | | | | | | | | | |
|---|---|---|---|---|---|---|---|---|---|---|
| **ZnO 111**[35,36] | ZnO coated with hydrophobic triethoxycapryl silane | metal-oxide | calibration | NM-111, BASF Z-Cote HP1 | JRC | 60-90 | 15 | sol ubl e | sol ubl e | 5.6 |

#### SI-5 Masks for image quantification

This image shows typical input fluorescence images and their automatically determined masks that are used in image quantification.

The cell-type segmentation into epithelial cells and macrophages is provided by cell-type-specific tracking probes, leading to reliable cell-type classification.

Visually, the masks of functional units and nanomaterial match the input fluorescence images to a high degree. The most common artefacts in these masks are:
- in very thin parts of the cell membrane, the signals are sometimes too low for detection (magenta arrow in frame b), which reduces the cell membrane mask for a few percents;
- for weakly scattering nanoparticles, some of their signal in cells (white arrows in frame d) can be overpowered by the scattering of cells themselves and is consequently filtered out from the masks, representing a negligible error in determination of nanomaterial amount.

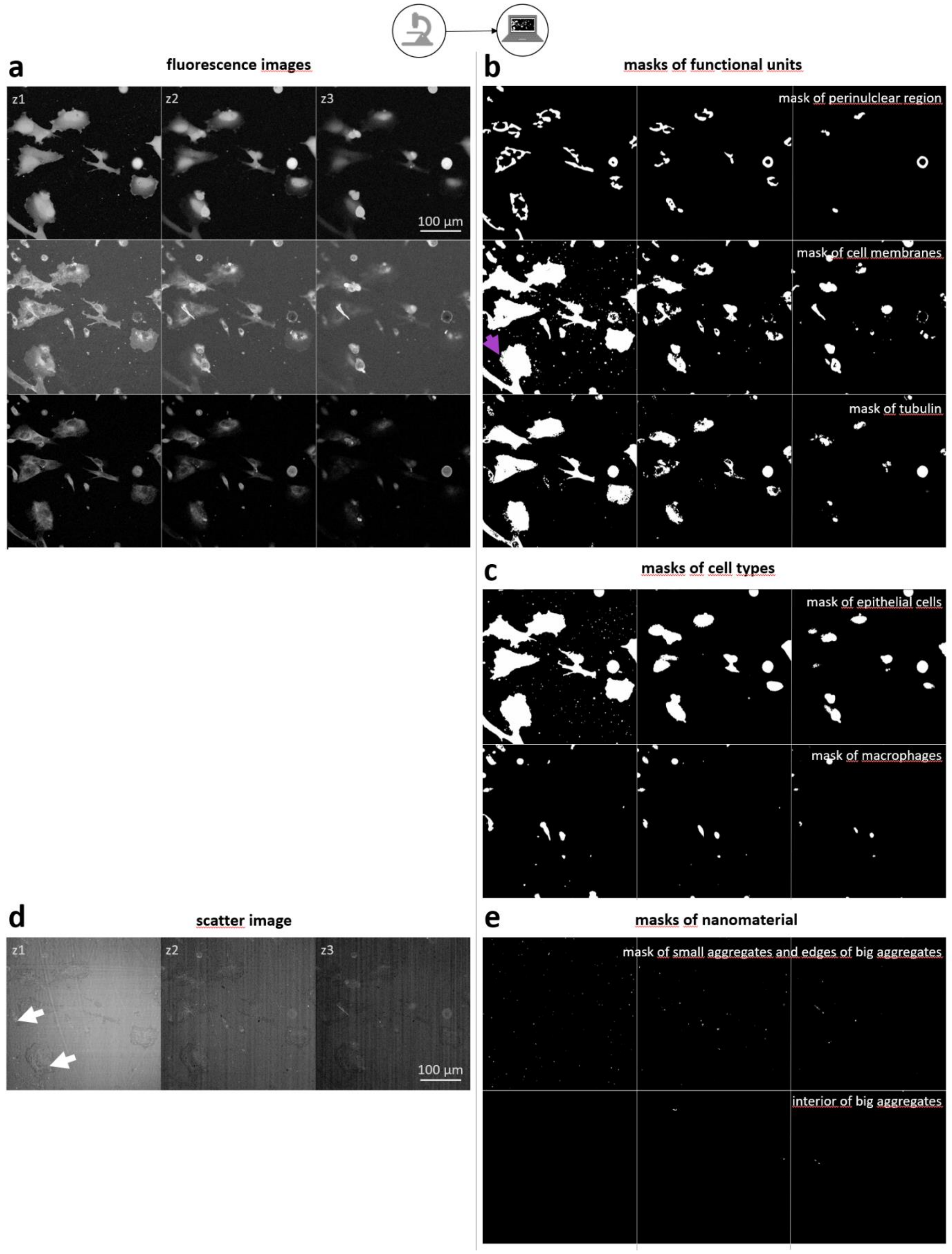

*FigureS 4. Imaging quality assurance.* ***a*** *fluorescence microscopy images recorded at three different focal planes (z1, z2, and z3) to assure correct assignement of material internalization and to avoid artefact due to focus drift, which was minimized by autofocus algorithm that kept constant focal planes,* ***b*** *examples of masks of biological observables,* ***c*** *masks of two different cell types assuring cell segmentation accuracy and assignement of materials to particular cell type,* ***d*** *scatter image (additional modality) for detection of the location and amount of material interacting with cells,* ***e*** *masks of detected materials at different z positions. Field of view FOV = 375 µm.*

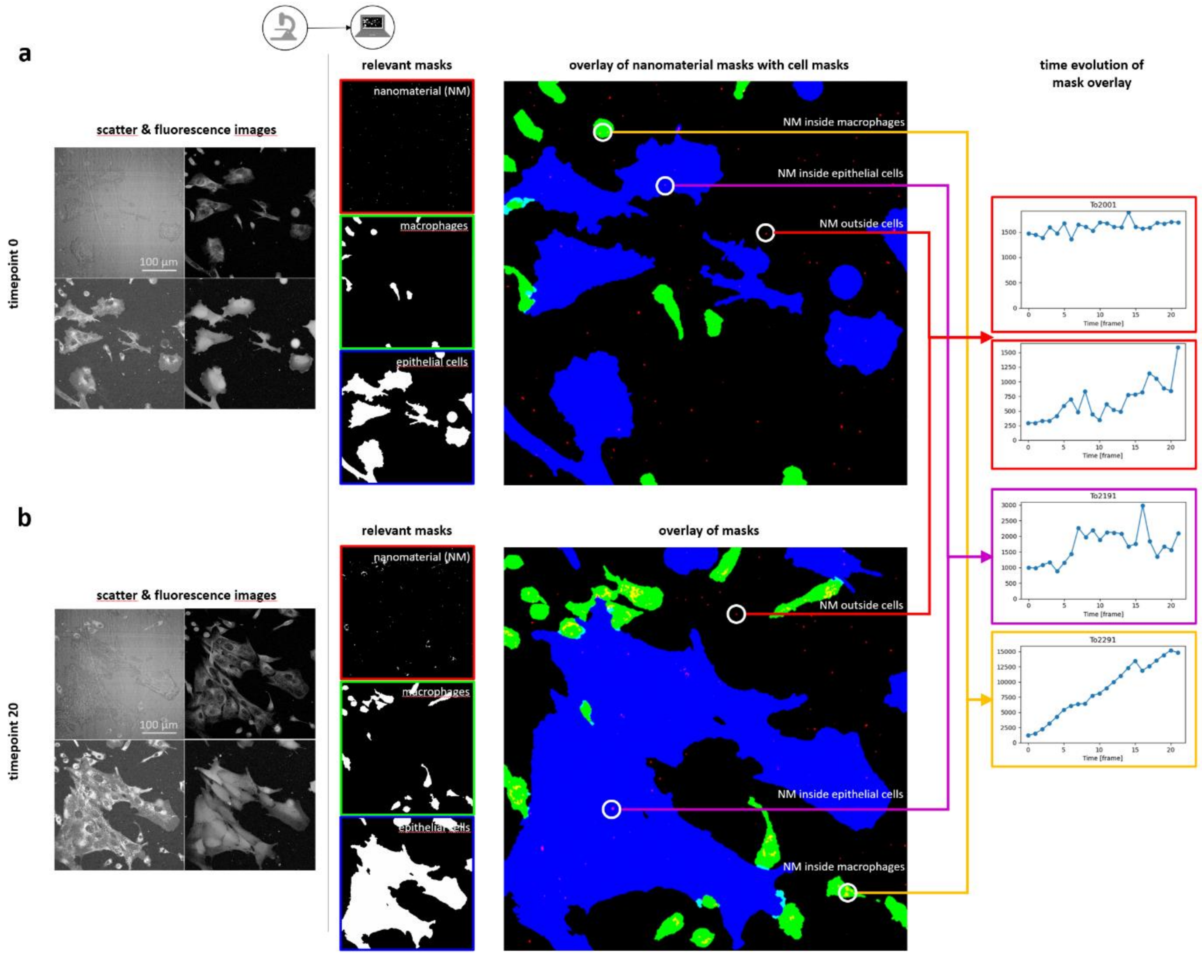

*FigureS 5. Imaging quality assurance – cell type segmentation, discrimination between internalized versus material outside cells.* ***a*** *confocal fluorescence and scattering microscopy images at time point 0,* ***b*** *at time point 20. Identified nanomaterial (red), identified macrophages (green), identified epithelial cells (blue). Field of view FOV = 375 µm. Right most column shows quantified time evolution of four different observables: To2001 – nanomaterial outside cells, To2191 – nanomaterial inside of epithelial cellsTo2291 – nanomaterial (toxicant) inside of immune cells.*

This image shows an example of determining nanomaterial uptake into different cell types. We use a 2D in vitro model and nanomaterial concentrations high enough to ensure that cells from each cell type are exposed to nanomaterial. The actual number of cells with internalised material depends on the cell type and nanomaterial type. In this case, internalised nanomaterial can be seen in most macrophages and more than half of epithelial cells at timepoint 20.

Material uptake analysis (observables *To2x91*) is performed on masks from unbinned images, with pixel size of 0.5 µm x 0.5 µm x 3 µm. The voxel from which the signal originates is 0.4 µm x 0.4 µm x 1.5 µm (i.e. PSF size for confocal imaging with pinhole 1.1 AU and a 0.8 NA 20x air objective). The total uptake into a cell type from the total area of 350 µm x 750 µm x 6 µm is binned together into one observable.

### SI-6 Materials and Methods

#### Animals

Animals were handled in a consistent manner for exposure to all 49 benchmark nanomaterials measured previously (as indicated by references next to material names in Table 1 above) and 7 additional industrial materials that were collected and measured during this study within the nanoPASS project. Female C57BL/6 mice aged 5-6 weeks were obtained from Taconic (Ry, Denmark) or Janvier (Janvier Labs, Le Genest-Saint-Isle, France) as described in more detail in Hadrup et al. (2025) [37]. Mice were acclimatized for 1-3 weeks before the experiment and were 8 weeks of age at the start of the study. All mice were given food (Altromin no. 1324) and water ad libitum. The mice were group housed in polypropylene cages with sawdust bedding and enrichment at controlled temperature and humidity and with a 12-h light:12-h dark cycle. The experiments were approved by the Danish Animal Experiments Inspectorate under the permit licence numbers stated in the individual papers referenced in SI-4, Table 1, and were carried out following the guidelines for the ethical conduct and care of animals used in research. All procedures complied with EC Directive 86/609/EEC and Danish laws regulating experiments on animals.

#### Preparation of stocks of nanomaterials for in vitro and in vivo exposure

Materials from the same stock (dry powder) were prepared either for *in vivo* or *in vitro* exposure. In most of the *in vivo* studies, water with 2% mouse serum (sibling serum) was used as a vehicle, but other validated vehicles were also used [38]. Vehicle-exposed negative controls and mice exposed to the benchmark particle carbon black Printex90 [39], were included in parallel. The stock particle suspensions were sonicated using a 400 W Branson Sonifier S-450D (Branson Ultrasonics Corp., Danbury, CT, USA) equipped with a disruptor horn (Model number: 101147-037). Samples were continuously cooled on ice during the sonication procedure. The stock suspension was used for the high dose, diluted 1:3 for the medium dose (0.054 mg) and diluted further 1:3 for the low dose (0.018 mg). Vehicle control solutions were prepared in parallel as described in more detail in Jackson et al. (2015) [40].

#### Characterization of nanomaterials

The hydrodynamic particle size distributions in the exposure media were determined by Dynamic Light Scattering (DLS) using a Malvern Zetasizer Nano ZS (Malvern Instruments Ltd, UK). Data were analyzed using the Dispersion Technology Software (DTS) version 5.0 (Malvern Instruments Ltd). Samples were measured at 25°C in 1 mL Malvern disposable polystyrene cuvettes, as described in more detail in Bourdon et al (2012) [30].

#### In vivo exposure of mice

The animal studies were conducted over a period of approximately 15 years, primarily using the same experimental setup as described [41]. Mice were exposed via a single intratracheal instillation under a brief isoflurane sedation [37,42]. During instillation the mice were placed on their backs on a 40 degree slope. The trachea was intubated using a 24 gauge BD Insyte catheter (Becton Dickinson, Denmark) with a shortened needle. The correct location of each intubation was tested using a highly sensitive pressure transducer (pneutachymeter). A 50 µl suspension was instilled followed by 150 µl air with a 250 µl SGE glass syringe (250F-LT-GT, MicroLab, Aarhus, Denmark). Control animals received vehicle instillations. The intubation catheter was held head up until proper breathing was assured. The animals did not show signs of respiratory distress, lethargy or other physical symptoms of exposure.

#### Collection of BAL cells and differential counting

Mice were euthanised by intraperitoneal injection of Zoletil Forte (250 mg/ml), Rompun (20 mg/ml), and Fentanyl (50 µg/ml) in sterile saline (0.1 ml/10 g body weight) in combination with heart blood withdrawal. To obtain the bronchoalveolar (BAL) fluid the lungs were flushed twice with sterile 0.9% sodium chloride through the trachea. The BAL fluid was kept on ice until separation of fluid and cells by centrifugation at 400×g for 10 min at 4 °C [43].

The separated BAL cells were resuspended in 100 µl HAMF12 medium containing 10% fetal bovine serum. 40 µl of the cell suspension was mixed with 160 µl medium containing 10% dimethyl sulfoxide (DMSO). The total number of cells and of dead cells was determined from 20 µl diluted cell suspension by NucleoCounter NC-200TM (Chemometec, Allerød, Denmark). 40 µl of the cell suspension was collected on microscope slides by centrifugation at 60×g for 4 min and where after the cells were fixed with 96% ethanol and stained with May–Grünwald–Giemsa. The samples were randomized and blinded before scoring of macrophages, neutrophils, lymphocytes, eosinophils, and epithelial cells, which were assessed by counting 200 cells/sample under light microscope (100× magnification).

#### In vitro exposure of InFiniteLungDT biological platform

The in vitro biological platform consists of a coculture of lung epithelial cells and lung immune cells, as well as both monocultures.

The MH-S cell line consists of mouse derived alveolar macrophages [44]. These contact-uninhibited, adherent cells closely mimic primary alveolar macrophages by retaining typical morphology, phagocytic capacity, esterase-positive/peroxidase-negative enzymatic profiles, and expression of Mac-1 and I-A surface antigens. They also exhibit Fc receptor-dependent phagocytosis, increased IL-1 production upon LPS stimulation, and polarization after exposure to stimuli, making them an excellent choice for studying lung pathology and immunological functions [45].

The LA-4 cell line consists of murine alveolar type II epithelial cells [46]. These cells replicate key features of type II pneumocytes, including contact-inhibited growth, a polygonal morphology, lamellar bodies, and high phospholipid synthesis [47]. They serve as an ideal model system for reproducing in vivo epithelial cell responses to metal-oxide nanoparticles [5].

#### SI-7 Materials can not be clustered by material family and specific surface area

**a** **each material family shown in different colour**

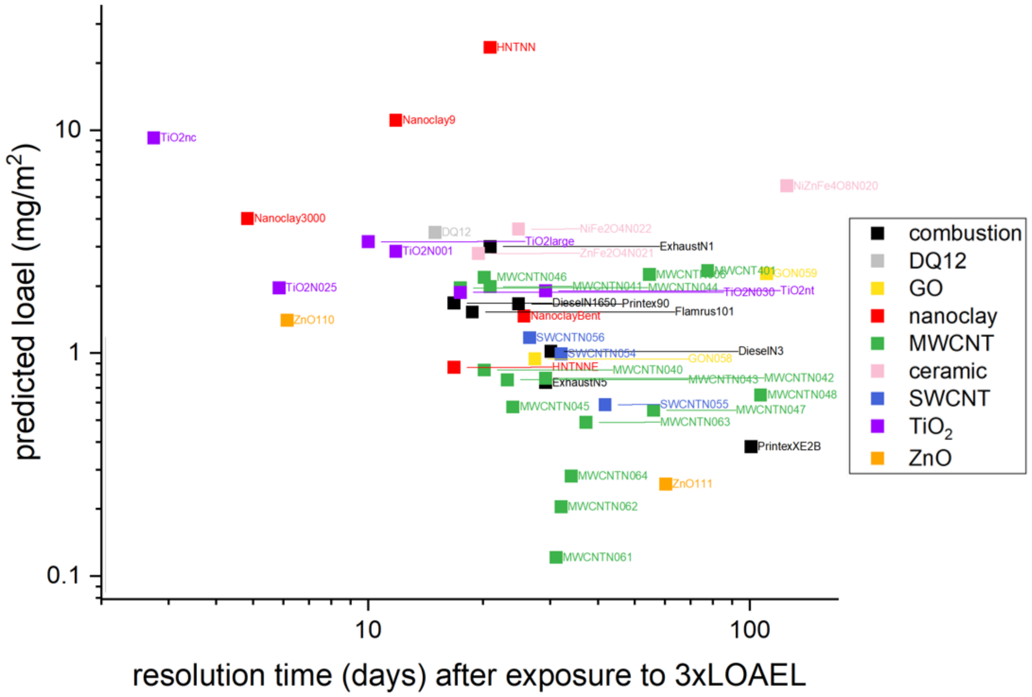

**b** **each material specific surface area (BET) shown in different colour**

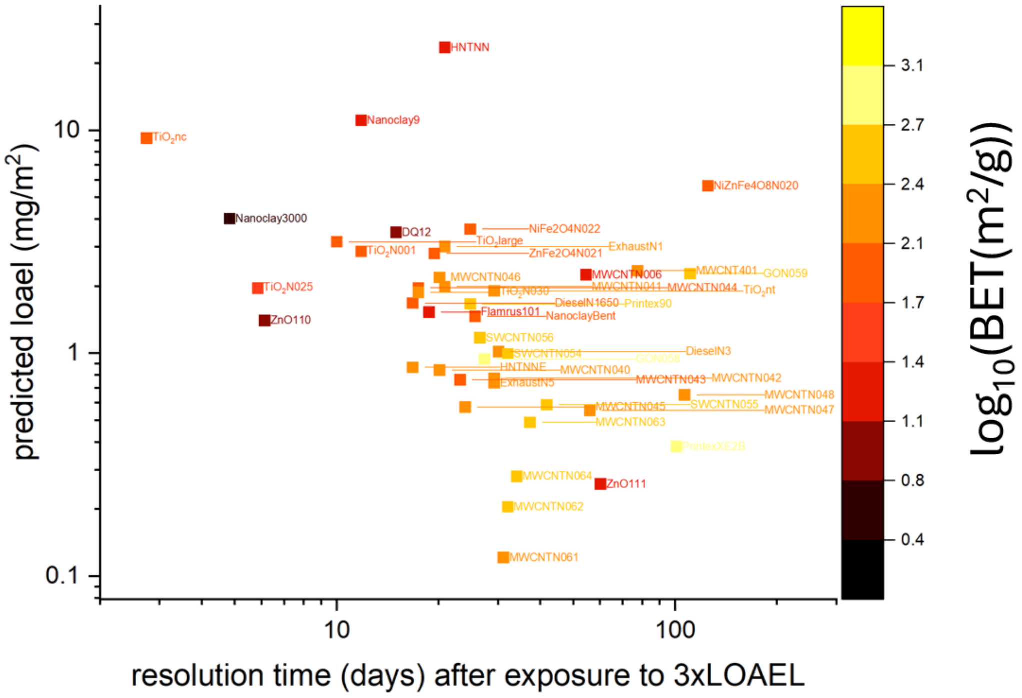

*FigureS 6. Absence of clustering of neutrophilic lung inflammation LOAEL and resolution times by:* ***a*** *material family – chemical/structural similarity (carbonaceous, metal oxides, $SiO_2$, $TiO_2$, MWCNTs, SWCNTs),* ***b*** *material specific surface measured by BET method.*

#### SI-8 ZnO100 and ZnO111 as example of solubility limit applicability domain case

We included two materials with high solubility (partial solubility):

1) ZnO110 with experimentally measured solubility 60% in $H_2O$, and 47% in C3A cell culture medium following 24 h of incubation at 37°C, 5% $CO_2$, and
2) ZnO111 with solubility 18% in $H_2O$; 39% in C3A cell culture medium over 24 hours [48].

As concluded by Kermanizadeh et al., “results suggest that about 50–60% of the added ZnO NMs were dissolved in the cell medium after 24 h. ” [48].

#### SI-9 Baseline phenotype of unexposed LA-4 and MH-S – integrity of inflammatory pathways

MH-S retain many of the properties of alveolar macrophages including phagocytosis, typical macrophage morphology and polarisation when exposed to stimuli. The MH-S cell line of murine alveolar macrophages (obtained from ATCC, CRL-2019 [44]) was originally obtained from a bronchoalveolar lavage of a 7-week male mouse by culturing the adherent alveolar macrophages for several days before infecting them with the SV40 virus [45]. SV40 viral transformation of alveolar macrophages collected from male BALB/c mice resulted in the establishment of the MH-S cell line [49]. This cell line maintains several characteristic traits of alveolar macrophages, such as (1) typical macrophage morphology with adhesive properties; (2) phagocytic capacity; (3) enzymatic profiles that are esterase-positive and peroxidase-negative; (4) positive expression of surface membrane antigens Mac-1 and I-A; and (5) Fc receptor-dependent phagocytosis of sheep red blood cells (SRBCs) coated with immunoglobulin G. Additionally, LPS stimulation markedly increased constitutive IL-1 production [50]. The contact-uninhibited, adherent MH-S share many properties with alveolar macrophages, including typical macrophage morphology, function and antigenic characteristics [45]. According to the creators of the cell line, "this cell line should facilitate studies where homogeneous populations of alveolar macrophages are desirable, especially those involved in determining the immunological functions of alveolar macrophages and their potential role in lung pathology" [45], and is as such an excellent choice for the planned experiments.

The LA-4 cell line of murine alveolar type II epithelial cells (obtained from ATCC, CCL-196 [46]) was originally derived from a benign urethane-induced mouse lung adenoma (tumor) of a 28-week-old mouse [47]. Similar to type II alveolar pneumocytes, these cells contain lamellar bodies (organelles in which surfactant is stored), have a characteristic polygonal-shaped morphology, and their growth is contact-inhibited (although they do gradually lose this inhibition when cultures reach confluency) [47]. They can also synthesize a large amount of phospholipids, albeit with a different phospholipid composition than pneumocytes type II in vivo [51]. In contrast to the normal diploid mouse cells with 40 chromosomes per cell [47], the LA-4 cells have on average 114 chromosomes (between 38 and 256 chromosomes) per cell [46]. In our case, they appeared to be the perfect model system because they could reproduce the in vivo observable events related to the response of epithelial cells to metal-oxide nanoparticles [5].

#### SI-10 Effective concentration phase space per ROI

The European Centre for Ecotoxicology and Toxicology of Chemicals (ECETOC) 'Nano Task Force' proposed the grouping of nanomaterials in the DF4nanoGrouping framework by their specific mode-of-action that results in a toxic effect is the crucial element for the grouping [52]. They concluded that no specific regulatory frameworks for nanomaterial grouping exist yet, and more data on biokinetics and in situ characteristics at toxicity sites are needed. It was emphasized that for meaningful risk assessment, grouping should integrate:

1) intrinsic characteristics,

2) life-cycle changes,

3) biokinetics (whether nanomaterial reaches the target site), and

4) observed biological effects, including cellular uptake, intracellular trafficking, and toxic responses [53].

Main drawback of existing in vitro studies on NMs is that the effective dose (the particle mass reaching the cultured cells) is not calculated nor measured and is rarely correlated with to doses in in vivo studies [54]. In contrast InDiniteLungDT measures 3) and 4) with the dose that reaches the cultured cells for each exposure separately.

The basic idea proposed by in 2007 by the US National Academy of Science [55] was that if a NAM is capable of predicting that some adverse outcome pathway is triggered at any dose, even a very at irrelevantly high one [56], the NAM could be helpful for regulatory use. However, this only identifies a trigger as potentially harmful, which is not particularly helpful for regulatory risk assessment, where determining the dose is crucial.

Effective concentration during in vitro experiment can vary over two orders of magnitude even though the nominal concentration is the same. In addition, effective concentration can change differently during each in vitro experiment. In some cases it can vary for less than a factor of 2 (blue circles in Figure below) or by a factor of 4 (blue stars in Figure below).

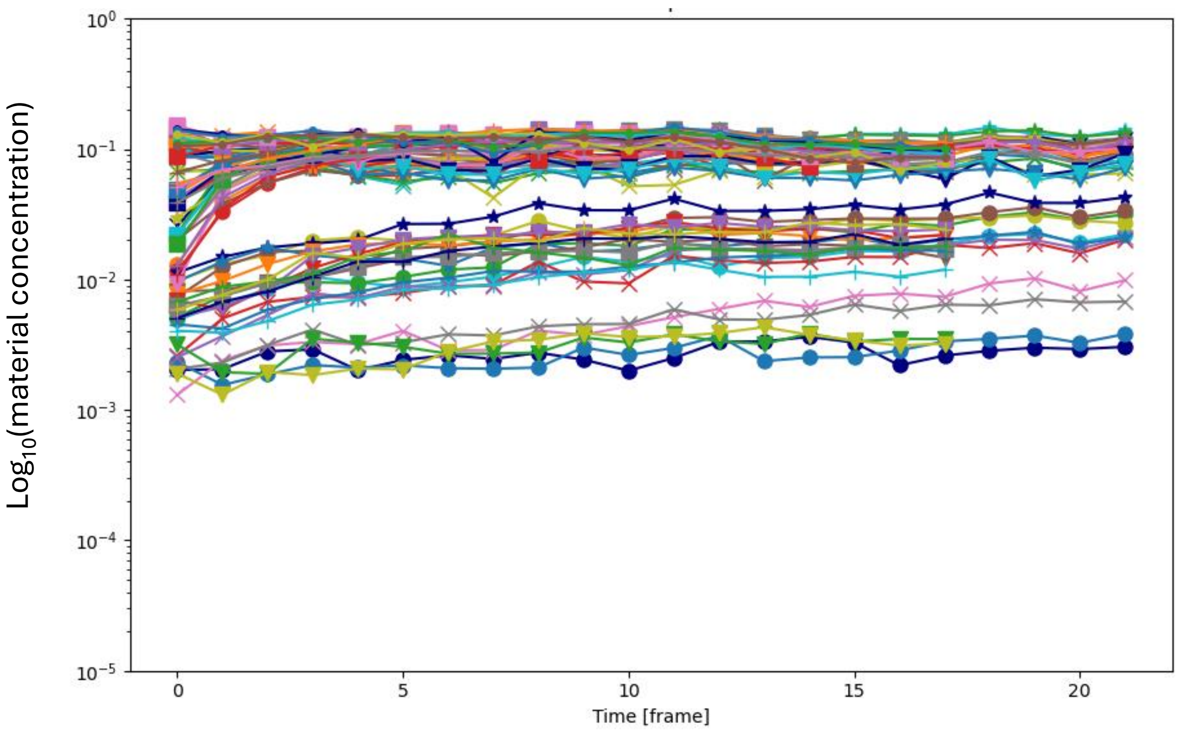

*FigureS 7. Amount of sedimentet MWCNTs that can interact with cells (effective dose) during in vitro measurement performed at the same nominal concentration. Each colour represents one region of interest during an in vitro experiment.*

#### SI-11 Predicted LOAELs/BMCs for individual post-exposure time windows

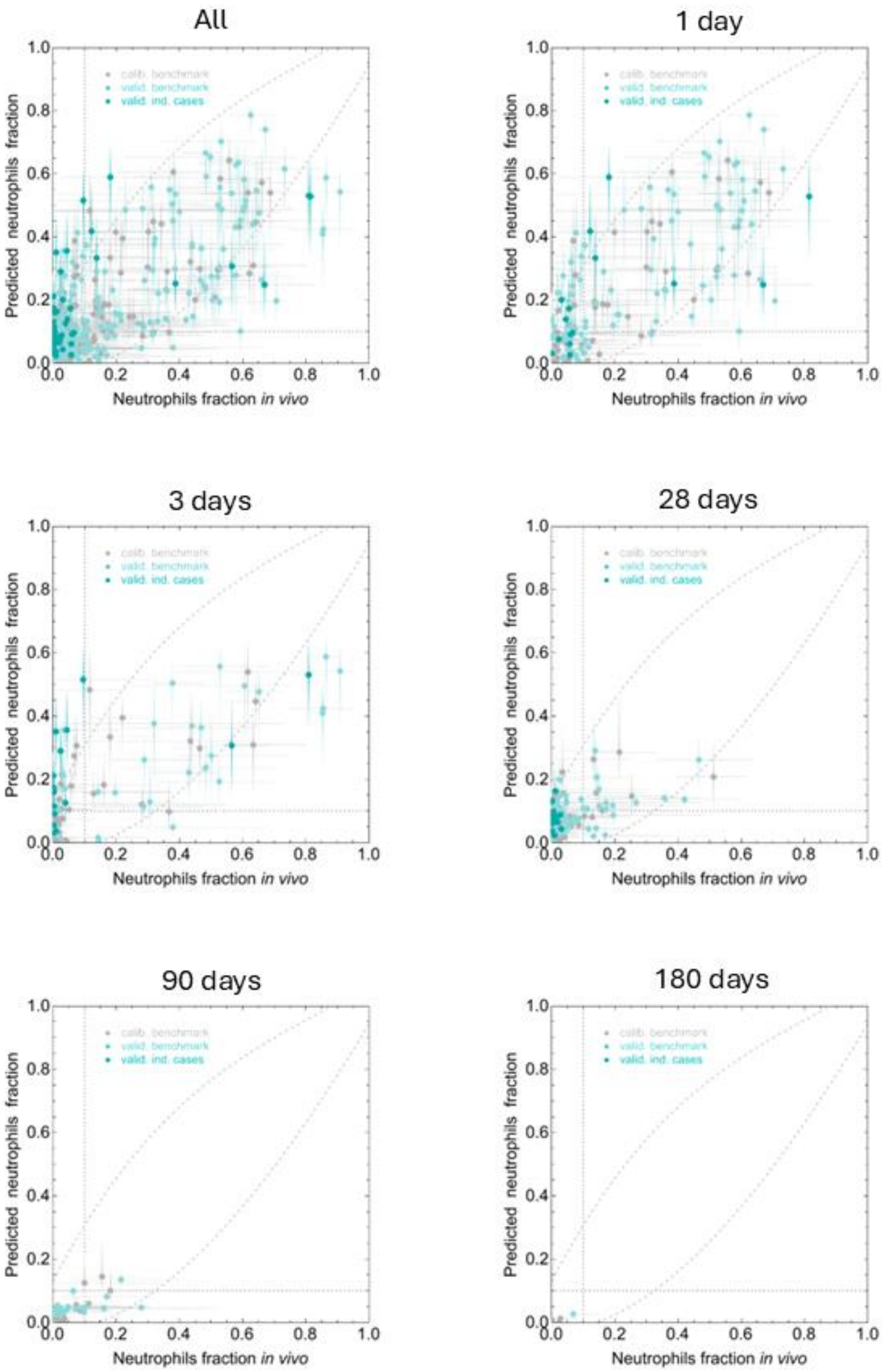

*FigureS 8. Predicted LOAELs/BMCs for individual post-exposure time windows, which show no notable prediction bias.*

#### SI-12 Lung burden clearance

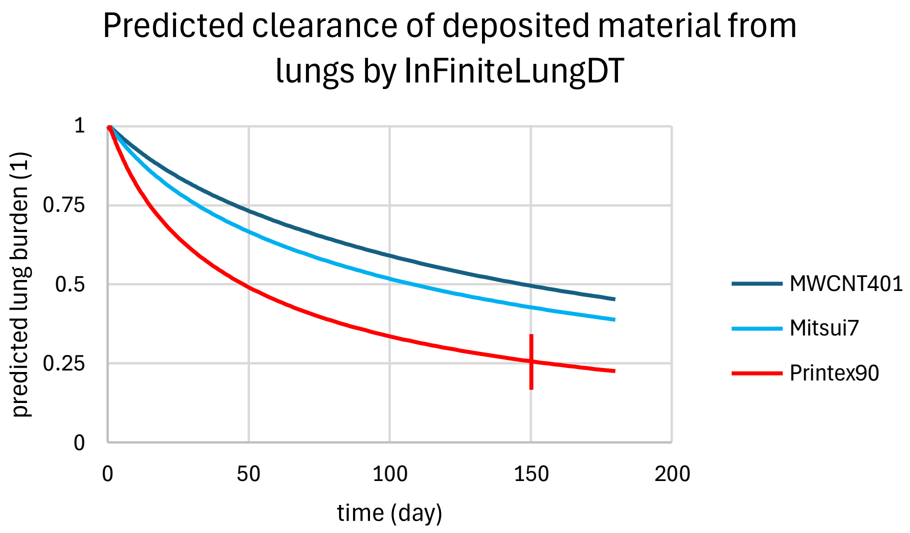

*FigureS 9. Lung burden clearance predicted by the digital twin based on in vitro measurements. The vertical line represent an uncertainty in the prediction.*

InfiniteLungDt predicts also lung burden at any post exposure time point, that can be used for a dose-range and post exposure time finding study.

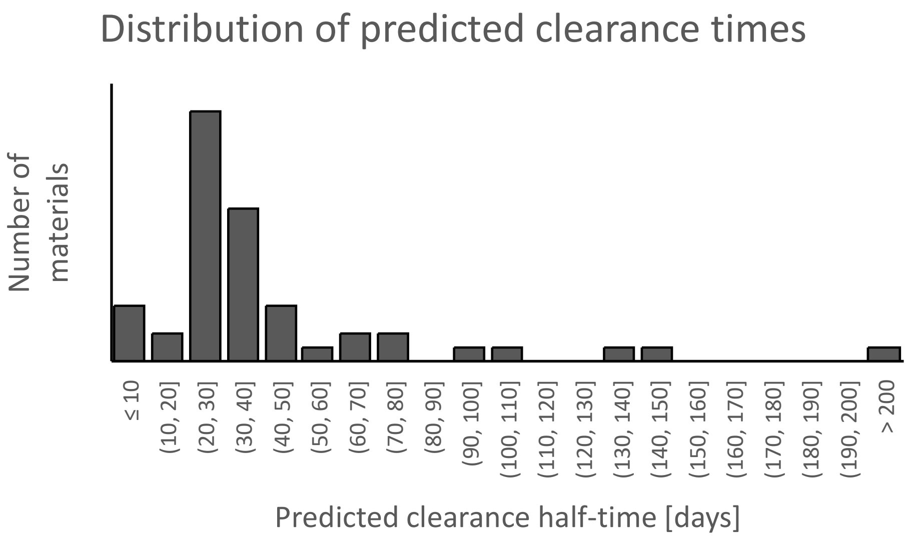

*FigureS 10. Distribution of predicted in vivo clearance times.*

The majority of clearance half-times, predicted by our model for the 49 nanomaterials included in this study, lie between 20 and 50 days, which is consistent with measured in vivo half-times from instillation studies performed with similar materials reviewed in Hadrup et al. [57,58]. However, the lack of in vivo data for these exact materials precludes a more quantitative assessment.

#### SI-13 Grouping of materials

##### Grouping methodology

The material-specific interaction couplings (section 3) are summed up within 12 classes of observables — which functional compartment drives which (see table below) — to reduce noise, and scaled to their own uncertainty determined from repeats of the reference material TiO2nt, so that every component of the representation is expressed in units of uncertainty (Figure 6a, $p_k$). The distance between any two materials (Figure 6b; $D_{ij}$) is then obtained as the sum of differences in all 12 dimensions (values are Gaussian-filtered to only distinguish close/far relationships, *G*), weighted by the 'fullness' of each material's MoA – i.e. same difference makes a larger distance for two materials with sparse MoA:

$$D_{ij} = \frac{\sum_k^{12} G(p_{ik} - p_{jk})}{\sqrt{\sum_k^{12} G(p_{ik}) \sum_k^{12} G(p_{jk})}}, \text{with } G(p) = 1 - \text{Exp}(-p^2/2)$$

| # | code | description |
|---|---|---|
| 1 | E+I ➔ E | epi response to epi-imu interaction |
| 2 | E ➔ E | epi-epi responses |
| 3 | E ➔ E+I | epi-imu interaction due to epi responses |
| 4 | T ➔ E | epi responses to toxicant |
| 5 | T ➔ I | imu responses to toxicant |
| 6 | T ➔ EC | extracellular debris due to toxicant |
| 7 | T ➔ E+I | epi-imu interaction due to toxicant |
| 8 | T ➔ T@EC | toxicant relocation into extracellular space and quarantining |
| 9 | T ➔ T@E | toxicant uptake and damage in epi |
| 10 | T ➔ T@I | toxicant uptake and damage in imu |
| 11 | E ➔ T | handling of toxicant and damage by epi |
| 12 | influx | predicted rate of neutrophils recruitment |

#### Case for 18 carbon nanotube materials grouped by early Mode-of-Action

The same grouping methodology as applied in the main manuscript (Figure 6) was applied to the subset of 18 single- and multi-walled carbon nanotube materials of the study. Panel a shows the similarity map, in which the distance between any two materials is their early Mode-of-Action (eMoA) distance, red links connect pairs below the grouping threshold (distance < 1), and node colour encodes the predicted acute LOAEL/BMC. Panel b shows the resulting grouping: 10 of the 18 materials fall into 5 groups of 2 members each, with the chosen reference material of each group labelled in red; the remaining 8 materials are not groupable at the threshold and would each require its own reference data. Carbon nanotubes are therefore not one mechanistic group: materials of nominally the same chemistry and morphology class separate by early Mode-of-Action, and the separation tracks the predicted LOAEL/BMC (node colour). This is the discrimination counterpart of the grouping demonstration in the main text, and it is the behaviour required of a method that is to support read-across without over-grouping.

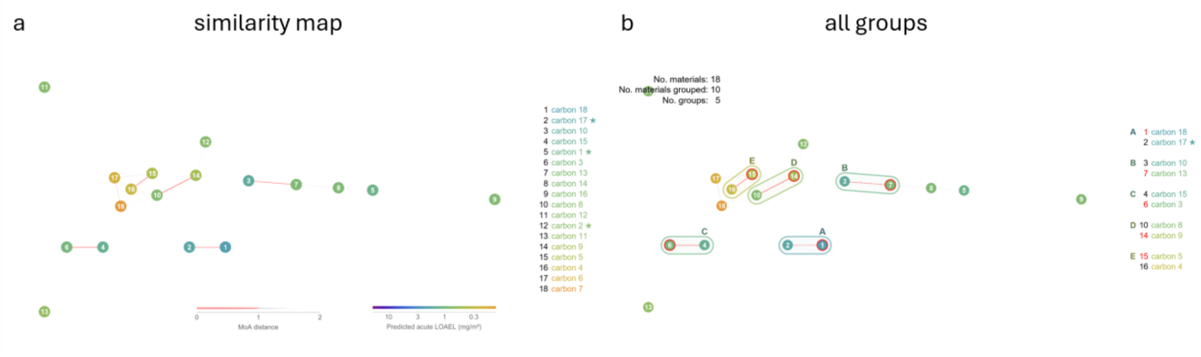

*FigureS 11. Grouping of 18 single- and multi-walled carbon nanotube materials by early Mode-of-Action. a similarity map (red links: eMoA distance below the grouping threshold of 1; grey links: above; node colour: predicted acute LOAEL/BMC). b grouping obtained at the threshold: 5 groups covering 10 of the 18 materials, chosen reference materials labelled in red.*

### SI-14 Rationale in InFiniteLung digital twin design

#### Why is predicting KE1497 important for chronic inflammation?

NEUTROPHIL INFLUX (KE1497 - Increased, recruitment of inflammatory cells) is known as a very **robust and consistent marker** of particle-induced inflammation in vivo [59].

There are many different proinflammatory cytokines that act as neutrophil chemo attractants and different particles initiates inflammation using different pathways that **all result in accumulation of neutrophils** in the lung [60,61]. Neutrophils not only mediate acute lung inflammation caused by acute infection but also play essential roles in other **chronic inflammatory diseases** by releasing proinflammatory mediators and activating other immune cells [62].

Measurement of neutrophil influx is mandated at different post-exposure times in OECD Test Guideline 413 (TG 413 sub-chronic inhalation toxicity, adopted 2018 to accommodate nanomaterial induced inhalation toxicity) and TG 412 (sub-acute inhalation toxicity). For this reason, **the largest number of in vivo measurements** exist in the literature for the lung neutrophil influx, which we could use to validate the accuracy (predictive capacity) with high statistical significance using one of the largest databases with more than 2600 animal instillation exposures at different doses and post-exposure time-points acquired in vivo in mice with matching 44 different nanomaterials for in vitro experiments.

#### How did we identify in vitro observables?

InFiniteLungDT monitors 10 key events, which are uniformly distributed within the network of human relevant inflammation related AOPs, listed in AOPWiki, this provides a solid ground for accurate prediction of chronic inflammation.

Altogether, InFiniteLungDT monitors 17 dynamic biological responses of cells (biological observables) which include direct material-induced damage of cellular organelles (functional units) or direct material co-localization with cellular organelles and cell types, and material-unspecific responses such as cell growth, material-independent responses of cellular organelles, cells (as a whole) and cell-cell communication via extracellular vesicle release and uptake.

To assure biological relevance of the biological platform of the InFiniteLungDT, we analyzed AOP Wiki, and identified 250 Key Events (KEs) associated with the keyword inflammation (blue dots and connections in Figure below), that are associated with 281 KEs directly upstream or downstream (gray dots and connections in the Figure below). Among them, the KE 1497 - neutrophils influx, is particularly robust and consistent marker of (also particle-induced) inflammation in vivo, which is a central endpoint predicted by InFiniteLungDT. To predict KE1497 we experimentally measure in vitro 9 KEs for each material separately thus being sensitive to app. 20% of all inflammation-associated KEs, because they are involved in 6 linear pathways (from each of the selected KEs to KE1497) or in 12953 cycles comprising at least one of the monitored KEs, covering 46 KEs.

From these 17 biological observables, InFiniteLungDT measures 10 key events (KEs) that are uniformly distributed within the network of KEs related to inflammation. According to AOPWiki, INFLAMMATION is associated with 250 upstream + 281 downstream KEY EVENTS (KEs), constituting several human-health-relevant AOPs. Among these KEs, neutrophil influx (KE1497 -

Increased, recruitment of inflammatory cells, the main results predicted by the *InFiniteLungDT*) is one of the central ones due to direct coupling with 46 KEs (6 pathways and 12953 cycle loops), reflecting the role of neutrophil influx as a very robust and consistent marker of inflammation observed *in vivo* [59].

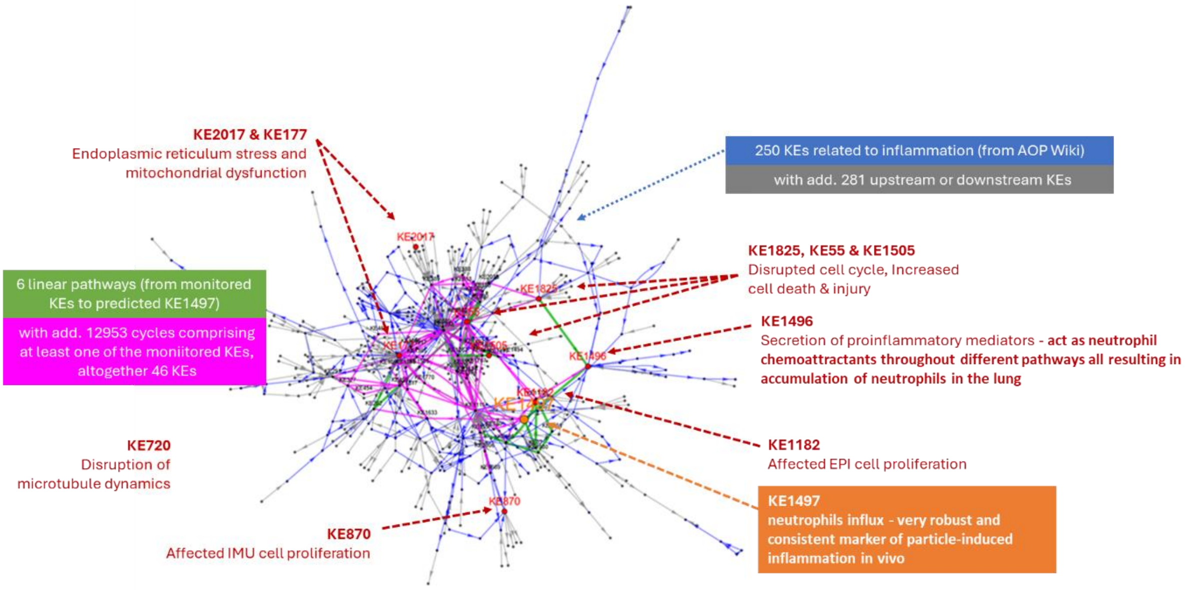

*FigureS 12. The biological responses of cell clusters quantified from in vitro readouts from the InFinite biological platform (in red) are connected within a large network of inflammation-related KEs listed in AOPWiki.*

#### Why does InFiniteLungDT biological platform comprise mouse epithelial and immune cell lines?

We selected cell lines for InFinite biological platform based on their ability to stably reproduce the largest number of inflammation-related key events described in paragraph 0 above. This ensures they are relevant for mouse, rat and human biology.

InFinite biological platform therefore consists of  murine alveolar co-culture of LA-4 epithelial cells and MH-S macrophages (both biosafety level 1), since we have previously shown that this system reproduces lung-inflammation-relevant gene transcription and proinflammatory cytokines secretion [5] and cancer-relevant whole-genome duplications observed in human patients [63]. Also, Alveolar macrophages have been shown to initiate recruitment of neutrophils after nanoparticle inhalation in vivo [60]. Thus, the coculture we selected should express upstream KEs leading to neutrophil influx (KE1497) observed in vivo.  See, SI-9 Baseline phenotype of unexposed LA-4 and MH-S – integrity of inflammatory pathways on page 83 for more detailed description.

#### Why are cell lines in InFinite biological platform grown at sub-confluent conditions?

Cells in cluster react faster and express the largest number of inflammation-related KEs, and provide that in vitro measurements can be performed in a uniform fashion across the whole plate (well-to-well), between plates and across multiple runs by using an automated algorithm for selection of reproducible regions of interest, according to GIVIMP guidelines.

In finite biological platform thus consists of sub-confluent cellular clusters (Submerged mono- and co-cultures are grown at a medium confluency (10-50% for LA-4 lung epithelial cells and 1-10% for MH-S alveolar immune cells - macrophages). This enables monitoring of KEs that are not observable in confluent conditions. Exponential growth is also the most robustly reproducible growth phase of cells in vitro. For example, proliferation of both cell types, cell mitosis, cell type to cell type communication via extracellular vesicle release and uptake, ... are only observable by dynamic (time-dependent) monitoring in InFinite biological platform.

#### Is InFinite biological platform relevant to human health?

InFinite biological platform is optimized to rodent and human relevant prediction of neutrophil influx. Since KEs that are listed in AOPWiki and measured by InFiniteLungDT are relevant both to rodents and humans. The InFinite biological platform expresses largest subgroup of 46 lung-relevant KEs known to drive neutrophils influx (KE1497), that are, according to AOPWiki, relevant for both mice (which is allowed by OECD TGs and for which we have large amount of in vivo data for calibration and validation) and humans (for which the prediction should finally be translatable) [64].

We tested the human relevance of the biological platform in Brown et al., where we documented whole-genome duplication and associated genomic signatures in tumours from a cohort of more than 1,800 patients, and demonstrated that combustion by-products are capable of inducing whole-genome duplications in cell culture, with mutational signatures in patient tumours consistent with exposure to those carcinogens [63]. LA-4 cells under particle exposure reproduce the same phenomenon. This shows correspondence between the cellular response of our murine cell substrate and a phenotype documented in human patients, established at the level of the underlying cellular response rather than reproducing a model that would reproduce human morphology or all the human specific biochemical pathways. InFiniteLungDT is, at least to our knowledge, the only in vitro cell model capable of reproducing this observation in human patinets.

#### Why have we selected multi-channel bi-modal live cell microscopy?

Only multi-channel bi-modal live cell microscopy can capture large number of different time evolutions of biological responses of different cell types in high-throughput manner, and without nanomaterial specific artefacts, which is necessary to propagate the in vitro measured KEs several months to years further in time and evaluate them against in vivo data. This provides cost-effective and timely prediction of long-term in vivo inflammation.

InFiniteLungDT monitoring is performed on a confocal microscope and relies on 4 physical channels: one back-scattered channel (at 488 nm) and 3 fluorescent channels (excitation at laser wavelengths of 488nm, 561nm and 640 nm). This enables simultaneous time-dependent monitoring of 9 inflammation relevant KEs:

- In the back-scattered channel, the material is tracked in label-free way delivering the mask of material for every image;
- In the first fluorescent channel, cell tracker is used to detect interior of epithelial cells;
- In the second fluorescent channel, second cell tracker is used to detect interior of immune cells together with tubulin label that is used to detect tubulin fibers in both type of cells;
- In the third fluorescent channel, membrane probe (cell mask) is used to localize all membrane-based cellular structures.

These readouts are automatically quantified from images acquired by 24-hour live-cell confocal microscopy of exposed/control (mono/co)-cultures via 3 fluorescence channels (monitoring fluorescently labelled cytosols, membranes, and tubulin) and 1 back-scattering channel (nanomaterial - particle tracking).

InFiniteLungDT monitors exclusively biological responses (and no material-related properties) throughout fluorescence and back-scattered modality, whereas some of these effects also include direct material-induced damage of cellular organelles (functional units) or direct material co-localisation with cellular organelles and cell types, and other effects include material-unspecific responses such as cell growth, material-independent responses of cellular organelles, cells (as a whole) and cell-cell communication. Material-dependent responses are monitored through label-free back-scattered modality, which guarantees that these effects are direct consequence of interaction between material and biological structures, while all others do not involve such direct interactions and are thus direct-interaction-free (because there is no colocalized material-signal from back-scattered modality).

By measuring biological responses to nanoforms, InFiniteLungDT minimizes assay interference. As was determined within the EU-funded FP7 NanoTEST project, conventional toxicity tests, designed for soluble chemicals and drugs (e.g. cytotoxicity assays (WST-1, MTT, lactate dehydrogenase, neutral red, propidium iodide, (3)H-thymidine incorporation), pro-inflammatory response evaluation (ELISA for GM-CSF, IL-6, and IL-8), and oxidative stress detection (monoBromoBimane, dichlorofluorescein, and NO assays)), often interfere with nanoforms (enhancement or quenching by the nanoform itself), thus requiring additional controls and substantial care with interpretationtes [65]. We instead measure biological response of the cells, similarly to JRC scientists in the frame of OECD Working Party of Manufactured Nanomaterials (WPMN) who showed that measuring biological response has several advantages over commonly used in vitro cytotoxicity assays, as it avoids test interferences [67].

#### Why is digital twin superior to pattern-matching AI approach?

The digital twin (mathematical model) is constructed by searching for the parameterization that can describe any response dynamics for all (exposure and post-exposure) times for all the mathematically relevant (and different) contexts. Digital Twin is therefore a virtual replica of a physical system/objects based on real-time measured data, ready to simulate time evolution of the system into the future under various scenaria.

This allows in silico prediction of the biological responses outside of the in vitro time window (propagation further in time) allowing that interaction couplings can be determined within the 24-hour in vitro experiment and then propagated up to 2 years further in time to be evaluated against in vivo post exposure observations at later times. This is crucial for chronic diseases where inflammation can persist years after initial nanomaterial exposure.

The most important property of mechanistically created digital twins, once the equations of motion are set up, is that the digital twin can be run from any state and boundary conditions, which in this case means it can be used to describe arbitrary exposure patterns.

Moreover, predictions for longer (subacute, sub-chronic and chronic) time windows are calibrated using responses at different time points from 1 day to 9 months post-exposure and at very different doses of 15 different materials that exploit very different triggering Mode-of-

Action and validated against responses spanning the same time and dose ranges triggered by even larger group of 29 diverse materials. This guarantees that digital twin creation covers very diverse ways of triggering and resolution and can then be mathematically applied to different exposure patterns from single, repeated or continuous, even for the previously untested materials, as well as mixtures of materials.

On the other hand AI methods suffer from overfitting and cannot be used to extrapolate much beyond time windows on which they were trained. In contrast to the pattern-based predictions of traditional machine learning methods, the integration of mechanistic models (open-box) into digital twins also addresses the **challenges of overfitting**, which are prevalent in systems characterized by vast parameter spaces such as machine learning methods, which learn patterns from data without necessarily incorporating underlying biological mechanisms [66]

#### Single versus continuous exposure – why were data from mice instillation experiments instead of rodent inhalation experiments used for the creation of the digital twin?

Single exposure (instillation) experiments enable that in vitro monitoring can be run at a single concentration, under condition that at this concentration all the interaction couplings between biological observables are significant. This cannot be achieved using continuous inhalation experimental data, due to explicit dependencies of interaction terms on dose-dependent and dose-independent interaction couplings. The single-exposure type of animal experiments combined with long post-exposure monitoring period (up to 9 months) can reliably unravel both phases of disease evolution: inflammation build-up and its resolution. On the other hand, repeated exposure effectively superimposes the responses to small, repeated doses counteracting the resolution of the responses due to earlier dose deliveries by the triggering of the new responses due to later dose deliveries. This makes single exposure experiments much more reliable source of data and far more superior to calibrate the creation of digital twins, both in terms of speed and mechanisms involved, covering local early Mode-of-Action as well as systemic responses.

Single exposure (instillation) experiments also allow more accurate determination of the local nanomaterial dose, which triggers the biological response of the cells and has to be matched in in vitro experiments.

#### Why were the 44 benchmark nanomaterials selected for the creation of the InFiniteLung digital twin?

We have used 44 different benchmark materials, from diverse material families, triggering orthogonal Mode-of-Action, since the largest database for single intratracheal mice exposure exists for them, and all the materials are commercially available or are held at JRC and NIST repository at larger quantities.

We have used the same materials (nanoforms from the same batch were distributed to all the partners), including standardized materials from JRC material repository, NIST standards or collected by NRCWE group in Denmark. For all the new materials, both in vivo and in vitro data has been also acquired for the same materials in recent EU projects (SmartNanoTox and NanoPASS). Thus, the validation does guarantee that the prediction is reliable.

The in vivo data were taken from the selected mice experiments because the latter were focused on post-exposure monitoring that is crucial for calibrating resolution dynamics (and not only triggering / building up the inflammatory responses). Altogether, we reused more than 3094 mice exposure data for creation of the InFiniteLungDT (mostly collected from various EU projects from the last 3 decades) – no comparably big alternative database of responses is available to our knowledge.

#### Reliability of the prediction regarding the nanomaterial assay interference

InFiniteLungDT method measures biological effects of nanoforms to avoid assay interference. Conventional toxicity tests interfere with nanoforms (enhancement or quenching by the nanoform itself), which precludes direct application of the tests to nanoforms in the same manner as for soluble chemicals and drugs, as was determined within the EU-funded FP7 NanoTEST project. Therefore we avoided the problematic in vitro cytotoxicity assays (WST-1, MTT, lactate dehydrogenase, neutral red, propidium iodide, (3)H-thymidine incorporation), pro-inflammatory response evaluation (ELISA for GM-CSF, IL-6, and IL-8), and oxidative stress detection (monoBromoBimane, dichlorofluorescein, and NO assays) [65]. We instead measure biological response of the cells, similarly to JRC scientists in the frame of OECD Working Party of Manufactured Nanomaterials (WPMN) who showed that measuring biological response has several advantages over commonly used in vitro cytotoxicity assays, as it avoids test interferences [67].

Therefore, InFiniteLungDT monitors exclusively biological responses (and no material-related properties) throughout fluorescence and back-scattered modality, whereas some of these effects also include direct material-induced damage of cellular organelles (functional units) or direct material co-localisation with cellular organelles and cell types, and other effects include material-unspecific responses such as cell growth, material-independent responses of cellular organelles, cells (as a whole) and cell-cell communication. Material-dependent responses are monitored through label-free back-scattered modality, which guarantees that these effects are direct consequence of interaction between material and biological structures, while all others do not involve such direct interactions and are thus direct-interaction-free (because there is no colocalized material-signal from back-scattered modality).

#### SI-15 In vitro measurements and time dependent quantifications of biological observables for each nanomaterial used in calibration and validation of InFiniteLungDT

For more detailed comparison of *in vitro* measured readouts by In Finite platform and quantification of biological observables elaborated for each material of this study, see the report of quantitative data of KE dynamics, extracted from images of time-lapse microscopy over 1–2 days, for benchmark and industrial materials on in vitro models for selected adverse outcomes.

For the purposes of providing time-lapse microscopy data, that is organized and documented according to FAIR principles, and formatted as needed for modelling, we are delivering the data repository, consisting of example microscopy images for each material (three time-points, all channels, all conditions, one biological replicate).

The full repository data is available to be accessed on the platform Zenodo, at the following link: https://doi.org/10.5281/zenodo.17696724
