## Supplementary figures and images for "InFiniteLungDT, an in-vitro-learned digital twin for the prediction of material-triggered chronic neutrophilic lung inflammation: mechanism-anchored pre-validation across materials, species and exposure scenarios"

### inhalation_dose-dependent_HiRes_plots.png

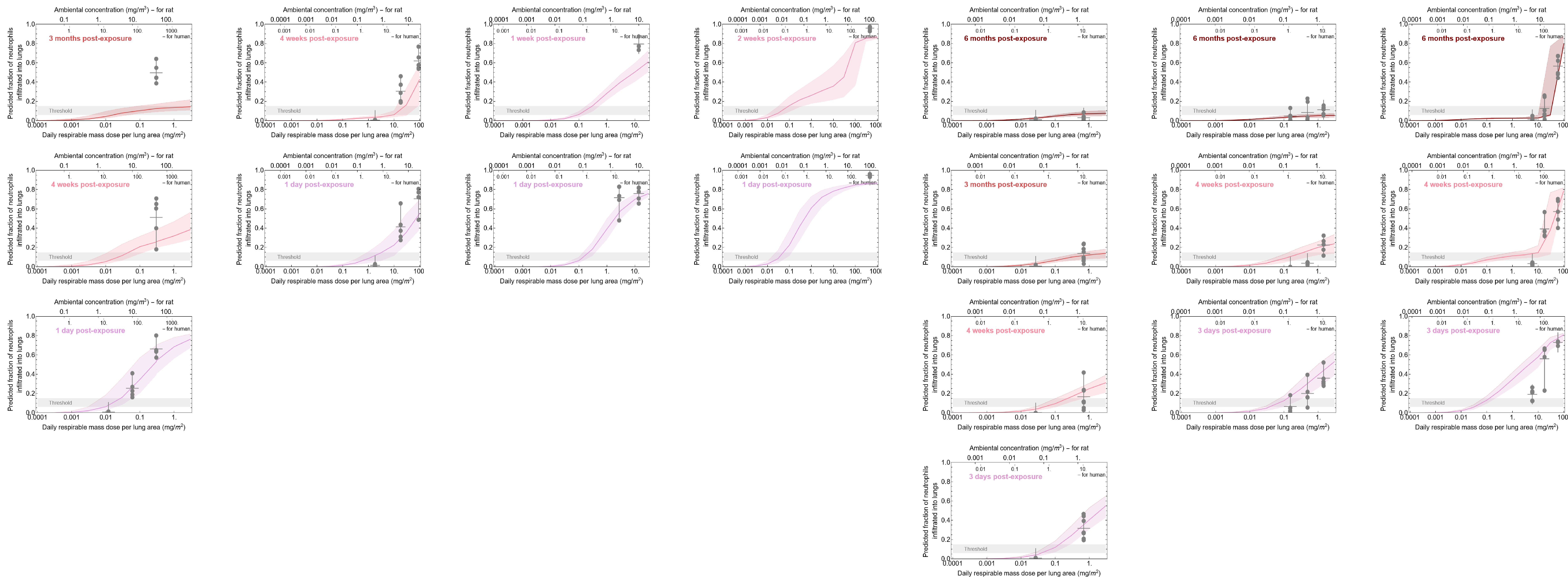

### inhalation_time-dependent_HiRes_plots.png

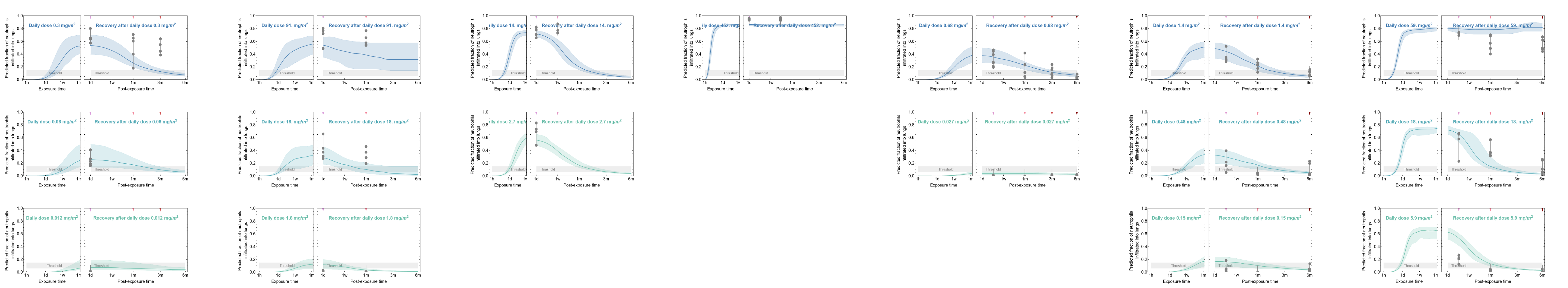

### instilation_dose-dependent_HiRes_plots.png

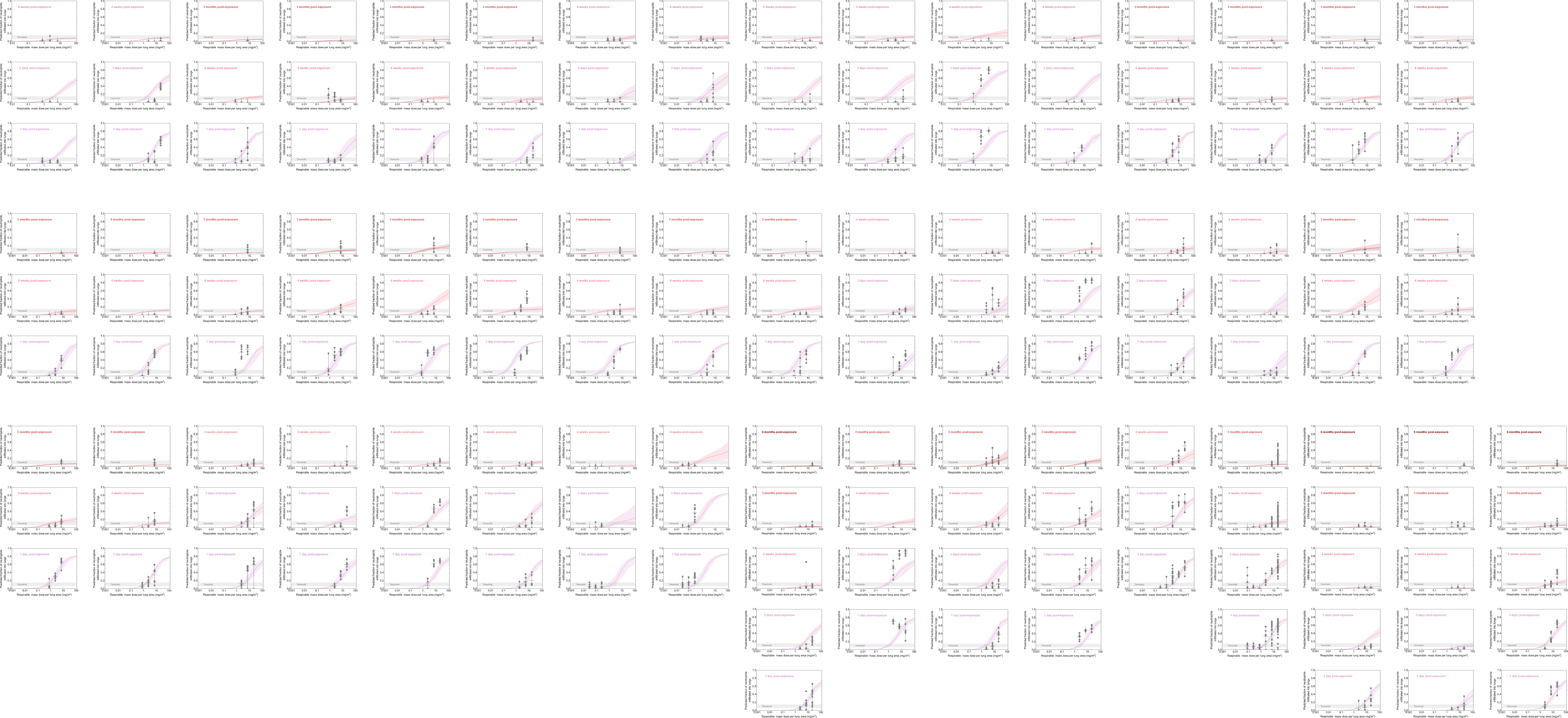

### instilation_time-dependent_HiRes_plots.png

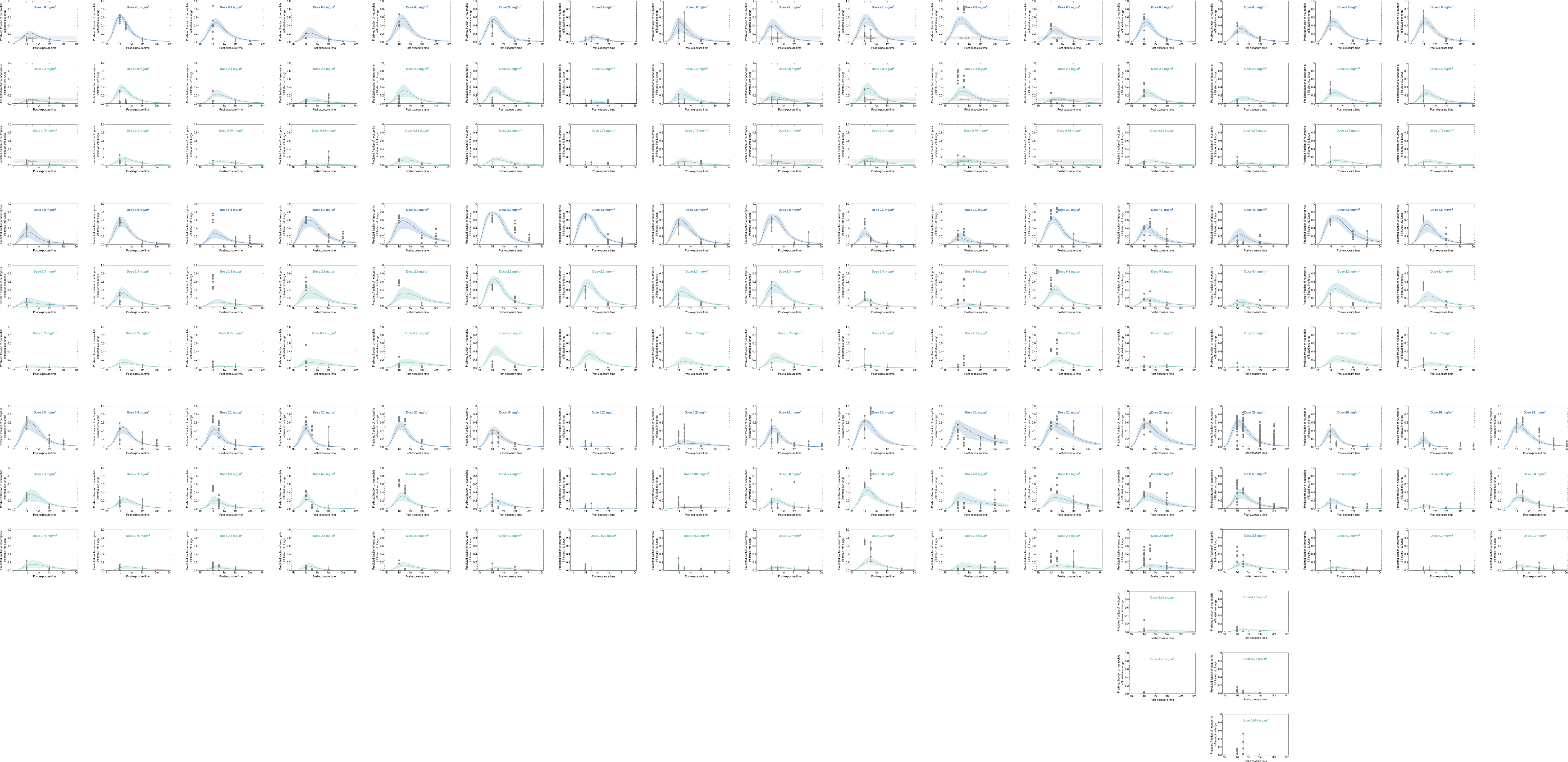
